# Enhancing sensitivity and controlling false discovery rate in somatic indel discovery

**DOI:** 10.1101/741256

**Authors:** Johannes Köster, Louis J. Dijkstra, Tobias Marschall, Alexander Schönhuth

## Abstract

As witnessed by various population-scale cancer genome sequencing projects, accurate discovery of somatic variants has become of central importance in modern cancer research. However, count statistics on somatic insertions and deletions (indels) discovered so far point out that large amounts of discoveries must have been missed. The reason is that the combination of uncertainties relating to, for example, gap and alignment ambiguities, twilight zone indels, cancer heterogeneity, sample purity, sampling and strand bias are hard to accurately quantify. Here, a unifying statistical model is provided whose dependency structures enable to accurately quantify all inherent uncertainties in short time. As major consequence, false discovery rate (FDR) in somatic indel discovery can now be controlled at utmost accuracy. As demonstrated on simulated and real data, this enables to dramatically increase the amount of true discoveries while safely suppressing the FDR. Specifically supported by workflow design, our approach can be integrated as a post-processing step in large-scale projects.

The software is publicly available at https://varlociraptor.github.io and can be easily installed via Bioconda^1^ [Grüning et al., 2018].

## 1 Introduction

Cancer is a genetic disorder in the first place; somatic mutations turn originally healthy cells into a heterogeneous mix of aberrantly evolving cell clones [Burrell et al., 2013]. Global consortia have launched population-scale sequencing projects concerned with the discovery and annotation of somatic variants in cancer genomes [The International Cancer Genome Consortium, 2010, Weinstein et al., 2013]. Potential benefits of the systematic analysis of somatic mutations include improved diagnosis, staging and therapy protocol selection in the clinic.

The fraction of somatic variants discovered, however, has left room for improvement across the whole range of possible variant types [Alioto et al., 2015]. Thereby, somatic insertions and deletions (indels)^2^ have proven to pose particular challenges when belonging to certain classes or length ranges [Hause et al., 2016, Maruvka et al., 2017]. Indels of length approximately 30 - 250 bp, termed “the next-generation sequencing (NGS) twilight zone of indels” in other contexts [Mandoiu and Zelikovsky, 2016, Marschall et al., 2013, Trappe et al., 2014], have resisted their discovery in particular also in somatic variant calling: while the COSMIC database^3^ counts 1,879,044 indels of length 1-30 bp, it only counts 17 793 indels of length 31-60 bp, 3758 indels of length 61-100 bp and 2483 indels of length 101-250 bp. The drop by two orders of magnitude from 1-30 to 30-60 bp, not followed by any such drops in further length bins, has so far not been supported by any reasonable biological interpretation. Our benchmark experiments demonstrate that this size range still corresponds to a blind spot in somatic mutation discovery. The most likely explanation is that the majority of somatic indels in that size range have remained undiscovered so far.

Therefore, the application of more sensitive^4^ somatic indel calling strategies most likely will induce striking changes in the spectrum of somatic indels so far detected. As indicated in earlier work [Maruvka et al., 2017], this has the potential to deepen our understanding of the origin and effects of somatic indels, beyond just balancing count statistics.

In this paper, we suggest such a sensitive strategy, and prove that with it we can make substantial progress in terms of eliminating the somatic indel discovery blind spot. To understand the issues that are characteristic of this blind spot, consider that somatic variant discovery (unlike germline variant discovery) is a two-step procedure: in a first step, one discovers putative variants in both the cancer and healthy (or control) genome of the individual analyzed. In the second step—which is unique to somatic variant discovery— one runs a differential analysis that classifies putative variants into somatic, because they only appear in the cancer genome, germline and healthy somatic, because variants appear in the control genome (where healthy somatic appear at subclonal levels), or just noise.

Already the first step (which also applies for generic germline variant discovery) is affected by major issues, where for example gap wander and annihilation (see [Lunter et al., 2008]) are well-known and notorious examples when determining gapped alignments in general. Issues become further aggravated when dealing with indels of 30-250 bp due to particularities of NGS read alignment and indel discovery tools. So, for determining twilight zone indels, one needs to make particular methodical efforts already when dealing with generic settings [Mandoiu and Zelikovsky, 2016, Marschall et al., 2012, 2013, Trappe et al., 2014].

In somatic indel discovery, we deal with an additional layer of issues due to cancer heterogeneity. The variant allele frequency (VAF), here the fraction of genome copies in the (tumor or control) sample affected by the variant, is either 0.0, 0.5, or 1.0 for germline variants, reflecting absence, hetero- or homozygosity. In contrast, allele frequencies of somatic variants vary across the whole range from 0.0 to 1.0, depending on the clonal structure of the tumor sample and its impurity (the ratio of healthy genome copies in the tumor sample). Usually, there is no prior information about the clonal structure available at the time of variant calling. Low-frequency variants (i.e. having a VAF close to zero) yield particularly weak, statistically uncertain signals. Of course, there are limits to somatic variant discovery, relative to VAF and sequencing depth. The methodical challenge is to not miss any discoveries that a statistically sound approach is able to reveal. The purpose of this paper is to do this: we would like to fathom the corresponding limits theoretically and to push them considerably in practice.

Certainly, there are still also improvements for the first step conceivable; however, the second (differential analysis) step has never been treated before with statistical rigor. So, the second step may have left room for improvements in particular. We recall that dealing with the various statistical uncertainties due to (as above-mentioned) cancer heterogeneity, gap placement, strand bias, etc., is the major challenge in the second step. For successful operation, one needs to accurately quantify all relevant uncertainties in the first place. As a consequence, one has a sound statistical account on whether putative variants are somatic, germline, healthy somatic, or just errors.

Key to success in increasing the number of true somatic variant discoveries is to establish statistically sound *false discovery rate (FDR)* control, as the canonical procedure to limit the number of false predictions when increasing the output. In an ideal setting, the user specifies the maximal ratio of false discoveries s/he is willing to deal with. Subsequently, a maximal set of discoveries is reported that ensures the FDR specified by the user. It is a well-known insight that accurate FDR control only works if uncertainties have been accurately quantified: otherwise inaccuracies propagate towards FDR control, implying insufficient control, too little discoveries, or both. This explains why working with accurately quantified uncertainties is imperative when seeking to safely increase the amount of true discoveries.

The desired accuracy in uncertainty quantification, however, comes at a cost: if data is uncertain, one deals with an exponential amount of possibly true data scenarios. This prevents naive approaches to work at the desired level of accuracy without serious runtime issues, which constitutes a common computational bottleneck in uncertainty quantification. In our setting, we will be dealing with at least 3^*n*^ possible scenarios for one putative indel where *n* corresponds to the sum of read coverages in the cancer and control genome at a particular locus. Nowadays, *n* ≤ 60 is not uncommon. This number of arithmetic operations is prohibitive when processing up to hundreds of thousands putative indel loci.

In the model presented in this paper, we overcome this bottleneck by presenting a Bayesian latent variable model whose conditional dependencies point out how to compute all of the relevant probabilities in time *linear, and not exponential* in the coverage of a putative indel locus. Thanks to its computational efficiency, the model captures and accurately quantifies all uncertainties involved in somatic indel discovery in a comprehensive manner.

In summary, we are able to compute all probabilities required for enforcing reliable FDR control at both the necessary speed and accuracy. Reliable, sound and accurate FDR control, in turn, gets us in position to substantially increase the recall in somatic indel discovery, without having to deal with losses or even improving in terms of precision. As was to be expected, improvements in the twilight zone turn out to be most dramatic: in comparison with state-of-the-art tools we double or even triple recall, while preserving (often better than just) operable precision.

## 2 Results

We have designed *Varlociraptor*, as a method to implement the improvements in the differential analysis step outlined in the Introduction. To the best of our knowledge, Varlociraptor is the first method that allows for accurate and statistically sound false discovery rate control in the discovery of somatic indels. As a consequence, the application of Varlociraptor leads to substantial increases in recall in somatic indel discovery. Varlociraptor doubles or even triples the number of true discoveries in comparison with state-of-the-art tools, while often also further improving on precision, or, at any rate, not incurring any kind of loss in precision. Varlociraptor also accurately estimates the variant allele frequency (VAF) for all somatic indels.

In the following, we provide a high-level description of the Varlociraptor workflow. We provide a brief explanation of how one can quantify all relevant uncertainties in linear runtime, as the key methodical breakthrough and the fundamental building block that underlies all that follows. We briefly illustrate how the Bayesian latent variable model that enables rapid quantification of uncertainties further immediately gives rise to the computation of all probabilities that are crucial in somatic variant calling. We further briefly address how Varlociraptor estimates variant allele frequencies (VAF’s). Finally, we explain how accurate FDR can be established. For details, we refer to the Methods section.

Subsequently, we analyze Varlociraptor’s performance in comparison with current state-of-the-art tools on simulated and real data. As pointed out above, we show that Varlociraptor indeed achieves (sometimes drastic) increases in recall, often accompanied by further increases in precision. We notice that the probabilities used for classifying putative variant calls allow for a clear distinction between true and false positives, which is of considerable value in classification practice. We then demonstrate that Varlociraptor indeed reliably controls FDR, thereby also providing the theoretical explanation for why Varlociraptor achieves superior performance rates in terms of recall and precision. Varlociraptor further accurately estimates all VAF’s. Turning our attention to real data, we conclude that Varlociraptor achieves superior concordance for variants of VAF at least 20%. For variants of VAF of less than 20%, Varlociraptor is the only tool that discovers considerable amounts of variants. The low coverage of reads supporting such calls delivers stringent statistical explanations for why concordance cannot be reached at rates that apply for calls above 20%. To corroborate that the majority of Varlociraptor’s calls are correct—just as we experienced on simulated data—we demonstrate that Varlociraptor’s count statistics agree with the theoretical expectation under neutral evolution.

### 2.1 Workflow

We first discuss how Varlociraptor embeds in a workflow for somatic variant calling and highlight the central difference to classical approaches.

The classical workflow for calling somatic variants (Figure 1a) starts with aligned reads from tumor and corresponding healthy sample of the same patient in BAM^5^ or CRAM^6^ format. First, variants are discovered and a differential analysis is performed to call variants as somatic or germline. Candidate variants are reported in VCF or BCF format^7^. In the following we will refer to VCF as a placeholder for VCF/BCF, and BAM as a placeholder for BAM/CRAM. Second, the candidate variants are filtered, usually by applying thresholds for various scores (e.g. variant quality, strand bias, coverage, minimum mapping quality, minimum number of supporting reads in healthy sample), in order to obtain final variant calls. If not just relying on some suggested defaults, finding those thresholds is often a tedious, study specific effort.

**Figure 1:**
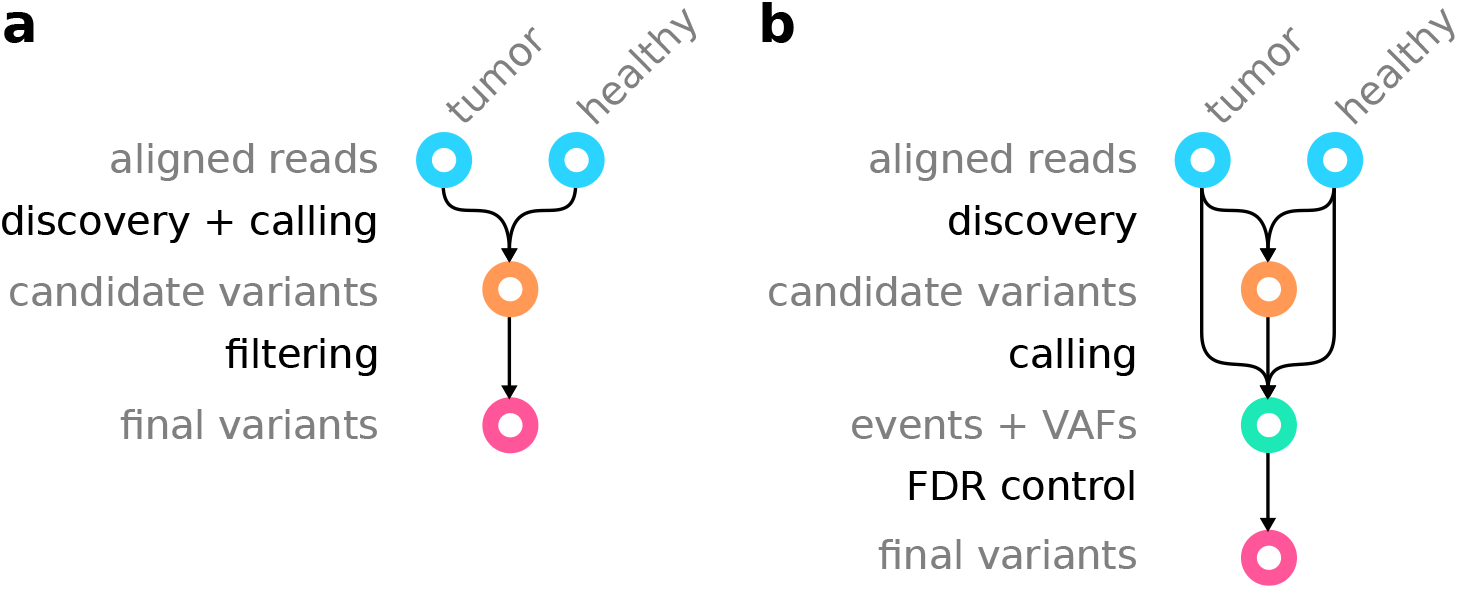
Difference between (a) the classic somatic variant calling workflow and (b) the Varlociraptor approach.

With Varlociraptor, we provide a new approach for calling and filtering, thereby separating variant discovery from calling (Figure 1b). The input for Varlociraptor are candidate variants from an external discovery step. Here, any variant calling tool can be applied.^8^ There are plenty of approaches that thoroughly deal with the discovery step. In particular, as it is already common practice within state of the art somatic variant calling pipelines (e.g. the Sarek pipeline^9^), it is possible to combine the candidate variants of different callers in order to obtain maximum sensitivity across all variant types and length ranges. However, instead of having to perform ad-hoc merging of finalized calls (e.g. Sarek^9^ performs majority voting), Varlociraptor provides a unified mechanism for assessing all candidate variants. During calling, Varlociraptor classifies variants into somatic tumor, somatic healthy, germline or absent variants, while providing posterior probabilities for each event (see section 2.2.2) along with maximum a posteriori estimates of the variant allele frequency (VAF) (see section 2.2.3), reported in BCF format. Finally, using the posterior probabilities, Varlociraptor can filter variants by simply controlling for a desired false discovery rate (FDR, see section 2.2.4), instead of requiring the adjustment of various thresholds. This becomes possible because Varlociraptor, as the first approach, integrates all known sources of uncertainty into a single, unified model.

In this work, we will evaluate Varlociraptor’s performance in direct comparison with the (usually ad-hoc) differential analysis routines provided by other tools. We will use the indels that are output by the competitor as per its first (discovery) step as input for Varlociraptor. This way we ensure that all tools receive input they are supposed to deal with, by their design, and therefore ensure maximum fairness.

### 2.2 Foundation of the Approach

#### 2.2.1 Efficient Computation of the Fundamental Likelihood Function

Let us fix a particular variant locus, as given by an entry in the VCF file that lists all candidate variants from Figure 1b. By *θ_h_* and *θ_c_* we denote the true, but unknown allele frequency of that (putative) variant among the healthy (*θ_h_*) and the cancerous (*θ_c_*) genome copies. While for *germline* variants *θ* ∈ {0,1/2,1} reflecting absence (artifacts or noise), hetero- and homozygosity of the variant, *θ* ∈ [0,1] for somatic variants. We model that *somatic healthy* variants usually show at subclonal rates by allowing only *θ* ∈ (0,1/2), i.e. the exclusive interval between 0 and 1/2. A variant evaluates as *somatic tumor* if and only if *θ_c_* > 0, while *θ_h_* = 0. Since we are most interested in these variants, one of the central goals is to conclude that *θ_c_* > 0, *θ_h_* = 0 for a particular putative variant with sufficiently large probability.

By 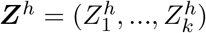 and 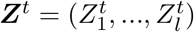, we denote the read data being associated with the variant locus in the healthy (h) and the tumor (t) sample^10^. Each of the 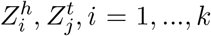, *j* = 1,…, *l* represents one (paired-end) read that became aligned across or nearby the given variant locus. This further means that *k* and *l* correspond to the sample-specific read coverages at that locus. For selecting reads via alignments, we use BWA-Mem [Li and Homer, 2010] in the following, although the choice of particular aligner is optional, as long as the aligner outputs a MAPQ value [Li et al., 2008b], which quantifies the certainty by which the sequenced fragment (represented by the read pair) stems from the locus under consideration.

Let further 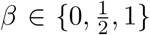 denote the *strand bias* affecting the particular variant locus. Thereby *β* = 0 and *β* = 1 denote that evidence about the putative variant occurs only in the reverse (*β* = 0) or in the forward (*β* = 1) strand. Both cases are indicators of sequencing or mapping artifacts. Therefore, no strand bias, i.e. *β* = 1/2 will subsequently be used to select for non-artifact variants.

We present a *Bayesian latent variable model* that enables to *efficiently compute*

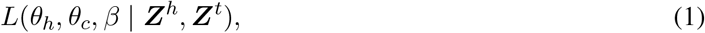

the likelihood of allele frequencies *θ_h_, θ_c_* and strand bias *β* given read data ***Z***^*h*^, ***Z***^*t*^ (see section 5.3). Straightforward approaches to computing (1) via *fully Bayesian inverse uncertainty quantification* [Liu et al., 2009], which is the canonical and approved way to computing (1) fail due to requiring exponential runtime. The conditional dependency structure of the statistical model we raise (see Methods, section 5.3), points out a way to compute (1) in runtime linear in *k* + *l*, as summarized by the following theorem.

##### Theorem 2.1.

*L*(*θ_h_, θ_c_, β* | ***Z***^*h*^, ***Z***^*t*^) *can be computed in O*(*k* + *l*) *arithmetic operations*.

Note that this is the best one can hope for; the insight is crucial in somatic variant calling practice, beyond establishing also a theoretical novelty, because Theorem 2.1 establishes that *L*(*θ_h_, θ_c_, β* | ***Z***^*h*^, ***Z***^*t*^) can be evaluated in short runtime at any (*θ_h_, θ_c_, β*). This in turn renders integration over *L*(*θ_h_, θ_c_, β* | ***Z***^*h*^, ***Z***^*t*^) computationally feasible, which facilitates solving the following three essential problems.

#### 2.2.2 Classification

Statistically sound classification in somatic variant calling requires, when given ***Z***^*h*^, ***Z***^*t*^, to compute the posterior probabilities for the following four cases, which refer to different combinations of *θ_c_, θ_h_* (see also Figure 8a in Methods).

- somatic tumor (st); *θ_c_* > 0, *θ_h_* = 0: the variant is somatic in the tumor, and does not appear in the healthy genome,
- germline (ge); *θ_h_* ∈ {1/2, 1}: a germline variant, where *θ_h_* = 1/2 reflects a heterozygous and *θ_h_* = 1 a homozygous variant,
- somatic healthy (sh); *θ_h_* ∈ (0, 1/2): a variant that is somatic but appears in the healthy genome, reflected by subclonal, non-germline variant allele frequencies, or
- absent (ab); *β* ∈ {0,1} or *θ_c_* = 0, *θ_h_* = 0: the variant reflects (strand bias) artifacts or noise

Note in particular that cases (st), (ge), (sh) imply that there is no strand bias, i.e. *β* = 1/2. Given the respective read data ***Z***^*h*^, ***Z***^*t*^ from the healthy and the tumor genome, the corresponding posterior probabilities compute as

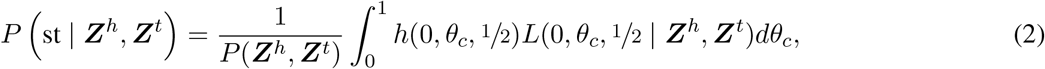

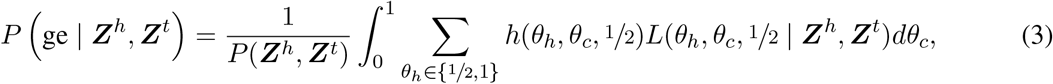

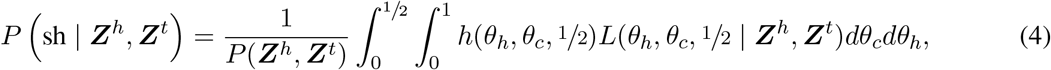

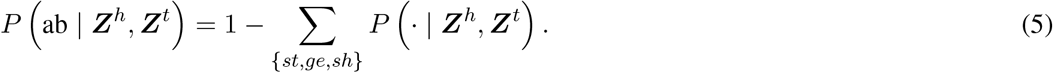

*h*(*θ_h_, θ_c_, β*) is the prior distribution of the three readouts *θ_h_, θ_c_, β*, which can be used to integrate prior knowledge about clonal structure or zygosity rates, if available. We consider the choice of *h* as an open question, that is most important for sparse data (i.e. very low coverage) It can be guided by the work of Williams et al. [2016], for example. For the evaluation conducted here, we use a uniform *h*. *P*(***Z***^*h*^, ***Z***^*t*^) is the marginal probability of the data, which acts as a normalization factor.

The integrals lack an analytic formula, but, supported by the efficient computation of (1), as established by theorem 2.1, can be approximated numerically using quadrature.

#### 2.2.3 Estimating allele frequencies for somatic tumor variants

Upon having determined that a variant is somatic tumor, implying *θ_c_* > 0, *θ_h_* = 0 and 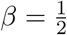, we would like to determine maximum a posteriori estimates for *θ_c_*, given the fragment data ***Z***^*h*^, ***Z***^*t*^. When using a uniform prior this is the same as computing the maximum likelihood estimate

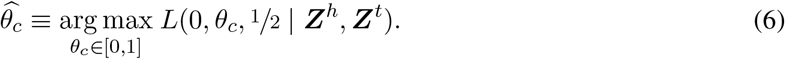

The likelihood function (1) is a higher-order polynomial in *θ_h_* and *θ_c_* for given *β*, as follows from the computations in section 5.3.4, which makes it infeasible to derive its maximum analytically. We can nevertheless prove the following theorem.

##### Theorem 2.2.

*For fixed θ_h_ and β, the logarithm of the likelihood function θ_c_* → *L*(*θ_h_, θ_c_, β* | ***Z***^*h*^, ***Z***^*t*^) *is concave on the unit interval* 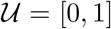. *Hence the likelihood function attains a unique global maximum* 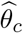 *on* [0,1].

This allows to report the corresponding global maximum 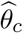 for *θ_h_* = 0 and 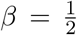, as a reasonable estimate for the VAF of a somatic cancer variant. See Theorem A.1 in Appendix A for a detailed technical exposition, including a proof and a list of extra conditions for this theorem necessary to hold, all of which apply in practice. Since the logarithm of the likelihood function is strictly concave, its maximum can be easily determined numerically.

#### 2.2.4 False Discovery Rate (FDR) Control

Once the event probabilities defined in section 2.2.2 are available, FDR can be controlled in a statistically sound way. In an ideal setting, the user specifies an FDR threshold *γ*, upon which a maximal amount of variants is output such that the ratio of false discoveries among the discoveries overall does not exceed *γ*. Only if the underlying model is statistically sound, maximal increases in terms of true discoveries among the output (sensitivity or recall) can be expected. Note that ad-hoc style FDR control procedures such as ‘call merging’ (raising only discoveries simultaneously supported by several variant callers), while indeed controlling FDR, usually incur serious losses in terms of discoveries, and tend to lead to discovery blind spots, thereby (while not being the main issue) contributing to the lack of ‘somatic twilight zone indels’ so far discovered. Moreover, it becomes next to impossible to fine-tune an analysis to a particular FDR acceptable in the specific context. First tries on FDR control procedures for variant calling have been reported before [DePristo et al., 2011, Marschall et al., 2012]. However, in this work, we present the first fully Bayesian approach, and also the first for the calling of somatic variants.

For a given set of putative somatic tumor variants *C*, each of which is annotated with *p_i_, i* = 1,…, |*C*|, the posterior probability (2) that variant *i* is indeed somatic tumor, we can calculate the expected FDR [Mueller et al., 2004] as

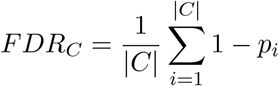

In order to both control FDR at *γ* and raise maximal output, one searches for the largest set of variants *C*\* ⊆ *C* such that *FDR_C_\** ≤ *γ*. One can efficiently implement this by sorting variants by 1 − *p_i_* in ascending order, and summing up 1 − *p_i_* in that order until this sum divided by the number of summands collected, has reached the threshold *γ*.

### 2.3 Data analysis reproducibility

The evaluation performed in this paper is available as a reproducible Snakemake [Köster and Rahmann, 2012] workflow archive^11^. In addition we provide a *Snakemake report* that allows to interactively explore all Figures shown in this article in the context of the workflow, the parameters, and the code used to generate them (see supplementary files).

### 2.4 Data

A central challenge when evaluating somatic indel calling is to sufficiently cover all relevant length ranges. Furthermore, the occurrence of already discovered true twilight zone indels is biased towards easily accessible, hence lesser uncertain regions of the human genome. However, somatic variant databases contain only few twilight zone indels (see section 1). Hence, we chose a dual approach based on designing a simulated dataset with the desired properties for assessing precision and recall, complemented by a concordance analysis on real data.

#### Simulated Data

We used a real genome (Venter’s genome [Levy et al., 2007]), previously approved for NGS benchmarking purposes [Marschall et al., 2012, 2013] as the control genome.

Our goal was to simulate a cancer genome that qualifies for statistically sound benchmarking. We therefore sampled 300 000 somatic point mutations, 150 000 insertions and 150 000 deletions, of which 279 509,139 491 and 139 532 in the autosomes, following the clonal structure described by Figure 2, so as to reflect a scenario that is realistic in terms of tumor evolution. To account for realistic proportions in terms of length, indels follow the length distribution of Venter’s germline indels (which roughly follows apower-law distribution). Allele frequencies range from 0.042 to 0.667 for the somatic indels. The majority of indels is of relatively low frequency and they were randomly placed across the genome. Both low frequencies (making indels hard to distinguish from noise) and random placement (rendering the calling particularly difficult in repetetive regions), lead to a particularly challenging benchmarking dataset.

**Figure 2:**
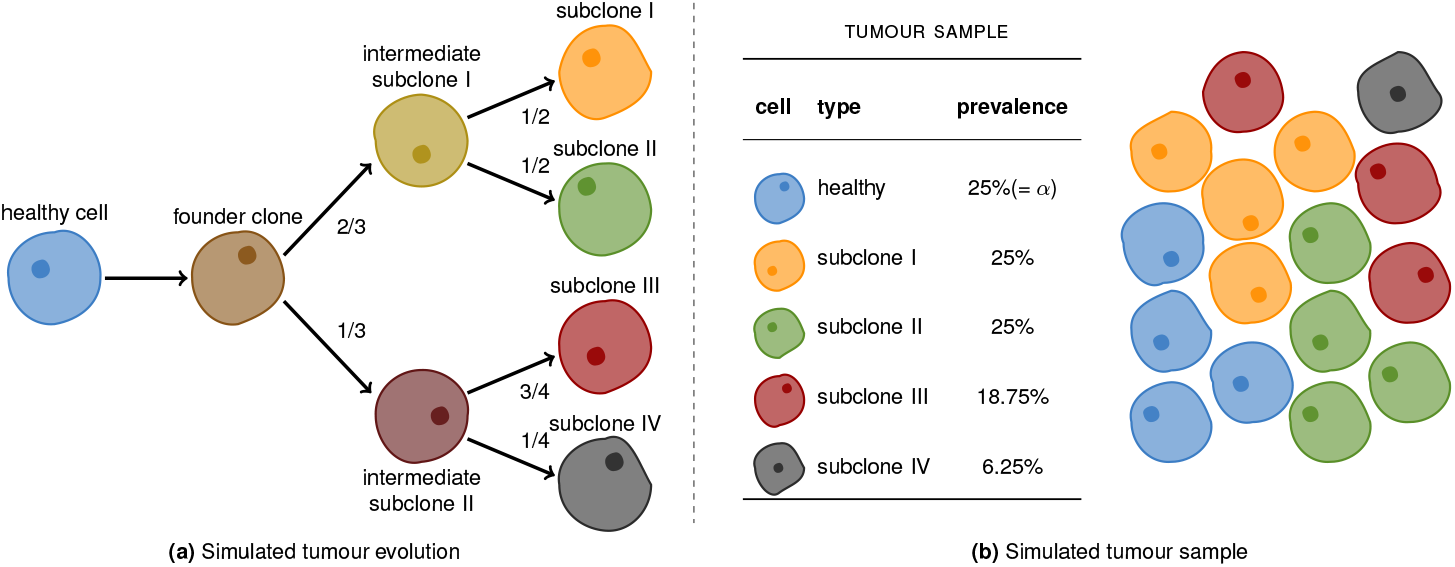
Simulated Cancer Clones: **(a)** The evolution of the cancer clones. **(b)** The simulated tumour sample. Each cell is assumed to be diploid. Cells of the same type share the same genetic code. The relative prevalences of the various cell types are shown in the table. The level of impurity (*α*) is 25%.

Reads were sampled using the Assemblathon read simulator SimSeq [Earl et al., 2011], at 30x and 40x for control and cancer genome, respectively. Subsequently, reads were aligned using BWA-MEM [Li and Durbin, 2009].

#### Real Data

To demonstrate the applicability in practice, we also evaluated real cancer-control genome pairs. Since, as of today, there are no real datasets with known ground truth available (where the lack of real twilight zone indel discoveries is of course an important factor) we opt for a concordance analysis, as a procedure that has been approved earlier. Namely, we analyzed the concordance of reported variants with respect to four replicates of cancer-control genome pairs, all of which have been sampled from the same tumor cell line (Melanoma cell line COLO829), albeit in different institutions [Craig et al., 2016]^12^. If discoveries referring to the four replicates agree to a sensible degree (considering that differences due to batch effects and independent progression are conceivable), one can conclude that performance is also of high quality on real datasets.

### 2.5 Tools

For generating lists of candidate indels in form of VCF files (see Figure 1), we chose Delly 0.7.7 [Rausch et al., 2012], Lancet 1.0.0 [Narzisi], Strelka 2.8.4 [Saunders et al., 2012], Manta 1.3.0 [Chen et al., 2016], and BPI 1.5^13^ without applying additional filtering. All these callers provide their own “ad-hoc” method of annotating somatic variants from tumor/normal sample pairs, which we applied for comparison. We provided BWA-MEM alignments with marked duplicates (via Picard-Tools^14^) as input for all tools. When subsequently running Varlociraptor, we used the output VCF files of the tools in combination with the BAM files that were the basis for generating the candidate variants.

### 2.6 Experiments

We first consider the simulated dataset (see section 2.4). Here, the true somatic variants are known from the simulation procedure. To classify predicted variants of the different callers into true and false positives, we matched them against the known truth using the `vcf–match` subcommand of RBT^15^ with parameters `--max-dist 50 --max-len-diff 50`. This means that we consider a predicted somatic variant to be a true positive if it’s position and length are within 50 bases of the true variant (which reflects approved evaluation practice, see [Wittler et al., 2015] for further reasoning). Thereby the position for a deletion is defined as its center point, i.e., ⌊(*e* − *s*)/2⌋ with *e* being the end position and *s* being the start position of the deletion. In the following we show the results for deletions. If results for insertions essentially deviate from the deletion results, we mention it in the corresponding section of the text. All results for insertions can be found in the supplement.

#### Varlociraptor achieves substantial increases in Recall without loosing Precision

In the following, *Recall* is defined to be the ratio of true variants that became predicted, while *Precision* is the ratio of correct predictions among the predictions overall. Figures 3 and S1 show Recall and Precision for all tools considered on the simulated data. Thereby, we juxtapose the tools’ Recall and Precision when run in standalone modus (dots or dotted lines) with Recall and Precision when postprocessing the respective output sheets of the tools with Varlociraptor (lines). The lines referring to Varlociraptor result from varying the posterior probability threshold: the greater the threshold the smaller the Recall. While the reduction in Recall is merely a consequence of reducing the output, a simultaneous increase in precision only shows if posterior probabilities make sense, as (first) essential evidence of the quality of the approach. Note that the only caller that offers to vary an output-specific threshold is Lancet (dotted orange line in Figures 3 and S1), by scoring variants with p-values^16^.

**Figure 3:**
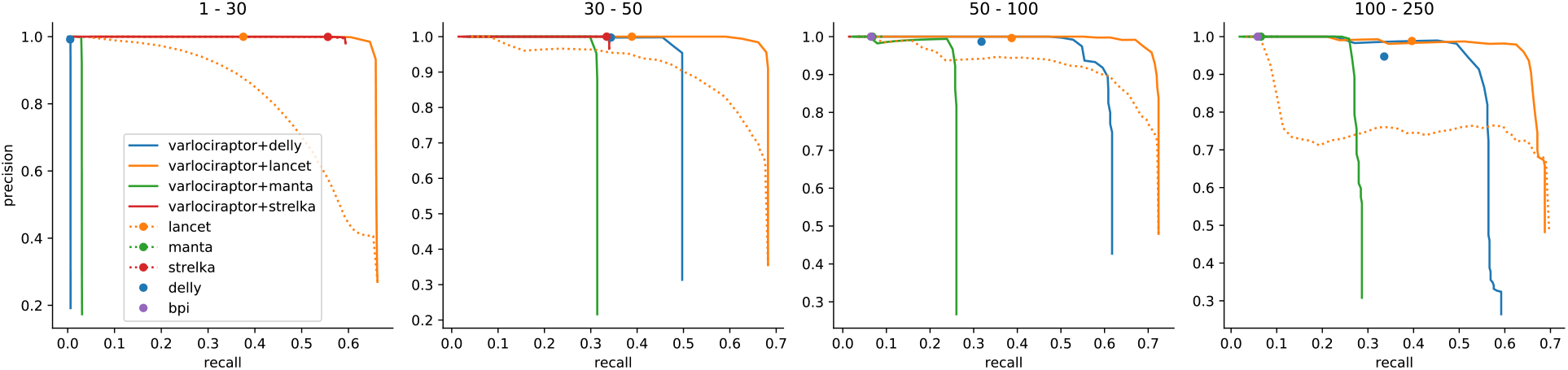
Recall and precision for calling somatic deletions. Results are grouped by deletion length, denoted as interval at the top of the plot. For our approach (Varlociraptor+*) curves are plotted by scanning over the posterior probability for having a somatic tumor variant. For other callers that provide a score to scan over (p-value for Lancet) we plot a dotted line. Ad-hoc results are shown as single dots. Results are shown if the prediction of the caller did provide at least 10 calls. The sharp curves for our approach reflect the favorable property of having a strong separation between the probabilities of true and false positives, see Figure 4.

The first, fundamental observation is that Precision indeed increases on increasing the posterior probability cutoff, across all size ranges and inputs. This points out that Varlociraptor’s posterior probabilities for a variant to be somatic indeed make sense.

When further comparing Varlociraptor’s (continuous) lines with the dots or dotted lines referring to the standalone modus of the other tools, it becomes immediately evident that Varlociraptor improves on the tools’ results: in all cases Varlociraptor’s lines are upper right of the tools’ dots or dotted lines. This means that Varlociraptor achieves better combinations of Recall and Precision in all cases. The most striking observation however is that Varlociraptor achieves substantial increases in recall in comparison to the standalone tools: in particular in the twilight zone (30-250 bp), Varlociraptor is able to double (Lancet, Delly) or even more than double (Manta) the Recall, while never incurring notable losses in Precision, but usually rather improving on Precision.

#### Posterior probabilities allow for a clear distinction between true and false positives

The precision-recall curves of Varlociraptor show a sharp turning point at their upper right. This indicates that Varlociraptor’s posterior probabilities allow for a clear and fine-grained distinction between true and false positives. Note that the vertical parts of these lines therefore also indicate the maximum recall one can achieve, the limit of which corresponds to the list of variant calls provided by the caller. In other words, Varlociraptor is able to identify all, or nearly all of the true variants that are provided as candidates. To further solidify this observation, we investigated the posterior probability distributions of Varlociraptor. Figures 4 (deletions) and S4 (insertions) show that Varlociraptor’s probabilities are indeed clearly separating true from false positives.

**Figure 4:**
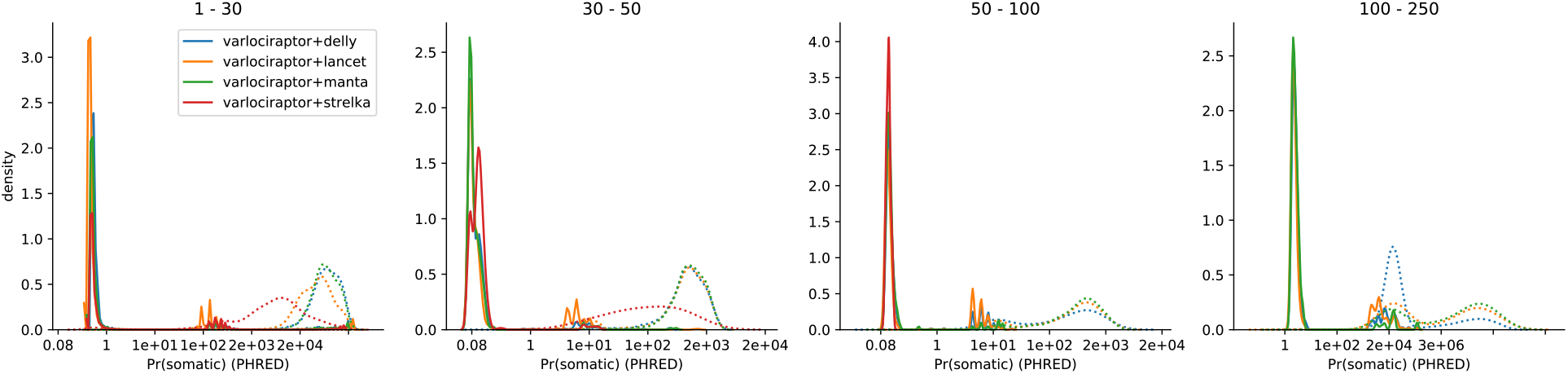
Posterior probability distributions for somatic deletions. Results are grouped by deletion length, denoted as interval at the top of the plot. The x-axis indicates the (PHRED-scaled) probability, and the y-axis indicates relative amounts of calls with this probability. The distributions of posteriors for true positive calls are shown as solid lines, the distributions of posteriors for false positive calls are shown as dotted lines.

#### Varlociraptor reliably controls false discovery rate (FDR)

Varlociraptor summarizes the uncertainty about a putative variant in terms of a single, and, as we pointed out so far, reliable quantity: an estimate of the posterior probability for the putative variant to be somatic in the tumor. It remains to determine which of the variants to output. This is a ubiquitous issue in sequencing based variant calling: optimally, one outputs maximum amounts of variants while ensuring that the mistaken predictions among the output variants do not exceed a preferably user-defined ratio. In other words, an optimal scenario is to employ a statistically sound routine that establishes *false discovery rate (FDR) control*.

Note that establishing statistically sound FDR control in variant discovery, to the best of our knowledge, amounts to a novelty: all of the state-of-the-art methods that we benchmarked against either cannot control FDR, or establish it through ad-hoc methods, such that theoretical guarantees cannot be provided.

For Varlociraptor, it is rather straightforward to establish FDR control: well-known theory [Mueller et al., 2004] points out how to achieve maximally large output at a desired level of FDR control, when working with Bayesian type posterior probabilities. See Figure 5 (deletions) and Figure S2 (insertions) for the evaluation of how Varlociraptor’s FDR control performs. In general, the closer to the diagonal, the better FDR control can be established by the user.

For deletions (Figure 5) Varlociraptor controls FDR, in the sense that across all deletion length ranges the curve is on or below the diagonal, and that deviations from the diagonal are small. The latter means that Varlociraptor tends to be slightly conservative in some combinations of length range and FDR threshold provided. One reason contributing to this is that, induced by too coarse MAPQ values or base quality scores, the resolution of posterior probabilities may be limited.

**Figure 5:**
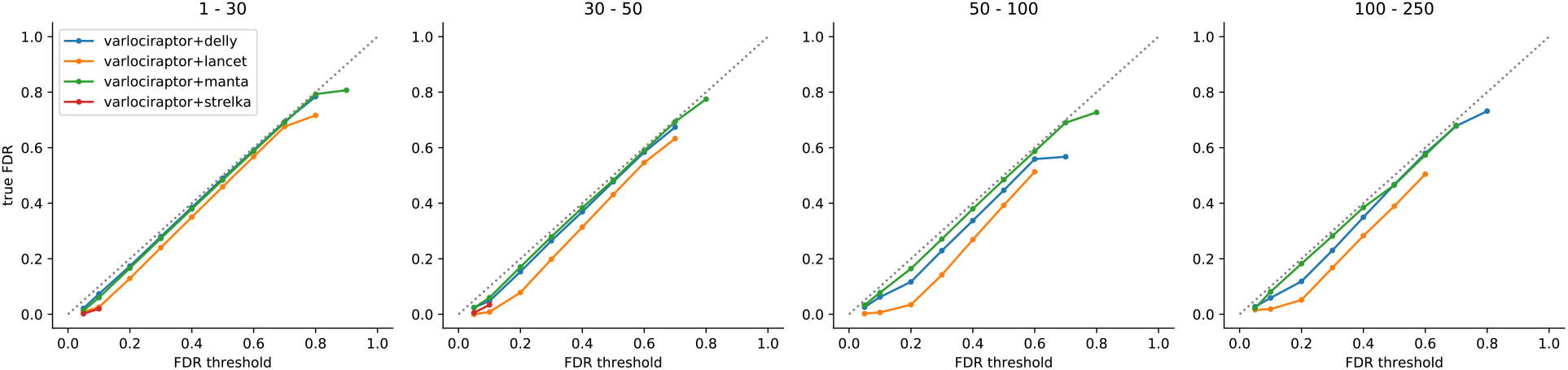
FDR control for somatic deletions. Results are grouped by deletion length, denoted as interval at the top of the plot. The axes denote the desired FDR, provided by the user as input (x-axis), and the true achieved FDR (y-axis). A perfect FDR control would keep the curve exactly on the dashed diagonal. Below the diagonal, the control is conservative. Above the diagonal, the FDR would be underestimated. Importantly, points below the diagonal mean that the true FDR is smaller than the threshold provided, which means that FDR control is still established; in this sense, points below the diagonal are preferable over points above the diagonal.

As for insertions (see Figure S2), which, as we recall, are generally more challenging, Varlociraptor equally achieves high-quality FDR control, across all length ranges and FDR thresholds. There is one caveat: for insertions of length 30-100 provided by Lancet [Narzisi], Varlociraptor’s FDR is slightly greater than the threshold specified by the user (and, although deviations are tiny, in this sense does not control FDR). An explanation for this to happen is the modus operandi of Lancet: Lancet bases insertion calls on microassemblies that are computed from all reads mapping to the variant locus. This approach is reasonable for large insertions, which do not fit into single reads, because their length either exceeds the read length, or are too long to show in single reads at full length. However, microassembly may also lead to false positives in repetitive areas where both alignments and assemblies can lead to ambiguities; note that Lancet only reaches a precision of 90% for larger insertions. Varlociraptor, at this point in time, cannot quantify all uncertainties emerging from microassemblies—we consider it highly interesting future work to also quantify uncertainties that are associated with microassemblies (see section 3).

#### Varlociraptor accurately estimates variant allele frequency (VAF)

Figures 6 (deletions) and S3 (insertions) show the difference between the true VAF’s and the ones predicted by Varlociraptor, henceforth referred to as prediction error. The first observation is that the prediction error is approximately centered around zero in all cases, which is the desired scenario. We further investigated the effect of sequencing depth on the accuracy of the VAF estimates. Figures S6 (deletions) and S7 (insertions) show how the prediction error varies relative to sequencing depth (number of non-ambiguously mapped fragments overlapping the variant locus), denoted by n and true VAF, denoted by *θ*\*. For each combination of *n* and *θ*\*, we determine an expected baseline error, modeling that one samples *n* fragments each of which stems from a variant-affected genome copy with probability *θ*\*. In other words, the expected baseline error is governed by a Binomial distribution *B*(*n, θ*\*). Accordingly, we determine

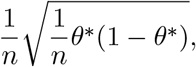

that is the standard deviation of *B*(*n, θ*\*), divided (normalized) by the depth, as the expected baseline error. This establishes the theoretical optimum of a VAF estimation procedure. In other words, no sound estimator can achieve smaller error in prediction. We see that, on average, our prediction is close to this theoretical optimum. Further, the accuracy of VAF estimates increases on increasing sequencing depth, which is the desired, logical behaviour. In summary, these results point out that the estimates are sound, and even close to what one can optimally achieve in theory. A possible explanation for the remaining small deviations from the theoretical optimum are reads that stem from different loci, and because of ambiguity in placement get aligned to the variant locus. If the read mapper is unable to reflect such ambiguity in the MAPQ score, for example because the true locus of origin is due to variation not properly represented in the reference sequence, our model is as well not able to properly reflect this in the allele frequency estimation. We expect such problems to mitigate with longer reads and more accurate (even non-linear) reference genomes in the future.

**Figure 6:**
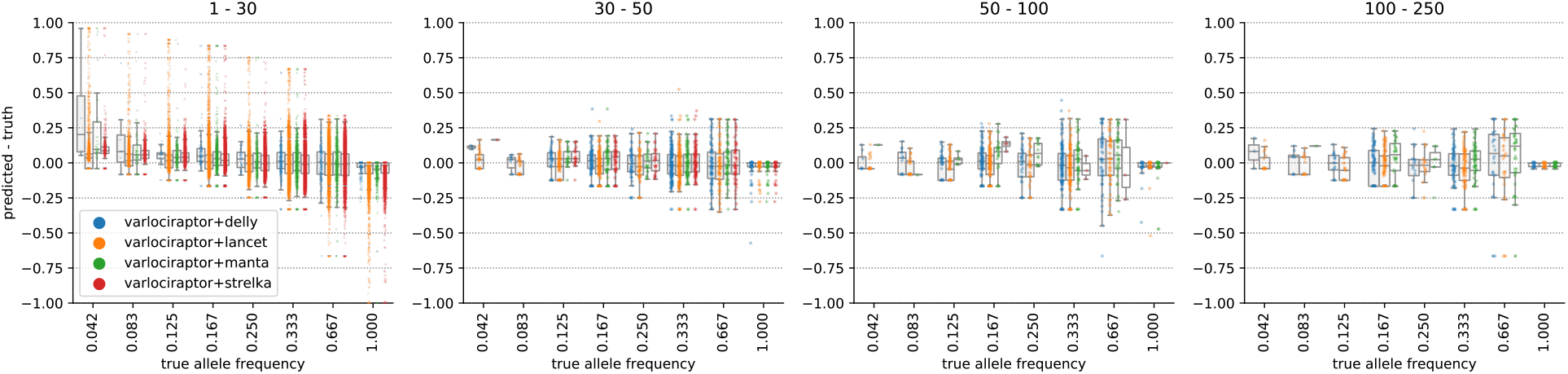
Allele frequency estimation for somatic deletions. Results are grouped by deletion length, denoted as interval at the top of the plot. The horizontal axis shows the true allele frequency, the vertical axis shows the error between predicted allele frequency and truth.

#### Varlociraptor achieves superior concordance above VAF of 20%

We applied all callers (see section 2.5) on the four replicates of cancer-control genome pairs described above (section 2.4), using their default parameters. In all analyses, because of the lack of prior knowledge available, Varlociraptor assumed a purity level *α* of 100%. We then performed a concordance analysis in the following way.

For each caller we collected calls for each of the four replicates, both when run in standalone fashion and when postprocessing calls with Varlociraptor. For each of the calling strategies, we then computed matchings across the four replicates as described at the beginning of section 2.6. For each calling strategy, we then constructed a graph where each node represents one variant call in one replicate and edges indicate that two calls (from different replicates) are matched. We then consider the connected components of this graph: any non-trivial connected component (that is any connected component consisting of more than one node) counts as *concordant call*. We then determine *concordance* as the ratio of concordant calls over all connected components

In Figures 7 and S5, we display for each possible VAF, the concordance of all calls with an at least as high VAF. In other words, for each VAF threshold *t* ∈ [0,1] we display the concordance all calls with a VAF ≥ *t*. To allow for a fair comparison, we use Varlociraptor’s VAF estimates for all calling strategies. It becomes immediately evident that Varlociraptor achieves superior concordance for VAFs of 20% and higher.

**Figure 7:**
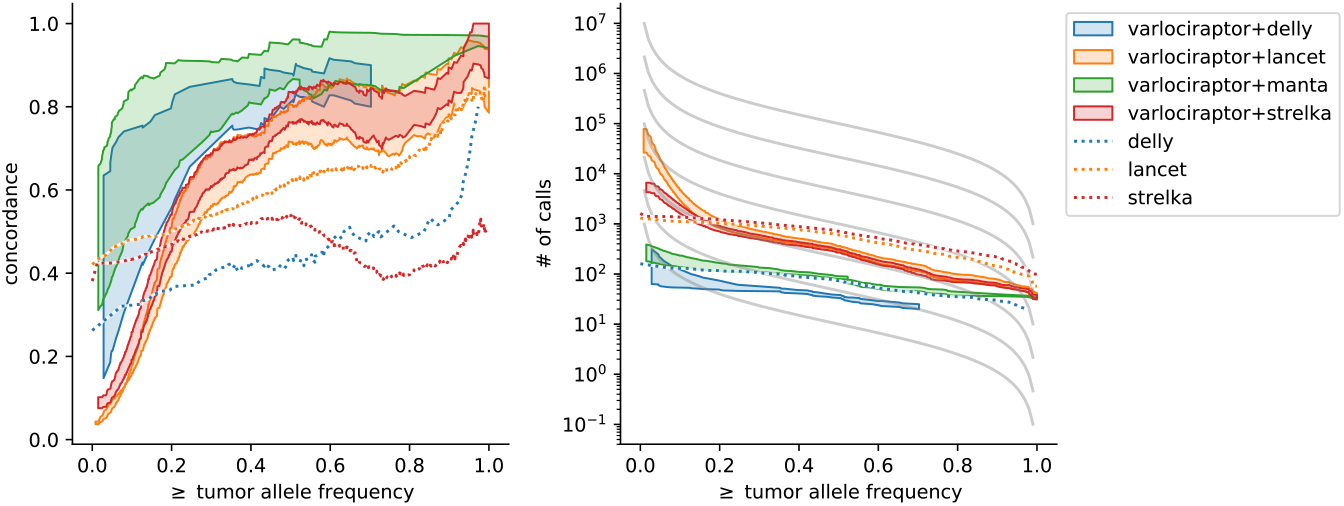
Concordance of somatic deletions on real data. For Varlociraptor, the interval between all calls with a posterior probability of at least 0.9 and at least 0.99 is shown as shaded area. Left: Concordance vs. minimum allele frequency. Right: Number of calls vs. minimum allele frequency. The different grey lines depict the theoretical expectation at different effective mutation rates according to Williams et al. [Williams et al., 2016] (see text).

#### Varlociraptor’s variant counts of VAF below 20% agree with with the theoretical expectation under neutral evolution

When inspecting all calls with a minimum VAF below 20% (i.e. *t* < 0.2, see above), Varlociraptor’s concordance drops below the rates achieved by callers run in standalone fashion. This does not mean, however, that Varlociraptor’s performance is worse. Note first that Strelka and Lancet, which appear to have superior concordance, raise only very little (and, as we will point out below, according to evolutionary models too little) discoveries. This can be seen in the right panels of Figures 7 and S5. In contrast, Varlociraptor raises substantially more discoveries at these frequencies. Second, for low frequency variants, data may not reach the necessary degree of certainty (which leads to a call by Varlociraptor) in sufficiently many of the four samples, and, since the four replicates were raised in different laboratories, low frequency variants unique to samples can be expected [Craig et al., 2016].

To explore this further and assess whether the larger numbers of low frequency calls provided by Varlociraptor are potentially correct, we compared the somatic variant count distributions with the theoretical expectation under neutral evolution (see gray lines in the right panels of Figures 7 and S5). For this, we employ the tumor evolution model by Williams et al. [Williams et al., 2016], and calculate the expected number of somatic variants of at least allele frequency *f* as

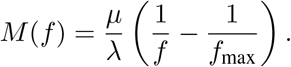

Here, *μ* is the somatic mutation rate, λ is the growth rate, and *f*_max_ is the maximum clonal allele frequency. Because there are no reliable estimates for *f*_max_ available, we set *f*_max_ = 1.0. We then plot the expected counts at various values of 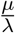 (the effective mutation rate). It can be seen that the counts of low frequency variants provided by Varlociraptor (shaded red and orange areas) are closer to the theoretical expectation than those of Lancet and Strelka (dotted red and orange lines). This is an indication that, by evolutionary principles, it is likely that many low frequency calls that were not recognized as concordant, are still correct.

## 3 Discussion

We have presented a statistical framework for the calling of somatic insertions and deletions from matched tumor-normal genome samples whose application has yielded substantial increases in terms of true discoveries, while safely limiting the amount of false discoveries. The framework is implemented in an easy-to-use open source software, called Varlociraptor^17^. In comparison with the state of the art, we have demonstrated to double, or even triple the amount of true discoveries while not increasing the false discovery rate, or suppressing it even further.

We have chosen to rely on external tools for the discovery of variants and focus on providing a sound and rigorous statistical treatment of the differential analysis (or the classification) of variants into the events relevant for somatic variant calling. An immediate benefit of this strategy is that Varlociraptor can be conveniently integrated in large-scale projects, as a post-processing step in production-quality somatic variant calling pipelines. This particularly applies when projects are run by large consortia that manage various, often heterogeneous combinations of variant callers: as per its design, Varlociraptor offers the first approach that is able to analyze sets of variants raised by different callers, all of which come with their own strengths, weaknesses, and blind spots, in a *statistically unifying* way. Because Varlociraptor preserves and combines the individual strengths of the callers while eliminating any of their particular weaknesses, it is able to raise call sheets of superior quality. In summary, the application of Varlociraptor has the potential to lead to substantial increases in true somatic indel discoveries—possibly even overwhelming in certain size ranges—in large-scale matched tumor-normal genome sequencing projects.

The technical challenge has been to overcome a well-known and notorious computational bottleneck that relates to the quantification of the uncertainties affecting the differential analysis. Thereby, not only ambiguities in terms of gap placement and aligning reads in general, which had been dealt with in the literature abundantly before, but also effects such as cancer heterogeneity (implying uncertain variant allele frequencies), purity of tumor samples, bias in terms of sampling indel-affected fragments, and strand bias, are major factors. We have presented, to the best of our knowledge, the first model that allows to capture all relevant effects and quantify all inherent uncertainties computationally efficiently. In particular, we want to stress that the presented model is the first approach that takes strand bias as a major source for systematic artifacts into a statistically comprehensive account, which is not possible if strand bias is quantified through independently raised auxiliary scores. An important aspect is the fact that the presented model can be considered as a “white box”, in the sense that all parameters have a direct biological interpretation. Hence, it supports the investigation of variant calls at different levels of detail, depending on the research question. First, one applies global filtering via FDR control. The remaining calls can optionally be investigated more closely, by looking at the estimated allele frequencies and their posterior distribution, the estimated sampling bias (see section 5.3.2 in Methods), the likelihoods and strand support of each fragment with associated mapping qualities, and finally, the read alignments themselves (if necessary). This becomes particularly important in the era of personalized medicine, where variant calls for individual patients might lead to therapy decisions, requiring upmost certainty and transparency in the decision process.

Because the statistical model comprehensively addresses all relevant effects and related uncertainties, filtering the posterior probabilities derived from the model gives immediate rise to accurate statistically sound (fully Bayesian) FDR control, for the first time in the variant calling field.

The advantage of such a straightforward and statistically interpretable filtering procedure becomes apparent when comparing it with the methodology of prior state-of-the-art approaches: so far, they have been relying on a variety of (often independently raised) scores, where combinations of (often manually) fine tuned default thresholds are supposed to ensure that predictions contain reasonably little amounts of false discoveries. Modifying these combinations of scores in order to change the default FDR, provided by the developers through the default settings, is tedious, difficult, and error-prone, if not entirely impossible.

The analysis of somatic variants often serves the purpose to yield further insight into the clonal structure of a tumor which includes to assess the allele frequencies of somatic variants appropriately. Varlociraptor, again as per its design, does not only assess the probability of individual putative somatic variants to be true discoveries, but also equips them with an estimate for their allele frequency. We have demonstrated that the corresponding VAF estimates are of almost optimal accuracy.

Still, as always, there is room for improvements. First, Varlociraptor so far only deals with insertions, deletions, and single nucleotide variants (which are not analyzed in this work). Note however the framework presented is generic in terms of its dependency structures. Therefore, efficient computation of the central likelihood function is also warranted for other variant classes; the only thing required is to adapt the computation of the typing uncertainty (see section 5.3.1 in Methods).

Second, we have been focusing on second-generation paired-end reads in this treatment, motivated by the fact that the vast majority of reads sequenced to date belongs to this class. However, our model is entirely agnostic to any particular choice of sequencing platform and can be used without any further adaptations: the only basic requirement is that the sequencing/mapping protocol in use yields (sufficiently reliable) MAPQ values. Any particular sequencing/mapping specific issues will be taken care of by the re-alignment step that Varlociraptor routinely makes use of for accurately quantifying alignment related uncertainties. When re-parameterizing the pair HMM that underlies the re-alignment step (which reflects a straightforward adaptation) to emerging long read technologies like Nanopore^18^ or SMRT sequencing^19^, Varlociraptor will be particularly apt for dealing with insertions and deletions within repeat regions.

Third, all quantities relating to uncertainties or spelled out probabilities that Varlociraptor (comprehensively) provides can be used further in a whole range of downstream analyses. Intriguing examples of such potential applications are the probabilistic assessment of larger somatic gains and losses, or the (partial) phasing of tumor subclones, which we will explore in future work.

Finally, it is important to note that the computational insights and the model presented in this work is, although motivated by, not at all limited to somatic variant calling. In fact, it turns out that it can be generalized towards arbitrary variant calling scenarios that require a differential analysis, where distinguishing between variants affecting the primary tumor and variants showing in metastases or relapse tumors, or distinguishing variants recurringly showing in different individual tumors from variants that do not, are immediate, relevant examples. The latest release of Varlociraptor already provides a *variant calling grammar*, as an interface for defining such scenarios^20^ and thereby a foundation for a unifying theory of variant calling, enabling us to explore entirely new fields of applications.

## Supporting information

Snakemake Report

## 4 Acknowledgements

We thank David Lähnemann for fruitful discussions and help with benchmarking and performance optimization. Johannes Köster was supported by NWO Veni grant 016.Veni.173.176.

## 5 Methods

### 5.1 Notation

We denote *observable variables* by Latin capital letters (e.g. *Z*). *Realizations* of these variables are denoted by small Latin letters (e.g. *z*). *Hidden/latent variables* are denoted by small Greek symbols. Vectors are denoted by boldface letters (e.g. ***Z*** = (*Z*_1_,…, *Z_k_*) or *z* = (*z*_1_,…, *z_k_*). We use super-/subscripts *h* and *t* for the healthy and the tumor sample and c to only refer to cancer cells within the tumor sample.

Let us fix a particular variant locus; we then denote the relevant read data in the healthy and the tumor sample by 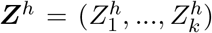 and 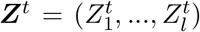, where each of the 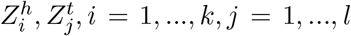 represents one (paired-end) read that (or parts of which) became aligned across or nearby the fixed variant locus. For selecting reads via alignments, we use BWA-Mem [Li and Homer, 2010] in the following, although the choice of particular aligner is free, as long as the aligner outputs a MAPQ value, which quantifies the certainty by which the reads stem from the locus under consideration.

By *variant allele frequency (VAF)*, we refer to the fraction of genome copies in the sample affected by the variant. We denote this (unknown) frequency in the healthy and the tumor sample by *θ_h_* and *θ_t_*, respectively. Since healthy cells are diploid, *θ_h_* is either 0,1/2, or 1 corresponding to absence, heterozygosity and homozygosity, when dealing with a *germline variant*. It is common that healthy cells, beyond germline variation, also exhibit somatic mutations. Somatic variants that affect healthy cells usually occur at subclonal rates, i.e. are generally not characteristic in terms of giving rise to subpopulations among the healthy cells. It is therefore reasonable to assume that somatic variants affect only less than half of the cells, reflected by allowing *θ_h_* ∈ (0, 0.5).

Prior to variant analysis, knowledge about *θ_t_* is usually not available, because only variant analysis itself can yield insight into the clonal structure of a tumor. It is therefore reasonable to assume that *θ_t_* ∈ [0, 1], that is we allow any possible VAF to apply for a somatic variant in a tumor cell.

A tumor genome sample can still contain non-negligible amounts of healthy cells, which affects our considerations. Therefore, let *α* ∈ [0,1] be the *purity* of the tumor genome sample, that is the relative amount of fragments in the sample that stem from a cancerous genome copy. Thus, 1 − *α* is the relative amount of fragments that stem from a healthy genome copy from within the tumor genome sample. For *θ_c_*, defined to be the VAF among only the cancer cells in the tumor sample, we obtain the relationship

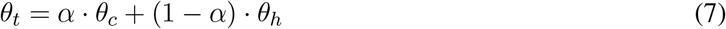

which, just as *θ_t_* can take values in all of the unit interval [0,1] for a somatic variant.

Finally, *strand bias* tends to introduce non-negligible issues in variant analysis. Our analyses are affected by strand bias as well. Strand bias refers to the fact that during the sequencing process, certain sequence motifs can cause the sequencing process to slow down, or even temporarily stall. If such delays occur, systematic sequencing artifacts affect the reads. Because the motif disappears when considering the complementary strand (unless the motif is a reverse complementary palindrome), artifact-inducing motifs usually affect reads from only one of the strands. This implies that the occurrence of artifacts depends on the origin of the read: artifacts can only affect reads from one of the strands, either the forward or the reverse strand, while reads sampled from the other strand are not affected by the sequencing artifact. See Allhoff et al. [2013] for more details.

In summary, there are three possible cases we need to deal with: *first*, a putative variant affects only reads sampled from the reverse strand, *second*, the putative variant affects reads from forward and reverse strands at equal rates, or *third*, the putative variant only affects reads sampled from the forward strand. The exclusive support of a variant by reads from only one of the two strands is indicative of an artifact [Li, 2014]. The goal is to remove such artifacts from the output.

We approach this by introducing a variable *β* taking values in 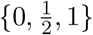 reflecting the three cases from above (so *β* = 1 reflects that the variant only shows on reads that stem from the forward strand, and so on). In other words, *β* reflects the probability that a read that is associated with the variant stems from the forward strand.

### 5.2 Motivation: Why Naive Approaches to Computing (1) Fail

To understand why *efficient computation of* (1) *is difficult*, consider that each of the reads 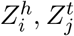 could

a. not stem from the particular variant locus,
b. stem from the locus, but is not affected by the variant,
c. stem from the locus, and is indeed affected by the variant.

We recall that it can be particularly difficult to be certain about (a), (b) or (c) when dealing with reads being associated with midsize indel loci (30-250 bp; sometimes termed the “NGS twilight zone”). Let *k* = |***Z***^*t*^| and *l* = |***Z***^*h*^| be the read coverage of the locus in the tumor and the healthy sample. Since there are 3 different possibilities—namely (a), (b) or (c)—for the overall *k* + *l* reads, we obtain that there are 3^*k*+1^ different scenarios that could reflect the truth, all of which apply with a particular probability. For computing (1) following a *fully Bayesian approach to inverse uncertainty quantification* [Liu et al., 2009]—which is the approved and canonical way to quantify uncertainties in our setting—one needs to integrate over all the possible *k* + *l* choices. In a naive approach, this translates into computing a sum with 3^*k*+1^ summands. Because *k* + *l* amounts to at least 60 to 70 in standard settings, naive approaches fail to compute the integral in human feasible runtime. This is further aggravated because one usually needs to consider hundreds of thousands of putative indel loci. *So, methodical efforts are required for uncertainty quantification in our setting*.

### 5.3 The Model

We present a graphical model that captures all dependency relationships among the variables relating to the computation of *L*(*θ_h_, θ_c_, β* | ***Z***^*h*^, ***Z***^*t*^) while taking all major uncertainties into account (see Figure 8 (**b**)).

**Figure 8:**
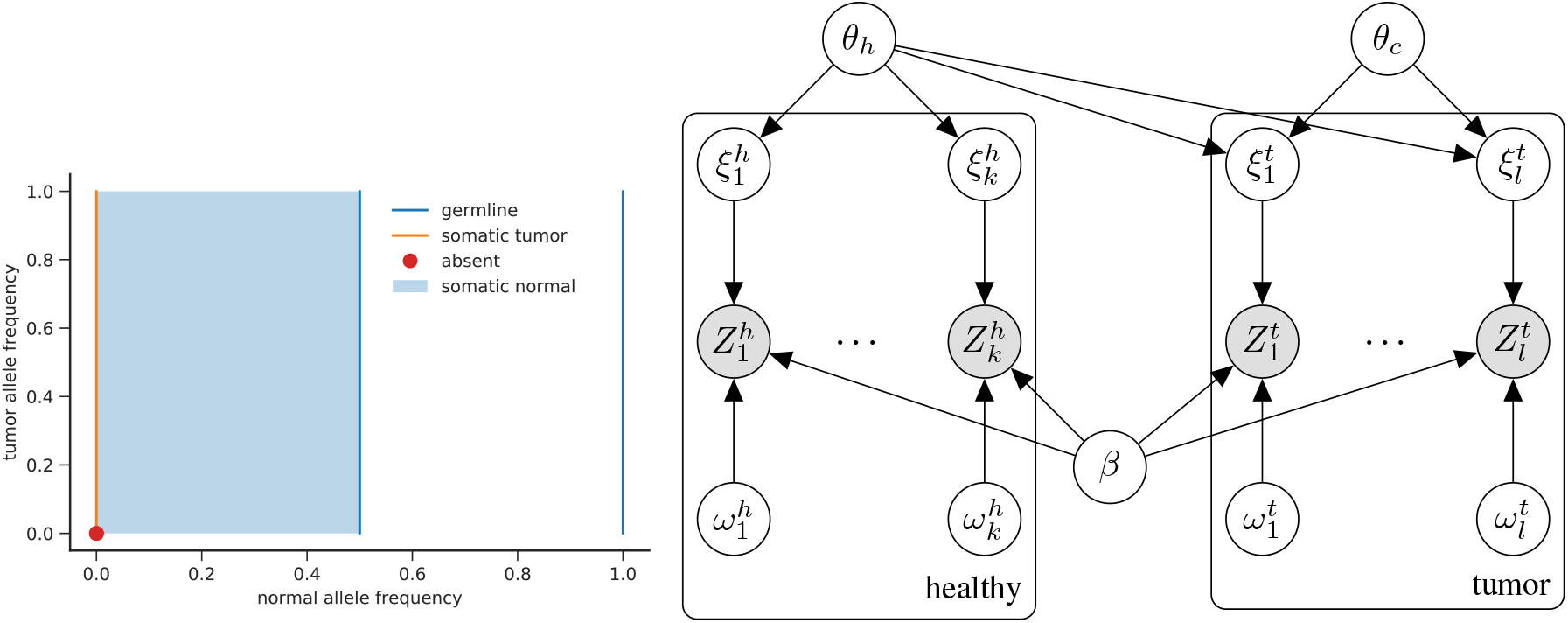
Left: Visualization of the parameter space Θ of the VAFs. *Orange*: somatic variants agree with (*θ_h_, θ_c_*) ∈ {0} × (0,1], which means that no healthy cells have the variant (*θ_h_* = 0), while some cancer clones do have the variant (*θ_c_* > 0). Germline variants (*blue*) are described by 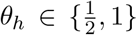 and absent variants (*red dot*) by *θ_h_* = 0, *θ_c_* = 0. Subclonal somatic variants in the healthy (normal) tissue are described by *θ_h_* ∈ (0,0.5). Right: Diagram of the model presented in section 5.3 (white circles=latent variables; grey circles=observable variables). Each column corresponds to one alignment 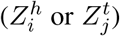 with its hyperparameters 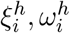 or 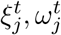. Due to (potential) sample impurity (denoted by *α* in the text), *θ_h_* has an influence on the alignments 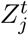 from the tumor sample.

Beyond observable variables reflecting observable read data 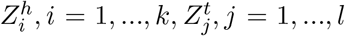 and latent variables *θ_h_, θ_c_, β* for allele frequencies and strand bias, we first introduce two additional latent variables. We denote with 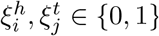 whether fragments 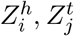 are associated with the variant 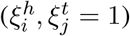 or not 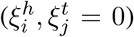. Further, 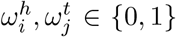 indicate whether reads indeed stem from the locus 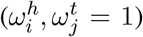 or not 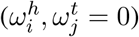.

Variables 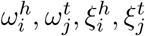 refer to the above-mentioned three cases (a), (b), (c). For example 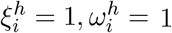 represents the case that read 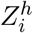 stems from the locus of interest and and is indeed associated with the variant (case (c) above). Note that the case *ω* = 0, which indicates that the read does not stem from the locus (a), renders specification of other variables obsolete, because this particular read cannot provide information about the variant. Since knowledge about realizations of the hyperparameters cannot be observed at the time of the analysis, they are latent variables.

In the following, we introduce the full model in three steps, thereby deriving the conditional dependencies between the different variables. *First*, we consider the basic model. *Second*, we introduce *impurity*, modeling the fact that the cancer genome sample also contains healthy genome copies. This implies to distinguish between the treatment of reads from healthy and tumor sample: while reads from the healthy sample still follow the basic model, processing reads from the tumor sample requires a modification to take impurity into account. *Third*, we introduce *strand bias*, which would affect both reads from the healthy and the tumor sample and thus requires to apply modifications for both parts.

**Part 1 – Foundations: Alignment and Typing Uncertainty.** We first model the two major sources of uncertainty, (a) *alignment uncertainty* and (b) *typing uncertainty*. Let *Z_i_, i* = 1,…, *k* be the observable alignment data. *Alignment uncertainty* is handled via

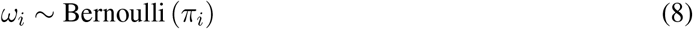

where *π_i_* reflects the probability that the *i*-th read has been aligned to the correct position in the genome. Estimates for *π_i_* are provided by the aligner of choice, via the reported MAPQ values [Li et al., 2008a]. *Typing uncertainty* can be modeled as

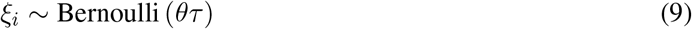

Thereby, the allele frequency *θ*, as per its definition, reflects the probability to sample a read from a variant-affected genome copy. Further, *τ* reflects the probability that, if sampled from the variant-affected copy, the read indeed covers the variant. That is, the product *θ_τ_* reflects to sample a fragment that is truly affected with the variant (and hence provides real evidence about the variant). See section 5.3.2 for how *τ* is computed. Note that *θ* and *τ* vary depending on whether *Z_i_* refers to fragment data from the healthy 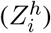 or the tumor genome 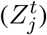. Appropriate choices for *θ, τ* are specified in the paragraph about impurity below.

Whether *ξ_i_* = 1 or *ξ_i_* = 0 is generally not immediately evident from the observed (paired-end) read *Z_i_*, leading to *typing uncertainty*. We define

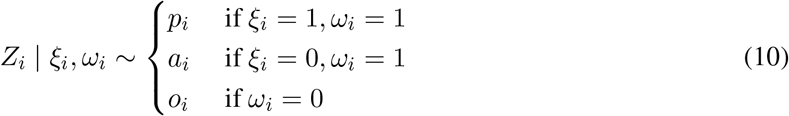

where *a_i_, p_i_, o_i_* are the probability distributions that reflect the situation that *Z_i_* stems from a genome copy in which the indel is either *p*resent or *a*bsent, or comes from an (unknown) *o*ther locus. In the last case, *Z_i_* is supposed to have no influence on the posterior probability distribution of *θ*.

In order to illustrate the nature of *p_i_, a_i_, o_i_*, let us consider a simplified case. The detailed definition can be found in section 5.3.1. We denote two different haplotypes *H*_ref_ and *H*_var_. The former represents thŧ reference sequence (no variant) and the latter the alternative (variant affected) sequence at the considered locus. Let *x* be a particular read. Let further

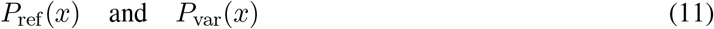

be the probabilities that *x* has been sampled from *H*_ref_ or from *H*_var_, respectively. Because *x* may contain errors, one needs to take the sequencing error profile of *x* into account when aiming at accurate computation of *P*_ref_(*x*) and *P*_var_(*x*). The error profile is provided via the base qualities reported by the sequencing machine^21^.

As described in earlier work [Poplin et al., 2018], probabilities (11) can be reliably computed by means of a Pair HMM whose parameters refer to the base quality profile of *x*, thereby appropriately accounting for the sequencing errors affecting *x*. Following this well-approved rationale, we model

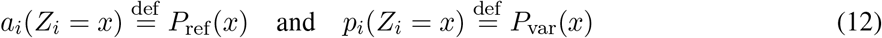

In a final remark, note that for that sake of a clear presentation, we have omitted the detail that *p_i_, a_i_* also reflect considerations about the fragment length distributions of the involved reads. See Subsection 5.3.1 for how *a_i_, p_i_* and *o_i_* are computed in full detail.

Finally, *o_i_* reflects the case that *x* does not stem from the locus. Without further knowledge available— which is the case at the time of analysis—it is reasonable to assume that *o_i_* is equal for all possible reads *x*. In other words, it is reasonable to assume that *o_i_* reflects a uniform distribution. The only detail to consider (which follows from the theoretical statements listed in section 5.3.4), is that the particular value *o*(*Z_i_* = *x*) needs to scale right relative to *p_i_*(*Z_i_* = *x*) and *a_i_*(*Z_i_* = *x*), see again section 5.3.1 for the respective details.

**Part 2 – Impurity.** Let *α* be the purity of the tumor genome sample. In turn, 1 − *α* is the ratio of fragments stemming from a healthy genome copy, when sampled from the tumor sample. Impurity does not affect the healthy sample, so (8), (9), (10) and (12) apply without further modifications for the healthy sample: variables can be indexed with super- or subscript *h* as they appear. For example 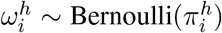 in (8) or 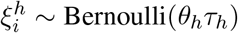 in (9).

However, variables referring to the tumor sample (indexed using super- or subscript *t*) require different treatment. Let 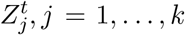 be the observable read data in the tumor sample. First note that impurity does not affect the degree of certainty of read alignments. So (8) applies without modifications:

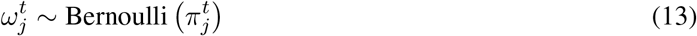

Modeling typing uncertainty however requires changes. Recalling (7), we compute

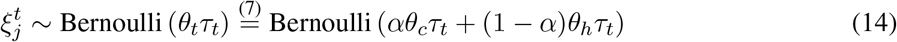

This reflects that if reads stem from a cancerous genome copy, which happens with probability *α*, variables *θ_c_, τ_t_* apply, and if stemming from a healthy genome copy, which happens with probability 1 − *α*, variables *θ_h_, τ_t_* apply. Note that although stemming from a healthy genome copy (thus *θ_h_*), still *τ_t_* applies, because the healthy genome copy was sampled from the tumor sample and τ is specific to the sample, healthy or tumor (see section 5.3.2 for details).

Equations (10) and (12) remain unchanged, that is super- and subscripts *t* can be introduced without further modifications, because impurity does not matter in these considerations.

**Part 3 – Strand Bias.** In the following, we will be dealing with paired-end read data. We will therefore be focusing on this case. Note however that our model also immediately applies for single-end and mate-pair read data, via some straightforward modifications.

For sake of simplicity, let us consider reads from only the healthy sample, and for the sake of a clear presentation, we omit super- and subscripts. That is, we write 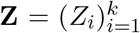 when meaning 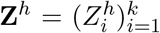 and we simply write *θ, τ* when meaning *θ_h_, τ_h_*. Extending the following arguments to the tumor sample is straightforward.

Following the usual protocols, in a paired-end read, one read end stems from the forward strand, while the other end stems from the reverse strand. When accounting for strand bias, it is important to track whether the forward or the reverse strand is associated with the variant, or even both of them (reflecting the situation of a self-overlapping read), because variants showing exclusively in only one of the types of ends are likely to reflect artifacts.

We model strand direction of read pairs by expanding **Z** into **Z** = (**R, S**), distinguishing between the read data themselves 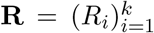 and variables 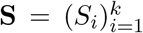. Each of the *S_i_* takes values in {−, +}, reflecting whether, the reverse (−) or the forward (+) end of read pair i are associated with the variant. For the sake of a clear presentation, we omit the case that a paired-end read has self-overlapping ends, such that it is possible that both of its ends are associated with the variant; see section 5.3.3 for the corresponding, straightforward modifications.

The value of *S_i_* only has meaning if the *i*-th read pair indeed stems from the variant haplotype, which refers to the case *ξ_i_* = 1, *ω_i_* = 1 in terms of the earlier latent variables. If *ω_i_* = 0, that is the *i*-th read pair does not stem from the variant locus, or if *ω_i_* = 1, *ξ_i_* = 0, that is, if the *i*-th read pair stems from the locus but is associated with the reference haplotype, strand bias cannot affect the *i*-th read pair. Realizations of *S_i_* are observable: we retrieve the corresponding values from the initial standard read aligner in the obvious way. Note that it is important to retrieve the realizations from the standard alignment, but not the re-alignment, because standard, reference-based alignments are a major source of strand bias artifacts.

Integrating the *S_i_*, the likelihood function extends to

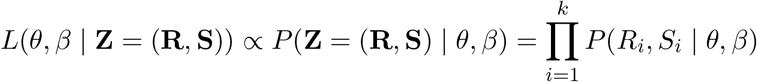

So, when treating strand bias we have to specify how to evaluate *P*(*R_i_, S_i_* | *θ, β*). We compute

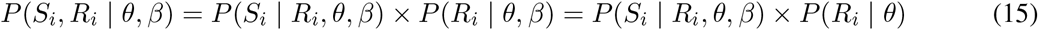

where the last equality follows from the fact that only *S_i_* depends on *β*, while *R_i_*, the observed read sequence itself, does not depend on *β*. Note that *R_i_* could be identical both for (+, −)-oriented and (−, +)-oriented read pairs, while only one of those orientations gives rise to a variant artifact. Computing *P*(*R_i_* | *θ*) is identical with the computations displayed for the cases not involving strand bias, see Part 1 and 2. So it remains to consider *P*(*S_i_* | *R_i_, θ, β*). We refer to section 5.3.3 where we elaborate on the corresponding details; here we conclude that (15) points out that strand bias can be handled efficiently by integrating additional factors *P*(*S_i_* | *R_i_, θ, β*) into the overall likelihood function.

As further outlined in the last paragraph of section 5.3.3, the factor *P*(*S_i_* | *R_i_, θ, β*) in (15) has no influence on the likelihood of *θ* if 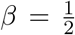. This is important, because only loci where *β* = 0 or 1 reflect strand bias artifacts and are to be removed from further considerations. Variants observed at loci where 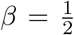 are no strand bias artifacts. Therefore, the desirable scenario is to deal with them as if strand bias was not to take into account. This means that 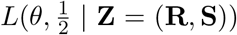 should be proportional to *L*(*θ* | **Z**), as treated before introducing strand bias; see again section 5.3.3 for further straightforward computations that prove this.

#### 5.3.1 Computing *a_i_, p_i_* and *o_i_*

We now specify distributions *a_i_* and *p_i_* for read pairs *Z_i_*. We note in passing that our model does in general not depend on a particular sequencing technology, so can also deal with single-end third-generation sequencing reads or other protocols; see the Discussion for some final remarks on that point.

As usual, let us fix a particular locus harboring a putative variant and denote the corresponding reference haplotype of that locus by *H*_ref_ and the alternative, variant-affected haplotype of the locus by *H*_var_. We compute *H*_var_ by application of the putative variant to the reference sequence (*H*_ref_) without any additional changes.

Let us consider a particular read pair *x* = (*x*_1_, *x*_2_), consisting of two ends *x*_1_, *x*_2_ (which in general are of equal length), that was found to align with the putative variant locus through use of a standard read aligner, (e.g. BWA-MEM Li [2013]). After having collected *x*, we discard all standard read alignment information about *x*, apart from one detail: we store *z*, the length of the alignment computed by the standard read aligner. Note that *z* evaluates as the distance between the rightmost alignment coordinate of the right end (*x*_2_) and the leftmost alignment coordinate of the left end (*x*_1_), including clipped bases.

Subsequently, we re-align *x* with both *H*_ref_ and *H*_var_ using a Pair HMM [Durbin et al., 1998], as it was initially suggested by Poplin et al. [2018]. The Pair HMM appropriately accounts for possible sequencing errors in *x*, because its parameters reflect the sequencing error (base quality) profile of *x*, and thereby reflects that *x* has been sampled as a fragment from *H*_ref_ or *H*_var_, by providing either *x* and *H*_ref_ or *x* and *H*_var_ as input pair of sequences.

We base all of our further considerations on the resulting re-alignments of *x* with *H*_ref_ and *H*_var_. The well-known justification for this is that such re-alignments avoid several biases standard read alignments (where only *H*_ref_ is considered) are affected with, and are therefore of considerably higher quality. When dealing with insertions and deletions, these biases include well-known effects referring to mistaken gap placement—note that placing gaps in alignments correctly has remained a notorious issue [Lunter et al., 2008]. Incomplete gaps (soft-/hard-clipped read alignments) output by the standard aligner add to these difficulties; therefore re-aligning clipped alignments can be of particular value. See Figure 9a for an illustration and Figure S8 for a read world example.

**Figure 9:**
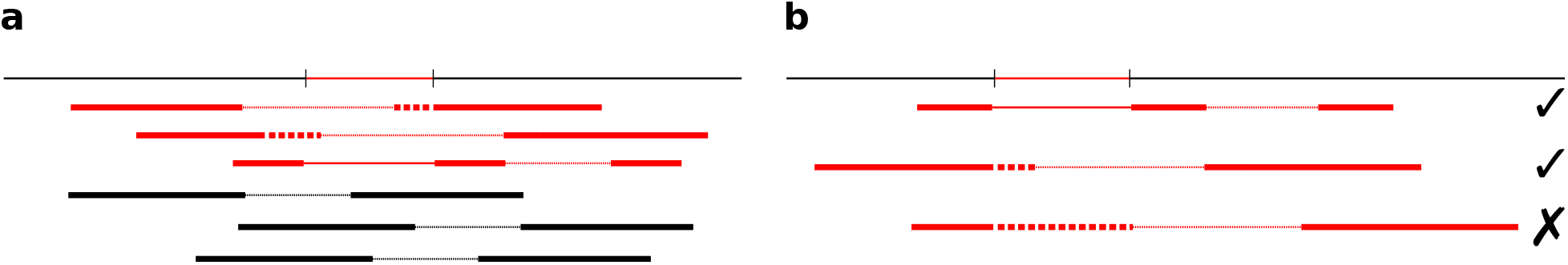
Dealing with evidence from alignments. The reference genome is displayed as a thin line at the top, with an example deletion highlighted in red. Paired-end reads are displayed as two connected thick bars. Deletions within the alignment of a read are displayed as thinner region. Soft-clips in the alignment are dotted. Fragment origin is encoded by color (red=variant; black=reference). (a) Reads coming from the variant allele can be aligned with soft-clipped ends, or with the variant being encoded in the alignment itself. Due to ambiguities, the positions are not necessarily perfectly aligned with the variant (see Figure S8 for a real example). (b) Depending on the size of the variant and properties of the read mapper on a specific genome, it can become less likely to obtain fragments from the variant allele. Here, the first and the second fragment are mappable, the third is not, because the soft-clip would become too large to be considered by the read mapper.

We derive values *a_i_*(*x*), *p_i_*(*x*), *o_i_*(*x*) from these re-alignments, reflecting probabilities that *x* has been sampled from *H*_ref_(*a_i_*), or from *H*_var_(*p_i_*), while *o_i_*, reflecting the probability that *x* does not stem from the locus, needs to scale right with *a_i_* and *p_i_*, in order to ensure numerical stability.

For aligning *x* with *H*_ref_ and *H*_var_ using a Pair HMM, we need to consider the issue that we need to specify exactly what *H*_ref_, *H*_var_ should look like. Of course, *H*_ref_, *H*_var_ cannot reflect whole-chromosome length sequences, because it is infeasible by runtime to align *x* against a full-length chromosome. Hence, we need to cut *H*_ref_ out of the full-length chromosome in an appropriate way. In particular, it is favorable that the parts of *H*_ref_ and *H*_var_ against which *x* is aligned (i.e., the parts provided as input to the Pair HMM) should be of equal length (or differ by one or two basepairs, but not differ by the length of the indel, for example), to avoid biases induced by the length of the indel under consideration.

To do this, we proceed in six steps:

1. Let *l* be the coordinate of the variant locus in the reference genome *G*. We determine *H*_ref_: = *G*[*l* − *h, l* + *h*], that is as a window of size 2*h* in the reference genome around the variant locus^22^. We then obtain *H*_var_ by applying the variant to *H*_ref_. Note that this is not yet the input for the Pair HMM.
2. Let *x*_1_ be the left end of the read pair *x* = (*x*_1_, *x*_2_). The following computations are executed also for *x*_2_, which happens entirely analogously. Taking *x*_1_ as the pattern and *H*_ref_ or *H*_var_, respectively, as the text, we apply Myers’ bitvector algorithm [Myers, 1999] and obtain coordinates *a_r_*, and *a_v_*, determined such that, relative to edit distance, the optimal occurrence of *x*_1_ in *H*_ref_ and *H*_var_ is *H*_ref_ [*a_r_, b_r_*] and *H*_var_[*a_v_, b_v_*], respectively. Note that by virtue of Myers’ bitvector algorithm, the lengths of *H*_ref_ [*a_r_, b_r_*] and *H*_var_[*a_v_, b_v_*], that is, *b_r_* − *a_r_* + 1 and *b_v_* − *a_v_* + 1, tend to differ only by at most 1 or two basepairs, which is the desired scenario. As above-mentioned, we also obtain such coordinates for the right end *x*_2_.
3. We finally specify a maximal edit distance *d*, and compute

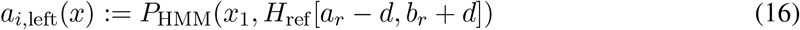

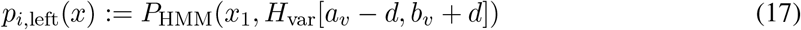

where *P*_HMM_(*x*_1_, *H*_ref_[*a_r_* − *d, b_r_* + *d*]) and *P*_HMM_(*x*_1_, *H*_var_[*a_v_* − *d, b_v_* + *d*]) are computed by means of the forward algorithm for Pair HMMs as defined by Durbin et al. [1998]. Again, probabilities *a*_*i*,right_(*x*), *p*_*i*,right_(*x*) referring to the right read end *x*_2_ are computed in the entirely analogous way, by replacing *x*_1_ with *x*_2_ and coordinates *a_r_, b_r_, a_v_, b_v_* with coordinates resulting from running Myers’ algorithm with *x*_2_ as pattern and *H*_ref_ and *H*_var_ as texts.
4. Let *f* specify the fragment length distribution referring to the sequencing library protocol in use. Note that *f* often is approximately Gaussian, hence can be specified by its mean *μ* and standard deviation *σ*. While, without any further adaptations, we can work with arbitrary (empirical) fragment length distributions, we make use of Gaussian distributions here. We recall that *z* is the length of the (standard) alignment of *x*. Note that *z* agrees with the length of the underlying fragment if *x* is not affected by the variant. If, however, *x* is affected by the variant, which is of length *δ*, then the length of the underlying fragment would evaluate as *z* + *δ*. Hence, we define

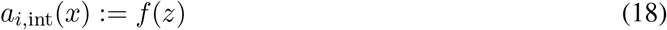

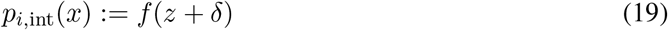

to reflect that *a*_*i*,int_ is supposed to reflect that alignment and fragment length agree, while *p*_*i*,int_ is supposed to reflect that they differ by the length of the variant.
5. We combine the evidence for the left and right read with the insert size by calculating

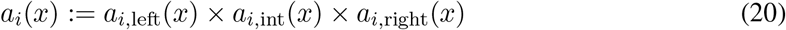

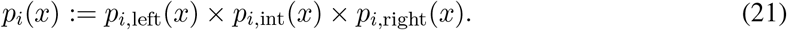
6. Finally, we denote

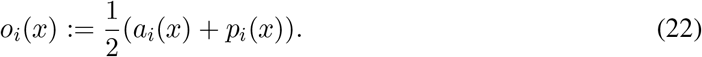 The formal justification for determining *o_i_*(*x*) as such follows from the proofs provided in section 5.3.4, which lists the formal statements that make the theoretical foundation of our model. Note that we discard all reads from being considered if neither the left read end (*x*_1_) nor the right read end (*x*_2_) overlap the variant locus as per their Pair HMM based re-alignments. Finally, let us revisit the decision to combine evidences of left and right read with the insert size. Splitting up of *a_i_,p_i_* into read end-specific factors would effectively weaken variant related signals from reads that can only be ambiguously placed, that is whose exact placement remains dubious even after re-alignment. To understand the advantage of combining evidences, consider a read *x* whose first end supports the variant, which yields large *p*_*i*,left_(*x*), but small *a*_*i*,left_(*x*), whereas the second end supports the reference, which yields large *a*_*i*,right_(*x*), but small *p*_*i*,right_(*x*). In consequence, overall, both *a_i_*(*x*) and *p_i_*(*x*) are approximately equal. The same argument holds, for example, if one or both reads support the variant, but the insert size does not. As follows from our model, reads *x* with (approximately) equal *a_i_*(*x*) and *p_i_*(*x*) yield an approximately uniform probability distribution with respect to *θ*, the allele frequency of the variant, which is just our prior distribution on *θ*. In other words, *x* does not contribute to making a statement in favor of a particular (range of) *θ*, which is the correct scenario.

#### 5.3.2 Sampling Probability

See Figure 9b for illustrations of the following. In the initial step, we select reads using a standard read aligner, which determines placement of reads by mapping them against the reference genome. Because differences between non-variant-affected reads and the reference sequence are small, the standard read aligner will be able to align all (or at least nearly all) of such reads successfully against the putative variant locus. The situation is different for reads that are affected by the variant. Because such reads differ from the reference sequence by, in our case, an insertion or deletion of considerable length, the standard read aligner may fail to align such reads successfully. Such failure occurs in particular if variant size and placement within the read interfere unfavorably with the alignment procedure. As a consequence, there can be a bias towards reads from the reference allele.

The effect was first systematically treated in [Sahlin et al., 2015]. The corresponding quantification means to assume that non-variant affected reads align against the locus with probability 1, while variant-affected reads align with *sampling probability τ*, which is smaller than one. The sampling probability *τ* is a locus-, donor genome-, and sequencing-specific parameter, because *τ* depends on the sequence context, which varies relative to the locus considered and also relative to the donor genome investigated as well as the used sequencing protocol (in terms of read length and targeted insert size). So, *τ* can be different for cancer and control genome, which we make explicit by dealing with *τ_h_* and *τ_t_*, referring to the healthy (*h*) and tumor (*t*) genome.

In the following, we briefly specify how *τ* is computed based on the considerations made by Sahlin et al. [2015]. First, *τ* depends on the fragment length distribution *f*, which can be retrieved from the read aligner; see also explanations referring to formulas (18) and (19) above. Further, *τ* depends on the number of bases *δ* the deletion or insertion covers in the donor genome (deletion: *δ* = 0; insertion: *δ* = length of insertion). It also depends on the number of bases the aligner requires to be aligned with the reference genome sequence upstream or downstream of the variant, in the upstream (left) or the downstream (right) read end, respectively, which are referred to as *k*_up_ and *k*_down_. This reflects that it is critical for a read aligner to get the outer ends of a paired-end read mapped. Values *k*_up_ and *k*_down_ depend on the read aligner and the variant. In the following, *k* is one of *k*_up_ or *k*_down_, depending on whether we consider information referring to the upstream or downstream read end; when dealing with single-end reads (see below), *k* is unique. *k* can be estimated by inspecting a representative subset of all aligned reads of a sample, and recording the maximum deletion and insertion size encoded in the CIGAR strings [Li et al., 2009] of this subset of reads, as well as the maximum soft clip size, retrieved from partial alignments from that representative subset of reads. If the variant is smaller than the maximum deletion or insertion encoded in the CIGAR strings, we can set *k* = 0 because the variant is small enough to be part of the alignment. Otherwise, *k* can be set to the read length minus the maximum soft clip size, that is, the smallest observed partial alignment.

Let further *o* denote the fragment length (which is unknown) and *f* denote the fragment length distribution (see section 5.3.1). Following Sahlin et al. [2015], we compute *τ* as

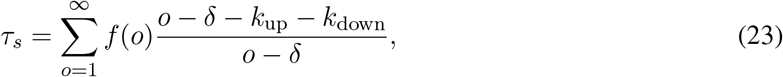

where *s = h, t* specifies the healthy or tumor sample, see above; note that all of *f, k*_up_ and *k*_down_ differ relative to the sample. The formula reflects that one sums over all possible fragment lengths, weighted by the probability to sample a fragment of that length, and calculates the probability to get a fragment of that length aligned, relative to the relevant parameters *δ, k*_up_ and *k*_down_. Note that in case of single end reads, (23) simplifies to

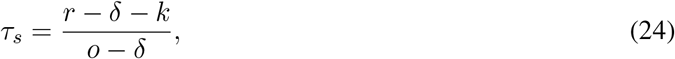

with *r* being the read length.

#### 5.3.3 Strand Bias: Technical Details

In the following, observable strand bias variables *S_i_* will take values in {−, +, ±} to also reflect the case that a paired-end read has self-overlapping reads both of which are affected by the variant (±). Note that existence of a ±-read rules out strand bias, because it means that read ends from both strands are affected by the variant.

Let us define

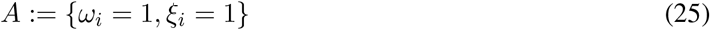

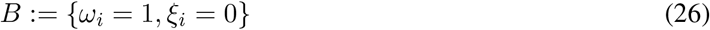

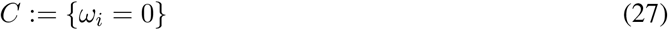

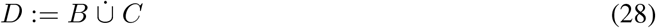

and recall that strand bias can only occur if event *A* applies, that is, if the read is indeed associated with the variant. Note that *A* and *D* span the entire space of cases (*ω_i_, ξ_i_*) ∈ {0,1} × {0,1}, which allows to apply the law of total probability. We obtain

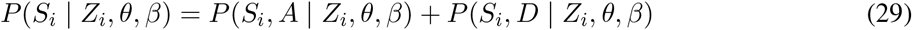

So we will continue to put further focus on the two summands in (29). Let *z_i_* be the observed realization (the read itself including its error profile) of *Z_i_*. For a clear presentation of what follows we define

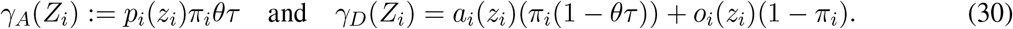

For the first summand from (29), we compute

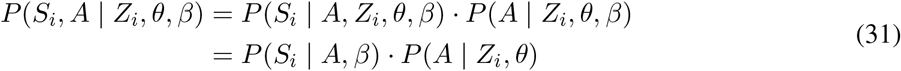

where the last equation follows from the fact that, on the one hand, given *A* and *β, S_i_* is independent of *Z_i_* and *θ* (note that all information *S_i_* depends on about *Z* and *θ* is captured by *A*) and, on the other hand, *A* is independent of *β*, the strand bias. We continue

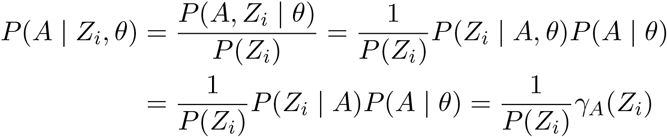

where the second to last equation follows from the dependency structure of the model, because *Z_i_* is independent of *θ* given *A*, and the last equation follows from the fundamental considerations of Part 1. Note at last that it is reasonable (and common in analogous settings) to assume that *P*(*Z_i_*), the prior probability to observe a read is constant across all reads, so 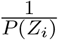 turns out to be a constant. In summary, we obtain

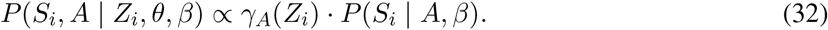

Recalling that read pairs were selected where at least one of the ends overlapped the locus, let *q*_1_ and *q*_2_ = 1 − *q*_1_ be the probabilities to have one or both reads overlapping the variant locus. Probabilities *q*_1_, *q*_2_ can be determined from the insert size distribution provided by the read mapper, in combination with the length of the read ends. Note that usually *q*_1_ >> *q*_2_, and that for single-end sequencing, obviously *q*_1_ = 1, *q*_2_ = 0. Our considerations are then finalized by providing the table

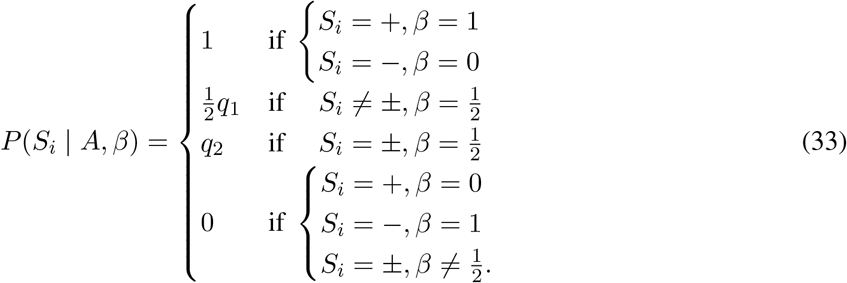

By entirely analogous considerations, we also obtain

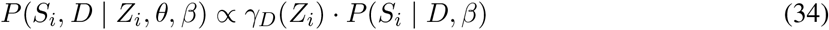

where here, however,

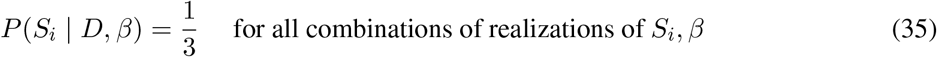

As an exemplary case, consider *γ_A_*(*Z_i_*) = 0, *γ_D_*(*Z_i_*) = 1, reflecting the case that the read is not associated with the variant with probability one. Here, we obtain that 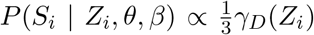 for all choices of *β*. This means that the *i*-th read assigns equal likelihood to all choices of *β*, which is the correct scenario, because the *i*-th read cannot provide any information about *β*.

Consider also the opposite case, *γ_A_*(*Z_i_*) = 1, *γ_D_*(*Z_i_*) = 0, reflecting full certainty about the read being associated with the variant. If for example *S_i_* = +, the likelihood for *β* = 0 is zero. Further (recalling that usually *q*_1_ >> *q*_2_, meaning that *q*_1_ is close to one), the likelihood for *β* = 1 is approximately double the amount as for 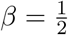. This again is the correct scenario.

Finally, we observe that if 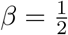, the factor *P*(*S_i_* | *Z_i_, θ, β*) in (15) has no influence on the likelihood of *θ*. This is important, because we will only consider loci where 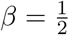 with large enough likelihood, and for such loci, we will base our further decisions on the evaluation of 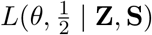. It is therefore essential that 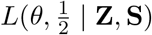 is proportional to *L*(*θ* |**Z**), that is when not considering strand bias. Considering strand bias should only lead to excluding variant artifacts—if variants are supposed to be real, strand bias considerations should not have an influence on decisions about whether variants are somatic or not, or, more specifically, about what their allele frequencies are.

To understand this, note that the only appearance of *θ* in the computation of *P*(*S_i_* | *Z_i_, θ, β*) is in the factors *γ_A_*(*Z_i_*) and *γ_D_*(*Z_i_*). Note further that these factors determine to what degree the *i*-th read makes a statement about *β*: the larger *γ_A_*(*Z_i_*) relative to *γ_D_*(*Z_i_*), the stronger the statement. If 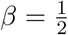, however, both *P*(*S_i_* | *A, β*) and *P*(*S_i_* | *D, β*) evaluate as 1, independently of the particular realization of *S_i_*. Therefore, if 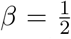, varying *θ* does not vary *P*(*S_i_*| *Z_i_, θ, β*). Of course, still, varying *θ* varies (15) because it varies *P*(*Z_i_* | *θ*). So, *θ* varies 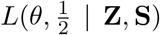, just as if *β* was not considered. In summary, this is the desired scenario.

#### 5.3.4 Statements

We finally obtain the result outlined in section 2.2 as a corollary to the following theorem.

##### Theorem 5.1.

*Let* 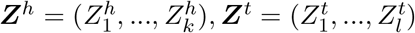 *be the observable read data from a healthy and a tumor sample, covering the locus of a putative variant. Then*

- *(i) The likelihood function*

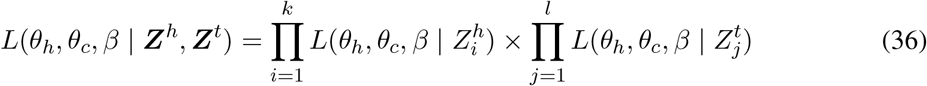

*factors into likelihood functions referring to individual read pairs*.
- *(ii) Let Z_i_ refer to any of the read data* 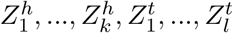 *and let ω_i_, ξ_i_ be its latent uncertainty hyperparameters. Then*

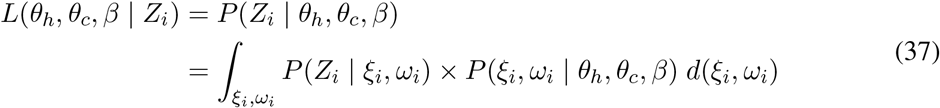

Proof. (*i*) follows immediately from the fact that the 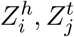 are conditionally independent given *θ_h_, θ_c_, β*; see Figure 8b. (*ii*) follows from application of the Chapman-Kolmogorov equation, in combination with the dependency relationships captured by our model, see again Figure 8b.

We distinguish between read data 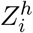 from the healthy sample and read data 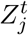 from the tumor sample. In the following, we make use of the fact that 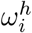 and *θ_h_* are independent (see Figure 8b). Let 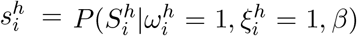 be the likelihood of the strand bias given read pair *i* as defined in equation (33). We compute the likelihood function for 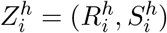, which does not depend on *θ_c_*, as

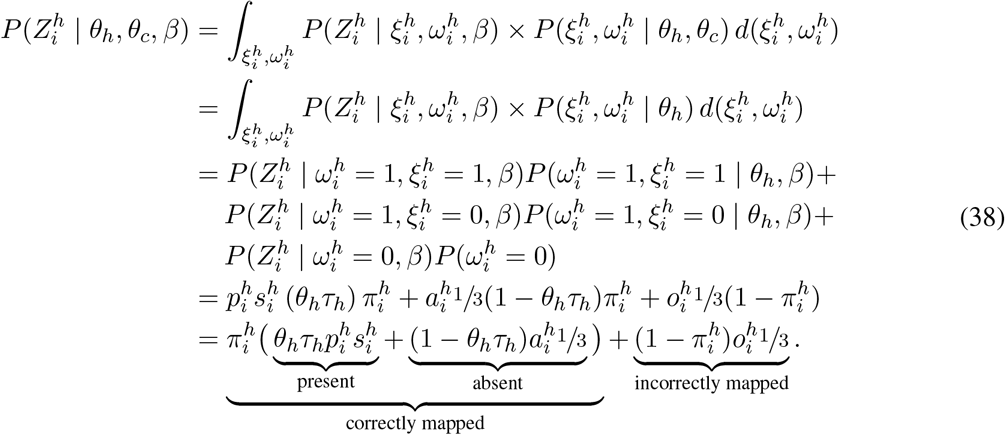

Analogously, while slightly more involved due to the purity considerations causing 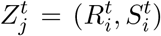 to depend on both *θ_h_* and *θ_c_*, we compute

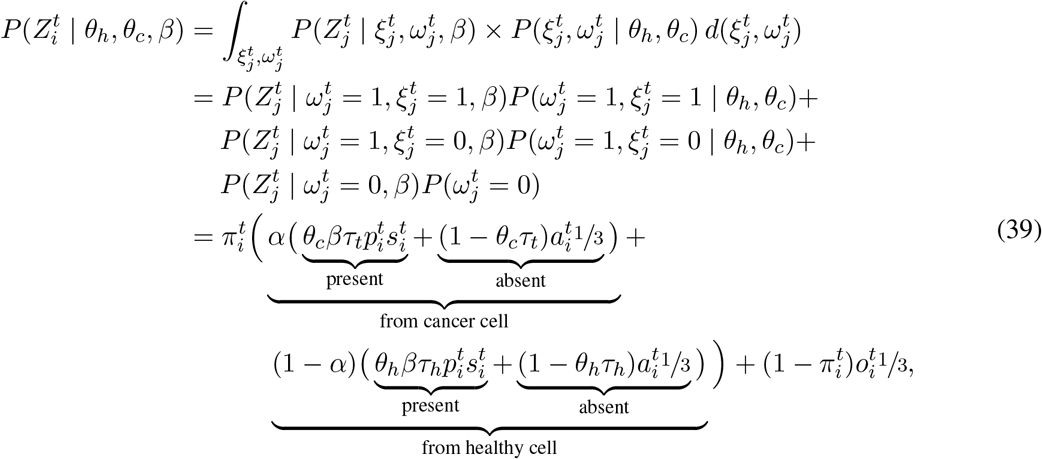

with 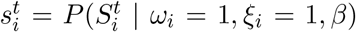 analogously to the healthy case above. As can be seen, the integral turns into a sum over the three cases {*ω_i_* = 0}, {*ω_i_* = 1, *ξ_i_* = 0}, {*ω_i_* = 1, *ξ_i_* = 1}, reflecting that *Z_i_* is either (1) incorrectly mapped, (2) correct and not affected by the variant, or (3) correct and affected by the variant. For computing *p_i_, a_i_*, and *o_i_*, we need a linear runtime in the length of the considered window, since we are using a banded pair HMM (see section 5.3.1). However, this only needs to be done once for all likelihood computations in the parameter space. We can therefore infer the following central corollary.

##### Corollary 1.

*L*(*θ_h_,θ_c_,β* | ***Z**^h^*, ***Z**^t^*) *can be computed in O*(*k + l*) *operations*.

## A Uniqueness and computation of the maximum likelihood estimate

The likelihood function of *θ_h_, θ_c_*, and *β* given the data ***Z**^h^* and ***Z**^t^* as shown in equation (1) is a higher-order polynomial, which makes it infeasible to derive its maximum analytically. We show in this section, however, that under weak conditions (as given in the following theorem) the likelihood function attains a unique global maximum on the unit interval for each value of *θ_h_* and *β*. We, in addition, show that the loglikelihood function is strictly concave, which simplifies the numerical maximization.

### Theorem A.1.

The likelihood function *L*(*θ_h_, θ_c_*, *β* | ***Z**^h^*, ***Z**^t^*) (*where θ_h_ and β are fixed*) *attains a unique global maximum* 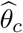 *on the unit interval* 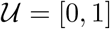 *when*

1. *the likelihood of θ_h_, β given the data from the healthy sample must be non-zero, i.e*.,

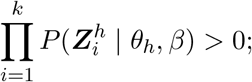
2. *the subset*

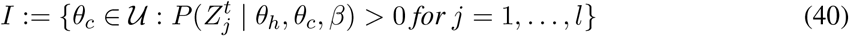

*is connected and non-empty;*
3. *the purity is greater than* 0, *i.e., α* > 0 (*otherwise the ‘tumor’ sample would not contain any cancer cells*);
4. *there exists read data* 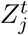 *for which the alignment probability* 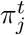 *is strictly larger than zero and p_j_* ≠ *a_j_* (*i.e., there must exist an observation that with non-zero probability stems from the locus of interest and provided information about the presence or absence of the indel of interest*).

*Proof*. The likelihood function with θh and β fixed can be written in the form

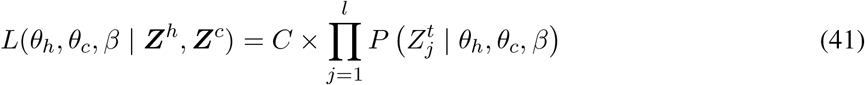

where *C* is the constant

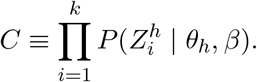

In the case that condition (*1*) is *not* met, *C* = 0. The likelihood *L*(*θ_h_, θ_c_, β* | *Z^h^, Z^c^*) equals zero for all *θ_c_* and, therefore, does not attain a unique global maximum.

Suppose condition (*1*) is met (*C* > 0). Let us consider condition (*2*). Note that *L*(*θ_h_, θ_c_, β* | ***Z**^h^, **Z**^c^*) = 0 when *θ_c_ ∉ I*, since for those *θ_c_*’s there exists an observation for which the 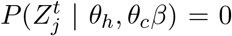. The likelihood *L* is by definition strictly larger than zero when *θ_c_* ∈ *I*. Since the function in eq. (41) is an *l*-th order polynomial and, therefore, continuous, it must attain a global maximum on the interval *I*.

Suppose condition (*2*) is met. The point 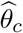 is a maximum of *L*(*θ_h_*, ·, · | ***Z**^h^, **Z**^c^*) if and only if it is a maximum of the loglikelihood function

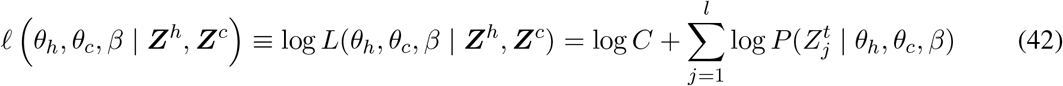

(with *θ_h_, β* fixed and *θ_c_ ∈ I*) since the logarithm is a monotonic transform. (Note that *ℓ* is only defined on the subset *I*). The second order derivative of the loglikelihood with respect to *θ_c_* is found to be

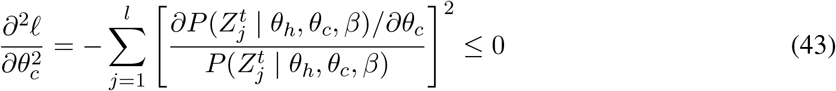

indicating that the loglikelihood function is concave. Note that it is strictly concave, i.e., 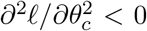, iff there exists an observation 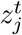 for which

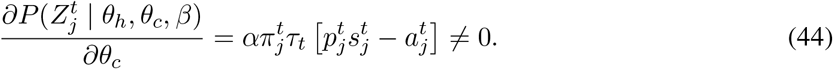

This inequality holds only when 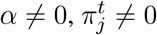 and 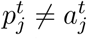, which constitutes conditions (*3*) and (*4*).

Suppose *I* is the non-empty closed set [*a, b*] on the unit interval. Since the loglikelihood is strictly concave when conditions (*3*) and (*4*) are met, it attains a unique global maximum 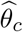 on *I*. Because the logarithm is a monotonic transformation, 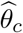 must be a unique global maximum of the likelihood function as well.

A similar reasoning holds when *I* is open or half-open. The maximum must lie on the interior of *I*, since the likelihood function is zero for those endpoints not in *I*. For example, when *I* is the open interval (*a, b*), then *L*(*θ_h_, a, β* | ***Z**^h^, **Z**^c^*) = *L*(*θ_h_, b, β* | ***Z**^h^, **Z**^c^*) = 0 while *L*(*θ_h_, θ_c_, β* | ***Z**^h^, **Z**^c^*) is strictly positive on I. The loglikelihood function is under conditions (*3*) and (*4*) strictly concave on *I*, therefore, the likelihood function attains a unique global maximum.?

## B Supplementary Figures

**Figure S1:**
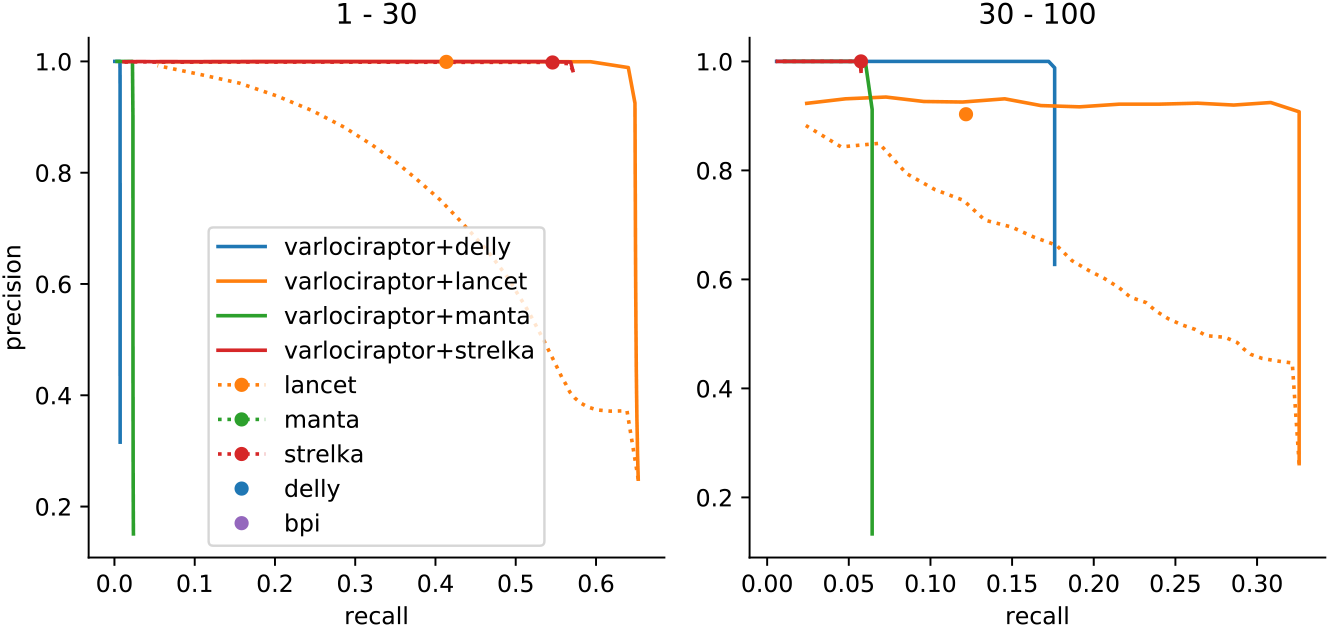
Recall and precision for calling somatic insertions. Results are grouped by deletion length, denoted as interval at the top of the plot. For our approach (Varlociraptor+*) curves are plotted by scanner over the posterior probability for having a somatic variant. For other callers that provide a score to scan over (p-value for Lancet) we plot a dotted line. Ad-hoc results are shown as single dots. Results are shown if the prediction of the caller did provide at least 10 calls. The sharp curves for our approach reflect the favorable property of having a strong separation between the probabilities of true and false positives, see Figure 4.

**Figure S2:**
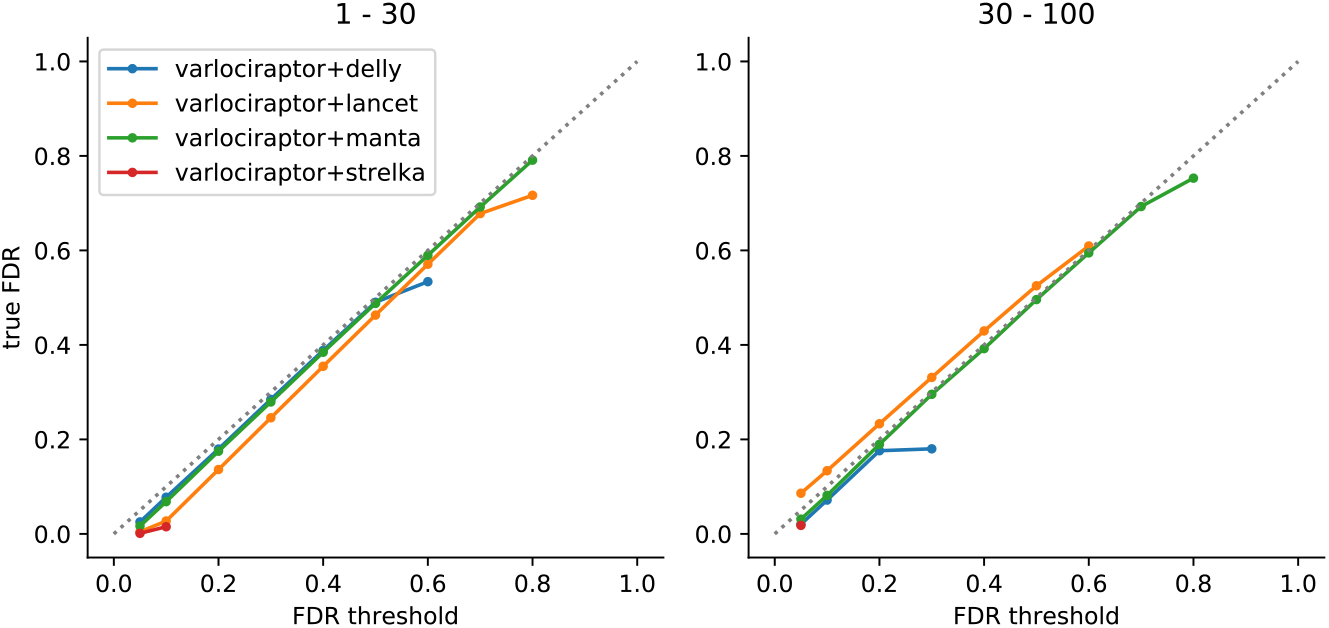
FDR control for somatic insertions. Results are grouped by deletion length, denoted as interval at the top of the plot. The axes denote the desired FDR, provided by the user as input (x-axis), and the true achieved FDR (y-axis). A perfect FDR control would keep the curve exactly on the dashed diagonal. Below the diagonal, the control is conservative. Above the diagonal, the FDR would be underestimated. Importantly, points below the diagonal mean that the true FDR is smaller than the threshold provided, which means that FDR control is still established; in this sense, points below the diagonal are preferable over points above the diagonal.

**Figure S3:**
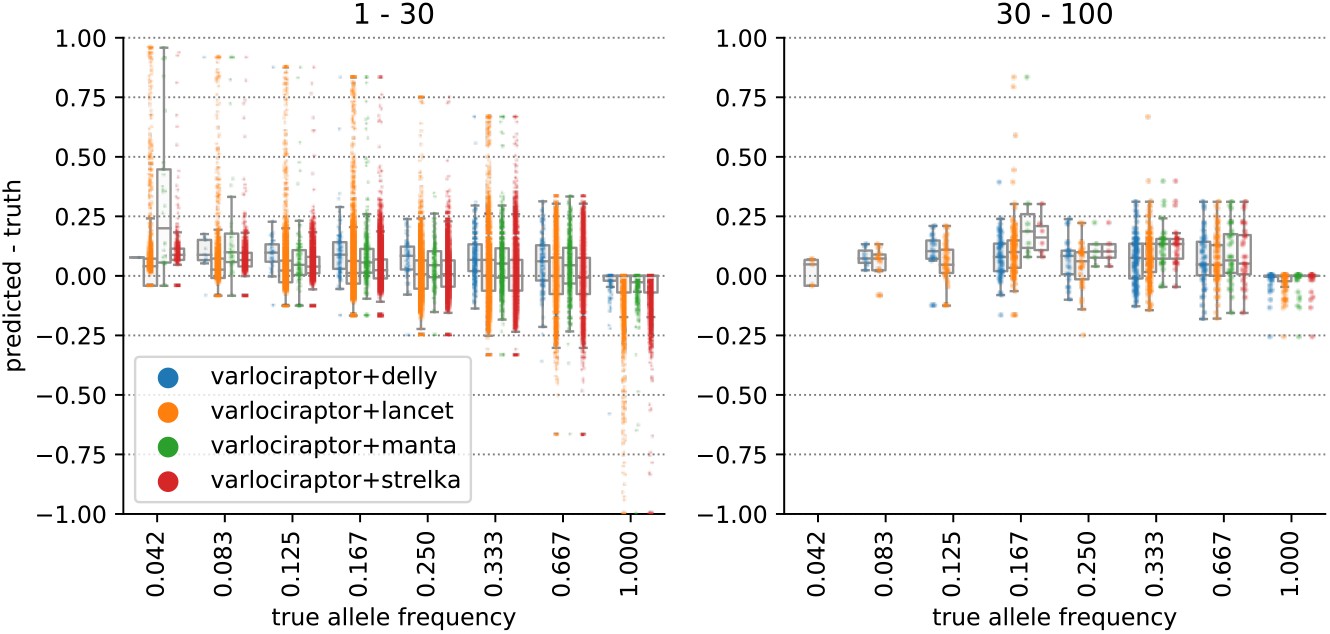
Allele frequency estimation for somatic insertions. Results are grouped by deletion length, denoted as interval at the top of the plot. The horizontal axis shows the true allele frequency, the vertical axis shows the error between predicted allele frequency and truth.

**Figure S4:**
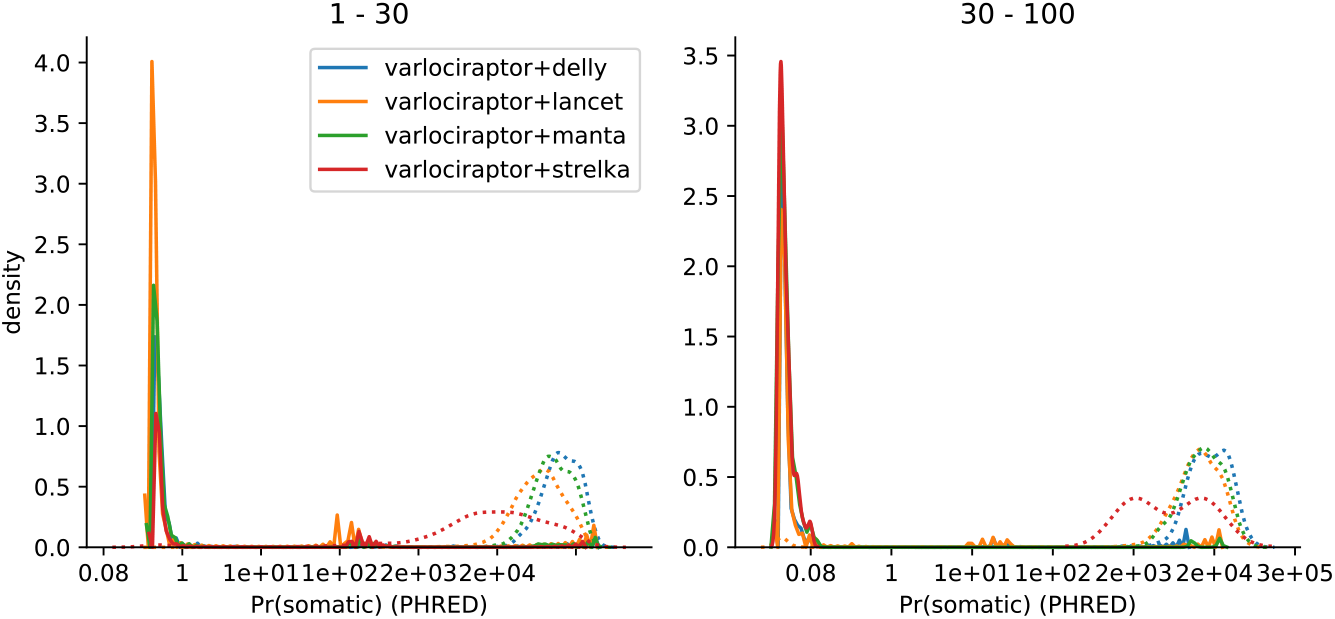
Posterior probability distributions for somatic insertions. Results are grouped by deletion length, denoted as interval at the top of the plot. The x-axis indicates the (PHRED-scaled) probability, and the y-axis indicates relative amounts of calls with this probability. The distributions of posteriors for true positive calls are shown as solid lines, the distributions of posteriors for false positive calls are shown as dotted lines.

**Figure S5:**
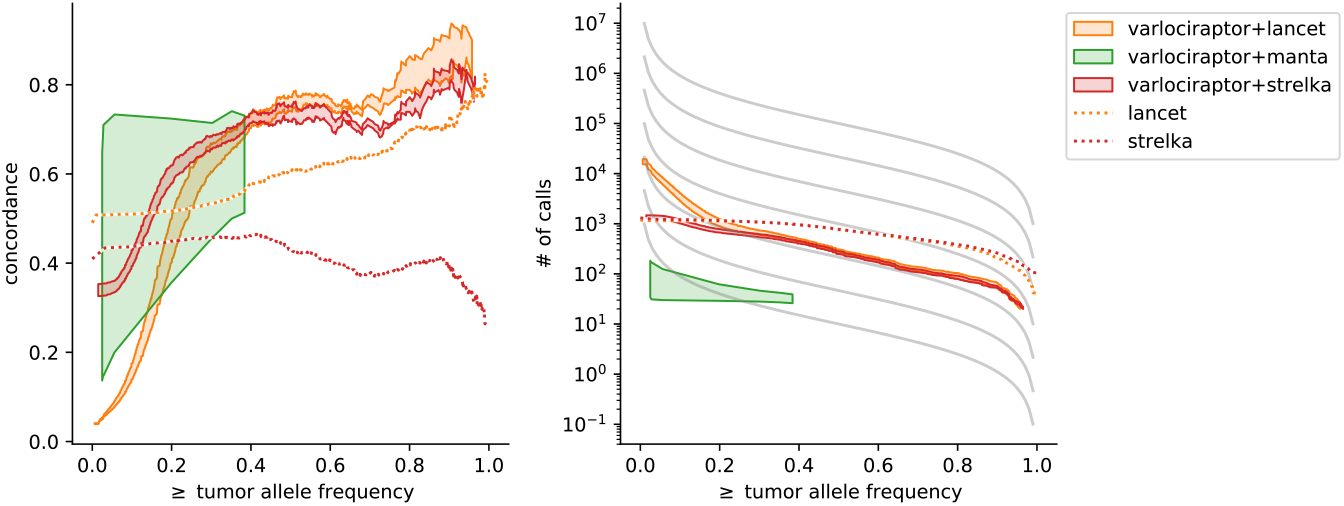
Concordance of somatic insertions on real data. For Varlociraptor, the interval between all calls with a posterior probability of at least 0.9 and at least 0.99 is shown as shaded area. Left: Concordance vs. minimum allele frequency. Right: Number of calls vs. minimum allele frequency. Grey lines depict the theoretical expectation according to Williams et al. [Williams et al., 2016]

**Figure S6:**
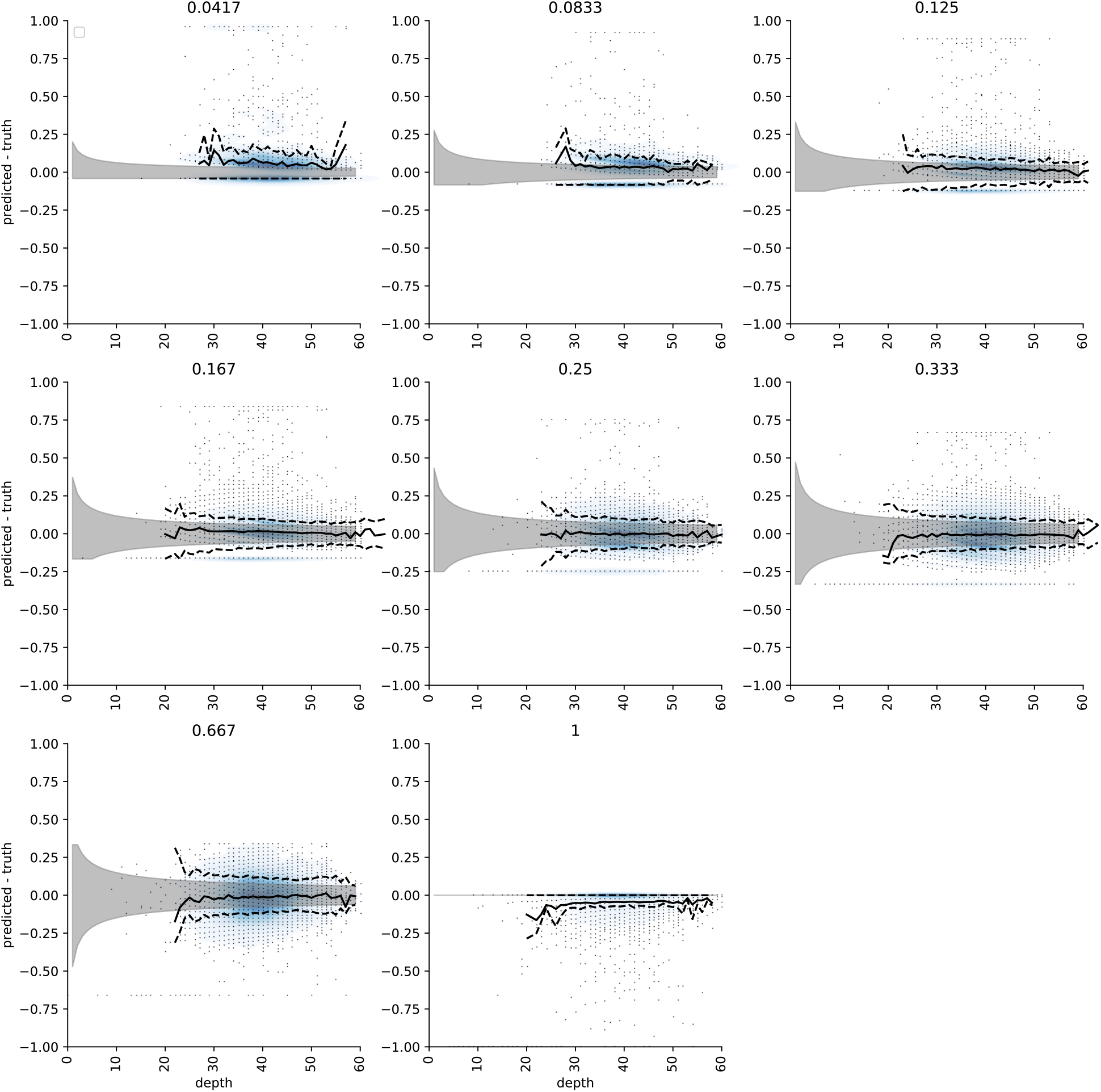
Allele frequency estimation error for somatic deletions compared to sequencing depth. Each plot shows the error (predicted - truth) for a particular true allele frequency (shown above the plot). Dots represent individual predictions, the blue shading shows a corresponding density estimate. The black line shows the mean, the dashed lines depict the standard deviation. The grey area represents the theoretically expected sampling error in an experiment with no further artifacts or biases (the theoretical optimum).

**Figure S7:**
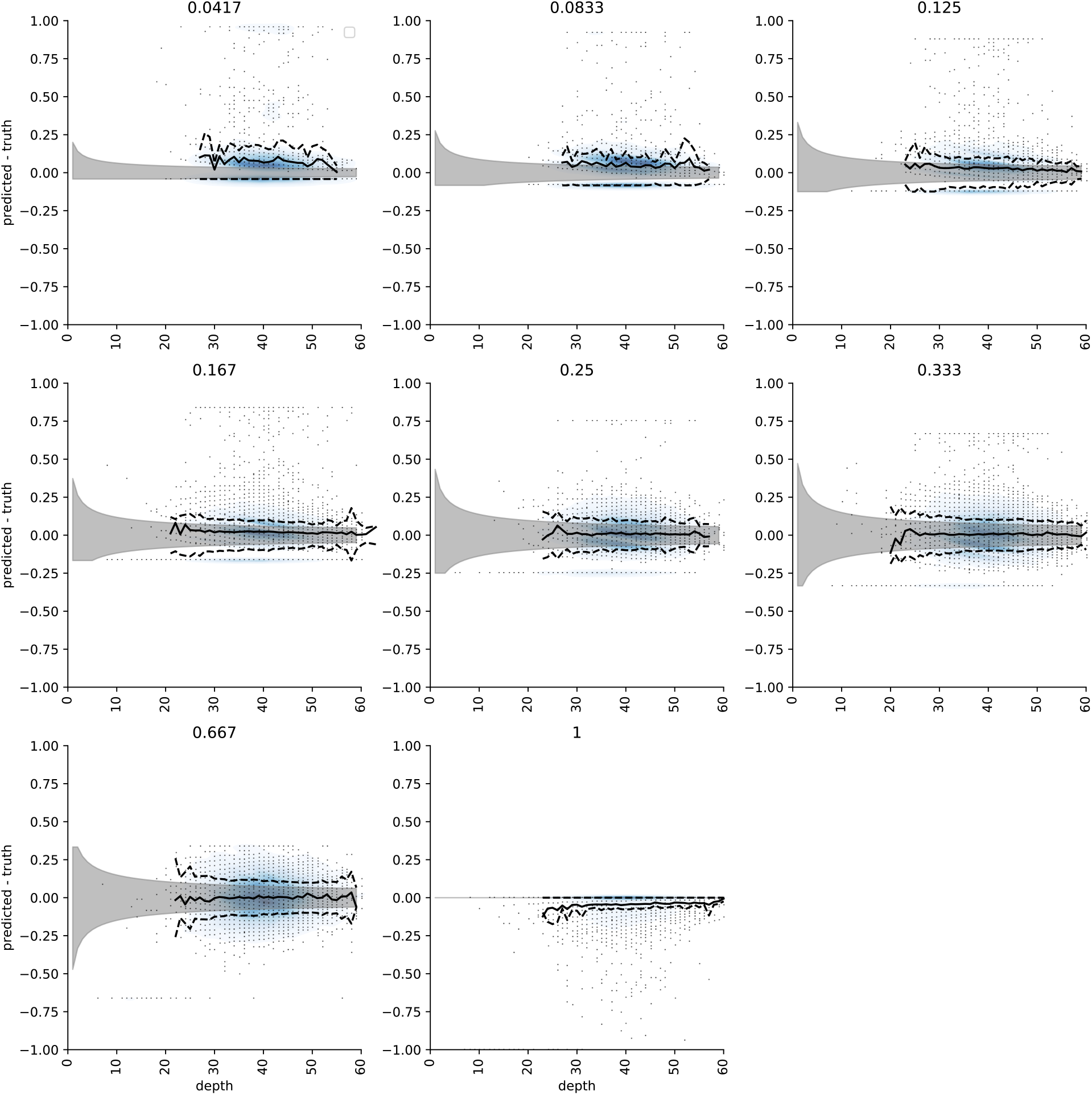
Allele frequency estimation error for somatic insertions compared to sequencing depth. Each plot shows the error (predicted - truth) for a particular true allele frequency (shown above the plot). Dots represent individual predictions, the blue shading shows a corresponding density estimate. The black line shows the mean, the dashed lines depict the standard deviation. The grey area represents the theoretically expected sampling error in an experiment with no further artifacts or biases (the theoretical optimum).

**Figure S8:**
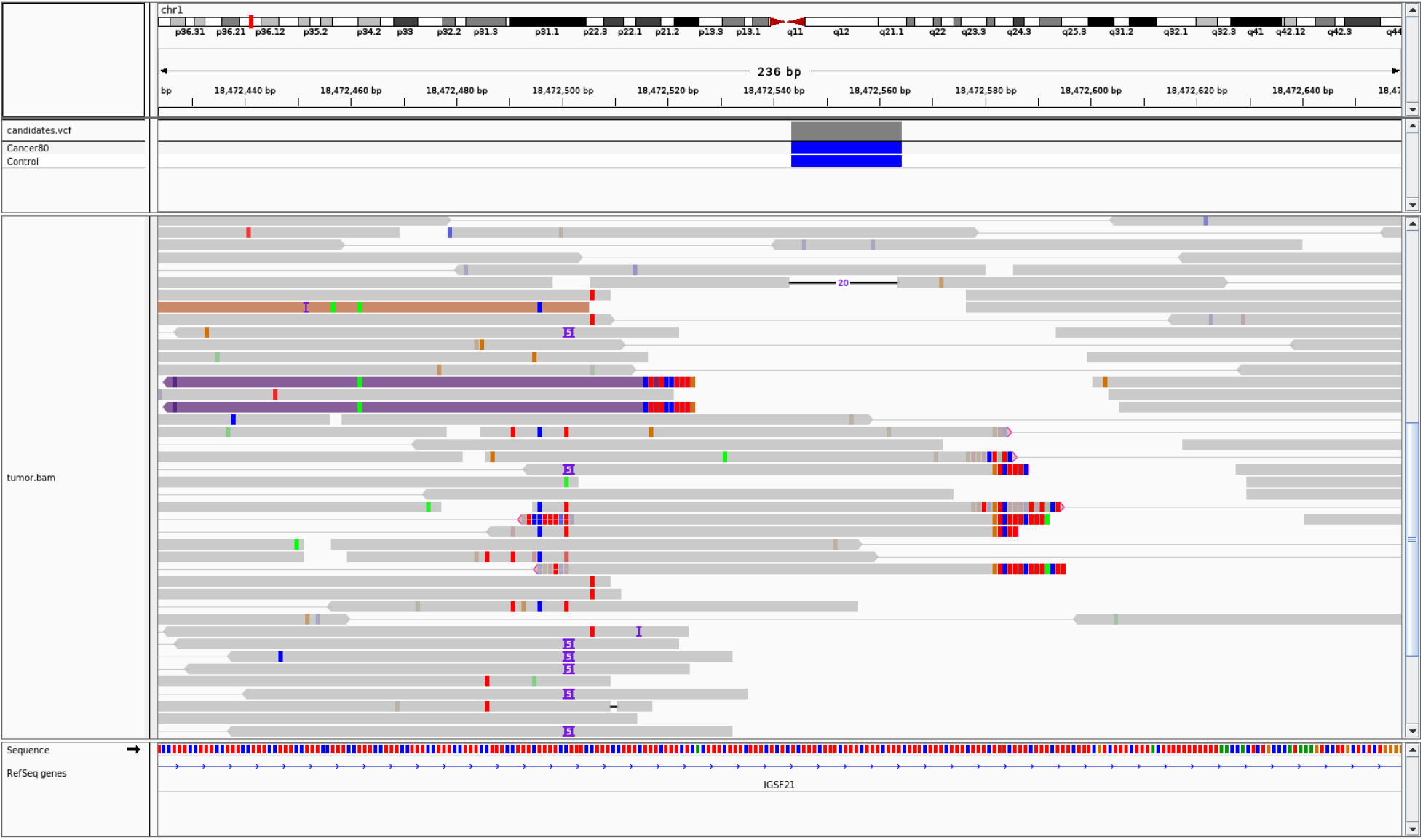
Example of a variant in a repetetive region that causes misplaced softclips, highlighting the need for a realignment against the variant allele (taken from our simulated dataset (see section 2.4). The clipped alignments (shown as mismatches at the read ends) should instead have a 20 bp deletion as the read at the top. Visualization was performed with IGV [Robinson et al., 2017].

## Footnotes

1 https://bioconda.github.io

2 Here, we refer to insertions and deletions of all possible sizes, ranging from 1 to thousands of base pairs.

3 https://cancer.sanger.ac.uk/cosmic; Release v88, data retrieved on 20th of March, 2019.

4 Here and in the following, *sensitivity or recall* denotes the ratio of true indels discovered over the overall amount of true indels, whereas *precision* denotes the ratio of true indels discovered over the overall amount of indels discovered. *False discovery rate (FDR)* is the ratio of mistaken discoveries over the overall amount of indels discovered (so precision = 1 - FDR).

5 https://samtools.github.io/hts-specs/SAMv1.pdf

6 https://samtools.github.io/hts-specs/CRAMv3.pdf

7 https://samtools.github.io/hts-specs/VCFv4.3.pdf

8 At the time of writing, the Varlociraptor implementation supports SNVs, insertions and deletions. However, the model presented here is agnostic of the variant type and we are actively working on adding support for all other types of variants in Varlociraptor. Therefore, while we are writing about indels in the following, keep in mind that the presented model can straightforwardly be applied to other variant types. Similarly, Varlociraptor currently supports single-end or paired-end short reads, while the model itself is agnostic of the sequencing protocol and technology. The implementation will be extended in the future.

9 https://github.com/SciLifeLab/Sarek

10 Note that we distinguish between the tumor sample, which is a mixture of healthy and cancer cells, and the cancer cells themselves. When referring to the latter, we use the subscript *c*, for the former, we use the subscript *t*.

11 https://doi.org/10.5281/zenodo.3361700

12 https://ega-archive.org/datasets/EGAD00001002142

13 https://github.com/hartwigmedical/hmftools/tree/master/break-point-inspector

14 https://broadinstitute.github.io/picard

15 https://github.com/rust-bio/rust-bio-tools

16 Lancet’s p-values are only one component of the ad-hoc filtering procedure performed by the tool, which relies on multiple scores. This explains while filtering based on the p-values alone (dotted orange line) yields suboptimal performance compared to the ad-hoc calls provided by Lancet (orange dot).

17 https://varlociraptor.github.io

18 https://nanoporetech.com

19 https://www.pacb.com/smrt-science/smrt-sequencing

20 https://varlociraptor.github.io/docs/calling#generic-variant-calling

21 Base qualities are reported along with read alignments in BAM files, see https://samtools.github.io/hts-specs/SAMv1.pdf.

22 Varlociraptor uses *h* = 64 by default.

