## Supplementary material for "Enhancing sensitivity and controlling false discovery rate in somatic indel discovery": Snakemake Report

Loading Snakemake Report...

Please enable Javascript in your browser to see this report.

Loading 6.0 MB. For large reports, this can take a while.

Snakemake Report

- Tue Aug 6 17:56:47 2019 CET
- Snakemake 5.5.3+6.g668fec20.dirty

- Workflow (current)
- Statistics
- Configuration
- Rules

###### Results

- Allele Frequency Estimation
- Concordance
- FDR Control
- Precision and Recall
- Score Distribution

### Workflow

Detailed software versions can be found under Rules.

### Results

###### Allele Frequency Estimation

| File | Size | Description | Job properties |
| --- | --- | --- | --- |
| simulated-bwa.DEL.svg | 328.3 kB | Allele frequency estimation error vs. true allele frequency on simulated data. | Rule \| plot\_allelefreq; Wildcards \| run=simulated-bwa, vartype=DEL; Params \| varlociraptor\_callers=['delly', 'lancet', 'manta', 'strelka'], len\_ranges=[[1, 30], [30, 50], [50, 100], [100, 250]] |
| simulated-bwa.DEL.svg | 1.9 MB | Allele frequency estimation error vs. true allele frequency on simulated data. The dashed lines depict the standard deviation, solid line depicts the mean. The grey area shows the standard deviation of a corresponding binomial sampling, which marks the maximum theoretically achievable accuracy. | Rule \| plot\_allelefreq\_scatter; Wildcards \| run=simulated-bwa, vartype=DEL; Params \| depth\_ranges=[[1, 20], [20, 40]], callers=['delly', 'lancet', 'manta', 'strelka'] |
| simulated-bwa.INS.svg | 185.2 kB | Allele frequency estimation error vs. true allele frequency on simulated data. | Rule \| plot\_allelefreq; Wildcards \| run=simulated-bwa, vartype=INS; Params \| varlociraptor\_callers=['delly', 'lancet', 'manta', 'strelka'], len\_ranges=[[1, 30], [30, 100]] |
| simulated-bwa.INS.svg | 2.1 MB | Allele frequency estimation error vs. true allele frequency on simulated data. The dashed lines depict the standard deviation, solid line depicts the mean. The grey area shows the standard deviation of a corresponding binomial sampling, which marks the maximum theoretically achievable accuracy. | Rule \| plot\_allelefreq\_scatter; Wildcards \| run=simulated-bwa, vartype=INS; Params \| depth\_ranges=[[1, 10], [10, 20], [20, 40], [40, 100]], callers=['delly', 'lancet', 'manta', 'strelka'] |

###### Concordance

| File | Size | Description | Job properties |
| --- | --- | --- | --- |
| colo1.DEL.concordance.svg | 563.4 kB | For concordance analysis, we ran varlociraptor and the other callers on four independent sequencings of the same tumor/normal pair. Called variants were matched against each other, yielding a graph. Connected components in this graph of more than one node we counted as concordant calls. Left plot shows concordance vs minimum allele frequency. Right plot shows number of calls vs minimum allele frequency. For Varlociraptor, results are shown as area between a minimum posterior probability (for somatic in tumor) of 0.9 and 0.98. For all other callers, results are shown as dotted lines (obtained with default parameters). The grey lines depict the theoretical expectation according to the model of neutral tumor evolution by Williams et al. See paper for details. | Rule \| plot\_concordance; Wildcards \| id=colo1, vartype=DEL; Params \| callers=['delly', 'lancet', 'manta', 'strelka', 'bpi'] |
| colo1.INS.concordance.svg | 428.9 kB | For concordance analysis, we ran varlociraptor and the other callers on four independent sequencings of the same tumor/normal pair. Called variants were matched against each other, yielding a graph. Connected components in this graph of more than one node we counted as concordant calls. Left plot shows concordance vs minimum allele frequency. Right plot shows number of calls vs minimum allele frequency. For Varlociraptor, results are shown as area between a minimum posterior probability (for somatic in tumor) of 0.9 and 0.98. For all other callers, results are shown as dotted lines (obtained with default parameters). The grey lines depict the theoretical expectation according to the model of neutral tumor evolution by Williams et al. See paper for details. | Rule \| plot\_concordance; Wildcards \| id=colo1, vartype=INS; Params \| callers=['delly', 'lancet', 'manta', 'strelka', 'bpi'] |

###### FDR Control

| File | Size | Description | Job properties |
| --- | --- | --- | --- |
| simulated-bwa.DEL.svg | 77.8 kB | Performance of FDR control on simulated data. Horizontal axis shows the applied FDR threshold, vertical axis shows the true FDR of the selected set of variants. Perfect FDR control would result in a line directly on the diagonal. Conservative control would lie in the area below the diagonal. Anything above the diagonal would be overly optimistic. | Rule \| plot\_fdr; Wildcards \| run=simulated-bwa, vartype=DEL; Params \| callers=['delly', 'lancet', 'manta', 'strelka'], purity=0.75, len\_ranges=[[1, 30], [30, 50], [50, 100], [100, 250]], fdrs=[1e-05, 0.0001, 0.001, 0.01, 0.05, 0.1, 0.2, 0.3, 0.4, 0.5, 0.6, 0.7, 0.8, 0.9] |
| simulated-bwa.INS.svg | 52.1 kB | Performance of FDR control on simulated data. Horizontal axis shows the applied FDR threshold, vertical axis shows the true FDR of the selected set of variants. Perfect FDR control would result in a line directly on the diagonal. Conservative control would lie in the area below the diagonal. Anything above the diagonal would be overly optimistic. | Rule \| plot\_fdr; Wildcards \| run=simulated-bwa, vartype=INS; Params \| callers=['delly', 'lancet', 'manta', 'strelka'], purity=0.75, len\_ranges=[[1, 30], [30, 100]], fdrs=[1e-05, 0.0001, 0.001, 0.01, 0.05, 0.1, 0.2, 0.3, 0.4, 0.5, 0.6, 0.7, 0.8, 0.9] |

###### Precision and Recall

| File | Size | Description | Job properties |
| --- | --- | --- | --- |
| simulated-bwa.DEL.svg | 100.9 kB | Precision and recall on simulated data. | Rule \| plot\_precision\_recall; Wildcards \| run=simulated-bwa, vartype=DEL; Params \| varlociraptor\_callers=['delly', 'lancet', 'manta', 'strelka'], default\_callers=['lancet', 'manta', 'strelka'], adhoc\_callers=['delly', 'lancet', 'manta', 'strelka', 'bpi'], len\_ranges=[[1, 30], [30, 50], [50, 100], [100, 250]] |
| simulated-bwa.INS.svg | 63.0 kB | Precision and recall on simulated data. | Rule \| plot\_precision\_recall; Wildcards \| run=simulated-bwa, vartype=INS; Params \| varlociraptor\_callers=['delly', 'lancet', 'manta', 'strelka'], default\_callers=['lancet', 'manta', 'strelka'], adhoc\_callers=['delly', 'lancet', 'manta', 'strelka', 'bpi'], len\_ranges=[[1, 30], [30, 100]] |

###### Score Distribution

| File | Size | Description | Job properties |
| --- | --- | --- | --- |
| simulated-bwa.DEL.svg | 114.6 kB | Distribution of true and false positives depending on posterior probability for somatic variant in the tumor as given by varlociraptor. Desirable is a good separation between the two distributions. | Rule \| plot\_score\_dist; Wildcards \| run=simulated-bwa, vartype=DEL; Params \| varlociraptor\_callers=['delly', 'lancet', 'manta', 'strelka'], len\_ranges=[[1, 30], [30, 50], [50, 100], [100, 250]] |
| simulated-bwa.INS.svg | 79.1 kB | Distribution of true and false positives depending on posterior probability for somatic variant in the tumor as given by varlociraptor. Desirable is a good separation between the two distributions. | Rule \| plot\_score\_dist; Wildcards \| run=simulated-bwa, vartype=INS; Params \| varlociraptor\_callers=['delly', 'lancet', 'manta', 'strelka'], len\_ranges=[[1, 30], [30, 100]] |

### Statistics

If the workflow has been executed in cluster/cloud, runtimes include the waiting time in the queue.

### Configuration

| File | Code |
| --- | --- |
| config.yaml | ``` 1 2 3 4 5 6 7 8 9 10 11 12 13 14 15 16 17 18 19 20 21 22 23 24 25 26 27 28 29 30 31 32 33 34 35 36 37 38 39 40 41 42 43 44 45 46 47 48 49 50 51 52 53 54 55 56 57 58 59 60 61 62 63 64 65 66 67 68 69 70 71 72 73 74 75 76 77 78 79 80 81 82 83 84 85 86 87 88 89 90 91 92 93 94 95 96 97 98 99 100 101 102 103 104 105 106 107 108 109 110 111 112 113 114 115 116 117 118 119 120 121 122 123 124 125 126 127 128 129 130 131 132 133 134 135 136 137 138 139 140 141 142 143 144 145 146 147 148 149 150 151 152 153 154 155 156 157 158 159 160 161 162 163 164 165 166 167 168 169 170 171 172 173 174 175 176 177 178 179 180 181 182 183 184 185 ``` \| ``` gdc-manifest: gdc_manifest_20171012_060907.txt runs: simulated-bwa: mapper: bwa dataset: simulated ref: hg18 purity: 0.75 COLO_829-GSC: dataset: COLO_829-GSC mapper: bwa ref: hg38 purity: 1.0 COLO_829-EBI: dataset: COLO_829-EBI mapper: bwa ref: hg38 purity: 1.0 COLO_829-TGen: dataset: COLO_829-TGen mapper: bwa ref: hg38 purity: 1.0 COLO_829-Ill: dataset: COLO_829-Ill mapper: bwa ref: hg38 purity: 1.0 datasets: simulated: tumor: bam: ../data/hiseq.Cancer80.wholegenome.bwamemm.cat.sorted.smtag.bam name: Cancer80 normal: bam: ../data/hiseq.Control.wholegenome.cov30.bwamemm.sorted.smtag.bam name: Control isize: mean: 312 sd: 15 truth: ../data/simulated-truth.indels.vcf COLO_829-GSC: tumor: bam: ../data/EGAD00001002142/EGAD00001002142/EGAZ00001226259_COLO_829_BCGSC_BCGSCPipe.bam name: Cancer normal: bam: ../data/EGAD00001002142/EGAD00001002142/EGAZ00001226245_COLO_829BL_BCGSC_BCGSCPipe.bam name: Control case: COLO_829 COLO_829-EBI: tumor: bam: ../data/EGAD00001002142/EGAD00001002142/EGAZ00001226269_COLO_829_EPleasance_BCGSCPipe.bam name: Cancer normal: bam: ../data/EGAD00001002142/EGAD00001002142/EGAZ00001226254_COLO_829BL_EPleasance_BCGSCPipe.bam name: Control case: COLO_829 COLO_829-Ill: tumor: bam: ../data/EGAD00001002142/EGAD00001002142/EGAZ00001226270_COLO_829_Illumina_BCGSCPipe.bam name: Cancer normal: bam: ../data/EGAD00001002142/EGAD00001002142/EGAZ00001226256_COLO_829BL_Illumina_BCGSCPipe.bam name: Control case: COLO_829 COLO_829-TGen: tumor: bam: ../data/EGAD00001002142/EGAD00001002142/EGAZ00001229972_COLO_829_TGEN_BCGSCPipe.bam name: Cancer normal: bam: ../data/EGAD00001002142/EGAD00001002142/EGAZ00001226249_COLO_829BL_TGEN_BCGSCPipe.bam name: Control case: COLO_829 ref: hg18: fasta: ../data/hg18.fasta date: 2006-03 chrom_prefix: chr hg38: fasta: ../data/Homo_sapiens.GRCh38.dna.primary_assembly.fa date: 2013-12 chrom_prefix: "" exons: resources/hg38.exons.bed caller: delly: params: "" adhoc: true genotypes: true info: - END - SOMATIC - SVLEN - SVTYPE #pindel: # params: "-M 1 -B 10000 -H 10 -x 4 -I false -a 2" lancet: params: "" score: QUAL invert: true adhoc: true genotypes: true info: - SOMATIC - SVLEN - SVTYPE manta: params: "" score: SOMATICSCORE invert: true adhoc: true useraw: true info: - SOMATIC - SOMATICSCORE - SVLEN - SVTYPE strelka: params: "" score: QSI invert: true adhoc: true info: - SOMATIC - QSI bpi: params: "" adhoc: true info: - SOMATIC - SVLEN - SVTYPE varlociraptor: params: "--indel-window 64 --omit-snvs" blacklist: - bpi score: PROB_SOMATIC_TUMOR genotypes: false info: - PROB_SOMATIC_TUMOR - PROB_GERMLINE_HET - PROB_GERMLINE_HOM - PROB_ABSENT - PROB_SOMATIC_NORMAL - PROB_ARTIFACT - SVLEN fmt: - AF - DP vcf-match-params: "--max-dist 50 --max-len-diff 50" len-ranges: DEL: - [1, 30] - [30, 50] - [50, 100] - [100, 250] INS: - [1, 30] - [30, 100] depth-ranges: DEL: - [1, 20] - [20, 40] INS: - [1, 10] - [10, 20] - [20, 40] - [40, 100] plots: concordance: colo1: - COLO_829-GSC - COLO_829-Ill - COLO_829-TGen - COLO_829-EBI known-truth: - simulated-bwa ``` |

### Rules

| Rule | Jobs | Output | Singularity | Conda environment | Code |
| --- | --- | --- | --- | --- | --- |
| plot\_precision\_recall | 2 | - plots/precision-recall/simulated-bwa.INS.svg - plots/precision-recall/simulated-bwa.DEL.svg |  | - python =3.6 - pandas =0.23 - matplotlib =3.0 - seaborn =0.9.0 - pysam =0.13.0 - svgutils =0.2 - pybedtools =0.7.10 - networkx =2.2 | ``` 1 2 3 4 5 6 7 8 9 10 11 12 13 14 15 16 17 18 19 20 21 22 23 24 25 26 27 28 29 30 31 32 33 34 35 36 37 38 39 40 41 42 43 44 45 46 47 48 49 50 51 52 53 54 55 56 57 58 59 60 61 62 63 64 65 66 67 68 69 70 71 72 73 74 75 76 77 78 79 80 81 82 83 84 85 86 87 88 89 90 91 92 93 94 95 96 97 98 99 100 101 102 103 104 105 106 107 108 109 110 111 112 113 114 115 116 117 ``` \| ``` from itertools import product import matplotlib matplotlib.use("agg") from matplotlib import pyplot as plt import seaborn as sns import pandas as pd import common import numpy as np import math from matplotlib.lines import Line2D MIN_CALLS = 10 vartype = snakemake.wildcards.vartype colors = common.get_colors(snakemake.config) def props(callers): return product(callers, snakemake.params.len_ranges) def plot_len_range(minlen, maxlen): truth = common.load_variants( snakemake.input.truth, minlen, maxlen, vartype=vartype) def plot(calls, label, color, line=True, style="-", invert=False, markersize=4): calls = pd.read_table(calls, index_col=0) if len(calls) < 10: return if line: thresholds = calls.score.quantile(np.linspace(0.0, 1.0, 50)) precision = [] recall = [] for t in thresholds: if invert: c = calls[calls.score >= t] else: c = calls[calls.score <= t] p = common.precision(c) r = common.recall(c, truth) print(label, t, c.shape[0], p, r) if len(c) < 10: print("skipping threshold: too few calls", c) continue precision.append(p) recall.append(r) if len(precision) <= 2: print("skipping curve because we have too few values") return else: precision = [common.precision(calls)] recall = [common.recall(calls, truth)] style = "." print(label, calls.shape[0], precision, recall) plt.plot( recall, precision, style, color=color, label=label, markersize=markersize) handles = [] for calls, (caller, len_range) in zip(snakemake.input.varlociraptor_calls, props(snakemake.params.varlociraptor_callers)): if len_range[0] != minlen and len_range[1] != maxlen: continue label = "varlociraptor+{}".format(caller) plot(calls, label, colors[caller]) handles.append(Line2D([0], [0], color=colors[caller], label=label)) for calls, (caller, len_range) in zip(snakemake.input.default_calls, props(snakemake.params.default_callers)): if len_range[0] != minlen and len_range[1] != maxlen: continue color = colors[caller] plot( calls, caller, color, style=":", invert=snakemake.config["caller"][caller].get("invert", False)) if caller in snakemake.params.adhoc_callers: handles.append(Line2D([0], [0], markersize=10, markerfacecolor=color, markeredgecolor=color, color=color, label=caller, marker=".", linestyle=":")) else: handles.append(Line2D([0], [0], color=color, label=caller, linestyle=":")) for calls, (caller, len_range) in zip(snakemake.input.adhoc_calls, props(snakemake.params.adhoc_callers)): if len_range[0] != minlen and len_range[1] != maxlen: continue color = colors[caller] plot(calls, caller, color, markersize=10, line=False) if caller not in snakemake.params.default_callers: handles.append(Line2D([0], [0], markersize=10, markerfacecolor=color, markeredgecolor=color, label=caller, marker=".", lw=0)) sns.despine() ax = plt.gca() return ax, handles common.plot_ranges( snakemake.params.len_ranges, plot_len_range, xlabel="recall", ylabel="precision") plt.savefig(snakemake.output[0], bbox_inches="tight") ``` |
| svg2pdf | 14 | - plots/precision-recall/simulated-bwa.INS.pdf - plots/precision-recall/simulated-bwa.DEL.pdf - plots/fdr-control/simulated-bwa.INS.pdf - plots/fdr-control/simulated-bwa.DEL.pdf - plots/allelefreqs/simulated-bwa.INS.pdf - plots/allelefreqs/simulated-bwa.DEL.pdf - plots/score-dist/simulated-bwa.INS.pdf - plots/score-dist/simulated-bwa.DEL.pdf - plots/allelefreq-recall/simulated-bwa.INS.pdf - plots/allelefreq-recall/simulated-bwa.DEL.pdf - plots/allelefreq-scatter/simulated-bwa.INS.pdf - plots/allelefreq-scatter/simulated-bwa.DEL.pdf - plots/concordance/colo1.INS.concordance.pdf - plots/concordance/colo1.DEL.concordance.pdf |  | - cairosvg =2.1.3 | ``` 1 ``` \| ``` cairosvg {input} -o {output} ``` |
| plot\_fdr | 2 | - plots/fdr-control/simulated-bwa.INS.svg - plots/fdr-control/simulated-bwa.DEL.svg |  | - python =3.6 - pandas =0.23 - matplotlib =3.0 - seaborn =0.9.0 - pysam =0.13.0 - svgutils =0.2 - pybedtools =0.7.10 - networkx =2.2 | ``` 1 2 3 4 5 6 7 8 9 10 11 12 13 14 15 16 17 18 19 20 21 22 23 24 25 26 27 28 29 30 31 32 33 34 35 36 37 38 39 40 41 42 43 44 45 46 47 48 49 50 51 52 53 54 55 56 57 58 59 60 61 62 63 64 65 66 ``` \| ``` from itertools import product import matplotlib matplotlib.use("agg") from matplotlib import pyplot as plt import seaborn as sns import pandas as pd import common import numpy as np MIN_CALLS = 100 colors = common.get_colors(snakemake.config) props = product(snakemake.params.callers, snakemake.params.len_ranges, snakemake.params.fdrs) calls = [] for _calls, (caller, len_range, fdr) in zip(snakemake.input.varlociraptor_calls, props): calls.append({"caller": caller, "len_range": len_range, "fdr": float(fdr), "calls": _calls}) calls = pd.DataFrame(calls) calls = calls.set_index("caller", drop=False) def plot_len_range(minlen, maxlen): def plot(caller): color = colors[caller] label = "varlociraptor+{}".format(caller) fdrs = [] alphas = [] calls_ = calls.loc[caller] calls_ = calls_[calls_["len_range"].map(lambda r: r == [minlen, maxlen])] calls_ = calls_.sort_values("fdr") for e in calls_.itertuples(): c = pd.read_table(e.calls) n = c.shape[0] if n < MIN_CALLS: continue true_fdr = 1.0 - common.precision(c) if fdrs and fdrs[-1] == true_fdr: continue fdrs.append(true_fdr) alphas.append(e.fdr) plt.plot(alphas, fdrs, ".-", color=color, label=label) for caller in calls.index.unique(): plot(caller) plt.plot([0, 1], [0, 1], ":", color="grey") sns.despine() ax = plt.gca() handles, _ = ax.get_legend_handles_labels() return ax, handles common.plot_ranges( snakemake.params.len_ranges, plot_len_range, xlabel="FDR threshold", ylabel="true FDR") plt.savefig(snakemake.output[0], bbox_inches="tight") ``` |
| plot\_allelefreq | 2 | - plots/allelefreqs/simulated-bwa.INS.svg - plots/allelefreqs/simulated-bwa.DEL.svg |  | - python =3.6 - pandas =0.23 - matplotlib =3.0 - seaborn =0.9.0 - pysam =0.13.0 - svgutils =0.2 - pybedtools =0.7.10 - networkx =2.2 | ``` 1 2 3 4 5 6 7 8 9 10 11 12 13 14 15 16 17 18 19 20 21 22 23 24 25 26 27 28 29 30 31 32 33 34 35 36 37 38 39 40 41 42 43 44 45 46 47 48 49 50 51 52 53 54 55 56 57 58 59 60 61 62 63 64 65 66 67 68 69 70 71 72 73 74 75 76 77 78 79 80 81 82 83 84 85 86 ``` \| ``` from itertools import product import math import matplotlib matplotlib.use("agg") from matplotlib import pyplot as plt import seaborn as sns import pandas as pd import common import numpy as np MIN_CALLS = 10 vartype = snakemake.wildcards.vartype colors = common.get_colors(snakemake.config) truth = common.load_variants(snakemake.input.truth, vartype=vartype) def props(callers): return product(callers, snakemake.params.len_ranges) def plot_len_range(minlen, maxlen): def plot(calls, colors): calls = calls[calls.is_tp] true_af = truth.loc[calls.MATCHING].reset_index().TAF calls = calls.reset_index() calls["error"] = calls.CASE_AF - true_af if calls.empty: return calls["true_af"] = true_af true_af = pd.Series(calls["true_af"].unique()).sort_values() # standard deviation when sampling in binomial process from allele freq # this is the expected sampling error within the correctly mapped fragments # sd = true_af.apply(lambda af: 1 / 40 * math.sqrt(40 * af * (1 - af))) # x = np.arange(len(true_af)) # offsets = [-0.5, 0.5] # y_upper = np.array([v for v in sd for o in offsets]) # y_lower = np.maximum(-y_upper, [-f for f in true_af for o in offsets]) # plt.fill_between([v + o for v in x for o in offsets], y_lower, y_upper, color="#EEEEEE", zorder=-5) calls["true_af"] = calls["true_af"].apply("{:.3f}".format) size = 1 if maxlen == 30 else 2 sns.stripplot("true_af", "error", hue="caller", data=calls, palette=colors, dodge=True, jitter=True, alpha=0.5, size=size, rasterized=True) sns.boxplot("true_af", "error", hue="caller", data=calls, color="white", fliersize=0, linewidth=1) handles, labels = plt.gca().get_legend_handles_labels() n = len(calls.caller.unique()) plt.ylim((-1,1)) plt.grid(axis="y", linestyle=":", color="grey") sns.despine() plt.xticks(rotation="vertical") ax = plt.gca() ax.legend().remove() return ax, handles[n:] all_calls, all_colors = load_calls(minlen, maxlen) return plot(all_calls, all_colors) def load_calls(minlen, maxlen): all_calls = [] all_colors = [] for calls, (caller, len_range) in zip(snakemake.input.varlociraptor_calls, props(snakemake.params.varlociraptor_callers)): if len_range[0] != minlen and len_range[1] != maxlen: continue label = "varlociraptor+{}".format(caller) calls = pd.read_table(calls) calls["caller"] = label if not calls.empty: all_calls.append(calls) all_colors.append(colors[caller]) all_calls = pd.concat(all_calls) return all_calls, all_colors common.plot_ranges( snakemake.params.len_ranges, plot_len_range, xlabel="true allele frequency", ylabel="predicted - truth") plt.savefig(snakemake.output[0], bbox_inches="tight") ``` |
| plot\_score\_dist | 2 | - plots/score-dist/simulated-bwa.INS.svg - plots/score-dist/simulated-bwa.DEL.svg |  | - python =3.6 - pandas =0.23 - matplotlib =3.0 - seaborn =0.9.0 - pysam =0.13.0 - svgutils =0.2 - pybedtools =0.7.10 - networkx =2.2 | ``` 1 2 3 4 5 6 7 8 9 10 11 12 13 14 15 16 17 18 19 20 21 22 23 24 25 26 27 28 29 30 31 32 33 34 35 36 37 38 39 40 41 42 43 44 45 46 47 ``` \| ``` from itertools import product import matplotlib matplotlib.use("agg") from matplotlib import pyplot as plt import seaborn as sns import pandas as pd import common import numpy as np import math vartype = snakemake.wildcards.vartype colors = common.get_colors(snakemake.config) def props(callers): return product(callers, snakemake.params.len_ranges) phred_to_log_factor = -0.23025850929940456 log_to_phred_factor = -4.3429448190325175 def plot_len_range(minlen, maxlen): for calls, (caller, len_range) in zip(snakemake.input.varlociraptor_calls, props(snakemake.params.varlociraptor_callers)): if len_range[0] != minlen and len_range[1] != maxlen: continue label = "varlociraptor+{}".format(caller) calls = pd.read_table(calls) calls["caller"] = label if not calls.empty: color = colors[caller] sns.kdeplot(calls[calls.is_tp].PROB_SOMATIC_TUMOR.map(np.log), color=color, label=label) sns.kdeplot(calls[~calls.is_tp].PROB_SOMATIC_TUMOR.map(np.log), color=color, linestyle=":", label="") ax = plt.gca() fmt_ticks = lambda ticks: ["{:.1g}".format(np.exp(t)) for t in ticks] ax.set_xticklabels(fmt_ticks(plt.xticks()[0])) ax.legend().remove() handles, _ = ax.get_legend_handles_labels() sns.despine() return ax, handles common.plot_ranges( snakemake.params.len_ranges, plot_len_range, xlabel="Pr(somatic) (PHRED)", ylabel="density") plt.savefig(snakemake.output[0], bbox_inches="tight") ``` |
| plot\_allelefreq\_recall | 2 | - plots/allelefreq-recall/simulated-bwa.INS.svg - plots/allelefreq-recall/simulated-bwa.DEL.svg |  | - python =3.6 - pandas =0.23 - matplotlib =3.0 - seaborn =0.9.0 - pysam =0.13.0 - svgutils =0.2 - pybedtools =0.7.10 - networkx =2.2 | ``` 1 2 3 4 5 6 7 8 9 10 11 12 13 14 15 16 17 18 19 20 21 22 23 24 25 26 27 28 29 30 31 32 33 34 35 36 37 38 39 40 41 42 43 44 45 46 47 48 49 50 51 52 53 54 55 56 57 58 59 60 61 62 63 64 65 66 67 68 69 70 71 72 73 74 75 76 77 78 79 80 81 82 83 84 85 86 87 88 89 90 91 92 93 94 95 96 ``` \| ``` from itertools import product import matplotlib matplotlib.use("agg") from matplotlib import pyplot as plt import seaborn as sns import pandas as pd import common import numpy as np import math from matplotlib.lines import Line2D MIN_CALLS = 10 vartype = snakemake.wildcards.vartype colors = common.get_colors(snakemake.config) def props(callers): return product(callers, snakemake.params.len_ranges) def plot_len_range(minlen, maxlen): truth = common.load_variants( snakemake.input.truth, minlen, maxlen, vartype=vartype) afs = pd.Series(truth.TAF.unique()).sort_values() def plot(calls, label, color, varlociraptor=True, style="-.", markersize=4): calls = pd.read_table(calls, index_col=0) if len(calls) < 10: return if varlociraptor: phred = lambda p: -10 * math.log10(p) def calc_recall(p): c = calls[calls.score <= phred(p)] return [common.recall(c, truth[truth.TAF >= af]) for af in afs] return plt.fill_between( afs, calc_recall(0.98 if maxlen > 30 else 0.99), calc_recall(0.9), color=color, label=label, alpha=0.6) else: recall = [common.recall(calls, truth[truth.TAF >= af]) for af in afs] # plot a white background first to increase visibility plt.plot(afs, recall, "-", color="white", alpha=0.8) return plt.plot( afs, recall, style, color=color, label=label)[0] handles = [] def register_handle(handle): if handle is not None: handles.append(handle) for calls, (caller, len_range) in zip(snakemake.input.varlociraptor_calls, props(snakemake.params.varlociraptor_callers)): if len_range[0] != minlen and len_range[1] != maxlen: continue label = "varlociraptor+{}".format(caller) handle = plot(calls, label, colors[caller], varlociraptor=True) register_handle(handle) #handles.append(Line2D([0], [0], color=colors[caller], label=label)) for calls, (caller, len_range) in zip(snakemake.input.adhoc_calls, props(snakemake.params.adhoc_callers)): if len_range[0] != minlen and len_range[1] != maxlen: continue color = colors[caller] handle = plot(calls, caller, color, style=":", varlociraptor=False) register_handle(handle) #handles.append(Line2D([0], [0], linestyle=":", color=color, label=caller)) sns.despine() ax = plt.gca() return ax, handles common.plot_ranges( snakemake.params.len_ranges, plot_len_range, xlabel="allele frequency", ylabel="recall") plt.savefig(snakemake.output[0], bbox_inches="tight") ``` |
| plot\_allelefreq\_scatter | 2 | - plots/allelefreq-scatter/simulated-bwa.INS.svg - plots/allelefreq-scatter/simulated-bwa.DEL.svg |  | - python =3.6 - pandas =0.23 - matplotlib =3.0 - seaborn =0.9.0 - pysam =0.13.0 - svgutils =0.2 - pybedtools =0.7.10 - networkx =2.2 | ``` 1 2 3 4 5 6 7 8 9 10 11 12 13 14 15 16 17 18 19 20 21 22 23 24 25 26 27 28 29 30 31 32 33 34 35 36 37 38 39 40 41 42 43 44 45 46 47 48 49 50 51 52 53 54 55 56 57 58 59 60 61 62 63 64 65 66 67 68 69 70 71 ``` \| ``` import math import matplotlib matplotlib.use("agg") from matplotlib import pyplot as plt import seaborn as sns import pandas as pd import common import numpy as np MIN_COUNT = 20 MAX_DEPTH = 60 vartype = snakemake.wildcards.vartype colors = common.get_colors(snakemake.config) truth = common.load_variants(snakemake.input.truth, vartype=vartype) all_calls = [] for caller, calls in zip(snakemake.params.callers, snakemake.input.calls): calls = pd.read_table(calls) calls.loc[:, "caller"] = caller all_calls.append(calls) all_calls = pd.concat(all_calls) def plot(af, _): constrain_lower = lambda error: np.maximum(error, -af) constrain_upper = lambda error: np.minimum(error, 1.0 - af) dp = all_calls["TUMOR_DP"] calls = all_calls[all_calls.is_tp] true_af = truth.loc[calls.MATCHING].reset_index().TAF calls = calls.reset_index() calls["true_af"] = true_af calls = calls[calls["true_af"] == af] calls["error"] = calls.CASE_AF - true_af sns.kdeplot(calls["TUMOR_DP"], calls["error"], cmap="Blues", n_levels=50, shade=True, alpha=0.7, shade_lowest=False) #alpha=0.5, clip=((0.0, 1.0), (0.0, af))) plt.plot(calls["TUMOR_DP"], calls["error"], ",", color="k", lw=0, alpha=1.0, rasterized=True) by_depth = calls.groupby("TUMOR_DP")["error"].describe().reset_index() by_depth["-std"] = constrain_lower(-by_depth["std"]) by_depth["std"] = constrain_upper(by_depth["std"]) by_depth = by_depth[by_depth["count"] >= MIN_COUNT] plt.plot(by_depth.TUMOR_DP, by_depth["std"], "--", color="k") plt.plot(by_depth.TUMOR_DP, by_depth["-std"], "--", color="k") plt.plot(by_depth.TUMOR_DP, by_depth["mean"], "-", color="k") depths = np.arange(0, MAX_DEPTH) # standard deviation when sampling in binomial process from allele freq # this is the expected sampling error within the correctly mapped fragments sd = np.array([1.0 / depth * math.sqrt(depth * af * (1.0 - af)) for depth in depths]) plt.fill_between(depths, constrain_lower(-sd), constrain_upper(sd), color="grey", alpha=0.5) sns.despine() plt.xticks(rotation="vertical") ax = plt.gca() ax.legend().remove() handles, labels = ax.get_legend_handles_labels() plt.ylim((-1.0, 1.0)) plt.xlim((0, MAX_DEPTH)) return ax, [] afs = [(af, af) for af in truth.TAF.sort_values().unique()] common.plot_ranges( afs, plot, "depth", "predicted - truth") plt.savefig(snakemake.output[0], bbox_inches="tight") ``` |
| plot\_concordance | 2 | - plots/concordance/colo1.INS.concordance.svg - plots/concordance/colo1.DEL.concordance.svg |  | - python =3.6 - pandas =0.23 - matplotlib =3.0 - seaborn =0.9.0 - pysam =0.13.0 - svgutils =0.2 - pybedtools =0.7.10 - networkx =2.2 | ``` 1 2 3 4 5 6 7 8 9 10 11 12 13 14 15 16 17 18 19 20 21 22 23 24 25 26 27 28 29 30 31 32 33 34 35 36 37 38 39 40 41 42 43 44 45 46 47 48 49 50 51 52 53 54 55 56 57 58 59 60 61 62 63 64 65 66 67 68 69 70 71 72 73 74 75 76 77 78 79 80 81 82 83 84 85 86 87 88 89 90 91 92 93 94 95 96 97 98 99 100 101 102 103 104 105 106 107 108 109 110 111 112 113 114 115 116 117 118 119 120 121 122 123 124 125 126 127 ``` \| ``` from itertools import product import matplotlib matplotlib.use("agg") from matplotlib import pyplot as plt import seaborn as sns import pandas as pd import common import numpy as np import math from matplotlib.lines import Line2D from matplotlib.colors import to_rgba class NotEnoughObservationsException(Exception): pass MIN_CALLS = 20 MAX_LEN = 1000 vartype = snakemake.wildcards.vartype colors = common.get_colors(snakemake.config) varlociraptor_calls_low = [pd.read_table(f) for f in snakemake.input.varlociraptor_calls_low] varlociraptor_calls_high = [pd.read_table(f) for f in snakemake.input.varlociraptor_calls_high] adhoc_calls = [pd.read_table(f) for f in snakemake.input.adhoc_calls] def expected_count(af, effective_mutation_rate): """Calculate the expected number of somatic variants greater than a given allele frequency given an effective mutation rate, according to the model of Williams et al. Nature Genetics 2016""" return effective_mutation_rate * (1.0 / af - 1.0) def expected_counts(afs, effective_mutation_rate): return [expected_count(af, effective_mutation_rate) for af in afs] def calc_concordance(calls): n = len(calls) return (calls["concordance_count"] > 1).sum() / n def plot_len_range(minlen, maxlen, yfunc=None, yscale=None, upper_bound=None): handles_varlociraptor = [] handles_adhoc = [] for i, caller in enumerate(snakemake.params.callers): def plot_calls(calls, label, color, style, calls_lower=None): def get_xy(calls, caseafs=None): svlen = calls.loc[:, calls.columns.str.startswith("SVLEN")].abs() # at least one of the calls has a valid svlen valid = ((svlen >= minlen) & (svlen <= maxlen)).sum(axis=1) >= 1 calls = calls[valid] if caseafs is None: caseafs = calls["max_case_af"].dropna().unique() y = [] _caseafs = [] for caseaf in sorted(caseafs): _calls = calls[calls["max_case_af"] >= caseaf] if upper_bound is not None: _calls = _calls[_calls["max_case_af"] <= caseaf + upper_bound] if len(_calls) < MIN_CALLS: continue _caseafs.append(caseaf) y.append(yfunc(_calls)) return _caseafs, y x, y = get_xy(calls) if not x: raise NotEnoughObservationsException() if calls_lower is not None: _, y2 = get_xy(calls_lower, caseafs=x) return plt.fill_between(x, y, y2, label=label, edgecolor=color, facecolor=to_rgba(color, alpha=0.2)) else: if style != "-": plt.plot(x, y, "-", color="white", alpha=0.8) return plt.plot(x, y, style, label=label, color=color)[0] color = colors[snakemake.params.callers[i]] try: handles_varlociraptor.append( plot_calls( varlociraptor_calls_high[i], "varlociraptor+{}".format(caller), color=color, style="-", calls_lower=varlociraptor_calls_low[i])) except NotEnoughObservationsException: # skip plot pass try: handles_adhoc.append(plot_calls(adhoc_calls[i], caller, color=color, style=":")) except NotEnoughObservationsException: # skip plot pass handles = handles_varlociraptor + handles_adhoc sns.despine() ax = plt.gca() if yscale is not None: ax.set_yscale(yscale) return ax, handles plt.figure(figsize=(10, 4)) plt.subplot(121) plot_len_range(1, MAX_LEN, yfunc=calc_concordance) plt.xlabel("$\geq$ tumor allele frequency") plt.ylabel("concordance") plt.subplot(122) for effective_mutation_rate in 10 ** np.linspace(1, 5, 7): afs = np.linspace(0.0, 1.0, 100, endpoint=False) plt.semilogy(afs, expected_counts(afs, effective_mutation_rate), "-", color="grey", alpha=0.4) ax, handles = plot_len_range(1, MAX_LEN, yfunc=lambda calls: len(calls), yscale="log") plt.xlabel("$\geq$ tumor allele frequency") plt.ylabel("# of calls") ax.legend(handles=handles, loc="upper left", bbox_to_anchor=(1.0, 1.0)) plt.tight_layout() plt.savefig(snakemake.output[0], bbox_inches="tight") ``` |
| obtain\_tp\_fp | 416 | - annotated-calls/varlociraptor-delly/simulated-bwa.INS.1-30.1.0.tsv - annotated-calls/varlociraptor-delly/simulated-bwa.INS.30-100.1.0.tsv - annotated-calls/varlociraptor-lancet/simulated-bwa.INS.1-30.1.0.tsv - annotated-calls/varlociraptor-lancet/simulated-bwa.INS.30-100.1.0.tsv - annotated-calls/varlociraptor-manta/simulated-bwa.INS.1-30.1.0.tsv - annotated-calls/varlociraptor-manta/simulated-bwa.INS.30-100.1.0.tsv - annotated-calls/varlociraptor-strelka/simulated-bwa.INS.1-30.1.0.tsv - annotated-calls/varlociraptor-strelka/simulated-bwa.INS.30-100.1.0.tsv - annotated-calls/default-lancet/simulated-bwa.INS.1-30.1.0.tsv - annotated-calls/default-lancet/simulated-bwa.INS.30-100.1.0.tsv - annotated-calls/default-manta/simulated-bwa.INS.1-30.1.0.tsv - annotated-calls/default-manta/simulated-bwa.INS.30-100.1.0.tsv - annotated-calls/default-strelka/simulated-bwa.INS.1-30.1.0.tsv - annotated-calls/default-strelka/simulated-bwa.INS.30-100.1.0.tsv - annotated-calls/adhoc-delly/simulated-bwa.INS.1-30.1.0.tsv - annotated-calls/adhoc-delly/simulated-bwa.INS.30-100.1.0.tsv - annotated-calls/adhoc-lancet/simulated-bwa.INS.1-30.1.0.tsv - annotated-calls/adhoc-lancet/simulated-bwa.INS.30-100.1.0.tsv - annotated-calls/adhoc-manta/simulated-bwa.INS.1-30.1.0.tsv - annotated-calls/adhoc-manta/simulated-bwa.INS.30-100.1.0.tsv - annotated-calls/adhoc-strelka/simulated-bwa.INS.1-30.1.0.tsv - annotated-calls/adhoc-strelka/simulated-bwa.INS.30-100.1.0.tsv - annotated-calls/adhoc-bpi/simulated-bwa.INS.1-30.1.0.tsv - annotated-calls/adhoc-bpi/simulated-bwa.INS.30-100.1.0.tsv - annotated-calls/varlociraptor-delly/simulated-bwa.DEL.1-30.1.0.tsv - annotated-calls/varlociraptor-delly/simulated-bwa.DEL.30-50.1.0.tsv - annotated-calls/varlociraptor-delly/simulated-bwa.DEL.50-100.1.0.tsv - annotated-calls/varlociraptor-delly/simulated-bwa.DEL.100-250.1.0.tsv - annotated-calls/varlociraptor-lancet/simulated-bwa.DEL.1-30.1.0.tsv - annotated-calls/varlociraptor-lancet/simulated-bwa.DEL.30-50.1.0.tsv - annotated-calls/varlociraptor-lancet/simulated-bwa.DEL.50-100.1.0.tsv - annotated-calls/varlociraptor-lancet/simulated-bwa.DEL.100-250.1.0.tsv - annotated-calls/varlociraptor-manta/simulated-bwa.DEL.1-30.1.0.tsv - annotated-calls/varlociraptor-manta/simulated-bwa.DEL.30-50.1.0.tsv - annotated-calls/varlociraptor-manta/simulated-bwa.DEL.50-100.1.0.tsv - annotated-calls/varlociraptor-manta/simulated-bwa.DEL.100-250.1.0.tsv - annotated-calls/varlociraptor-strelka/simulated-bwa.DEL.1-30.1.0.tsv - annotated-calls/varlociraptor-strelka/simulated-bwa.DEL.30-50.1.0.tsv - annotated-calls/varlociraptor-strelka/simulated-bwa.DEL.50-100.1.0.tsv - annotated-calls/varlociraptor-strelka/simulated-bwa.DEL.100-250.1.0.tsv - annotated-calls/default-lancet/simulated-bwa.DEL.1-30.1.0.tsv - annotated-calls/default-lancet/simulated-bwa.DEL.30-50.1.0.tsv - annotated-calls/default-lancet/simulated-bwa.DEL.50-100.1.0.tsv - annotated-calls/default-lancet/simulated-bwa.DEL.100-250.1.0.tsv - annotated-calls/default-manta/simulated-bwa.DEL.1-30.1.0.tsv - annotated-calls/default-manta/simulated-bwa.DEL.30-50.1.0.tsv - annotated-calls/default-manta/simulated-bwa.DEL.50-100.1.0.tsv - annotated-calls/default-manta/simulated-bwa.DEL.100-250.1.0.tsv - annotated-calls/default-strelka/simulated-bwa.DEL.1-30.1.0.tsv - annotated-calls/default-strelka/simulated-bwa.DEL.30-50.1.0.tsv - annotated-calls/default-strelka/simulated-bwa.DEL.50-100.1.0.tsv - annotated-calls/default-strelka/simulated-bwa.DEL.100-250.1.0.tsv - annotated-calls/adhoc-delly/simulated-bwa.DEL.1-30.1.0.tsv - annotated-calls/adhoc-delly/simulated-bwa.DEL.30-50.1.0.tsv - annotated-calls/adhoc-delly/simulated-bwa.DEL.50-100.1.0.tsv - annotated-calls/adhoc-delly/simulated-bwa.DEL.100-250.1.0.tsv - annotated-calls/adhoc-lancet/simulated-bwa.DEL.1-30.1.0.tsv - annotated-calls/adhoc-lancet/simulated-bwa.DEL.30-50.1.0.tsv - annotated-calls/adhoc-lancet/simulated-bwa.DEL.50-100.1.0.tsv - annotated-calls/adhoc-lancet/simulated-bwa.DEL.100-250.1.0.tsv - annotated-calls/adhoc-manta/simulated-bwa.DEL.1-30.1.0.tsv - annotated-calls/adhoc-manta/simulated-bwa.DEL.30-50.1.0.tsv - annotated-calls/adhoc-manta/simulated-bwa.DEL.50-100.1.0.tsv - annotated-calls/adhoc-manta/simulated-bwa.DEL.100-250.1.0.tsv - annotated-calls/adhoc-strelka/simulated-bwa.DEL.1-30.1.0.tsv - annotated-calls/adhoc-strelka/simulated-bwa.DEL.30-50.1.0.tsv - annotated-calls/adhoc-strelka/simulated-bwa.DEL.50-100.1.0.tsv - annotated-calls/adhoc-strelka/simulated-bwa.DEL.100-250.1.0.tsv - annotated-calls/adhoc-bpi/simulated-bwa.DEL.1-30.1.0.tsv - annotated-calls/adhoc-bpi/simulated-bwa.DEL.30-50.1.0.tsv - annotated-calls/adhoc-bpi/simulated-bwa.DEL.50-100.1.0.tsv - annotated-calls/adhoc-bpi/simulated-bwa.DEL.100-250.1.0.tsv - annotated-calls/varlociraptor-delly/simulated-bwa.INS.1-30.1e-05.tsv - annotated-calls/varlociraptor-delly/simulated-bwa.INS.1-30.0.0001.tsv - annotated-calls/varlociraptor-delly/simulated-bwa.INS.1-30.0.001.tsv - annotated-calls/varlociraptor-delly/simulated-bwa.INS.1-30.0.01.tsv - annotated-calls/varlociraptor-delly/simulated-bwa.INS.1-30.0.05.tsv - annotated-calls/varlociraptor-delly/simulated-bwa.INS.1-30.0.1.tsv - annotated-calls/varlociraptor-delly/simulated-bwa.INS.1-30.0.2.tsv - annotated-calls/varlociraptor-delly/simulated-bwa.INS.1-30.0.3.tsv - annotated-calls/varlociraptor-delly/simulated-bwa.INS.1-30.0.4.tsv - annotated-calls/varlociraptor-delly/simulated-bwa.INS.1-30.0.5.tsv - annotated-calls/varlociraptor-delly/simulated-bwa.INS.1-30.0.6.tsv - annotated-calls/varlociraptor-delly/simulated-bwa.INS.1-30.0.7.tsv - annotated-calls/varlociraptor-delly/simulated-bwa.INS.1-30.0.8.tsv - annotated-calls/varlociraptor-delly/simulated-bwa.INS.1-30.0.9.tsv - annotated-calls/varlociraptor-delly/simulated-bwa.INS.30-100.1e-05.tsv - annotated-calls/varlociraptor-delly/simulated-bwa.INS.30-100.0.0001.tsv - annotated-calls/varlociraptor-delly/simulated-bwa.INS.30-100.0.001.tsv - annotated-calls/varlociraptor-delly/simulated-bwa.INS.30-100.0.01.tsv - annotated-calls/varlociraptor-delly/simulated-bwa.INS.30-100.0.05.tsv - annotated-calls/varlociraptor-delly/simulated-bwa.INS.30-100.0.1.tsv - annotated-calls/varlociraptor-delly/simulated-bwa.INS.30-100.0.2.tsv - annotated-calls/varlociraptor-delly/simulated-bwa.INS.30-100.0.3.tsv - annotated-calls/varlociraptor-delly/simulated-bwa.INS.30-100.0.4.tsv - annotated-calls/varlociraptor-delly/simulated-bwa.INS.30-100.0.5.tsv - annotated-calls/varlociraptor-delly/simulated-bwa.INS.30-100.0.6.tsv - annotated-calls/varlociraptor-delly/simulated-bwa.INS.30-100.0.7.tsv - annotated-calls/varlociraptor-delly/simulated-bwa.INS.30-100.0.8.tsv - annotated-calls/varlociraptor-delly/simulated-bwa.INS.30-100.0.9.tsv - annotated-calls/varlociraptor-lancet/simulated-bwa.INS.1-30.1e-05.tsv - annotated-calls/varlociraptor-lancet/simulated-bwa.INS.1-30.0.0001.tsv - annotated-calls/varlociraptor-lancet/simulated-bwa.INS.1-30.0.001.tsv - annotated-calls/varlociraptor-lancet/simulated-bwa.INS.1-30.0.01.tsv - annotated-calls/varlociraptor-lancet/simulated-bwa.INS.1-30.0.05.tsv - annotated-calls/varlociraptor-lancet/simulated-bwa.INS.1-30.0.1.tsv - annotated-calls/varlociraptor-lancet/simulated-bwa.INS.1-30.0.2.tsv - annotated-calls/varlociraptor-lancet/simulated-bwa.INS.1-30.0.3.tsv - annotated-calls/varlociraptor-lancet/simulated-bwa.INS.1-30.0.4.tsv - annotated-calls/varlociraptor-lancet/simulated-bwa.INS.1-30.0.5.tsv - annotated-calls/varlociraptor-lancet/simulated-bwa.INS.1-30.0.6.tsv - annotated-calls/varlociraptor-lancet/simulated-bwa.INS.1-30.0.7.tsv - annotated-calls/varlociraptor-lancet/simulated-bwa.INS.1-30.0.8.tsv - annotated-calls/varlociraptor-lancet/simulated-bwa.INS.1-30.0.9.tsv - annotated-calls/varlociraptor-lancet/simulated-bwa.INS.30-100.1e-05.tsv - annotated-calls/varlociraptor-lancet/simulated-bwa.INS.30-100.0.0001.tsv - annotated-calls/varlociraptor-lancet/simulated-bwa.INS.30-100.0.001.tsv - annotated-calls/varlociraptor-lancet/simulated-bwa.INS.30-100.0.01.tsv - annotated-calls/varlociraptor-lancet/simulated-bwa.INS.30-100.0.05.tsv - annotated-calls/varlociraptor-lancet/simulated-bwa.INS.30-100.0.1.tsv - annotated-calls/varlociraptor-lancet/simulated-bwa.INS.30-100.0.2.tsv - annotated-calls/varlociraptor-lancet/simulated-bwa.INS.30-100.0.3.tsv - annotated-calls/varlociraptor-lancet/simulated-bwa.INS.30-100.0.4.tsv - annotated-calls/varlociraptor-lancet/simulated-bwa.INS.30-100.0.5.tsv - annotated-calls/varlociraptor-lancet/simulated-bwa.INS.30-100.0.6.tsv - annotated-calls/varlociraptor-lancet/simulated-bwa.INS.30-100.0.7.tsv - annotated-calls/varlociraptor-lancet/simulated-bwa.INS.30-100.0.8.tsv - annotated-calls/varlociraptor-lancet/simulated-bwa.INS.30-100.0.9.tsv - annotated-calls/varlociraptor-manta/simulated-bwa.INS.1-30.1e-05.tsv - annotated-calls/varlociraptor-manta/simulated-bwa.INS.1-30.0.0001.tsv - annotated-calls/varlociraptor-manta/simulated-bwa.INS.1-30.0.001.tsv - annotated-calls/varlociraptor-manta/simulated-bwa.INS.1-30.0.01.tsv - annotated-calls/varlociraptor-manta/simulated-bwa.INS.1-30.0.05.tsv - annotated-calls/varlociraptor-manta/simulated-bwa.INS.1-30.0.1.tsv - annotated-calls/varlociraptor-manta/simulated-bwa.INS.1-30.0.2.tsv - annotated-calls/varlociraptor-manta/simulated-bwa.INS.1-30.0.3.tsv - annotated-calls/varlociraptor-manta/simulated-bwa.INS.1-30.0.4.tsv - annotated-calls/varlociraptor-manta/simulated-bwa.INS.1-30.0.5.tsv - annotated-calls/varlociraptor-manta/simulated-bwa.INS.1-30.0.6.tsv - annotated-calls/varlociraptor-manta/simulated-bwa.INS.1-30.0.7.tsv - annotated-calls/varlociraptor-manta/simulated-bwa.INS.1-30.0.8.tsv - annotated-calls/varlociraptor-manta/simulated-bwa.INS.1-30.0.9.tsv - annotated-calls/varlociraptor-manta/simulated-bwa.INS.30-100.1e-05.tsv - annotated-calls/varlociraptor-manta/simulated-bwa.INS.30-100.0.0001.tsv - annotated-calls/varlociraptor-manta/simulated-bwa.INS.30-100.0.001.tsv - annotated-calls/varlociraptor-manta/simulated-bwa.INS.30-100.0.01.tsv - annotated-calls/varlociraptor-manta/simulated-bwa.INS.30-100.0.05.tsv - annotated-calls/varlociraptor-manta/simulated-bwa.INS.30-100.0.1.tsv - annotated-calls/varlociraptor-manta/simulated-bwa.INS.30-100.0.2.tsv - annotated-calls/varlociraptor-manta/simulated-bwa.INS.30-100.0.3.tsv - annotated-calls/varlociraptor-manta/simulated-bwa.INS.30-100.0.4.tsv - annotated-calls/varlociraptor-manta/simulated-bwa.INS.30-100.0.5.tsv - annotated-calls/varlociraptor-manta/simulated-bwa.INS.30-100.0.6.tsv - annotated-calls/varlociraptor-manta/simulated-bwa.INS.30-100.0.7.tsv - annotated-calls/varlociraptor-manta/simulated-bwa.INS.30-100.0.8.tsv - annotated-calls/varlociraptor-manta/simulated-bwa.INS.30-100.0.9.tsv - annotated-calls/varlociraptor-strelka/simulated-bwa.INS.1-30.1e-05.tsv - annotated-calls/varlociraptor-strelka/simulated-bwa.INS.1-30.0.0001.tsv - annotated-calls/varlociraptor-strelka/simulated-bwa.INS.1-30.0.001.tsv - annotated-calls/varlociraptor-strelka/simulated-bwa.INS.1-30.0.01.tsv - annotated-calls/varlociraptor-strelka/simulated-bwa.INS.1-30.0.05.tsv - annotated-calls/varlociraptor-strelka/simulated-bwa.INS.1-30.0.1.tsv - annotated-calls/varlociraptor-strelka/simulated-bwa.INS.1-30.0.2.tsv - annotated-calls/varlociraptor-strelka/simulated-bwa.INS.1-30.0.3.tsv - annotated-calls/varlociraptor-strelka/simulated-bwa.INS.1-30.0.4.tsv - annotated-calls/varlociraptor-strelka/simulated-bwa.INS.1-30.0.5.tsv - annotated-calls/varlociraptor-strelka/simulated-bwa.INS.1-30.0.6.tsv - annotated-calls/varlociraptor-strelka/simulated-bwa.INS.1-30.0.7.tsv - annotated-calls/varlociraptor-strelka/simulated-bwa.INS.1-30.0.8.tsv - annotated-calls/varlociraptor-strelka/simulated-bwa.INS.1-30.0.9.tsv - annotated-calls/varlociraptor-strelka/simulated-bwa.INS.30-100.1e-05.tsv - annotated-calls/varlociraptor-strelka/simulated-bwa.INS.30-100.0.0001.tsv - annotated-calls/varlociraptor-strelka/simulated-bwa.INS.30-100.0.001.tsv - annotated-calls/varlociraptor-strelka/simulated-bwa.INS.30-100.0.01.tsv - annotated-calls/varlociraptor-strelka/simulated-bwa.INS.30-100.0.05.tsv - annotated-calls/varlociraptor-strelka/simulated-bwa.INS.30-100.0.1.tsv - annotated-calls/varlociraptor-strelka/simulated-bwa.INS.30-100.0.2.tsv - annotated-calls/varlociraptor-strelka/simulated-bwa.INS.30-100.0.3.tsv - annotated-calls/varlociraptor-strelka/simulated-bwa.INS.30-100.0.4.tsv - annotated-calls/varlociraptor-strelka/simulated-bwa.INS.30-100.0.5.tsv - annotated-calls/varlociraptor-strelka/simulated-bwa.INS.30-100.0.6.tsv - annotated-calls/varlociraptor-strelka/simulated-bwa.INS.30-100.0.7.tsv - annotated-calls/varlociraptor-strelka/simulated-bwa.INS.30-100.0.8.tsv - annotated-calls/varlociraptor-strelka/simulated-bwa.INS.30-100.0.9.tsv - annotated-calls/varlociraptor-delly/simulated-bwa.DEL.1-30.1e-05.tsv - annotated-calls/varlociraptor-delly/simulated-bwa.DEL.1-30.0.0001.tsv - annotated-calls/varlociraptor-delly/simulated-bwa.DEL.1-30.0.001.tsv - annotated-calls/varlociraptor-delly/simulated-bwa.DEL.1-30.0.01.tsv - annotated-calls/varlociraptor-delly/simulated-bwa.DEL.1-30.0.05.tsv - annotated-calls/varlociraptor-delly/simulated-bwa.DEL.1-30.0.1.tsv - annotated-calls/varlociraptor-delly/simulated-bwa.DEL.1-30.0.2.tsv - annotated-calls/varlociraptor-delly/simulated-bwa.DEL.1-30.0.3.tsv - annotated-calls/varlociraptor-delly/simulated-bwa.DEL.1-30.0.4.tsv - annotated-calls/varlociraptor-delly/simulated-bwa.DEL.1-30.0.5.tsv - annotated-calls/varlociraptor-delly/simulated-bwa.DEL.1-30.0.6.tsv - annotated-calls/varlociraptor-delly/simulated-bwa.DEL.1-30.0.7.tsv - annotated-calls/varlociraptor-delly/simulated-bwa.DEL.1-30.0.8.tsv - annotated-calls/varlociraptor-delly/simulated-bwa.DEL.1-30.0.9.tsv - annotated-calls/varlociraptor-delly/simulated-bwa.DEL.30-50.1e-05.tsv - annotated-calls/varlociraptor-delly/simulated-bwa.DEL.30-50.0.0001.tsv - annotated-calls/varlociraptor-delly/simulated-bwa.DEL.30-50.0.001.tsv - annotated-calls/varlociraptor-delly/simulated-bwa.DEL.30-50.0.01.tsv - annotated-calls/varlociraptor-delly/simulated-bwa.DEL.30-50.0.05.tsv - annotated-calls/varlociraptor-delly/simulated-bwa.DEL.30-50.0.1.tsv - annotated-calls/varlociraptor-delly/simulated-bwa.DEL.30-50.0.2.tsv - annotated-calls/varlociraptor-delly/simulated-bwa.DEL.30-50.0.3.tsv - annotated-calls/varlociraptor-delly/simulated-bwa.DEL.30-50.0.4.tsv - annotated-calls/varlociraptor-delly/simulated-bwa.DEL.30-50.0.5.tsv - annotated-calls/varlociraptor-delly/simulated-bwa.DEL.30-50.0.6.tsv - annotated-calls/varlociraptor-delly/simulated-bwa.DEL.30-50.0.7.tsv - annotated-calls/varlociraptor-delly/simulated-bwa.DEL.30-50.0.8.tsv - annotated-calls/varlociraptor-delly/simulated-bwa.DEL.30-50.0.9.tsv - annotated-calls/varlociraptor-delly/simulated-bwa.DEL.50-100.1e-05.tsv - annotated-calls/varlociraptor-delly/simulated-bwa.DEL.50-100.0.0001.tsv - annotated-calls/varlociraptor-delly/simulated-bwa.DEL.50-100.0.001.tsv - annotated-calls/varlociraptor-delly/simulated-bwa.DEL.50-100.0.01.tsv - annotated-calls/varlociraptor-delly/simulated-bwa.DEL.50-100.0.05.tsv - annotated-calls/varlociraptor-delly/simulated-bwa.DEL.50-100.0.1.tsv - annotated-calls/varlociraptor-delly/simulated-bwa.DEL.50-100.0.2.tsv - annotated-calls/varlociraptor-delly/simulated-bwa.DEL.50-100.0.3.tsv - annotated-calls/varlociraptor-delly/simulated-bwa.DEL.50-100.0.4.tsv - annotated-calls/varlociraptor-delly/simulated-bwa.DEL.50-100.0.5.tsv - annotated-calls/varlociraptor-delly/simulated-bwa.DEL.50-100.0.6.tsv - annotated-calls/varlociraptor-delly/simulated-bwa.DEL.50-100.0.7.tsv - annotated-calls/varlociraptor-delly/simulated-bwa.DEL.50-100.0.8.tsv - annotated-calls/varlociraptor-delly/simulated-bwa.DEL.50-100.0.9.tsv - annotated-calls/varlociraptor-delly/simulated-bwa.DEL.100-250.1e-05.tsv - annotated-calls/varlociraptor-delly/simulated-bwa.DEL.100-250.0.0001.tsv - annotated-calls/varlociraptor-delly/simulated-bwa.DEL.100-250.0.001.tsv - annotated-calls/varlociraptor-delly/simulated-bwa.DEL.100-250.0.01.tsv - annotated-calls/varlociraptor-delly/simulated-bwa.DEL.100-250.0.05.tsv - annotated-calls/varlociraptor-delly/simulated-bwa.DEL.100-250.0.1.tsv - annotated-calls/varlociraptor-delly/simulated-bwa.DEL.100-250.0.2.tsv - annotated-calls/varlociraptor-delly/simulated-bwa.DEL.100-250.0.3.tsv - annotated-calls/varlociraptor-delly/simulated-bwa.DEL.100-250.0.4.tsv - annotated-calls/varlociraptor-delly/simulated-bwa.DEL.100-250.0.5.tsv - annotated-calls/varlociraptor-delly/simulated-bwa.DEL.100-250.0.6.tsv - annotated-calls/varlociraptor-delly/simulated-bwa.DEL.100-250.0.7.tsv - annotated-calls/varlociraptor-delly/simulated-bwa.DEL.100-250.0.8.tsv - annotated-calls/varlociraptor-delly/simulated-bwa.DEL.100-250.0.9.tsv - annotated-calls/varlociraptor-lancet/simulated-bwa.DEL.1-30.1e-05.tsv - annotated-calls/varlociraptor-lancet/simulated-bwa.DEL.1-30.0.0001.tsv - annotated-calls/varlociraptor-lancet/simulated-bwa.DEL.1-30.0.001.tsv - annotated-calls/varlociraptor-lancet/simulated-bwa.DEL.1-30.0.01.tsv - annotated-calls/varlociraptor-lancet/simulated-bwa.DEL.1-30.0.05.tsv - annotated-calls/varlociraptor-lancet/simulated-bwa.DEL.1-30.0.1.tsv - annotated-calls/varlociraptor-lancet/simulated-bwa.DEL.1-30.0.2.tsv - annotated-calls/varlociraptor-lancet/simulated-bwa.DEL.1-30.0.3.tsv - annotated-calls/varlociraptor-lancet/simulated-bwa.DEL.1-30.0.4.tsv - annotated-calls/varlociraptor-lancet/simulated-bwa.DEL.1-30.0.5.tsv - annotated-calls/varlociraptor-lancet/simulated-bwa.DEL.1-30.0.6.tsv - annotated-calls/varlociraptor-lancet/simulated-bwa.DEL.1-30.0.7.tsv - annotated-calls/varlociraptor-lancet/simulated-bwa.DEL.1-30.0.8.tsv - annotated-calls/varlociraptor-lancet/simulated-bwa.DEL.1-30.0.9.tsv - annotated-calls/varlociraptor-lancet/simulated-bwa.DEL.30-50.1e-05.tsv - annotated-calls/varlociraptor-lancet/simulated-bwa.DEL.30-50.0.0001.tsv - annotated-calls/varlociraptor-lancet/simulated-bwa.DEL.30-50.0.001.tsv - annotated-calls/varlociraptor-lancet/simulated-bwa.DEL.30-50.0.01.tsv - annotated-calls/varlociraptor-lancet/simulated-bwa.DEL.30-50.0.05.tsv - annotated-calls/varlociraptor-lancet/simulated-bwa.DEL.30-50.0.1.tsv - annotated-calls/varlociraptor-lancet/simulated-bwa.DEL.30-50.0.2.tsv - annotated-calls/varlociraptor-lancet/simulated-bwa.DEL.30-50.0.3.tsv - annotated-calls/varlociraptor-lancet/simulated-bwa.DEL.30-50.0.4.tsv - annotated-calls/varlociraptor-lancet/simulated-bwa.DEL.30-50.0.5.tsv - annotated-calls/varlociraptor-lancet/simulated-bwa.DEL.30-50.0.6.tsv - annotated-calls/varlociraptor-lancet/simulated-bwa.DEL.30-50.0.7.tsv - annotated-calls/varlociraptor-lancet/simulated-bwa.DEL.30-50.0.8.tsv - annotated-calls/varlociraptor-lancet/simulated-bwa.DEL.30-50.0.9.tsv - annotated-calls/varlociraptor-lancet/simulated-bwa.DEL.50-100.1e-05.tsv - annotated-calls/varlociraptor-lancet/simulated-bwa.DEL.50-100.0.0001.tsv - annotated-calls/varlociraptor-lancet/simulated-bwa.DEL.50-100.0.001.tsv - annotated-calls/varlociraptor-lancet/simulated-bwa.DEL.50-100.0.01.tsv - annotated-calls/varlociraptor-lancet/simulated-bwa.DEL.50-100.0.05.tsv - annotated-calls/varlociraptor-lancet/simulated-bwa.DEL.50-100.0.1.tsv - annotated-calls/varlociraptor-lancet/simulated-bwa.DEL.50-100.0.2.tsv - annotated-calls/varlociraptor-lancet/simulated-bwa.DEL.50-100.0.3.tsv - annotated-calls/varlociraptor-lancet/simulated-bwa.DEL.50-100.0.4.tsv - annotated-calls/varlociraptor-lancet/simulated-bwa.DEL.50-100.0.5.tsv - annotated-calls/varlociraptor-lancet/simulated-bwa.DEL.50-100.0.6.tsv - annotated-calls/varlociraptor-lancet/simulated-bwa.DEL.50-100.0.7.tsv - annotated-calls/varlociraptor-lancet/simulated-bwa.DEL.50-100.0.8.tsv - annotated-calls/varlociraptor-lancet/simulated-bwa.DEL.50-100.0.9.tsv - annotated-calls/varlociraptor-lancet/simulated-bwa.DEL.100-250.1e-05.tsv - annotated-calls/varlociraptor-lancet/simulated-bwa.DEL.100-250.0.0001.tsv - annotated-calls/varlociraptor-lancet/simulated-bwa.DEL.100-250.0.001.tsv - annotated-calls/varlociraptor-lancet/simulated-bwa.DEL.100-250.0.01.tsv - annotated-calls/varlociraptor-lancet/simulated-bwa.DEL.100-250.0.05.tsv - annotated-calls/varlociraptor-lancet/simulated-bwa.DEL.100-250.0.1.tsv - annotated-calls/varlociraptor-lancet/simulated-bwa.DEL.100-250.0.2.tsv - annotated-calls/varlociraptor-lancet/simulated-bwa.DEL.100-250.0.3.tsv - annotated-calls/varlociraptor-lancet/simulated-bwa.DEL.100-250.0.4.tsv - annotated-calls/varlociraptor-lancet/simulated-bwa.DEL.100-250.0.5.tsv - annotated-calls/varlociraptor-lancet/simulated-bwa.DEL.100-250.0.6.tsv - annotated-calls/varlociraptor-lancet/simulated-bwa.DEL.100-250.0.7.tsv - annotated-calls/varlociraptor-lancet/simulated-bwa.DEL.100-250.0.8.tsv - annotated-calls/varlociraptor-lancet/simulated-bwa.DEL.100-250.0.9.tsv - annotated-calls/varlociraptor-manta/simulated-bwa.DEL.1-30.1e-05.tsv - annotated-calls/varlociraptor-manta/simulated-bwa.DEL.1-30.0.0001.tsv - annotated-calls/varlociraptor-manta/simulated-bwa.DEL.1-30.0.001.tsv - annotated-calls/varlociraptor-manta/simulated-bwa.DEL.1-30.0.01.tsv - annotated-calls/varlociraptor-manta/simulated-bwa.DEL.1-30.0.05.tsv - annotated-calls/varlociraptor-manta/simulated-bwa.DEL.1-30.0.1.tsv - annotated-calls/varlociraptor-manta/simulated-bwa.DEL.1-30.0.2.tsv - annotated-calls/varlociraptor-manta/simulated-bwa.DEL.1-30.0.3.tsv - annotated-calls/varlociraptor-manta/simulated-bwa.DEL.1-30.0.4.tsv - annotated-calls/varlociraptor-manta/simulated-bwa.DEL.1-30.0.5.tsv - annotated-calls/varlociraptor-manta/simulated-bwa.DEL.1-30.0.6.tsv - annotated-calls/varlociraptor-manta/simulated-bwa.DEL.1-30.0.7.tsv - annotated-calls/varlociraptor-manta/simulated-bwa.DEL.1-30.0.8.tsv - annotated-calls/varlociraptor-manta/simulated-bwa.DEL.1-30.0.9.tsv - annotated-calls/varlociraptor-manta/simulated-bwa.DEL.30-50.1e-05.tsv - annotated-calls/varlociraptor-manta/simulated-bwa.DEL.30-50.0.0001.tsv - annotated-calls/varlociraptor-manta/simulated-bwa.DEL.30-50.0.001.tsv - annotated-calls/varlociraptor-manta/simulated-bwa.DEL.30-50.0.01.tsv - annotated-calls/varlociraptor-manta/simulated-bwa.DEL.30-50.0.05.tsv - annotated-calls/varlociraptor-manta/simulated-bwa.DEL.30-50.0.1.tsv - annotated-calls/varlociraptor-manta/simulated-bwa.DEL.30-50.0.2.tsv - annotated-calls/varlociraptor-manta/simulated-bwa.DEL.30-50.0.3.tsv - annotated-calls/varlociraptor-manta/simulated-bwa.DEL.30-50.0.4.tsv - annotated-calls/varlociraptor-manta/simulated-bwa.DEL.30-50.0.5.tsv - annotated-calls/varlociraptor-manta/simulated-bwa.DEL.30-50.0.6.tsv - annotated-calls/varlociraptor-manta/simulated-bwa.DEL.30-50.0.7.tsv - annotated-calls/varlociraptor-manta/simulated-bwa.DEL.30-50.0.8.tsv - annotated-calls/varlociraptor-manta/simulated-bwa.DEL.30-50.0.9.tsv - annotated-calls/varlociraptor-manta/simulated-bwa.DEL.50-100.1e-05.tsv - annotated-calls/varlociraptor-manta/simulated-bwa.DEL.50-100.0.0001.tsv - annotated-calls/varlociraptor-manta/simulated-bwa.DEL.50-100.0.001.tsv - annotated-calls/varlociraptor-manta/simulated-bwa.DEL.50-100.0.01.tsv - annotated-calls/varlociraptor-manta/simulated-bwa.DEL.50-100.0.05.tsv - annotated-calls/varlociraptor-manta/simulated-bwa.DEL.50-100.0.1.tsv - annotated-calls/varlociraptor-manta/simulated-bwa.DEL.50-100.0.2.tsv - annotated-calls/varlociraptor-manta/simulated-bwa.DEL.50-100.0.3.tsv - annotated-calls/varlociraptor-manta/simulated-bwa.DEL.50-100.0.4.tsv - annotated-calls/varlociraptor-manta/simulated-bwa.DEL.50-100.0.5.tsv - annotated-calls/varlociraptor-manta/simulated-bwa.DEL.50-100.0.6.tsv - annotated-calls/varlociraptor-manta/simulated-bwa.DEL.50-100.0.7.tsv - annotated-calls/varlociraptor-manta/simulated-bwa.DEL.50-100.0.8.tsv - annotated-calls/varlociraptor-manta/simulated-bwa.DEL.50-100.0.9.tsv - annotated-calls/varlociraptor-manta/simulated-bwa.DEL.100-250.1e-05.tsv - annotated-calls/varlociraptor-manta/simulated-bwa.DEL.100-250.0.0001.tsv - annotated-calls/varlociraptor-manta/simulated-bwa.DEL.100-250.0.001.tsv - annotated-calls/varlociraptor-manta/simulated-bwa.DEL.100-250.0.01.tsv - annotated-calls/varlociraptor-manta/simulated-bwa.DEL.100-250.0.05.tsv - annotated-calls/varlociraptor-manta/simulated-bwa.DEL.100-250.0.1.tsv - annotated-calls/varlociraptor-manta/simulated-bwa.DEL.100-250.0.2.tsv - annotated-calls/varlociraptor-manta/simulated-bwa.DEL.100-250.0.3.tsv - annotated-calls/varlociraptor-manta/simulated-bwa.DEL.100-250.0.4.tsv - annotated-calls/varlociraptor-manta/simulated-bwa.DEL.100-250.0.5.tsv - annotated-calls/varlociraptor-manta/simulated-bwa.DEL.100-250.0.6.tsv - annotated-calls/varlociraptor-manta/simulated-bwa.DEL.100-250.0.7.tsv - annotated-calls/varlociraptor-manta/simulated-bwa.DEL.100-250.0.8.tsv - annotated-calls/varlociraptor-manta/simulated-bwa.DEL.100-250.0.9.tsv - annotated-calls/varlociraptor-strelka/simulated-bwa.DEL.1-30.1e-05.tsv - annotated-calls/varlociraptor-strelka/simulated-bwa.DEL.1-30.0.0001.tsv - annotated-calls/varlociraptor-strelka/simulated-bwa.DEL.1-30.0.001.tsv - annotated-calls/varlociraptor-strelka/simulated-bwa.DEL.1-30.0.01.tsv - annotated-calls/varlociraptor-strelka/simulated-bwa.DEL.1-30.0.05.tsv - annotated-calls/varlociraptor-strelka/simulated-bwa.DEL.1-30.0.1.tsv - annotated-calls/varlociraptor-strelka/simulated-bwa.DEL.1-30.0.2.tsv - annotated-calls/varlociraptor-strelka/simulated-bwa.DEL.1-30.0.3.tsv - annotated-calls/varlociraptor-strelka/simulated-bwa.DEL.1-30.0.4.tsv - annotated-calls/varlociraptor-strelka/simulated-bwa.DEL.1-30.0.5.tsv - annotated-calls/varlociraptor-strelka/simulated-bwa.DEL.1-30.0.6.tsv - annotated-calls/varlociraptor-strelka/simulated-bwa.DEL.1-30.0.7.tsv - annotated-calls/varlociraptor-strelka/simulated-bwa.DEL.1-30.0.8.tsv - annotated-calls/varlociraptor-strelka/simulated-bwa.DEL.1-30.0.9.tsv - annotated-calls/varlociraptor-strelka/simulated-bwa.DEL.30-50.1e-05.tsv - annotated-calls/varlociraptor-strelka/simulated-bwa.DEL.30-50.0.0001.tsv - annotated-calls/varlociraptor-strelka/simulated-bwa.DEL.30-50.0.001.tsv - annotated-calls/varlociraptor-strelka/simulated-bwa.DEL.30-50.0.01.tsv - annotated-calls/varlociraptor-strelka/simulated-bwa.DEL.30-50.0.05.tsv - annotated-calls/varlociraptor-strelka/simulated-bwa.DEL.30-50.0.1.tsv - annotated-calls/varlociraptor-strelka/simulated-bwa.DEL.30-50.0.2.tsv - annotated-calls/varlociraptor-strelka/simulated-bwa.DEL.30-50.0.3.tsv - annotated-calls/varlociraptor-strelka/simulated-bwa.DEL.30-50.0.4.tsv - annotated-calls/varlociraptor-strelka/simulated-bwa.DEL.30-50.0.5.tsv - annotated-calls/varlociraptor-strelka/simulated-bwa.DEL.30-50.0.6.tsv - annotated-calls/varlociraptor-strelka/simulated-bwa.DEL.30-50.0.7.tsv - annotated-calls/varlociraptor-strelka/simulated-bwa.DEL.30-50.0.8.tsv - annotated-calls/varlociraptor-strelka/simulated-bwa.DEL.30-50.0.9.tsv - annotated-calls/varlociraptor-strelka/simulated-bwa.DEL.50-100.1e-05.tsv - annotated-calls/varlociraptor-strelka/simulated-bwa.DEL.50-100.0.0001.tsv - annotated-calls/varlociraptor-strelka/simulated-bwa.DEL.50-100.0.001.tsv - annotated-calls/varlociraptor-strelka/simulated-bwa.DEL.50-100.0.01.tsv - annotated-calls/varlociraptor-strelka/simulated-bwa.DEL.50-100.0.05.tsv - annotated-calls/varlociraptor-strelka/simulated-bwa.DEL.50-100.0.1.tsv - annotated-calls/varlociraptor-strelka/simulated-bwa.DEL.50-100.0.2.tsv - annotated-calls/varlociraptor-strelka/simulated-bwa.DEL.50-100.0.3.tsv - annotated-calls/varlociraptor-strelka/simulated-bwa.DEL.50-100.0.4.tsv - annotated-calls/varlociraptor-strelka/simulated-bwa.DEL.50-100.0.5.tsv - annotated-calls/varlociraptor-strelka/simulated-bwa.DEL.50-100.0.6.tsv - annotated-calls/varlociraptor-strelka/simulated-bwa.DEL.50-100.0.7.tsv - annotated-calls/varlociraptor-strelka/simulated-bwa.DEL.50-100.0.8.tsv - annotated-calls/varlociraptor-strelka/simulated-bwa.DEL.50-100.0.9.tsv - annotated-calls/varlociraptor-strelka/simulated-bwa.DEL.100-250.1e-05.tsv - annotated-calls/varlociraptor-strelka/simulated-bwa.DEL.100-250.0.0001.tsv - annotated-calls/varlociraptor-strelka/simulated-bwa.DEL.100-250.0.001.tsv - annotated-calls/varlociraptor-strelka/simulated-bwa.DEL.100-250.0.01.tsv - annotated-calls/varlociraptor-strelka/simulated-bwa.DEL.100-250.0.05.tsv - annotated-calls/varlociraptor-strelka/simulated-bwa.DEL.100-250.0.1.tsv - annotated-calls/varlociraptor-strelka/simulated-bwa.DEL.100-250.0.2.tsv - annotated-calls/varlociraptor-strelka/simulated-bwa.DEL.100-250.0.3.tsv - annotated-calls/varlociraptor-strelka/simulated-bwa.DEL.100-250.0.4.tsv - annotated-calls/varlociraptor-strelka/simulated-bwa.DEL.100-250.0.5.tsv - annotated-calls/varlociraptor-strelka/simulated-bwa.DEL.100-250.0.6.tsv - annotated-calls/varlociraptor-strelka/simulated-bwa.DEL.100-250.0.7.tsv - annotated-calls/varlociraptor-strelka/simulated-bwa.DEL.100-250.0.8.tsv - annotated-calls/varlociraptor-strelka/simulated-bwa.DEL.100-250.0.9.tsv - annotated-calls/varlociraptor-delly/simulated-bwa.INS.1-250.1.0.tsv - annotated-calls/varlociraptor-lancet/simulated-bwa.INS.1-250.1.0.tsv - annotated-calls/varlociraptor-manta/simulated-bwa.INS.1-250.1.0.tsv - annotated-calls/varlociraptor-strelka/simulated-bwa.INS.1-250.1.0.tsv - annotated-calls/varlociraptor-delly/simulated-bwa.DEL.1-250.1.0.tsv - annotated-calls/varlociraptor-lancet/simulated-bwa.DEL.1-250.1.0.tsv - annotated-calls/varlociraptor-manta/simulated-bwa.DEL.1-250.1.0.tsv - annotated-calls/varlociraptor-strelka/simulated-bwa.DEL.1-250.1.0.tsv |  | - python =3.6 - pandas =0.23 - matplotlib =3.0 - seaborn =0.9.0 - pysam =0.13.0 - svgutils =0.2 - pybedtools =0.7.10 - networkx =2.2 | ``` 1 2 3 4 5 6 7 8 9 10 11 12 13 14 15 16 17 18 19 20 21 22 23 24 25 26 ``` \| ``` import pandas as pd import numpy as np from common import load_variants minlen = int(snakemake.wildcards.minlen) maxlen = int(snakemake.wildcards.maxlen) vartype = snakemake.wildcards.vartype if snakemake.wildcards.mode == "varlociraptor": score = snakemake.config["caller"]["varlociraptor"]["score"] # calls are already filtered by FDR control step minlen = None maxlen = None elif snakemake.wildcards.mode == "default": score = snakemake.config["caller"][snakemake.wildcards.caller]["score"] else: score = None calls = load_variants(snakemake.input.calls, vartype=vartype, minlen=minlen, maxlen=maxlen) calls["is_tp"] = calls["MATCHING"] >= 0 calls["score"] = calls[score] if score else np.nan calls.to_csv(snakemake.output[0], sep="\t") ``` |
| aggregate\_concordance | 30 | - aggregated-concordance/varlociraptor-delly-0.9/colo1.INS.tsv - aggregated-concordance/varlociraptor-lancet-0.9/colo1.INS.tsv - aggregated-concordance/varlociraptor-manta-0.9/colo1.INS.tsv - aggregated-concordance/varlociraptor-strelka-0.9/colo1.INS.tsv - aggregated-concordance/varlociraptor-bpi-0.9/colo1.INS.tsv - aggregated-concordance/varlociraptor-delly-0.98/colo1.INS.tsv - aggregated-concordance/varlociraptor-lancet-0.98/colo1.INS.tsv - aggregated-concordance/varlociraptor-manta-0.98/colo1.INS.tsv - aggregated-concordance/varlociraptor-strelka-0.98/colo1.INS.tsv - aggregated-concordance/varlociraptor-bpi-0.98/colo1.INS.tsv - aggregated-concordance/adhoc-delly-default/colo1.INS.tsv - aggregated-concordance/adhoc-lancet-default/colo1.INS.tsv - aggregated-concordance/adhoc-manta-default/colo1.INS.tsv - aggregated-concordance/adhoc-strelka-default/colo1.INS.tsv - aggregated-concordance/adhoc-bpi-default/colo1.INS.tsv - aggregated-concordance/varlociraptor-delly-0.9/colo1.DEL.tsv - aggregated-concordance/varlociraptor-lancet-0.9/colo1.DEL.tsv - aggregated-concordance/varlociraptor-manta-0.9/colo1.DEL.tsv - aggregated-concordance/varlociraptor-strelka-0.9/colo1.DEL.tsv - aggregated-concordance/varlociraptor-bpi-0.9/colo1.DEL.tsv - aggregated-concordance/varlociraptor-delly-0.98/colo1.DEL.tsv - aggregated-concordance/varlociraptor-lancet-0.98/colo1.DEL.tsv - aggregated-concordance/varlociraptor-manta-0.98/colo1.DEL.tsv - aggregated-concordance/varlociraptor-strelka-0.98/colo1.DEL.tsv - aggregated-concordance/varlociraptor-bpi-0.98/colo1.DEL.tsv - aggregated-concordance/adhoc-delly-default/colo1.DEL.tsv - aggregated-concordance/adhoc-lancet-default/colo1.DEL.tsv - aggregated-concordance/adhoc-manta-default/colo1.DEL.tsv - aggregated-concordance/adhoc-strelka-default/colo1.DEL.tsv - aggregated-concordance/adhoc-bpi-default/colo1.DEL.tsv |  | - python =3.6 - pandas =0.23 - matplotlib =3.0 - seaborn =0.9.0 - pysam =0.13.0 - svgutils =0.2 - pybedtools =0.7.10 - networkx =2.2 | ``` 1 2 3 4 5 6 7 8 9 10 11 12 13 14 15 16 17 18 19 20 21 22 23 24 25 26 27 28 29 30 31 32 33 34 35 36 37 38 39 40 41 42 43 44 45 46 47 48 49 50 51 52 53 54 55 56 57 58 59 60 61 62 63 64 65 66 67 68 69 70 71 72 73 74 75 76 77 ``` \| ``` from common import load_variants import networkx as nx import pandas as pd import numpy as np vartype = snakemake.wildcards.vartype index_cols = ["CHROM", "POS", "SVLEN"] if vartype == "INS" or vartype == "DEL" else ["CHROM", "POS", "ALT"] all_variants = [load_variants(f, vartype=vartype) for f in snakemake.input.calls] G = nx.Graph() for calls, (i, j) in zip(all_variants, snakemake.params.dataset_combinations): calls["component"] = None for call in calls.itertuples(): a = (i, call.Index) G.add_node(a) if call.MATCHING >= 0: b = (j, call.MATCHING) G.add_node(b) G.add_edge(a, b) # get a set of calls for each dataset (we don't need all pairwise comparisons for that) representatives = {snakemake.params.dataset_combinations[i][0]: calls for i, calls in enumerate(all_variants)} if snakemake.wildcards.mode != "varlociraptor": varlociraptor_variants = [load_variants(f, vartype=vartype) for f in snakemake.input.varlociraptor_calls] for calls in varlociraptor_variants: calls.set_index(index_cols, inplace=True) varlociraptor_representatives = {snakemake.params.dataset_combinations[i][0]: calls for i, calls in enumerate(varlociraptor_variants)} # annotate calls with their component, i.e. their equivalence class for component_id, component in enumerate(nx.connected_components(G)): for i, k in component: representatives[i].loc[k, "component"] = component_id for calls in representatives.values(): calls["component"] = calls["component"].astype(np.float32) calls.set_index("component", inplace=True) # join calls based on their equivalence class aggregated = None suffix = "_{}".format dataset_name = lambda i: snakemake.params.datasets[i] is_varlociraptor = False for dataset_id, calls in representatives.items(): cols = list(index_cols) if "CASE_AF" in calls.columns: cols.extend(["CASE_AF", "PROB_SOMATIC_TUMOR"]) is_varlociraptor = True calls = calls[cols] if snakemake.wildcards.mode != "varlociraptor": caseaf = calls.set_index(cols, drop=False).join(varlociraptor_representatives[dataset_id][["CASE_AF"]], how="left")["CASE_AF"] caseaf = caseaf[~caseaf.index.duplicated()] calls["CASE_AF"] = caseaf.values calls.columns = [c + suffix(dataset_name(dataset_id)) for c in calls.columns] if aggregated is None: aggregated = calls else: aggregated = aggregated.join(calls, how="outer", lsuffix="", rsuffix="") # Forget the component id. Otherwise, we might run into errors with duplicate elements # in the index below. These can occur if there are multiple ambiguous calls. aggregated.reset_index(inplace=True, drop=True) pos_cols = aggregated.columns[aggregated.columns.str.startswith("POS_")] is_called = (~aggregated[pos_cols].isnull()).astype(int) is_called.columns = pos_cols.str.replace("POS_", "") aggregated = aggregated.join(is_called, lsuffix="", rsuffix="") aggregated.insert(len(aggregated.columns), "concordance_count", is_called.sum(axis=1)) aggregated["max_case_af"] = aggregated[aggregated.columns[aggregated.columns.str.startswith("CASE_AF")]].max(axis=1) if is_varlociraptor: aggregated["max_prob_somatic_tumor"] = aggregated[aggregated.columns[aggregated.columns.str.startswith("PROB_SOMATIC")]].min(axis=1) aggregated.to_csv(snakemake.output[0], sep="\t", index=False) ``` |
| varlociraptor\_calls\_to\_tsv | 368 | - matched-calls/varlociraptor-delly/simulated-bwa.INS.1-30.1.0.tsv - matched-calls/varlociraptor-delly/simulated-bwa.INS.30-100.1.0.tsv - matched-calls/varlociraptor-lancet/simulated-bwa.INS.1-30.1.0.tsv - matched-calls/varlociraptor-lancet/simulated-bwa.INS.30-100.1.0.tsv - matched-calls/varlociraptor-manta/simulated-bwa.INS.1-30.1.0.tsv - matched-calls/varlociraptor-manta/simulated-bwa.INS.30-100.1.0.tsv - matched-calls/varlociraptor-strelka/simulated-bwa.INS.1-30.1.0.tsv - matched-calls/varlociraptor-strelka/simulated-bwa.INS.30-100.1.0.tsv - matched-calls/varlociraptor-delly/simulated-bwa.DEL.1-30.1.0.tsv - matched-calls/varlociraptor-delly/simulated-bwa.DEL.30-50.1.0.tsv - matched-calls/varlociraptor-delly/simulated-bwa.DEL.50-100.1.0.tsv - matched-calls/varlociraptor-delly/simulated-bwa.DEL.100-250.1.0.tsv - matched-calls/varlociraptor-lancet/simulated-bwa.DEL.1-30.1.0.tsv - matched-calls/varlociraptor-lancet/simulated-bwa.DEL.30-50.1.0.tsv - matched-calls/varlociraptor-lancet/simulated-bwa.DEL.50-100.1.0.tsv - matched-calls/varlociraptor-lancet/simulated-bwa.DEL.100-250.1.0.tsv - matched-calls/varlociraptor-manta/simulated-bwa.DEL.1-30.1.0.tsv - matched-calls/varlociraptor-manta/simulated-bwa.DEL.30-50.1.0.tsv - matched-calls/varlociraptor-manta/simulated-bwa.DEL.50-100.1.0.tsv - matched-calls/varlociraptor-manta/simulated-bwa.DEL.100-250.1.0.tsv - matched-calls/varlociraptor-strelka/simulated-bwa.DEL.1-30.1.0.tsv - matched-calls/varlociraptor-strelka/simulated-bwa.DEL.30-50.1.0.tsv - matched-calls/varlociraptor-strelka/simulated-bwa.DEL.50-100.1.0.tsv - matched-calls/varlociraptor-strelka/simulated-bwa.DEL.100-250.1.0.tsv - matched-calls/varlociraptor-delly/simulated-bwa.INS.1-30.1e-05.tsv - matched-calls/varlociraptor-delly/simulated-bwa.INS.1-30.0.0001.tsv - matched-calls/varlociraptor-delly/simulated-bwa.INS.1-30.0.001.tsv - matched-calls/varlociraptor-delly/simulated-bwa.INS.1-30.0.01.tsv - matched-calls/varlociraptor-delly/simulated-bwa.INS.1-30.0.05.tsv - matched-calls/varlociraptor-delly/simulated-bwa.INS.1-30.0.1.tsv - matched-calls/varlociraptor-delly/simulated-bwa.INS.1-30.0.2.tsv - matched-calls/varlociraptor-delly/simulated-bwa.INS.1-30.0.3.tsv - matched-calls/varlociraptor-delly/simulated-bwa.INS.1-30.0.4.tsv - matched-calls/varlociraptor-delly/simulated-bwa.INS.1-30.0.5.tsv - matched-calls/varlociraptor-delly/simulated-bwa.INS.1-30.0.6.tsv - matched-calls/varlociraptor-delly/simulated-bwa.INS.1-30.0.7.tsv - matched-calls/varlociraptor-delly/simulated-bwa.INS.1-30.0.8.tsv - matched-calls/varlociraptor-delly/simulated-bwa.INS.1-30.0.9.tsv - matched-calls/varlociraptor-delly/simulated-bwa.INS.30-100.1e-05.tsv - matched-calls/varlociraptor-delly/simulated-bwa.INS.30-100.0.0001.tsv - matched-calls/varlociraptor-delly/simulated-bwa.INS.30-100.0.001.tsv - matched-calls/varlociraptor-delly/simulated-bwa.INS.30-100.0.01.tsv - matched-calls/varlociraptor-delly/simulated-bwa.INS.30-100.0.05.tsv - matched-calls/varlociraptor-delly/simulated-bwa.INS.30-100.0.1.tsv - matched-calls/varlociraptor-delly/simulated-bwa.INS.30-100.0.2.tsv - matched-calls/varlociraptor-delly/simulated-bwa.INS.30-100.0.3.tsv - matched-calls/varlociraptor-delly/simulated-bwa.INS.30-100.0.4.tsv - matched-calls/varlociraptor-delly/simulated-bwa.INS.30-100.0.5.tsv - matched-calls/varlociraptor-delly/simulated-bwa.INS.30-100.0.6.tsv - matched-calls/varlociraptor-delly/simulated-bwa.INS.30-100.0.7.tsv - matched-calls/varlociraptor-delly/simulated-bwa.INS.30-100.0.8.tsv - matched-calls/varlociraptor-delly/simulated-bwa.INS.30-100.0.9.tsv - matched-calls/varlociraptor-lancet/simulated-bwa.INS.1-30.1e-05.tsv - matched-calls/varlociraptor-lancet/simulated-bwa.INS.1-30.0.0001.tsv - matched-calls/varlociraptor-lancet/simulated-bwa.INS.1-30.0.001.tsv - matched-calls/varlociraptor-lancet/simulated-bwa.INS.1-30.0.01.tsv - matched-calls/varlociraptor-lancet/simulated-bwa.INS.1-30.0.05.tsv - matched-calls/varlociraptor-lancet/simulated-bwa.INS.1-30.0.1.tsv - matched-calls/varlociraptor-lancet/simulated-bwa.INS.1-30.0.2.tsv - matched-calls/varlociraptor-lancet/simulated-bwa.INS.1-30.0.3.tsv - matched-calls/varlociraptor-lancet/simulated-bwa.INS.1-30.0.4.tsv - matched-calls/varlociraptor-lancet/simulated-bwa.INS.1-30.0.5.tsv - matched-calls/varlociraptor-lancet/simulated-bwa.INS.1-30.0.6.tsv - matched-calls/varlociraptor-lancet/simulated-bwa.INS.1-30.0.7.tsv - matched-calls/varlociraptor-lancet/simulated-bwa.INS.1-30.0.8.tsv - matched-calls/varlociraptor-lancet/simulated-bwa.INS.1-30.0.9.tsv - matched-calls/varlociraptor-lancet/simulated-bwa.INS.30-100.1e-05.tsv - matched-calls/varlociraptor-lancet/simulated-bwa.INS.30-100.0.0001.tsv - matched-calls/varlociraptor-lancet/simulated-bwa.INS.30-100.0.001.tsv - matched-calls/varlociraptor-lancet/simulated-bwa.INS.30-100.0.01.tsv - matched-calls/varlociraptor-lancet/simulated-bwa.INS.30-100.0.05.tsv - matched-calls/varlociraptor-lancet/simulated-bwa.INS.30-100.0.1.tsv - matched-calls/varlociraptor-lancet/simulated-bwa.INS.30-100.0.2.tsv - matched-calls/varlociraptor-lancet/simulated-bwa.INS.30-100.0.3.tsv - matched-calls/varlociraptor-lancet/simulated-bwa.INS.30-100.0.4.tsv - matched-calls/varlociraptor-lancet/simulated-bwa.INS.30-100.0.5.tsv - matched-calls/varlociraptor-lancet/simulated-bwa.INS.30-100.0.6.tsv - matched-calls/varlociraptor-lancet/simulated-bwa.INS.30-100.0.7.tsv - matched-calls/varlociraptor-lancet/simulated-bwa.INS.30-100.0.8.tsv - matched-calls/varlociraptor-lancet/simulated-bwa.INS.30-100.0.9.tsv - matched-calls/varlociraptor-manta/simulated-bwa.INS.1-30.1e-05.tsv - matched-calls/varlociraptor-manta/simulated-bwa.INS.1-30.0.0001.tsv - matched-calls/varlociraptor-manta/simulated-bwa.INS.1-30.0.001.tsv - matched-calls/varlociraptor-manta/simulated-bwa.INS.1-30.0.01.tsv - matched-calls/varlociraptor-manta/simulated-bwa.INS.1-30.0.05.tsv - matched-calls/varlociraptor-manta/simulated-bwa.INS.1-30.0.1.tsv - matched-calls/varlociraptor-manta/simulated-bwa.INS.1-30.0.2.tsv - matched-calls/varlociraptor-manta/simulated-bwa.INS.1-30.0.3.tsv - matched-calls/varlociraptor-manta/simulated-bwa.INS.1-30.0.4.tsv - matched-calls/varlociraptor-manta/simulated-bwa.INS.1-30.0.5.tsv - matched-calls/varlociraptor-manta/simulated-bwa.INS.1-30.0.6.tsv - matched-calls/varlociraptor-manta/simulated-bwa.INS.1-30.0.7.tsv - matched-calls/varlociraptor-manta/simulated-bwa.INS.1-30.0.8.tsv - matched-calls/varlociraptor-manta/simulated-bwa.INS.1-30.0.9.tsv - matched-calls/varlociraptor-manta/simulated-bwa.INS.30-100.1e-05.tsv - matched-calls/varlociraptor-manta/simulated-bwa.INS.30-100.0.0001.tsv - matched-calls/varlociraptor-manta/simulated-bwa.INS.30-100.0.001.tsv - matched-calls/varlociraptor-manta/simulated-bwa.INS.30-100.0.01.tsv - matched-calls/varlociraptor-manta/simulated-bwa.INS.30-100.0.05.tsv - matched-calls/varlociraptor-manta/simulated-bwa.INS.30-100.0.1.tsv - matched-calls/varlociraptor-manta/simulated-bwa.INS.30-100.0.2.tsv - matched-calls/varlociraptor-manta/simulated-bwa.INS.30-100.0.3.tsv - matched-calls/varlociraptor-manta/simulated-bwa.INS.30-100.0.4.tsv - matched-calls/varlociraptor-manta/simulated-bwa.INS.30-100.0.5.tsv - matched-calls/varlociraptor-manta/simulated-bwa.INS.30-100.0.6.tsv - matched-calls/varlociraptor-manta/simulated-bwa.INS.30-100.0.7.tsv - matched-calls/varlociraptor-manta/simulated-bwa.INS.30-100.0.8.tsv - matched-calls/varlociraptor-manta/simulated-bwa.INS.30-100.0.9.tsv - matched-calls/varlociraptor-strelka/simulated-bwa.INS.1-30.1e-05.tsv - matched-calls/varlociraptor-strelka/simulated-bwa.INS.1-30.0.0001.tsv - matched-calls/varlociraptor-strelka/simulated-bwa.INS.1-30.0.001.tsv - matched-calls/varlociraptor-strelka/simulated-bwa.INS.1-30.0.01.tsv - matched-calls/varlociraptor-strelka/simulated-bwa.INS.1-30.0.05.tsv - matched-calls/varlociraptor-strelka/simulated-bwa.INS.1-30.0.1.tsv - matched-calls/varlociraptor-strelka/simulated-bwa.INS.1-30.0.2.tsv - matched-calls/varlociraptor-strelka/simulated-bwa.INS.1-30.0.3.tsv - matched-calls/varlociraptor-strelka/simulated-bwa.INS.1-30.0.4.tsv - matched-calls/varlociraptor-strelka/simulated-bwa.INS.1-30.0.5.tsv - matched-calls/varlociraptor-strelka/simulated-bwa.INS.1-30.0.6.tsv - matched-calls/varlociraptor-strelka/simulated-bwa.INS.1-30.0.7.tsv - matched-calls/varlociraptor-strelka/simulated-bwa.INS.1-30.0.8.tsv - matched-calls/varlociraptor-strelka/simulated-bwa.INS.1-30.0.9.tsv - matched-calls/varlociraptor-strelka/simulated-bwa.INS.30-100.1e-05.tsv - matched-calls/varlociraptor-strelka/simulated-bwa.INS.30-100.0.0001.tsv - matched-calls/varlociraptor-strelka/simulated-bwa.INS.30-100.0.001.tsv - matched-calls/varlociraptor-strelka/simulated-bwa.INS.30-100.0.01.tsv - matched-calls/varlociraptor-strelka/simulated-bwa.INS.30-100.0.05.tsv - matched-calls/varlociraptor-strelka/simulated-bwa.INS.30-100.0.1.tsv - matched-calls/varlociraptor-strelka/simulated-bwa.INS.30-100.0.2.tsv - matched-calls/varlociraptor-strelka/simulated-bwa.INS.30-100.0.3.tsv - matched-calls/varlociraptor-strelka/simulated-bwa.INS.30-100.0.4.tsv - matched-calls/varlociraptor-strelka/simulated-bwa.INS.30-100.0.5.tsv - matched-calls/varlociraptor-strelka/simulated-bwa.INS.30-100.0.6.tsv - matched-calls/varlociraptor-strelka/simulated-bwa.INS.30-100.0.7.tsv - matched-calls/varlociraptor-strelka/simulated-bwa.INS.30-100.0.8.tsv - matched-calls/varlociraptor-strelka/simulated-bwa.INS.30-100.0.9.tsv - matched-calls/varlociraptor-delly/simulated-bwa.DEL.1-30.1e-05.tsv - matched-calls/varlociraptor-delly/simulated-bwa.DEL.1-30.0.0001.tsv - matched-calls/varlociraptor-delly/simulated-bwa.DEL.1-30.0.001.tsv - matched-calls/varlociraptor-delly/simulated-bwa.DEL.1-30.0.01.tsv - matched-calls/varlociraptor-delly/simulated-bwa.DEL.1-30.0.05.tsv - matched-calls/varlociraptor-delly/simulated-bwa.DEL.1-30.0.1.tsv - matched-calls/varlociraptor-delly/simulated-bwa.DEL.1-30.0.2.tsv - matched-calls/varlociraptor-delly/simulated-bwa.DEL.1-30.0.3.tsv - matched-calls/varlociraptor-delly/simulated-bwa.DEL.1-30.0.4.tsv - matched-calls/varlociraptor-delly/simulated-bwa.DEL.1-30.0.5.tsv - matched-calls/varlociraptor-delly/simulated-bwa.DEL.1-30.0.6.tsv - matched-calls/varlociraptor-delly/simulated-bwa.DEL.1-30.0.7.tsv - matched-calls/varlociraptor-delly/simulated-bwa.DEL.1-30.0.8.tsv - matched-calls/varlociraptor-delly/simulated-bwa.DEL.1-30.0.9.tsv - matched-calls/varlociraptor-delly/simulated-bwa.DEL.30-50.1e-05.tsv - matched-calls/varlociraptor-delly/simulated-bwa.DEL.30-50.0.0001.tsv - matched-calls/varlociraptor-delly/simulated-bwa.DEL.30-50.0.001.tsv - matched-calls/varlociraptor-delly/simulated-bwa.DEL.30-50.0.01.tsv - matched-calls/varlociraptor-delly/simulated-bwa.DEL.30-50.0.05.tsv - matched-calls/varlociraptor-delly/simulated-bwa.DEL.30-50.0.1.tsv - matched-calls/varlociraptor-delly/simulated-bwa.DEL.30-50.0.2.tsv - matched-calls/varlociraptor-delly/simulated-bwa.DEL.30-50.0.3.tsv - matched-calls/varlociraptor-delly/simulated-bwa.DEL.30-50.0.4.tsv - matched-calls/varlociraptor-delly/simulated-bwa.DEL.30-50.0.5.tsv - matched-calls/varlociraptor-delly/simulated-bwa.DEL.30-50.0.6.tsv - matched-calls/varlociraptor-delly/simulated-bwa.DEL.30-50.0.7.tsv - matched-calls/varlociraptor-delly/simulated-bwa.DEL.30-50.0.8.tsv - matched-calls/varlociraptor-delly/simulated-bwa.DEL.30-50.0.9.tsv - matched-calls/varlociraptor-delly/simulated-bwa.DEL.50-100.1e-05.tsv - matched-calls/varlociraptor-delly/simulated-bwa.DEL.50-100.0.0001.tsv - matched-calls/varlociraptor-delly/simulated-bwa.DEL.50-100.0.001.tsv - matched-calls/varlociraptor-delly/simulated-bwa.DEL.50-100.0.01.tsv - matched-calls/varlociraptor-delly/simulated-bwa.DEL.50-100.0.05.tsv - matched-calls/varlociraptor-delly/simulated-bwa.DEL.50-100.0.1.tsv - matched-calls/varlociraptor-delly/simulated-bwa.DEL.50-100.0.2.tsv - matched-calls/varlociraptor-delly/simulated-bwa.DEL.50-100.0.3.tsv - matched-calls/varlociraptor-delly/simulated-bwa.DEL.50-100.0.4.tsv - matched-calls/varlociraptor-delly/simulated-bwa.DEL.50-100.0.5.tsv - matched-calls/varlociraptor-delly/simulated-bwa.DEL.50-100.0.6.tsv - matched-calls/varlociraptor-delly/simulated-bwa.DEL.50-100.0.7.tsv - matched-calls/varlociraptor-delly/simulated-bwa.DEL.50-100.0.8.tsv - matched-calls/varlociraptor-delly/simulated-bwa.DEL.50-100.0.9.tsv - matched-calls/varlociraptor-delly/simulated-bwa.DEL.100-250.1e-05.tsv - matched-calls/varlociraptor-delly/simulated-bwa.DEL.100-250.0.0001.tsv - matched-calls/varlociraptor-delly/simulated-bwa.DEL.100-250.0.001.tsv - matched-calls/varlociraptor-delly/simulated-bwa.DEL.100-250.0.01.tsv - matched-calls/varlociraptor-delly/simulated-bwa.DEL.100-250.0.05.tsv - matched-calls/varlociraptor-delly/simulated-bwa.DEL.100-250.0.1.tsv - matched-calls/varlociraptor-delly/simulated-bwa.DEL.100-250.0.2.tsv - matched-calls/varlociraptor-delly/simulated-bwa.DEL.100-250.0.3.tsv - matched-calls/varlociraptor-delly/simulated-bwa.DEL.100-250.0.4.tsv - matched-calls/varlociraptor-delly/simulated-bwa.DEL.100-250.0.5.tsv - matched-calls/varlociraptor-delly/simulated-bwa.DEL.100-250.0.6.tsv - matched-calls/varlociraptor-delly/simulated-bwa.DEL.100-250.0.7.tsv - matched-calls/varlociraptor-delly/simulated-bwa.DEL.100-250.0.8.tsv - matched-calls/varlociraptor-delly/simulated-bwa.DEL.100-250.0.9.tsv - matched-calls/varlociraptor-lancet/simulated-bwa.DEL.1-30.1e-05.tsv - matched-calls/varlociraptor-lancet/simulated-bwa.DEL.1-30.0.0001.tsv - matched-calls/varlociraptor-lancet/simulated-bwa.DEL.1-30.0.001.tsv - matched-calls/varlociraptor-lancet/simulated-bwa.DEL.1-30.0.01.tsv - matched-calls/varlociraptor-lancet/simulated-bwa.DEL.1-30.0.05.tsv - matched-calls/varlociraptor-lancet/simulated-bwa.DEL.1-30.0.1.tsv - matched-calls/varlociraptor-lancet/simulated-bwa.DEL.1-30.0.2.tsv - matched-calls/varlociraptor-lancet/simulated-bwa.DEL.1-30.0.3.tsv - matched-calls/varlociraptor-lancet/simulated-bwa.DEL.1-30.0.4.tsv - matched-calls/varlociraptor-lancet/simulated-bwa.DEL.1-30.0.5.tsv - matched-calls/varlociraptor-lancet/simulated-bwa.DEL.1-30.0.6.tsv - matched-calls/varlociraptor-lancet/simulated-bwa.DEL.1-30.0.7.tsv - matched-calls/varlociraptor-lancet/simulated-bwa.DEL.1-30.0.8.tsv - matched-calls/varlociraptor-lancet/simulated-bwa.DEL.1-30.0.9.tsv - matched-calls/varlociraptor-lancet/simulated-bwa.DEL.30-50.1e-05.tsv - matched-calls/varlociraptor-lancet/simulated-bwa.DEL.30-50.0.0001.tsv - matched-calls/varlociraptor-lancet/simulated-bwa.DEL.30-50.0.001.tsv - matched-calls/varlociraptor-lancet/simulated-bwa.DEL.30-50.0.01.tsv - matched-calls/varlociraptor-lancet/simulated-bwa.DEL.30-50.0.05.tsv - matched-calls/varlociraptor-lancet/simulated-bwa.DEL.30-50.0.1.tsv - matched-calls/varlociraptor-lancet/simulated-bwa.DEL.30-50.0.2.tsv - matched-calls/varlociraptor-lancet/simulated-bwa.DEL.30-50.0.3.tsv - matched-calls/varlociraptor-lancet/simulated-bwa.DEL.30-50.0.4.tsv - matched-calls/varlociraptor-lancet/simulated-bwa.DEL.30-50.0.5.tsv - matched-calls/varlociraptor-lancet/simulated-bwa.DEL.30-50.0.6.tsv - matched-calls/varlociraptor-lancet/simulated-bwa.DEL.30-50.0.7.tsv - matched-calls/varlociraptor-lancet/simulated-bwa.DEL.30-50.0.8.tsv - matched-calls/varlociraptor-lancet/simulated-bwa.DEL.30-50.0.9.tsv - matched-calls/varlociraptor-lancet/simulated-bwa.DEL.50-100.1e-05.tsv - matched-calls/varlociraptor-lancet/simulated-bwa.DEL.50-100.0.0001.tsv - matched-calls/varlociraptor-lancet/simulated-bwa.DEL.50-100.0.001.tsv - matched-calls/varlociraptor-lancet/simulated-bwa.DEL.50-100.0.01.tsv - matched-calls/varlociraptor-lancet/simulated-bwa.DEL.50-100.0.05.tsv - matched-calls/varlociraptor-lancet/simulated-bwa.DEL.50-100.0.1.tsv - matched-calls/varlociraptor-lancet/simulated-bwa.DEL.50-100.0.2.tsv - matched-calls/varlociraptor-lancet/simulated-bwa.DEL.50-100.0.3.tsv - matched-calls/varlociraptor-lancet/simulated-bwa.DEL.50-100.0.4.tsv - matched-calls/varlociraptor-lancet/simulated-bwa.DEL.50-100.0.5.tsv - matched-calls/varlociraptor-lancet/simulated-bwa.DEL.50-100.0.6.tsv - matched-calls/varlociraptor-lancet/simulated-bwa.DEL.50-100.0.7.tsv - matched-calls/varlociraptor-lancet/simulated-bwa.DEL.50-100.0.8.tsv - matched-calls/varlociraptor-lancet/simulated-bwa.DEL.50-100.0.9.tsv - matched-calls/varlociraptor-lancet/simulated-bwa.DEL.100-250.1e-05.tsv - matched-calls/varlociraptor-lancet/simulated-bwa.DEL.100-250.0.0001.tsv - matched-calls/varlociraptor-lancet/simulated-bwa.DEL.100-250.0.001.tsv - matched-calls/varlociraptor-lancet/simulated-bwa.DEL.100-250.0.01.tsv - matched-calls/varlociraptor-lancet/simulated-bwa.DEL.100-250.0.05.tsv - matched-calls/varlociraptor-lancet/simulated-bwa.DEL.100-250.0.1.tsv - matched-calls/varlociraptor-lancet/simulated-bwa.DEL.100-250.0.2.tsv - matched-calls/varlociraptor-lancet/simulated-bwa.DEL.100-250.0.3.tsv - matched-calls/varlociraptor-lancet/simulated-bwa.DEL.100-250.0.4.tsv - matched-calls/varlociraptor-lancet/simulated-bwa.DEL.100-250.0.5.tsv - matched-calls/varlociraptor-lancet/simulated-bwa.DEL.100-250.0.6.tsv - matched-calls/varlociraptor-lancet/simulated-bwa.DEL.100-250.0.7.tsv - matched-calls/varlociraptor-lancet/simulated-bwa.DEL.100-250.0.8.tsv - matched-calls/varlociraptor-lancet/simulated-bwa.DEL.100-250.0.9.tsv - matched-calls/varlociraptor-manta/simulated-bwa.DEL.1-30.1e-05.tsv - matched-calls/varlociraptor-manta/simulated-bwa.DEL.1-30.0.0001.tsv - matched-calls/varlociraptor-manta/simulated-bwa.DEL.1-30.0.001.tsv - matched-calls/varlociraptor-manta/simulated-bwa.DEL.1-30.0.01.tsv - matched-calls/varlociraptor-manta/simulated-bwa.DEL.1-30.0.05.tsv - matched-calls/varlociraptor-manta/simulated-bwa.DEL.1-30.0.1.tsv - matched-calls/varlociraptor-manta/simulated-bwa.DEL.1-30.0.2.tsv - matched-calls/varlociraptor-manta/simulated-bwa.DEL.1-30.0.3.tsv - matched-calls/varlociraptor-manta/simulated-bwa.DEL.1-30.0.4.tsv - matched-calls/varlociraptor-manta/simulated-bwa.DEL.1-30.0.5.tsv - matched-calls/varlociraptor-manta/simulated-bwa.DEL.1-30.0.6.tsv - matched-calls/varlociraptor-manta/simulated-bwa.DEL.1-30.0.7.tsv - matched-calls/varlociraptor-manta/simulated-bwa.DEL.1-30.0.8.tsv - matched-calls/varlociraptor-manta/simulated-bwa.DEL.1-30.0.9.tsv - matched-calls/varlociraptor-manta/simulated-bwa.DEL.30-50.1e-05.tsv - matched-calls/varlociraptor-manta/simulated-bwa.DEL.30-50.0.0001.tsv - matched-calls/varlociraptor-manta/simulated-bwa.DEL.30-50.0.001.tsv - matched-calls/varlociraptor-manta/simulated-bwa.DEL.30-50.0.01.tsv - matched-calls/varlociraptor-manta/simulated-bwa.DEL.30-50.0.05.tsv - matched-calls/varlociraptor-manta/simulated-bwa.DEL.30-50.0.1.tsv - matched-calls/varlociraptor-manta/simulated-bwa.DEL.30-50.0.2.tsv - matched-calls/varlociraptor-manta/simulated-bwa.DEL.30-50.0.3.tsv - matched-calls/varlociraptor-manta/simulated-bwa.DEL.30-50.0.4.tsv - matched-calls/varlociraptor-manta/simulated-bwa.DEL.30-50.0.5.tsv - matched-calls/varlociraptor-manta/simulated-bwa.DEL.30-50.0.6.tsv - matched-calls/varlociraptor-manta/simulated-bwa.DEL.30-50.0.7.tsv - matched-calls/varlociraptor-manta/simulated-bwa.DEL.30-50.0.8.tsv - matched-calls/varlociraptor-manta/simulated-bwa.DEL.30-50.0.9.tsv - matched-calls/varlociraptor-manta/simulated-bwa.DEL.50-100.1e-05.tsv - matched-calls/varlociraptor-manta/simulated-bwa.DEL.50-100.0.0001.tsv - matched-calls/varlociraptor-manta/simulated-bwa.DEL.50-100.0.001.tsv - matched-calls/varlociraptor-manta/simulated-bwa.DEL.50-100.0.01.tsv - matched-calls/varlociraptor-manta/simulated-bwa.DEL.50-100.0.05.tsv - matched-calls/varlociraptor-manta/simulated-bwa.DEL.50-100.0.1.tsv - matched-calls/varlociraptor-manta/simulated-bwa.DEL.50-100.0.2.tsv - matched-calls/varlociraptor-manta/simulated-bwa.DEL.50-100.0.3.tsv - matched-calls/varlociraptor-manta/simulated-bwa.DEL.50-100.0.4.tsv - matched-calls/varlociraptor-manta/simulated-bwa.DEL.50-100.0.5.tsv - matched-calls/varlociraptor-manta/simulated-bwa.DEL.50-100.0.6.tsv - matched-calls/varlociraptor-manta/simulated-bwa.DEL.50-100.0.7.tsv - matched-calls/varlociraptor-manta/simulated-bwa.DEL.50-100.0.8.tsv - matched-calls/varlociraptor-manta/simulated-bwa.DEL.50-100.0.9.tsv - matched-calls/varlociraptor-manta/simulated-bwa.DEL.100-250.1e-05.tsv - matched-calls/varlociraptor-manta/simulated-bwa.DEL.100-250.0.0001.tsv - matched-calls/varlociraptor-manta/simulated-bwa.DEL.100-250.0.001.tsv - matched-calls/varlociraptor-manta/simulated-bwa.DEL.100-250.0.01.tsv - matched-calls/varlociraptor-manta/simulated-bwa.DEL.100-250.0.05.tsv - matched-calls/varlociraptor-manta/simulated-bwa.DEL.100-250.0.1.tsv - matched-calls/varlociraptor-manta/simulated-bwa.DEL.100-250.0.2.tsv - matched-calls/varlociraptor-manta/simulated-bwa.DEL.100-250.0.3.tsv - matched-calls/varlociraptor-manta/simulated-bwa.DEL.100-250.0.4.tsv - matched-calls/varlociraptor-manta/simulated-bwa.DEL.100-250.0.5.tsv - matched-calls/varlociraptor-manta/simulated-bwa.DEL.100-250.0.6.tsv - matched-calls/varlociraptor-manta/simulated-bwa.DEL.100-250.0.7.tsv - matched-calls/varlociraptor-manta/simulated-bwa.DEL.100-250.0.8.tsv - matched-calls/varlociraptor-manta/simulated-bwa.DEL.100-250.0.9.tsv - matched-calls/varlociraptor-strelka/simulated-bwa.DEL.1-30.1e-05.tsv - matched-calls/varlociraptor-strelka/simulated-bwa.DEL.1-30.0.0001.tsv - matched-calls/varlociraptor-strelka/simulated-bwa.DEL.1-30.0.001.tsv - matched-calls/varlociraptor-strelka/simulated-bwa.DEL.1-30.0.01.tsv - matched-calls/varlociraptor-strelka/simulated-bwa.DEL.1-30.0.05.tsv - matched-calls/varlociraptor-strelka/simulated-bwa.DEL.1-30.0.1.tsv - matched-calls/varlociraptor-strelka/simulated-bwa.DEL.1-30.0.2.tsv - matched-calls/varlociraptor-strelka/simulated-bwa.DEL.1-30.0.3.tsv - matched-calls/varlociraptor-strelka/simulated-bwa.DEL.1-30.0.4.tsv - matched-calls/varlociraptor-strelka/simulated-bwa.DEL.1-30.0.5.tsv - matched-calls/varlociraptor-strelka/simulated-bwa.DEL.1-30.0.6.tsv - matched-calls/varlociraptor-strelka/simulated-bwa.DEL.1-30.0.7.tsv - matched-calls/varlociraptor-strelka/simulated-bwa.DEL.1-30.0.8.tsv - matched-calls/varlociraptor-strelka/simulated-bwa.DEL.1-30.0.9.tsv - matched-calls/varlociraptor-strelka/simulated-bwa.DEL.30-50.1e-05.tsv - matched-calls/varlociraptor-strelka/simulated-bwa.DEL.30-50.0.0001.tsv - matched-calls/varlociraptor-strelka/simulated-bwa.DEL.30-50.0.001.tsv - matched-calls/varlociraptor-strelka/simulated-bwa.DEL.30-50.0.01.tsv - matched-calls/varlociraptor-strelka/simulated-bwa.DEL.30-50.0.05.tsv - matched-calls/varlociraptor-strelka/simulated-bwa.DEL.30-50.0.1.tsv - matched-calls/varlociraptor-strelka/simulated-bwa.DEL.30-50.0.2.tsv - matched-calls/varlociraptor-strelka/simulated-bwa.DEL.30-50.0.3.tsv - matched-calls/varlociraptor-strelka/simulated-bwa.DEL.30-50.0.4.tsv - matched-calls/varlociraptor-strelka/simulated-bwa.DEL.30-50.0.5.tsv - matched-calls/varlociraptor-strelka/simulated-bwa.DEL.30-50.0.6.tsv - matched-calls/varlociraptor-strelka/simulated-bwa.DEL.30-50.0.7.tsv - matched-calls/varlociraptor-strelka/simulated-bwa.DEL.30-50.0.8.tsv - matched-calls/varlociraptor-strelka/simulated-bwa.DEL.30-50.0.9.tsv - matched-calls/varlociraptor-strelka/simulated-bwa.DEL.50-100.1e-05.tsv - matched-calls/varlociraptor-strelka/simulated-bwa.DEL.50-100.0.0001.tsv - matched-calls/varlociraptor-strelka/simulated-bwa.DEL.50-100.0.001.tsv - matched-calls/varlociraptor-strelka/simulated-bwa.DEL.50-100.0.01.tsv - matched-calls/varlociraptor-strelka/simulated-bwa.DEL.50-100.0.05.tsv - matched-calls/varlociraptor-strelka/simulated-bwa.DEL.50-100.0.1.tsv - matched-calls/varlociraptor-strelka/simulated-bwa.DEL.50-100.0.2.tsv - matched-calls/varlociraptor-strelka/simulated-bwa.DEL.50-100.0.3.tsv - matched-calls/varlociraptor-strelka/simulated-bwa.DEL.50-100.0.4.tsv - matched-calls/varlociraptor-strelka/simulated-bwa.DEL.50-100.0.5.tsv - matched-calls/varlociraptor-strelka/simulated-bwa.DEL.50-100.0.6.tsv - matched-calls/varlociraptor-strelka/simulated-bwa.DEL.50-100.0.7.tsv - matched-calls/varlociraptor-strelka/simulated-bwa.DEL.50-100.0.8.tsv - matched-calls/varlociraptor-strelka/simulated-bwa.DEL.50-100.0.9.tsv - matched-calls/varlociraptor-strelka/simulated-bwa.DEL.100-250.1e-05.tsv - matched-calls/varlociraptor-strelka/simulated-bwa.DEL.100-250.0.0001.tsv - matched-calls/varlociraptor-strelka/simulated-bwa.DEL.100-250.0.001.tsv - matched-calls/varlociraptor-strelka/simulated-bwa.DEL.100-250.0.01.tsv - matched-calls/varlociraptor-strelka/simulated-bwa.DEL.100-250.0.05.tsv - matched-calls/varlociraptor-strelka/simulated-bwa.DEL.100-250.0.1.tsv - matched-calls/varlociraptor-strelka/simulated-bwa.DEL.100-250.0.2.tsv - matched-calls/varlociraptor-strelka/simulated-bwa.DEL.100-250.0.3.tsv - matched-calls/varlociraptor-strelka/simulated-bwa.DEL.100-250.0.4.tsv - matched-calls/varlociraptor-strelka/simulated-bwa.DEL.100-250.0.5.tsv - matched-calls/varlociraptor-strelka/simulated-bwa.DEL.100-250.0.6.tsv - matched-calls/varlociraptor-strelka/simulated-bwa.DEL.100-250.0.7.tsv - matched-calls/varlociraptor-strelka/simulated-bwa.DEL.100-250.0.8.tsv - matched-calls/varlociraptor-strelka/simulated-bwa.DEL.100-250.0.9.tsv - matched-calls/varlociraptor-delly/simulated-bwa.INS.1-250.1.0.tsv - matched-calls/varlociraptor-lancet/simulated-bwa.INS.1-250.1.0.tsv - matched-calls/varlociraptor-manta/simulated-bwa.INS.1-250.1.0.tsv - matched-calls/varlociraptor-strelka/simulated-bwa.INS.1-250.1.0.tsv - matched-calls/varlociraptor-delly/simulated-bwa.DEL.1-250.1.0.tsv - matched-calls/varlociraptor-lancet/simulated-bwa.DEL.1-250.1.0.tsv - matched-calls/varlociraptor-manta/simulated-bwa.DEL.1-250.1.0.tsv - matched-calls/varlociraptor-strelka/simulated-bwa.DEL.1-250.1.0.tsv |  | - rust-bio-tools =0.5.0 - bedtools =2.27.1 - bcftools =1.8 | ``` 1 ``` \| ``` rbt vcf-to-txt {params.gt} {params.tags} --info MATCHING < {input} > {output} ``` |
| other\_calls\_to\_tsv | 8 | - matched-calls/default-lancet/simulated-bwa.all.tsv - matched-calls/default-manta/simulated-bwa.all.tsv - matched-calls/default-strelka/simulated-bwa.all.tsv - matched-calls/adhoc-delly/simulated-bwa.all.tsv - matched-calls/adhoc-lancet/simulated-bwa.all.tsv - matched-calls/adhoc-manta/simulated-bwa.all.tsv - matched-calls/adhoc-strelka/simulated-bwa.all.tsv - matched-calls/adhoc-bpi/simulated-bwa.all.tsv |  | - rust-bio-tools =0.2.5 - bedtools =2.27.1 - bcftools =1.8 | ``` 1 ``` \| ``` rbt vcf-to-txt {params.gt} {params.tags} --info MATCHING < {input} > {output} ``` |
| concordance\_to\_tsv | 60 | - concordance/varlociraptor-delly-0.9/colo1.0-vs-1.tsv - concordance/varlociraptor-delly-0.9/colo1.1-vs-2.tsv - concordance/varlociraptor-delly-0.9/colo1.2-vs-0.tsv - concordance/varlociraptor-delly-0.9/colo1.2-vs-3.tsv - concordance/varlociraptor-delly-0.9/colo1.3-vs-0.tsv - concordance/varlociraptor-delly-0.9/colo1.3-vs-1.tsv - concordance/varlociraptor-lancet-0.9/colo1.0-vs-1.tsv - concordance/varlociraptor-lancet-0.9/colo1.1-vs-2.tsv - concordance/varlociraptor-lancet-0.9/colo1.2-vs-0.tsv - concordance/varlociraptor-lancet-0.9/colo1.2-vs-3.tsv - concordance/varlociraptor-lancet-0.9/colo1.3-vs-0.tsv - concordance/varlociraptor-lancet-0.9/colo1.3-vs-1.tsv - concordance/varlociraptor-manta-0.9/colo1.0-vs-1.tsv - concordance/varlociraptor-manta-0.9/colo1.1-vs-2.tsv - concordance/varlociraptor-manta-0.9/colo1.2-vs-0.tsv - concordance/varlociraptor-manta-0.9/colo1.2-vs-3.tsv - concordance/varlociraptor-manta-0.9/colo1.3-vs-0.tsv - concordance/varlociraptor-manta-0.9/colo1.3-vs-1.tsv - concordance/varlociraptor-strelka-0.9/colo1.0-vs-1.tsv - concordance/varlociraptor-strelka-0.9/colo1.1-vs-2.tsv - concordance/varlociraptor-strelka-0.9/colo1.2-vs-0.tsv - concordance/varlociraptor-strelka-0.9/colo1.2-vs-3.tsv - concordance/varlociraptor-strelka-0.9/colo1.3-vs-0.tsv - concordance/varlociraptor-strelka-0.9/colo1.3-vs-1.tsv - concordance/varlociraptor-bpi-0.9/colo1.0-vs-1.tsv - concordance/varlociraptor-bpi-0.9/colo1.1-vs-2.tsv - concordance/varlociraptor-bpi-0.9/colo1.2-vs-0.tsv - concordance/varlociraptor-bpi-0.9/colo1.2-vs-3.tsv - concordance/varlociraptor-bpi-0.9/colo1.3-vs-0.tsv - concordance/varlociraptor-bpi-0.9/colo1.3-vs-1.tsv - concordance/varlociraptor-delly-0.98/colo1.0-vs-1.tsv - concordance/varlociraptor-delly-0.98/colo1.1-vs-2.tsv - concordance/varlociraptor-delly-0.98/colo1.2-vs-0.tsv - concordance/varlociraptor-delly-0.98/colo1.2-vs-3.tsv - concordance/varlociraptor-delly-0.98/colo1.3-vs-0.tsv - concordance/varlociraptor-delly-0.98/colo1.3-vs-1.tsv - concordance/varlociraptor-lancet-0.98/colo1.0-vs-1.tsv - concordance/varlociraptor-lancet-0.98/colo1.1-vs-2.tsv - concordance/varlociraptor-lancet-0.98/colo1.2-vs-0.tsv - concordance/varlociraptor-lancet-0.98/colo1.2-vs-3.tsv - concordance/varlociraptor-lancet-0.98/colo1.3-vs-0.tsv - concordance/varlociraptor-lancet-0.98/colo1.3-vs-1.tsv - concordance/varlociraptor-manta-0.98/colo1.0-vs-1.tsv - concordance/varlociraptor-manta-0.98/colo1.1-vs-2.tsv - concordance/varlociraptor-manta-0.98/colo1.2-vs-0.tsv - concordance/varlociraptor-manta-0.98/colo1.2-vs-3.tsv - concordance/varlociraptor-manta-0.98/colo1.3-vs-0.tsv - concordance/varlociraptor-manta-0.98/colo1.3-vs-1.tsv - concordance/varlociraptor-strelka-0.98/colo1.0-vs-1.tsv - concordance/varlociraptor-strelka-0.98/colo1.1-vs-2.tsv - concordance/varlociraptor-strelka-0.98/colo1.2-vs-0.tsv - concordance/varlociraptor-strelka-0.98/colo1.2-vs-3.tsv - concordance/varlociraptor-strelka-0.98/colo1.3-vs-0.tsv - concordance/varlociraptor-strelka-0.98/colo1.3-vs-1.tsv - concordance/varlociraptor-bpi-0.98/colo1.0-vs-1.tsv - concordance/varlociraptor-bpi-0.98/colo1.1-vs-2.tsv - concordance/varlociraptor-bpi-0.98/colo1.2-vs-0.tsv - concordance/varlociraptor-bpi-0.98/colo1.2-vs-3.tsv - concordance/varlociraptor-bpi-0.98/colo1.3-vs-0.tsv - concordance/varlociraptor-bpi-0.98/colo1.3-vs-1.tsv |  | - rust-bio-tools =0.5.0 - bedtools =2.27.1 - bcftools =1.8 | ``` 1 ``` \| ``` rbt vcf-to-txt {params.gt} {params.tags} --info MATCHING < {input} > {output} ``` |
| concordance\_to\_tsv | 30 | - concordance/adhoc-delly-default/colo1.0-vs-1.tsv - concordance/adhoc-delly-default/colo1.1-vs-2.tsv - concordance/adhoc-delly-default/colo1.2-vs-0.tsv - concordance/adhoc-delly-default/colo1.2-vs-3.tsv - concordance/adhoc-delly-default/colo1.3-vs-0.tsv - concordance/adhoc-delly-default/colo1.3-vs-1.tsv - concordance/adhoc-lancet-default/colo1.0-vs-1.tsv - concordance/adhoc-lancet-default/colo1.1-vs-2.tsv - concordance/adhoc-lancet-default/colo1.2-vs-0.tsv - concordance/adhoc-lancet-default/colo1.2-vs-3.tsv - concordance/adhoc-lancet-default/colo1.3-vs-0.tsv - concordance/adhoc-lancet-default/colo1.3-vs-1.tsv - concordance/adhoc-manta-default/colo1.0-vs-1.tsv - concordance/adhoc-manta-default/colo1.1-vs-2.tsv - concordance/adhoc-manta-default/colo1.2-vs-0.tsv - concordance/adhoc-manta-default/colo1.2-vs-3.tsv - concordance/adhoc-manta-default/colo1.3-vs-0.tsv - concordance/adhoc-manta-default/colo1.3-vs-1.tsv - concordance/adhoc-strelka-default/colo1.0-vs-1.tsv - concordance/adhoc-strelka-default/colo1.1-vs-2.tsv - concordance/adhoc-strelka-default/colo1.2-vs-0.tsv - concordance/adhoc-strelka-default/colo1.2-vs-3.tsv - concordance/adhoc-strelka-default/colo1.3-vs-0.tsv - concordance/adhoc-strelka-default/colo1.3-vs-1.tsv - concordance/adhoc-bpi-default/colo1.0-vs-1.tsv - concordance/adhoc-bpi-default/colo1.1-vs-2.tsv - concordance/adhoc-bpi-default/colo1.2-vs-0.tsv - concordance/adhoc-bpi-default/colo1.2-vs-3.tsv - concordance/adhoc-bpi-default/colo1.3-vs-0.tsv - concordance/adhoc-bpi-default/colo1.3-vs-1.tsv |  | - rust-bio-tools =0.2.5 - bedtools =2.27.1 - bcftools =1.8 | ``` 1 ``` \| ``` rbt vcf-to-txt {params.gt} {params.tags} --info MATCHING < {input} > {output} ``` |
| varlociraptor\_all\_calls\_to\_tsv | 20 | - varlociraptor-delly/COLO\_829-GSC.all.tsv - varlociraptor-delly/COLO\_829-Ill.all.tsv - varlociraptor-delly/COLO\_829-TGen.all.tsv - varlociraptor-delly/COLO\_829-EBI.all.tsv - varlociraptor-lancet/COLO\_829-GSC.all.tsv - varlociraptor-lancet/COLO\_829-Ill.all.tsv - varlociraptor-lancet/COLO\_829-TGen.all.tsv - varlociraptor-lancet/COLO\_829-EBI.all.tsv - varlociraptor-manta/COLO\_829-GSC.all.tsv - varlociraptor-manta/COLO\_829-Ill.all.tsv - varlociraptor-manta/COLO\_829-TGen.all.tsv - varlociraptor-manta/COLO\_829-EBI.all.tsv - varlociraptor-strelka/COLO\_829-GSC.all.tsv - varlociraptor-strelka/COLO\_829-Ill.all.tsv - varlociraptor-strelka/COLO\_829-TGen.all.tsv - varlociraptor-strelka/COLO\_829-EBI.all.tsv - varlociraptor-bpi/COLO\_829-GSC.all.tsv - varlociraptor-bpi/COLO\_829-Ill.all.tsv - varlociraptor-bpi/COLO\_829-TGen.all.tsv - varlociraptor-bpi/COLO\_829-EBI.all.tsv |  | - rust-bio-tools =0.5.0 - bedtools =2.27.1 - bcftools =1.8 | ``` 1 ``` \| ``` rbt vcf-to-txt {params.gt} {params.tags} < {input} > {output} ``` |
| match\_varlociraptor\_calls | 368 | - matched-calls/varlociraptor-delly/simulated-bwa.INS.1-30.1.0.bcf - matched-calls/varlociraptor-delly/simulated-bwa.INS.30-100.1.0.bcf - matched-calls/varlociraptor-lancet/simulated-bwa.INS.1-30.1.0.bcf - matched-calls/varlociraptor-lancet/simulated-bwa.INS.30-100.1.0.bcf - matched-calls/varlociraptor-manta/simulated-bwa.INS.1-30.1.0.bcf - matched-calls/varlociraptor-manta/simulated-bwa.INS.30-100.1.0.bcf - matched-calls/varlociraptor-strelka/simulated-bwa.INS.1-30.1.0.bcf - matched-calls/varlociraptor-strelka/simulated-bwa.INS.30-100.1.0.bcf - matched-calls/varlociraptor-delly/simulated-bwa.DEL.1-30.1.0.bcf - matched-calls/varlociraptor-delly/simulated-bwa.DEL.30-50.1.0.bcf - matched-calls/varlociraptor-delly/simulated-bwa.DEL.50-100.1.0.bcf - matched-calls/varlociraptor-delly/simulated-bwa.DEL.100-250.1.0.bcf - matched-calls/varlociraptor-lancet/simulated-bwa.DEL.1-30.1.0.bcf - matched-calls/varlociraptor-lancet/simulated-bwa.DEL.30-50.1.0.bcf - matched-calls/varlociraptor-lancet/simulated-bwa.DEL.50-100.1.0.bcf - matched-calls/varlociraptor-lancet/simulated-bwa.DEL.100-250.1.0.bcf - matched-calls/varlociraptor-manta/simulated-bwa.DEL.1-30.1.0.bcf - matched-calls/varlociraptor-manta/simulated-bwa.DEL.30-50.1.0.bcf - matched-calls/varlociraptor-manta/simulated-bwa.DEL.50-100.1.0.bcf - matched-calls/varlociraptor-manta/simulated-bwa.DEL.100-250.1.0.bcf - matched-calls/varlociraptor-strelka/simulated-bwa.DEL.1-30.1.0.bcf - matched-calls/varlociraptor-strelka/simulated-bwa.DEL.30-50.1.0.bcf - matched-calls/varlociraptor-strelka/simulated-bwa.DEL.50-100.1.0.bcf - matched-calls/varlociraptor-strelka/simulated-bwa.DEL.100-250.1.0.bcf - matched-calls/varlociraptor-delly/simulated-bwa.INS.1-30.1e-05.bcf - matched-calls/varlociraptor-delly/simulated-bwa.INS.1-30.0.0001.bcf - matched-calls/varlociraptor-delly/simulated-bwa.INS.1-30.0.001.bcf - matched-calls/varlociraptor-delly/simulated-bwa.INS.1-30.0.01.bcf - matched-calls/varlociraptor-delly/simulated-bwa.INS.1-30.0.05.bcf - matched-calls/varlociraptor-delly/simulated-bwa.INS.1-30.0.1.bcf - matched-calls/varlociraptor-delly/simulated-bwa.INS.1-30.0.2.bcf - matched-calls/varlociraptor-delly/simulated-bwa.INS.1-30.0.3.bcf - matched-calls/varlociraptor-delly/simulated-bwa.INS.1-30.0.4.bcf - matched-calls/varlociraptor-delly/simulated-bwa.INS.1-30.0.5.bcf - matched-calls/varlociraptor-delly/simulated-bwa.INS.1-30.0.6.bcf - matched-calls/varlociraptor-delly/simulated-bwa.INS.1-30.0.7.bcf - matched-calls/varlociraptor-delly/simulated-bwa.INS.1-30.0.8.bcf - matched-calls/varlociraptor-delly/simulated-bwa.INS.1-30.0.9.bcf - matched-calls/varlociraptor-delly/simulated-bwa.INS.30-100.1e-05.bcf - matched-calls/varlociraptor-delly/simulated-bwa.INS.30-100.0.0001.bcf - matched-calls/varlociraptor-delly/simulated-bwa.INS.30-100.0.001.bcf - matched-calls/varlociraptor-delly/simulated-bwa.INS.30-100.0.01.bcf - matched-calls/varlociraptor-delly/simulated-bwa.INS.30-100.0.05.bcf - matched-calls/varlociraptor-delly/simulated-bwa.INS.30-100.0.1.bcf - matched-calls/varlociraptor-delly/simulated-bwa.INS.30-100.0.2.bcf - matched-calls/varlociraptor-delly/simulated-bwa.INS.30-100.0.3.bcf - matched-calls/varlociraptor-delly/simulated-bwa.INS.30-100.0.4.bcf - matched-calls/varlociraptor-delly/simulated-bwa.INS.30-100.0.5.bcf - matched-calls/varlociraptor-delly/simulated-bwa.INS.30-100.0.6.bcf - matched-calls/varlociraptor-delly/simulated-bwa.INS.30-100.0.7.bcf - matched-calls/varlociraptor-delly/simulated-bwa.INS.30-100.0.8.bcf - matched-calls/varlociraptor-delly/simulated-bwa.INS.30-100.0.9.bcf - matched-calls/varlociraptor-lancet/simulated-bwa.INS.1-30.1e-05.bcf - matched-calls/varlociraptor-lancet/simulated-bwa.INS.1-30.0.0001.bcf - matched-calls/varlociraptor-lancet/simulated-bwa.INS.1-30.0.001.bcf - matched-calls/varlociraptor-lancet/simulated-bwa.INS.1-30.0.01.bcf - matched-calls/varlociraptor-lancet/simulated-bwa.INS.1-30.0.05.bcf - matched-calls/varlociraptor-lancet/simulated-bwa.INS.1-30.0.1.bcf - matched-calls/varlociraptor-lancet/simulated-bwa.INS.1-30.0.2.bcf - matched-calls/varlociraptor-lancet/simulated-bwa.INS.1-30.0.3.bcf - matched-calls/varlociraptor-lancet/simulated-bwa.INS.1-30.0.4.bcf - matched-calls/varlociraptor-lancet/simulated-bwa.INS.1-30.0.5.bcf - matched-calls/varlociraptor-lancet/simulated-bwa.INS.1-30.0.6.bcf - matched-calls/varlociraptor-lancet/simulated-bwa.INS.1-30.0.7.bcf - matched-calls/varlociraptor-lancet/simulated-bwa.INS.1-30.0.8.bcf - matched-calls/varlociraptor-lancet/simulated-bwa.INS.1-30.0.9.bcf - matched-calls/varlociraptor-lancet/simulated-bwa.INS.30-100.1e-05.bcf - matched-calls/varlociraptor-lancet/simulated-bwa.INS.30-100.0.0001.bcf - matched-calls/varlociraptor-lancet/simulated-bwa.INS.30-100.0.001.bcf - matched-calls/varlociraptor-lancet/simulated-bwa.INS.30-100.0.01.bcf - matched-calls/varlociraptor-lancet/simulated-bwa.INS.30-100.0.05.bcf - matched-calls/varlociraptor-lancet/simulated-bwa.INS.30-100.0.1.bcf - matched-calls/varlociraptor-lancet/simulated-bwa.INS.30-100.0.2.bcf - matched-calls/varlociraptor-lancet/simulated-bwa.INS.30-100.0.3.bcf - matched-calls/varlociraptor-lancet/simulated-bwa.INS.30-100.0.4.bcf - matched-calls/varlociraptor-lancet/simulated-bwa.INS.30-100.0.5.bcf - matched-calls/varlociraptor-lancet/simulated-bwa.INS.30-100.0.6.bcf - matched-calls/varlociraptor-lancet/simulated-bwa.INS.30-100.0.7.bcf - matched-calls/varlociraptor-lancet/simulated-bwa.INS.30-100.0.8.bcf - matched-calls/varlociraptor-lancet/simulated-bwa.INS.30-100.0.9.bcf - matched-calls/varlociraptor-manta/simulated-bwa.INS.1-30.1e-05.bcf - matched-calls/varlociraptor-manta/simulated-bwa.INS.1-30.0.0001.bcf - matched-calls/varlociraptor-manta/simulated-bwa.INS.1-30.0.001.bcf - matched-calls/varlociraptor-manta/simulated-bwa.INS.1-30.0.01.bcf - matched-calls/varlociraptor-manta/simulated-bwa.INS.1-30.0.05.bcf - matched-calls/varlociraptor-manta/simulated-bwa.INS.1-30.0.1.bcf - matched-calls/varlociraptor-manta/simulated-bwa.INS.1-30.0.2.bcf - matched-calls/varlociraptor-manta/simulated-bwa.INS.1-30.0.3.bcf - matched-calls/varlociraptor-manta/simulated-bwa.INS.1-30.0.4.bcf - matched-calls/varlociraptor-manta/simulated-bwa.INS.1-30.0.5.bcf - matched-calls/varlociraptor-manta/simulated-bwa.INS.1-30.0.6.bcf - matched-calls/varlociraptor-manta/simulated-bwa.INS.1-30.0.7.bcf - matched-calls/varlociraptor-manta/simulated-bwa.INS.1-30.0.8.bcf - matched-calls/varlociraptor-manta/simulated-bwa.INS.1-30.0.9.bcf - matched-calls/varlociraptor-manta/simulated-bwa.INS.30-100.1e-05.bcf - matched-calls/varlociraptor-manta/simulated-bwa.INS.30-100.0.0001.bcf - matched-calls/varlociraptor-manta/simulated-bwa.INS.30-100.0.001.bcf - matched-calls/varlociraptor-manta/simulated-bwa.INS.30-100.0.01.bcf - matched-calls/varlociraptor-manta/simulated-bwa.INS.30-100.0.05.bcf - matched-calls/varlociraptor-manta/simulated-bwa.INS.30-100.0.1.bcf - matched-calls/varlociraptor-manta/simulated-bwa.INS.30-100.0.2.bcf - matched-calls/varlociraptor-manta/simulated-bwa.INS.30-100.0.3.bcf - matched-calls/varlociraptor-manta/simulated-bwa.INS.30-100.0.4.bcf - matched-calls/varlociraptor-manta/simulated-bwa.INS.30-100.0.5.bcf - matched-calls/varlociraptor-manta/simulated-bwa.INS.30-100.0.6.bcf - matched-calls/varlociraptor-manta/simulated-bwa.INS.30-100.0.7.bcf - matched-calls/varlociraptor-manta/simulated-bwa.INS.30-100.0.8.bcf - matched-calls/varlociraptor-manta/simulated-bwa.INS.30-100.0.9.bcf - matched-calls/varlociraptor-strelka/simulated-bwa.INS.1-30.1e-05.bcf - matched-calls/varlociraptor-strelka/simulated-bwa.INS.1-30.0.0001.bcf - matched-calls/varlociraptor-strelka/simulated-bwa.INS.1-30.0.001.bcf - matched-calls/varlociraptor-strelka/simulated-bwa.INS.1-30.0.01.bcf - matched-calls/varlociraptor-strelka/simulated-bwa.INS.1-30.0.05.bcf - matched-calls/varlociraptor-strelka/simulated-bwa.INS.1-30.0.1.bcf - matched-calls/varlociraptor-strelka/simulated-bwa.INS.1-30.0.2.bcf - matched-calls/varlociraptor-strelka/simulated-bwa.INS.1-30.0.3.bcf - matched-calls/varlociraptor-strelka/simulated-bwa.INS.1-30.0.4.bcf - matched-calls/varlociraptor-strelka/simulated-bwa.INS.1-30.0.5.bcf - matched-calls/varlociraptor-strelka/simulated-bwa.INS.1-30.0.6.bcf - matched-calls/varlociraptor-strelka/simulated-bwa.INS.1-30.0.7.bcf - matched-calls/varlociraptor-strelka/simulated-bwa.INS.1-30.0.8.bcf - matched-calls/varlociraptor-strelka/simulated-bwa.INS.1-30.0.9.bcf - matched-calls/varlociraptor-strelka/simulated-bwa.INS.30-100.1e-05.bcf - matched-calls/varlociraptor-strelka/simulated-bwa.INS.30-100.0.0001.bcf - matched-calls/varlociraptor-strelka/simulated-bwa.INS.30-100.0.001.bcf - matched-calls/varlociraptor-strelka/simulated-bwa.INS.30-100.0.01.bcf - matched-calls/varlociraptor-strelka/simulated-bwa.INS.30-100.0.05.bcf - matched-calls/varlociraptor-strelka/simulated-bwa.INS.30-100.0.1.bcf - matched-calls/varlociraptor-strelka/simulated-bwa.INS.30-100.0.2.bcf - matched-calls/varlociraptor-strelka/simulated-bwa.INS.30-100.0.3.bcf - matched-calls/varlociraptor-strelka/simulated-bwa.INS.30-100.0.4.bcf - matched-calls/varlociraptor-strelka/simulated-bwa.INS.30-100.0.5.bcf - matched-calls/varlociraptor-strelka/simulated-bwa.INS.30-100.0.6.bcf - matched-calls/varlociraptor-strelka/simulated-bwa.INS.30-100.0.7.bcf - matched-calls/varlociraptor-strelka/simulated-bwa.INS.30-100.0.8.bcf - matched-calls/varlociraptor-strelka/simulated-bwa.INS.30-100.0.9.bcf - matched-calls/varlociraptor-delly/simulated-bwa.DEL.1-30.1e-05.bcf - matched-calls/varlociraptor-delly/simulated-bwa.DEL.1-30.0.0001.bcf - matched-calls/varlociraptor-delly/simulated-bwa.DEL.1-30.0.001.bcf - matched-calls/varlociraptor-delly/simulated-bwa.DEL.1-30.0.01.bcf - matched-calls/varlociraptor-delly/simulated-bwa.DEL.1-30.0.05.bcf - matched-calls/varlociraptor-delly/simulated-bwa.DEL.1-30.0.1.bcf - matched-calls/varlociraptor-delly/simulated-bwa.DEL.1-30.0.2.bcf - matched-calls/varlociraptor-delly/simulated-bwa.DEL.1-30.0.3.bcf - matched-calls/varlociraptor-delly/simulated-bwa.DEL.1-30.0.4.bcf - matched-calls/varlociraptor-delly/simulated-bwa.DEL.1-30.0.5.bcf - matched-calls/varlociraptor-delly/simulated-bwa.DEL.1-30.0.6.bcf - matched-calls/varlociraptor-delly/simulated-bwa.DEL.1-30.0.7.bcf - matched-calls/varlociraptor-delly/simulated-bwa.DEL.1-30.0.8.bcf - matched-calls/varlociraptor-delly/simulated-bwa.DEL.1-30.0.9.bcf - matched-calls/varlociraptor-delly/simulated-bwa.DEL.30-50.1e-05.bcf - matched-calls/varlociraptor-delly/simulated-bwa.DEL.30-50.0.0001.bcf - matched-calls/varlociraptor-delly/simulated-bwa.DEL.30-50.0.001.bcf - matched-calls/varlociraptor-delly/simulated-bwa.DEL.30-50.0.01.bcf - matched-calls/varlociraptor-delly/simulated-bwa.DEL.30-50.0.05.bcf - matched-calls/varlociraptor-delly/simulated-bwa.DEL.30-50.0.1.bcf - matched-calls/varlociraptor-delly/simulated-bwa.DEL.30-50.0.2.bcf - matched-calls/varlociraptor-delly/simulated-bwa.DEL.30-50.0.3.bcf - matched-calls/varlociraptor-delly/simulated-bwa.DEL.30-50.0.4.bcf - matched-calls/varlociraptor-delly/simulated-bwa.DEL.30-50.0.5.bcf - matched-calls/varlociraptor-delly/simulated-bwa.DEL.30-50.0.6.bcf - matched-calls/varlociraptor-delly/simulated-bwa.DEL.30-50.0.7.bcf - matched-calls/varlociraptor-delly/simulated-bwa.DEL.30-50.0.8.bcf - matched-calls/varlociraptor-delly/simulated-bwa.DEL.30-50.0.9.bcf - matched-calls/varlociraptor-delly/simulated-bwa.DEL.50-100.1e-05.bcf - matched-calls/varlociraptor-delly/simulated-bwa.DEL.50-100.0.0001.bcf - matched-calls/varlociraptor-delly/simulated-bwa.DEL.50-100.0.001.bcf - matched-calls/varlociraptor-delly/simulated-bwa.DEL.50-100.0.01.bcf - matched-calls/varlociraptor-delly/simulated-bwa.DEL.50-100.0.05.bcf - matched-calls/varlociraptor-delly/simulated-bwa.DEL.50-100.0.1.bcf - matched-calls/varlociraptor-delly/simulated-bwa.DEL.50-100.0.2.bcf - matched-calls/varlociraptor-delly/simulated-bwa.DEL.50-100.0.3.bcf - matched-calls/varlociraptor-delly/simulated-bwa.DEL.50-100.0.4.bcf - matched-calls/varlociraptor-delly/simulated-bwa.DEL.50-100.0.5.bcf - matched-calls/varlociraptor-delly/simulated-bwa.DEL.50-100.0.6.bcf - matched-calls/varlociraptor-delly/simulated-bwa.DEL.50-100.0.7.bcf - matched-calls/varlociraptor-delly/simulated-bwa.DEL.50-100.0.8.bcf - matched-calls/varlociraptor-delly/simulated-bwa.DEL.50-100.0.9.bcf - matched-calls/varlociraptor-delly/simulated-bwa.DEL.100-250.1e-05.bcf - matched-calls/varlociraptor-delly/simulated-bwa.DEL.100-250.0.0001.bcf - matched-calls/varlociraptor-delly/simulated-bwa.DEL.100-250.0.001.bcf - matched-calls/varlociraptor-delly/simulated-bwa.DEL.100-250.0.01.bcf - matched-calls/varlociraptor-delly/simulated-bwa.DEL.100-250.0.05.bcf - matched-calls/varlociraptor-delly/simulated-bwa.DEL.100-250.0.1.bcf - matched-calls/varlociraptor-delly/simulated-bwa.DEL.100-250.0.2.bcf - matched-calls/varlociraptor-delly/simulated-bwa.DEL.100-250.0.3.bcf - matched-calls/varlociraptor-delly/simulated-bwa.DEL.100-250.0.4.bcf - matched-calls/varlociraptor-delly/simulated-bwa.DEL.100-250.0.5.bcf - matched-calls/varlociraptor-delly/simulated-bwa.DEL.100-250.0.6.bcf - matched-calls/varlociraptor-delly/simulated-bwa.DEL.100-250.0.7.bcf - matched-calls/varlociraptor-delly/simulated-bwa.DEL.100-250.0.8.bcf - matched-calls/varlociraptor-delly/simulated-bwa.DEL.100-250.0.9.bcf - matched-calls/varlociraptor-lancet/simulated-bwa.DEL.1-30.1e-05.bcf - matched-calls/varlociraptor-lancet/simulated-bwa.DEL.1-30.0.0001.bcf - matched-calls/varlociraptor-lancet/simulated-bwa.DEL.1-30.0.001.bcf - matched-calls/varlociraptor-lancet/simulated-bwa.DEL.1-30.0.01.bcf - matched-calls/varlociraptor-lancet/simulated-bwa.DEL.1-30.0.05.bcf - matched-calls/varlociraptor-lancet/simulated-bwa.DEL.1-30.0.1.bcf - matched-calls/varlociraptor-lancet/simulated-bwa.DEL.1-30.0.2.bcf - matched-calls/varlociraptor-lancet/simulated-bwa.DEL.1-30.0.3.bcf - matched-calls/varlociraptor-lancet/simulated-bwa.DEL.1-30.0.4.bcf - matched-calls/varlociraptor-lancet/simulated-bwa.DEL.1-30.0.5.bcf - matched-calls/varlociraptor-lancet/simulated-bwa.DEL.1-30.0.6.bcf - matched-calls/varlociraptor-lancet/simulated-bwa.DEL.1-30.0.7.bcf - matched-calls/varlociraptor-lancet/simulated-bwa.DEL.1-30.0.8.bcf - matched-calls/varlociraptor-lancet/simulated-bwa.DEL.1-30.0.9.bcf - matched-calls/varlociraptor-lancet/simulated-bwa.DEL.30-50.1e-05.bcf - matched-calls/varlociraptor-lancet/simulated-bwa.DEL.30-50.0.0001.bcf - matched-calls/varlociraptor-lancet/simulated-bwa.DEL.30-50.0.001.bcf - matched-calls/varlociraptor-lancet/simulated-bwa.DEL.30-50.0.01.bcf - matched-calls/varlociraptor-lancet/simulated-bwa.DEL.30-50.0.05.bcf - matched-calls/varlociraptor-lancet/simulated-bwa.DEL.30-50.0.1.bcf - matched-calls/varlociraptor-lancet/simulated-bwa.DEL.30-50.0.2.bcf - matched-calls/varlociraptor-lancet/simulated-bwa.DEL.30-50.0.3.bcf - matched-calls/varlociraptor-lancet/simulated-bwa.DEL.30-50.0.4.bcf - matched-calls/varlociraptor-lancet/simulated-bwa.DEL.30-50.0.5.bcf - matched-calls/varlociraptor-lancet/simulated-bwa.DEL.30-50.0.6.bcf - matched-calls/varlociraptor-lancet/simulated-bwa.DEL.30-50.0.7.bcf - matched-calls/varlociraptor-lancet/simulated-bwa.DEL.30-50.0.8.bcf - matched-calls/varlociraptor-lancet/simulated-bwa.DEL.30-50.0.9.bcf - matched-calls/varlociraptor-lancet/simulated-bwa.DEL.50-100.1e-05.bcf - matched-calls/varlociraptor-lancet/simulated-bwa.DEL.50-100.0.0001.bcf - matched-calls/varlociraptor-lancet/simulated-bwa.DEL.50-100.0.001.bcf - matched-calls/varlociraptor-lancet/simulated-bwa.DEL.50-100.0.01.bcf - matched-calls/varlociraptor-lancet/simulated-bwa.DEL.50-100.0.05.bcf - matched-calls/varlociraptor-lancet/simulated-bwa.DEL.50-100.0.1.bcf - matched-calls/varlociraptor-lancet/simulated-bwa.DEL.50-100.0.2.bcf - matched-calls/varlociraptor-lancet/simulated-bwa.DEL.50-100.0.3.bcf - matched-calls/varlociraptor-lancet/simulated-bwa.DEL.50-100.0.4.bcf - matched-calls/varlociraptor-lancet/simulated-bwa.DEL.50-100.0.5.bcf - matched-calls/varlociraptor-lancet/simulated-bwa.DEL.50-100.0.6.bcf - matched-calls/varlociraptor-lancet/simulated-bwa.DEL.50-100.0.7.bcf - matched-calls/varlociraptor-lancet/simulated-bwa.DEL.50-100.0.8.bcf - matched-calls/varlociraptor-lancet/simulated-bwa.DEL.50-100.0.9.bcf - matched-calls/varlociraptor-lancet/simulated-bwa.DEL.100-250.1e-05.bcf - matched-calls/varlociraptor-lancet/simulated-bwa.DEL.100-250.0.0001.bcf - matched-calls/varlociraptor-lancet/simulated-bwa.DEL.100-250.0.001.bcf - matched-calls/varlociraptor-lancet/simulated-bwa.DEL.100-250.0.01.bcf - matched-calls/varlociraptor-lancet/simulated-bwa.DEL.100-250.0.05.bcf - matched-calls/varlociraptor-lancet/simulated-bwa.DEL.100-250.0.1.bcf - matched-calls/varlociraptor-lancet/simulated-bwa.DEL.100-250.0.2.bcf - matched-calls/varlociraptor-lancet/simulated-bwa.DEL.100-250.0.3.bcf - matched-calls/varlociraptor-lancet/simulated-bwa.DEL.100-250.0.4.bcf - matched-calls/varlociraptor-lancet/simulated-bwa.DEL.100-250.0.5.bcf - matched-calls/varlociraptor-lancet/simulated-bwa.DEL.100-250.0.6.bcf - matched-calls/varlociraptor-lancet/simulated-bwa.DEL.100-250.0.7.bcf - matched-calls/varlociraptor-lancet/simulated-bwa.DEL.100-250.0.8.bcf - matched-calls/varlociraptor-lancet/simulated-bwa.DEL.100-250.0.9.bcf - matched-calls/varlociraptor-manta/simulated-bwa.DEL.1-30.1e-05.bcf - matched-calls/varlociraptor-manta/simulated-bwa.DEL.1-30.0.0001.bcf - matched-calls/varlociraptor-manta/simulated-bwa.DEL.1-30.0.001.bcf - matched-calls/varlociraptor-manta/simulated-bwa.DEL.1-30.0.01.bcf - matched-calls/varlociraptor-manta/simulated-bwa.DEL.1-30.0.05.bcf - matched-calls/varlociraptor-manta/simulated-bwa.DEL.1-30.0.1.bcf - matched-calls/varlociraptor-manta/simulated-bwa.DEL.1-30.0.2.bcf - matched-calls/varlociraptor-manta/simulated-bwa.DEL.1-30.0.3.bcf - matched-calls/varlociraptor-manta/simulated-bwa.DEL.1-30.0.4.bcf - matched-calls/varlociraptor-manta/simulated-bwa.DEL.1-30.0.5.bcf - matched-calls/varlociraptor-manta/simulated-bwa.DEL.1-30.0.6.bcf - matched-calls/varlociraptor-manta/simulated-bwa.DEL.1-30.0.7.bcf - matched-calls/varlociraptor-manta/simulated-bwa.DEL.1-30.0.8.bcf - matched-calls/varlociraptor-manta/simulated-bwa.DEL.1-30.0.9.bcf - matched-calls/varlociraptor-manta/simulated-bwa.DEL.30-50.1e-05.bcf - matched-calls/varlociraptor-manta/simulated-bwa.DEL.30-50.0.0001.bcf - matched-calls/varlociraptor-manta/simulated-bwa.DEL.30-50.0.001.bcf - matched-calls/varlociraptor-manta/simulated-bwa.DEL.30-50.0.01.bcf - matched-calls/varlociraptor-manta/simulated-bwa.DEL.30-50.0.05.bcf - matched-calls/varlociraptor-manta/simulated-bwa.DEL.30-50.0.1.bcf - matched-calls/varlociraptor-manta/simulated-bwa.DEL.30-50.0.2.bcf - matched-calls/varlociraptor-manta/simulated-bwa.DEL.30-50.0.3.bcf - matched-calls/varlociraptor-manta/simulated-bwa.DEL.30-50.0.4.bcf - matched-calls/varlociraptor-manta/simulated-bwa.DEL.30-50.0.5.bcf - matched-calls/varlociraptor-manta/simulated-bwa.DEL.30-50.0.6.bcf - matched-calls/varlociraptor-manta/simulated-bwa.DEL.30-50.0.7.bcf - matched-calls/varlociraptor-manta/simulated-bwa.DEL.30-50.0.8.bcf - matched-calls/varlociraptor-manta/simulated-bwa.DEL.30-50.0.9.bcf - matched-calls/varlociraptor-manta/simulated-bwa.DEL.50-100.1e-05.bcf - matched-calls/varlociraptor-manta/simulated-bwa.DEL.50-100.0.0001.bcf - matched-calls/varlociraptor-manta/simulated-bwa.DEL.50-100.0.001.bcf - matched-calls/varlociraptor-manta/simulated-bwa.DEL.50-100.0.01.bcf - matched-calls/varlociraptor-manta/simulated-bwa.DEL.50-100.0.05.bcf - matched-calls/varlociraptor-manta/simulated-bwa.DEL.50-100.0.1.bcf - matched-calls/varlociraptor-manta/simulated-bwa.DEL.50-100.0.2.bcf - matched-calls/varlociraptor-manta/simulated-bwa.DEL.50-100.0.3.bcf - matched-calls/varlociraptor-manta/simulated-bwa.DEL.50-100.0.4.bcf - matched-calls/varlociraptor-manta/simulated-bwa.DEL.50-100.0.5.bcf - matched-calls/varlociraptor-manta/simulated-bwa.DEL.50-100.0.6.bcf - matched-calls/varlociraptor-manta/simulated-bwa.DEL.50-100.0.7.bcf - matched-calls/varlociraptor-manta/simulated-bwa.DEL.50-100.0.8.bcf - matched-calls/varlociraptor-manta/simulated-bwa.DEL.50-100.0.9.bcf - matched-calls/varlociraptor-manta/simulated-bwa.DEL.100-250.1e-05.bcf - matched-calls/varlociraptor-manta/simulated-bwa.DEL.100-250.0.0001.bcf - matched-calls/varlociraptor-manta/simulated-bwa.DEL.100-250.0.001.bcf - matched-calls/varlociraptor-manta/simulated-bwa.DEL.100-250.0.01.bcf - matched-calls/varlociraptor-manta/simulated-bwa.DEL.100-250.0.05.bcf - matched-calls/varlociraptor-manta/simulated-bwa.DEL.100-250.0.1.bcf - matched-calls/varlociraptor-manta/simulated-bwa.DEL.100-250.0.2.bcf - matched-calls/varlociraptor-manta/simulated-bwa.DEL.100-250.0.3.bcf - matched-calls/varlociraptor-manta/simulated-bwa.DEL.100-250.0.4.bcf - matched-calls/varlociraptor-manta/simulated-bwa.DEL.100-250.0.5.bcf - matched-calls/varlociraptor-manta/simulated-bwa.DEL.100-250.0.6.bcf - matched-calls/varlociraptor-manta/simulated-bwa.DEL.100-250.0.7.bcf - matched-calls/varlociraptor-manta/simulated-bwa.DEL.100-250.0.8.bcf - matched-calls/varlociraptor-manta/simulated-bwa.DEL.100-250.0.9.bcf - matched-calls/varlociraptor-strelka/simulated-bwa.DEL.1-30.1e-05.bcf - matched-calls/varlociraptor-strelka/simulated-bwa.DEL.1-30.0.0001.bcf - matched-calls/varlociraptor-strelka/simulated-bwa.DEL.1-30.0.001.bcf - matched-calls/varlociraptor-strelka/simulated-bwa.DEL.1-30.0.01.bcf - matched-calls/varlociraptor-strelka/simulated-bwa.DEL.1-30.0.05.bcf - matched-calls/varlociraptor-strelka/simulated-bwa.DEL.1-30.0.1.bcf - matched-calls/varlociraptor-strelka/simulated-bwa.DEL.1-30.0.2.bcf - matched-calls/varlociraptor-strelka/simulated-bwa.DEL.1-30.0.3.bcf - matched-calls/varlociraptor-strelka/simulated-bwa.DEL.1-30.0.4.bcf - matched-calls/varlociraptor-strelka/simulated-bwa.DEL.1-30.0.5.bcf - matched-calls/varlociraptor-strelka/simulated-bwa.DEL.1-30.0.6.bcf - matched-calls/varlociraptor-strelka/simulated-bwa.DEL.1-30.0.7.bcf - matched-calls/varlociraptor-strelka/simulated-bwa.DEL.1-30.0.8.bcf - matched-calls/varlociraptor-strelka/simulated-bwa.DEL.1-30.0.9.bcf - matched-calls/varlociraptor-strelka/simulated-bwa.DEL.30-50.1e-05.bcf - matched-calls/varlociraptor-strelka/simulated-bwa.DEL.30-50.0.0001.bcf - matched-calls/varlociraptor-strelka/simulated-bwa.DEL.30-50.0.001.bcf - matched-calls/varlociraptor-strelka/simulated-bwa.DEL.30-50.0.01.bcf - matched-calls/varlociraptor-strelka/simulated-bwa.DEL.30-50.0.05.bcf - matched-calls/varlociraptor-strelka/simulated-bwa.DEL.30-50.0.1.bcf - matched-calls/varlociraptor-strelka/simulated-bwa.DEL.30-50.0.2.bcf - matched-calls/varlociraptor-strelka/simulated-bwa.DEL.30-50.0.3.bcf - matched-calls/varlociraptor-strelka/simulated-bwa.DEL.30-50.0.4.bcf - matched-calls/varlociraptor-strelka/simulated-bwa.DEL.30-50.0.5.bcf - matched-calls/varlociraptor-strelka/simulated-bwa.DEL.30-50.0.6.bcf - matched-calls/varlociraptor-strelka/simulated-bwa.DEL.30-50.0.7.bcf - matched-calls/varlociraptor-strelka/simulated-bwa.DEL.30-50.0.8.bcf - matched-calls/varlociraptor-strelka/simulated-bwa.DEL.30-50.0.9.bcf - matched-calls/varlociraptor-strelka/simulated-bwa.DEL.50-100.1e-05.bcf - matched-calls/varlociraptor-strelka/simulated-bwa.DEL.50-100.0.0001.bcf - matched-calls/varlociraptor-strelka/simulated-bwa.DEL.50-100.0.001.bcf - matched-calls/varlociraptor-strelka/simulated-bwa.DEL.50-100.0.01.bcf - matched-calls/varlociraptor-strelka/simulated-bwa.DEL.50-100.0.05.bcf - matched-calls/varlociraptor-strelka/simulated-bwa.DEL.50-100.0.1.bcf - matched-calls/varlociraptor-strelka/simulated-bwa.DEL.50-100.0.2.bcf - matched-calls/varlociraptor-strelka/simulated-bwa.DEL.50-100.0.3.bcf - matched-calls/varlociraptor-strelka/simulated-bwa.DEL.50-100.0.4.bcf - matched-calls/varlociraptor-strelka/simulated-bwa.DEL.50-100.0.5.bcf - matched-calls/varlociraptor-strelka/simulated-bwa.DEL.50-100.0.6.bcf - matched-calls/varlociraptor-strelka/simulated-bwa.DEL.50-100.0.7.bcf - matched-calls/varlociraptor-strelka/simulated-bwa.DEL.50-100.0.8.bcf - matched-calls/varlociraptor-strelka/simulated-bwa.DEL.50-100.0.9.bcf - matched-calls/varlociraptor-strelka/simulated-bwa.DEL.100-250.1e-05.bcf - matched-calls/varlociraptor-strelka/simulated-bwa.DEL.100-250.0.0001.bcf - matched-calls/varlociraptor-strelka/simulated-bwa.DEL.100-250.0.001.bcf - matched-calls/varlociraptor-strelka/simulated-bwa.DEL.100-250.0.01.bcf - matched-calls/varlociraptor-strelka/simulated-bwa.DEL.100-250.0.05.bcf - matched-calls/varlociraptor-strelka/simulated-bwa.DEL.100-250.0.1.bcf - matched-calls/varlociraptor-strelka/simulated-bwa.DEL.100-250.0.2.bcf - matched-calls/varlociraptor-strelka/simulated-bwa.DEL.100-250.0.3.bcf - matched-calls/varlociraptor-strelka/simulated-bwa.DEL.100-250.0.4.bcf - matched-calls/varlociraptor-strelka/simulated-bwa.DEL.100-250.0.5.bcf - matched-calls/varlociraptor-strelka/simulated-bwa.DEL.100-250.0.6.bcf - matched-calls/varlociraptor-strelka/simulated-bwa.DEL.100-250.0.7.bcf - matched-calls/varlociraptor-strelka/simulated-bwa.DEL.100-250.0.8.bcf - matched-calls/varlociraptor-strelka/simulated-bwa.DEL.100-250.0.9.bcf - matched-calls/varlociraptor-delly/simulated-bwa.INS.1-250.1.0.bcf - matched-calls/varlociraptor-lancet/simulated-bwa.INS.1-250.1.0.bcf - matched-calls/varlociraptor-manta/simulated-bwa.INS.1-250.1.0.bcf - matched-calls/varlociraptor-strelka/simulated-bwa.INS.1-250.1.0.bcf - matched-calls/varlociraptor-delly/simulated-bwa.DEL.1-250.1.0.bcf - matched-calls/varlociraptor-lancet/simulated-bwa.DEL.1-250.1.0.bcf - matched-calls/varlociraptor-manta/simulated-bwa.DEL.1-250.1.0.bcf - matched-calls/varlociraptor-strelka/simulated-bwa.DEL.1-250.1.0.bcf |  | - rust-bio-tools =0.5.0 - bedtools =2.27.1 - bcftools =1.8 | ``` 1 ``` \| ``` rbt vcf-match {params} {input.truth} < {input.calls} > {output} ``` |
| match\_other\_calls | 8 | - matched-calls/default-lancet/simulated-bwa.all.bcf - matched-calls/default-manta/simulated-bwa.all.bcf - matched-calls/default-strelka/simulated-bwa.all.bcf - matched-calls/adhoc-delly/simulated-bwa.all.bcf - matched-calls/adhoc-lancet/simulated-bwa.all.bcf - matched-calls/adhoc-manta/simulated-bwa.all.bcf - matched-calls/adhoc-strelka/simulated-bwa.all.bcf - matched-calls/adhoc-bpi/simulated-bwa.all.bcf |  |  | ``` 1 ``` \| ``` rbt vcf-match {params} {input.truth} < {input.calls} > {output} ``` |
| concordance\_match | 60 | - concordance/varlociraptor-delly-0.9/colo1.0-vs-1.bcf - concordance/varlociraptor-delly-0.9/colo1.1-vs-2.bcf - concordance/varlociraptor-delly-0.9/colo1.2-vs-0.bcf - concordance/varlociraptor-delly-0.9/colo1.2-vs-3.bcf - concordance/varlociraptor-delly-0.9/colo1.3-vs-0.bcf - concordance/varlociraptor-delly-0.9/colo1.3-vs-1.bcf - concordance/varlociraptor-lancet-0.9/colo1.0-vs-1.bcf - concordance/varlociraptor-lancet-0.9/colo1.1-vs-2.bcf - concordance/varlociraptor-lancet-0.9/colo1.2-vs-0.bcf - concordance/varlociraptor-lancet-0.9/colo1.2-vs-3.bcf - concordance/varlociraptor-lancet-0.9/colo1.3-vs-0.bcf - concordance/varlociraptor-lancet-0.9/colo1.3-vs-1.bcf - concordance/varlociraptor-manta-0.9/colo1.0-vs-1.bcf - concordance/varlociraptor-manta-0.9/colo1.1-vs-2.bcf - concordance/varlociraptor-manta-0.9/colo1.2-vs-0.bcf - concordance/varlociraptor-manta-0.9/colo1.2-vs-3.bcf - concordance/varlociraptor-manta-0.9/colo1.3-vs-0.bcf - concordance/varlociraptor-manta-0.9/colo1.3-vs-1.bcf - concordance/varlociraptor-strelka-0.9/colo1.0-vs-1.bcf - concordance/varlociraptor-strelka-0.9/colo1.1-vs-2.bcf - concordance/varlociraptor-strelka-0.9/colo1.2-vs-0.bcf - concordance/varlociraptor-strelka-0.9/colo1.2-vs-3.bcf - concordance/varlociraptor-strelka-0.9/colo1.3-vs-0.bcf - concordance/varlociraptor-strelka-0.9/colo1.3-vs-1.bcf - concordance/varlociraptor-bpi-0.9/colo1.0-vs-1.bcf - concordance/varlociraptor-bpi-0.9/colo1.1-vs-2.bcf - concordance/varlociraptor-bpi-0.9/colo1.2-vs-0.bcf - concordance/varlociraptor-bpi-0.9/colo1.2-vs-3.bcf - concordance/varlociraptor-bpi-0.9/colo1.3-vs-0.bcf - concordance/varlociraptor-bpi-0.9/colo1.3-vs-1.bcf - concordance/varlociraptor-delly-0.98/colo1.0-vs-1.bcf - concordance/varlociraptor-delly-0.98/colo1.1-vs-2.bcf - concordance/varlociraptor-delly-0.98/colo1.2-vs-0.bcf - concordance/varlociraptor-delly-0.98/colo1.2-vs-3.bcf - concordance/varlociraptor-delly-0.98/colo1.3-vs-0.bcf - concordance/varlociraptor-delly-0.98/colo1.3-vs-1.bcf - concordance/varlociraptor-lancet-0.98/colo1.0-vs-1.bcf - concordance/varlociraptor-lancet-0.98/colo1.1-vs-2.bcf - concordance/varlociraptor-lancet-0.98/colo1.2-vs-0.bcf - concordance/varlociraptor-lancet-0.98/colo1.2-vs-3.bcf - concordance/varlociraptor-lancet-0.98/colo1.3-vs-0.bcf - concordance/varlociraptor-lancet-0.98/colo1.3-vs-1.bcf - concordance/varlociraptor-manta-0.98/colo1.0-vs-1.bcf - concordance/varlociraptor-manta-0.98/colo1.1-vs-2.bcf - concordance/varlociraptor-manta-0.98/colo1.2-vs-0.bcf - concordance/varlociraptor-manta-0.98/colo1.2-vs-3.bcf - concordance/varlociraptor-manta-0.98/colo1.3-vs-0.bcf - concordance/varlociraptor-manta-0.98/colo1.3-vs-1.bcf - concordance/varlociraptor-strelka-0.98/colo1.0-vs-1.bcf - concordance/varlociraptor-strelka-0.98/colo1.1-vs-2.bcf - concordance/varlociraptor-strelka-0.98/colo1.2-vs-0.bcf - concordance/varlociraptor-strelka-0.98/colo1.2-vs-3.bcf - concordance/varlociraptor-strelka-0.98/colo1.3-vs-0.bcf - concordance/varlociraptor-strelka-0.98/colo1.3-vs-1.bcf - concordance/varlociraptor-bpi-0.98/colo1.0-vs-1.bcf - concordance/varlociraptor-bpi-0.98/colo1.1-vs-2.bcf - concordance/varlociraptor-bpi-0.98/colo1.2-vs-0.bcf - concordance/varlociraptor-bpi-0.98/colo1.2-vs-3.bcf - concordance/varlociraptor-bpi-0.98/colo1.3-vs-0.bcf - concordance/varlociraptor-bpi-0.98/colo1.3-vs-1.bcf |  | - rust-bio-tools =0.5.0 - bedtools =2.27.1 - bcftools =1.8 | ``` 1 ``` \| ``` rbt vcf-match {params.match} {params.bcfs[1]} < {params.bcfs[0]} > {output} ``` |
| concordance\_match | 30 | - concordance/adhoc-delly-default/colo1.0-vs-1.bcf - concordance/adhoc-delly-default/colo1.1-vs-2.bcf - concordance/adhoc-delly-default/colo1.2-vs-0.bcf - concordance/adhoc-delly-default/colo1.2-vs-3.bcf - concordance/adhoc-delly-default/colo1.3-vs-0.bcf - concordance/adhoc-delly-default/colo1.3-vs-1.bcf - concordance/adhoc-lancet-default/colo1.0-vs-1.bcf - concordance/adhoc-lancet-default/colo1.1-vs-2.bcf - concordance/adhoc-lancet-default/colo1.2-vs-0.bcf - concordance/adhoc-lancet-default/colo1.2-vs-3.bcf - concordance/adhoc-lancet-default/colo1.3-vs-0.bcf - concordance/adhoc-lancet-default/colo1.3-vs-1.bcf - concordance/adhoc-manta-default/colo1.0-vs-1.bcf - concordance/adhoc-manta-default/colo1.1-vs-2.bcf - concordance/adhoc-manta-default/colo1.2-vs-0.bcf - concordance/adhoc-manta-default/colo1.2-vs-3.bcf - concordance/adhoc-manta-default/colo1.3-vs-0.bcf - concordance/adhoc-manta-default/colo1.3-vs-1.bcf - concordance/adhoc-strelka-default/colo1.0-vs-1.bcf - concordance/adhoc-strelka-default/colo1.1-vs-2.bcf - concordance/adhoc-strelka-default/colo1.2-vs-0.bcf - concordance/adhoc-strelka-default/colo1.2-vs-3.bcf - concordance/adhoc-strelka-default/colo1.3-vs-0.bcf - concordance/adhoc-strelka-default/colo1.3-vs-1.bcf - concordance/adhoc-bpi-default/colo1.0-vs-1.bcf - concordance/adhoc-bpi-default/colo1.1-vs-2.bcf - concordance/adhoc-bpi-default/colo1.2-vs-0.bcf - concordance/adhoc-bpi-default/colo1.2-vs-3.bcf - concordance/adhoc-bpi-default/colo1.3-vs-0.bcf - concordance/adhoc-bpi-default/colo1.3-vs-1.bcf |  | - rust-bio-tools =0.2.5 - bedtools =2.27.1 - bcftools =1.8 | ``` 1 ``` \| ``` rbt vcf-match {params.match} {params.bcfs[1]} < {params.bcfs[0]} > {output} ``` |
| varlociraptor\_merge | 24 | - varlociraptor-delly/COLO\_829-GSC.all.bcf - varlociraptor-delly/COLO\_829-Ill.all.bcf - varlociraptor-delly/COLO\_829-TGen.all.bcf - varlociraptor-delly/COLO\_829-EBI.all.bcf - varlociraptor-lancet/COLO\_829-GSC.all.bcf - varlociraptor-lancet/COLO\_829-Ill.all.bcf - varlociraptor-lancet/COLO\_829-TGen.all.bcf - varlociraptor-lancet/COLO\_829-EBI.all.bcf - varlociraptor-manta/COLO\_829-GSC.all.bcf - varlociraptor-manta/COLO\_829-Ill.all.bcf - varlociraptor-manta/COLO\_829-TGen.all.bcf - varlociraptor-manta/COLO\_829-EBI.all.bcf - varlociraptor-strelka/COLO\_829-GSC.all.bcf - varlociraptor-strelka/COLO\_829-Ill.all.bcf - varlociraptor-strelka/COLO\_829-TGen.all.bcf - varlociraptor-strelka/COLO\_829-EBI.all.bcf - varlociraptor-bpi/COLO\_829-GSC.all.bcf - varlociraptor-bpi/COLO\_829-Ill.all.bcf - varlociraptor-bpi/COLO\_829-TGen.all.bcf - varlociraptor-bpi/COLO\_829-EBI.all.bcf - varlociraptor-delly/simulated-bwa.all.bcf - varlociraptor-lancet/simulated-bwa.all.bcf - varlociraptor-manta/simulated-bwa.all.bcf - varlociraptor-strelka/simulated-bwa.all.bcf |  | - bcftools ==1.5 | ``` 1 2 3 4 5 6 7 8 9 10 11 12 ``` \| ``` __author__ = "Johannes Köster" __copyright__ = "Copyright 2016, Johannes Köster" __email__ = "" __license__ = "MIT" from snakemake.shell import shell shell( "bcftools concat {snakemake.params} -o {snakemake.output[0]} " "{snakemake.input}") ``` |
| varlociraptor\_control\_fdr | 368 | - varlociraptor-delly/simulated-bwa.INS.1-30.1.0.bcf - varlociraptor-delly/simulated-bwa.INS.30-100.1.0.bcf - varlociraptor-lancet/simulated-bwa.INS.1-30.1.0.bcf - varlociraptor-lancet/simulated-bwa.INS.30-100.1.0.bcf - varlociraptor-manta/simulated-bwa.INS.1-30.1.0.bcf - varlociraptor-manta/simulated-bwa.INS.30-100.1.0.bcf - varlociraptor-strelka/simulated-bwa.INS.1-30.1.0.bcf - varlociraptor-strelka/simulated-bwa.INS.30-100.1.0.bcf - varlociraptor-delly/simulated-bwa.DEL.1-30.1.0.bcf - varlociraptor-delly/simulated-bwa.DEL.30-50.1.0.bcf - varlociraptor-delly/simulated-bwa.DEL.50-100.1.0.bcf - varlociraptor-delly/simulated-bwa.DEL.100-250.1.0.bcf - varlociraptor-lancet/simulated-bwa.DEL.1-30.1.0.bcf - varlociraptor-lancet/simulated-bwa.DEL.30-50.1.0.bcf - varlociraptor-lancet/simulated-bwa.DEL.50-100.1.0.bcf - varlociraptor-lancet/simulated-bwa.DEL.100-250.1.0.bcf - varlociraptor-manta/simulated-bwa.DEL.1-30.1.0.bcf - varlociraptor-manta/simulated-bwa.DEL.30-50.1.0.bcf - varlociraptor-manta/simulated-bwa.DEL.50-100.1.0.bcf - varlociraptor-manta/simulated-bwa.DEL.100-250.1.0.bcf - varlociraptor-strelka/simulated-bwa.DEL.1-30.1.0.bcf - varlociraptor-strelka/simulated-bwa.DEL.30-50.1.0.bcf - varlociraptor-strelka/simulated-bwa.DEL.50-100.1.0.bcf - varlociraptor-strelka/simulated-bwa.DEL.100-250.1.0.bcf - varlociraptor-delly/simulated-bwa.INS.1-30.1e-05.bcf - varlociraptor-delly/simulated-bwa.INS.1-30.0.0001.bcf - varlociraptor-delly/simulated-bwa.INS.1-30.0.001.bcf - varlociraptor-delly/simulated-bwa.INS.1-30.0.01.bcf - varlociraptor-delly/simulated-bwa.INS.1-30.0.05.bcf - varlociraptor-delly/simulated-bwa.INS.1-30.0.1.bcf - varlociraptor-delly/simulated-bwa.INS.1-30.0.2.bcf - varlociraptor-delly/simulated-bwa.INS.1-30.0.3.bcf - varlociraptor-delly/simulated-bwa.INS.1-30.0.4.bcf - varlociraptor-delly/simulated-bwa.INS.1-30.0.5.bcf - varlociraptor-delly/simulated-bwa.INS.1-30.0.6.bcf - varlociraptor-delly/simulated-bwa.INS.1-30.0.7.bcf - varlociraptor-delly/simulated-bwa.INS.1-30.0.8.bcf - varlociraptor-delly/simulated-bwa.INS.1-30.0.9.bcf - varlociraptor-delly/simulated-bwa.INS.30-100.1e-05.bcf - varlociraptor-delly/simulated-bwa.INS.30-100.0.0001.bcf - varlociraptor-delly/simulated-bwa.INS.30-100.0.001.bcf - varlociraptor-delly/simulated-bwa.INS.30-100.0.01.bcf - varlociraptor-delly/simulated-bwa.INS.30-100.0.05.bcf - varlociraptor-delly/simulated-bwa.INS.30-100.0.1.bcf - varlociraptor-delly/simulated-bwa.INS.30-100.0.2.bcf - varlociraptor-delly/simulated-bwa.INS.30-100.0.3.bcf - varlociraptor-delly/simulated-bwa.INS.30-100.0.4.bcf - varlociraptor-delly/simulated-bwa.INS.30-100.0.5.bcf - varlociraptor-delly/simulated-bwa.INS.30-100.0.6.bcf - varlociraptor-delly/simulated-bwa.INS.30-100.0.7.bcf - varlociraptor-delly/simulated-bwa.INS.30-100.0.8.bcf - varlociraptor-delly/simulated-bwa.INS.30-100.0.9.bcf - varlociraptor-lancet/simulated-bwa.INS.1-30.1e-05.bcf - varlociraptor-lancet/simulated-bwa.INS.1-30.0.0001.bcf - varlociraptor-lancet/simulated-bwa.INS.1-30.0.001.bcf - varlociraptor-lancet/simulated-bwa.INS.1-30.0.01.bcf - varlociraptor-lancet/simulated-bwa.INS.1-30.0.05.bcf - varlociraptor-lancet/simulated-bwa.INS.1-30.0.1.bcf - varlociraptor-lancet/simulated-bwa.INS.1-30.0.2.bcf - varlociraptor-lancet/simulated-bwa.INS.1-30.0.3.bcf - varlociraptor-lancet/simulated-bwa.INS.1-30.0.4.bcf - varlociraptor-lancet/simulated-bwa.INS.1-30.0.5.bcf - varlociraptor-lancet/simulated-bwa.INS.1-30.0.6.bcf - varlociraptor-lancet/simulated-bwa.INS.1-30.0.7.bcf - varlociraptor-lancet/simulated-bwa.INS.1-30.0.8.bcf - varlociraptor-lancet/simulated-bwa.INS.1-30.0.9.bcf - varlociraptor-lancet/simulated-bwa.INS.30-100.1e-05.bcf - varlociraptor-lancet/simulated-bwa.INS.30-100.0.0001.bcf - varlociraptor-lancet/simulated-bwa.INS.30-100.0.001.bcf - varlociraptor-lancet/simulated-bwa.INS.30-100.0.01.bcf - varlociraptor-lancet/simulated-bwa.INS.30-100.0.05.bcf - varlociraptor-lancet/simulated-bwa.INS.30-100.0.1.bcf - varlociraptor-lancet/simulated-bwa.INS.30-100.0.2.bcf - varlociraptor-lancet/simulated-bwa.INS.30-100.0.3.bcf - varlociraptor-lancet/simulated-bwa.INS.30-100.0.4.bcf - varlociraptor-lancet/simulated-bwa.INS.30-100.0.5.bcf - varlociraptor-lancet/simulated-bwa.INS.30-100.0.6.bcf - varlociraptor-lancet/simulated-bwa.INS.30-100.0.7.bcf - varlociraptor-lancet/simulated-bwa.INS.30-100.0.8.bcf - varlociraptor-lancet/simulated-bwa.INS.30-100.0.9.bcf - varlociraptor-manta/simulated-bwa.INS.1-30.1e-05.bcf - varlociraptor-manta/simulated-bwa.INS.1-30.0.0001.bcf - varlociraptor-manta/simulated-bwa.INS.1-30.0.001.bcf - varlociraptor-manta/simulated-bwa.INS.1-30.0.01.bcf - varlociraptor-manta/simulated-bwa.INS.1-30.0.05.bcf - varlociraptor-manta/simulated-bwa.INS.1-30.0.1.bcf - varlociraptor-manta/simulated-bwa.INS.1-30.0.2.bcf - varlociraptor-manta/simulated-bwa.INS.1-30.0.3.bcf - varlociraptor-manta/simulated-bwa.INS.1-30.0.4.bcf - varlociraptor-manta/simulated-bwa.INS.1-30.0.5.bcf - varlociraptor-manta/simulated-bwa.INS.1-30.0.6.bcf - varlociraptor-manta/simulated-bwa.INS.1-30.0.7.bcf - varlociraptor-manta/simulated-bwa.INS.1-30.0.8.bcf - varlociraptor-manta/simulated-bwa.INS.1-30.0.9.bcf - varlociraptor-manta/simulated-bwa.INS.30-100.1e-05.bcf - varlociraptor-manta/simulated-bwa.INS.30-100.0.0001.bcf - varlociraptor-manta/simulated-bwa.INS.30-100.0.001.bcf - varlociraptor-manta/simulated-bwa.INS.30-100.0.01.bcf - varlociraptor-manta/simulated-bwa.INS.30-100.0.05.bcf - varlociraptor-manta/simulated-bwa.INS.30-100.0.1.bcf - varlociraptor-manta/simulated-bwa.INS.30-100.0.2.bcf - varlociraptor-manta/simulated-bwa.INS.30-100.0.3.bcf - varlociraptor-manta/simulated-bwa.INS.30-100.0.4.bcf - varlociraptor-manta/simulated-bwa.INS.30-100.0.5.bcf - varlociraptor-manta/simulated-bwa.INS.30-100.0.6.bcf - varlociraptor-manta/simulated-bwa.INS.30-100.0.7.bcf - varlociraptor-manta/simulated-bwa.INS.30-100.0.8.bcf - varlociraptor-manta/simulated-bwa.INS.30-100.0.9.bcf - varlociraptor-strelka/simulated-bwa.INS.1-30.1e-05.bcf - varlociraptor-strelka/simulated-bwa.INS.1-30.0.0001.bcf - varlociraptor-strelka/simulated-bwa.INS.1-30.0.001.bcf - varlociraptor-strelka/simulated-bwa.INS.1-30.0.01.bcf - varlociraptor-strelka/simulated-bwa.INS.1-30.0.05.bcf - varlociraptor-strelka/simulated-bwa.INS.1-30.0.1.bcf - varlociraptor-strelka/simulated-bwa.INS.1-30.0.2.bcf - varlociraptor-strelka/simulated-bwa.INS.1-30.0.3.bcf - varlociraptor-strelka/simulated-bwa.INS.1-30.0.4.bcf - varlociraptor-strelka/simulated-bwa.INS.1-30.0.5.bcf - varlociraptor-strelka/simulated-bwa.INS.1-30.0.6.bcf - varlociraptor-strelka/simulated-bwa.INS.1-30.0.7.bcf - varlociraptor-strelka/simulated-bwa.INS.1-30.0.8.bcf - varlociraptor-strelka/simulated-bwa.INS.1-30.0.9.bcf - varlociraptor-strelka/simulated-bwa.INS.30-100.1e-05.bcf - varlociraptor-strelka/simulated-bwa.INS.30-100.0.0001.bcf - varlociraptor-strelka/simulated-bwa.INS.30-100.0.001.bcf - varlociraptor-strelka/simulated-bwa.INS.30-100.0.01.bcf - varlociraptor-strelka/simulated-bwa.INS.30-100.0.05.bcf - varlociraptor-strelka/simulated-bwa.INS.30-100.0.1.bcf - varlociraptor-strelka/simulated-bwa.INS.30-100.0.2.bcf - varlociraptor-strelka/simulated-bwa.INS.30-100.0.3.bcf - varlociraptor-strelka/simulated-bwa.INS.30-100.0.4.bcf - varlociraptor-strelka/simulated-bwa.INS.30-100.0.5.bcf - varlociraptor-strelka/simulated-bwa.INS.30-100.0.6.bcf - varlociraptor-strelka/simulated-bwa.INS.30-100.0.7.bcf - varlociraptor-strelka/simulated-bwa.INS.30-100.0.8.bcf - varlociraptor-strelka/simulated-bwa.INS.30-100.0.9.bcf - varlociraptor-delly/simulated-bwa.DEL.1-30.1e-05.bcf - varlociraptor-delly/simulated-bwa.DEL.1-30.0.0001.bcf - varlociraptor-delly/simulated-bwa.DEL.1-30.0.001.bcf - varlociraptor-delly/simulated-bwa.DEL.1-30.0.01.bcf - varlociraptor-delly/simulated-bwa.DEL.1-30.0.05.bcf - varlociraptor-delly/simulated-bwa.DEL.1-30.0.1.bcf - varlociraptor-delly/simulated-bwa.DEL.1-30.0.2.bcf - varlociraptor-delly/simulated-bwa.DEL.1-30.0.3.bcf - varlociraptor-delly/simulated-bwa.DEL.1-30.0.4.bcf - varlociraptor-delly/simulated-bwa.DEL.1-30.0.5.bcf - varlociraptor-delly/simulated-bwa.DEL.1-30.0.6.bcf - varlociraptor-delly/simulated-bwa.DEL.1-30.0.7.bcf - varlociraptor-delly/simulated-bwa.DEL.1-30.0.8.bcf - varlociraptor-delly/simulated-bwa.DEL.1-30.0.9.bcf - varlociraptor-delly/simulated-bwa.DEL.30-50.1e-05.bcf - varlociraptor-delly/simulated-bwa.DEL.30-50.0.0001.bcf - varlociraptor-delly/simulated-bwa.DEL.30-50.0.001.bcf - varlociraptor-delly/simulated-bwa.DEL.30-50.0.01.bcf - varlociraptor-delly/simulated-bwa.DEL.30-50.0.05.bcf - varlociraptor-delly/simulated-bwa.DEL.30-50.0.1.bcf - varlociraptor-delly/simulated-bwa.DEL.30-50.0.2.bcf - varlociraptor-delly/simulated-bwa.DEL.30-50.0.3.bcf - varlociraptor-delly/simulated-bwa.DEL.30-50.0.4.bcf - varlociraptor-delly/simulated-bwa.DEL.30-50.0.5.bcf - varlociraptor-delly/simulated-bwa.DEL.30-50.0.6.bcf - varlociraptor-delly/simulated-bwa.DEL.30-50.0.7.bcf - varlociraptor-delly/simulated-bwa.DEL.30-50.0.8.bcf - varlociraptor-delly/simulated-bwa.DEL.30-50.0.9.bcf - varlociraptor-delly/simulated-bwa.DEL.50-100.1e-05.bcf - varlociraptor-delly/simulated-bwa.DEL.50-100.0.0001.bcf - varlociraptor-delly/simulated-bwa.DEL.50-100.0.001.bcf - varlociraptor-delly/simulated-bwa.DEL.50-100.0.01.bcf - varlociraptor-delly/simulated-bwa.DEL.50-100.0.05.bcf - varlociraptor-delly/simulated-bwa.DEL.50-100.0.1.bcf - varlociraptor-delly/simulated-bwa.DEL.50-100.0.2.bcf - varlociraptor-delly/simulated-bwa.DEL.50-100.0.3.bcf - varlociraptor-delly/simulated-bwa.DEL.50-100.0.4.bcf - varlociraptor-delly/simulated-bwa.DEL.50-100.0.5.bcf - varlociraptor-delly/simulated-bwa.DEL.50-100.0.6.bcf - varlociraptor-delly/simulated-bwa.DEL.50-100.0.7.bcf - varlociraptor-delly/simulated-bwa.DEL.50-100.0.8.bcf - varlociraptor-delly/simulated-bwa.DEL.50-100.0.9.bcf - varlociraptor-delly/simulated-bwa.DEL.100-250.1e-05.bcf - varlociraptor-delly/simulated-bwa.DEL.100-250.0.0001.bcf - varlociraptor-delly/simulated-bwa.DEL.100-250.0.001.bcf - varlociraptor-delly/simulated-bwa.DEL.100-250.0.01.bcf - varlociraptor-delly/simulated-bwa.DEL.100-250.0.05.bcf - varlociraptor-delly/simulated-bwa.DEL.100-250.0.1.bcf - varlociraptor-delly/simulated-bwa.DEL.100-250.0.2.bcf - varlociraptor-delly/simulated-bwa.DEL.100-250.0.3.bcf - varlociraptor-delly/simulated-bwa.DEL.100-250.0.4.bcf - varlociraptor-delly/simulated-bwa.DEL.100-250.0.5.bcf - varlociraptor-delly/simulated-bwa.DEL.100-250.0.6.bcf - varlociraptor-delly/simulated-bwa.DEL.100-250.0.7.bcf - varlociraptor-delly/simulated-bwa.DEL.100-250.0.8.bcf - varlociraptor-delly/simulated-bwa.DEL.100-250.0.9.bcf - varlociraptor-lancet/simulated-bwa.DEL.1-30.1e-05.bcf - varlociraptor-lancet/simulated-bwa.DEL.1-30.0.0001.bcf - varlociraptor-lancet/simulated-bwa.DEL.1-30.0.001.bcf - varlociraptor-lancet/simulated-bwa.DEL.1-30.0.01.bcf - varlociraptor-lancet/simulated-bwa.DEL.1-30.0.05.bcf - varlociraptor-lancet/simulated-bwa.DEL.1-30.0.1.bcf - varlociraptor-lancet/simulated-bwa.DEL.1-30.0.2.bcf - varlociraptor-lancet/simulated-bwa.DEL.1-30.0.3.bcf - varlociraptor-lancet/simulated-bwa.DEL.1-30.0.4.bcf - varlociraptor-lancet/simulated-bwa.DEL.1-30.0.5.bcf - varlociraptor-lancet/simulated-bwa.DEL.1-30.0.6.bcf - varlociraptor-lancet/simulated-bwa.DEL.1-30.0.7.bcf - varlociraptor-lancet/simulated-bwa.DEL.1-30.0.8.bcf - varlociraptor-lancet/simulated-bwa.DEL.1-30.0.9.bcf - varlociraptor-lancet/simulated-bwa.DEL.30-50.1e-05.bcf - varlociraptor-lancet/simulated-bwa.DEL.30-50.0.0001.bcf - varlociraptor-lancet/simulated-bwa.DEL.30-50.0.001.bcf - varlociraptor-lancet/simulated-bwa.DEL.30-50.0.01.bcf - varlociraptor-lancet/simulated-bwa.DEL.30-50.0.05.bcf - varlociraptor-lancet/simulated-bwa.DEL.30-50.0.1.bcf - varlociraptor-lancet/simulated-bwa.DEL.30-50.0.2.bcf - varlociraptor-lancet/simulated-bwa.DEL.30-50.0.3.bcf - varlociraptor-lancet/simulated-bwa.DEL.30-50.0.4.bcf - varlociraptor-lancet/simulated-bwa.DEL.30-50.0.5.bcf - varlociraptor-lancet/simulated-bwa.DEL.30-50.0.6.bcf - varlociraptor-lancet/simulated-bwa.DEL.30-50.0.7.bcf - varlociraptor-lancet/simulated-bwa.DEL.30-50.0.8.bcf - varlociraptor-lancet/simulated-bwa.DEL.30-50.0.9.bcf - varlociraptor-lancet/simulated-bwa.DEL.50-100.1e-05.bcf - varlociraptor-lancet/simulated-bwa.DEL.50-100.0.0001.bcf - varlociraptor-lancet/simulated-bwa.DEL.50-100.0.001.bcf - varlociraptor-lancet/simulated-bwa.DEL.50-100.0.01.bcf - varlociraptor-lancet/simulated-bwa.DEL.50-100.0.05.bcf - varlociraptor-lancet/simulated-bwa.DEL.50-100.0.1.bcf - varlociraptor-lancet/simulated-bwa.DEL.50-100.0.2.bcf - varlociraptor-lancet/simulated-bwa.DEL.50-100.0.3.bcf - varlociraptor-lancet/simulated-bwa.DEL.50-100.0.4.bcf - varlociraptor-lancet/simulated-bwa.DEL.50-100.0.5.bcf - varlociraptor-lancet/simulated-bwa.DEL.50-100.0.6.bcf - varlociraptor-lancet/simulated-bwa.DEL.50-100.0.7.bcf - varlociraptor-lancet/simulated-bwa.DEL.50-100.0.8.bcf - varlociraptor-lancet/simulated-bwa.DEL.50-100.0.9.bcf - varlociraptor-lancet/simulated-bwa.DEL.100-250.1e-05.bcf - varlociraptor-lancet/simulated-bwa.DEL.100-250.0.0001.bcf - varlociraptor-lancet/simulated-bwa.DEL.100-250.0.001.bcf - varlociraptor-lancet/simulated-bwa.DEL.100-250.0.01.bcf - varlociraptor-lancet/simulated-bwa.DEL.100-250.0.05.bcf - varlociraptor-lancet/simulated-bwa.DEL.100-250.0.1.bcf - varlociraptor-lancet/simulated-bwa.DEL.100-250.0.2.bcf - varlociraptor-lancet/simulated-bwa.DEL.100-250.0.3.bcf - varlociraptor-lancet/simulated-bwa.DEL.100-250.0.4.bcf - varlociraptor-lancet/simulated-bwa.DEL.100-250.0.5.bcf - varlociraptor-lancet/simulated-bwa.DEL.100-250.0.6.bcf - varlociraptor-lancet/simulated-bwa.DEL.100-250.0.7.bcf - varlociraptor-lancet/simulated-bwa.DEL.100-250.0.8.bcf - varlociraptor-lancet/simulated-bwa.DEL.100-250.0.9.bcf - varlociraptor-manta/simulated-bwa.DEL.1-30.1e-05.bcf - varlociraptor-manta/simulated-bwa.DEL.1-30.0.0001.bcf - varlociraptor-manta/simulated-bwa.DEL.1-30.0.001.bcf - varlociraptor-manta/simulated-bwa.DEL.1-30.0.01.bcf - varlociraptor-manta/simulated-bwa.DEL.1-30.0.05.bcf - varlociraptor-manta/simulated-bwa.DEL.1-30.0.1.bcf - varlociraptor-manta/simulated-bwa.DEL.1-30.0.2.bcf - varlociraptor-manta/simulated-bwa.DEL.1-30.0.3.bcf - varlociraptor-manta/simulated-bwa.DEL.1-30.0.4.bcf - varlociraptor-manta/simulated-bwa.DEL.1-30.0.5.bcf - varlociraptor-manta/simulated-bwa.DEL.1-30.0.6.bcf - varlociraptor-manta/simulated-bwa.DEL.1-30.0.7.bcf - varlociraptor-manta/simulated-bwa.DEL.1-30.0.8.bcf - varlociraptor-manta/simulated-bwa.DEL.1-30.0.9.bcf - varlociraptor-manta/simulated-bwa.DEL.30-50.1e-05.bcf - varlociraptor-manta/simulated-bwa.DEL.30-50.0.0001.bcf - varlociraptor-manta/simulated-bwa.DEL.30-50.0.001.bcf - varlociraptor-manta/simulated-bwa.DEL.30-50.0.01.bcf - varlociraptor-manta/simulated-bwa.DEL.30-50.0.05.bcf - varlociraptor-manta/simulated-bwa.DEL.30-50.0.1.bcf - varlociraptor-manta/simulated-bwa.DEL.30-50.0.2.bcf - varlociraptor-manta/simulated-bwa.DEL.30-50.0.3.bcf - varlociraptor-manta/simulated-bwa.DEL.30-50.0.4.bcf - varlociraptor-manta/simulated-bwa.DEL.30-50.0.5.bcf - varlociraptor-manta/simulated-bwa.DEL.30-50.0.6.bcf - varlociraptor-manta/simulated-bwa.DEL.30-50.0.7.bcf - varlociraptor-manta/simulated-bwa.DEL.30-50.0.8.bcf - varlociraptor-manta/simulated-bwa.DEL.30-50.0.9.bcf - varlociraptor-manta/simulated-bwa.DEL.50-100.1e-05.bcf - varlociraptor-manta/simulated-bwa.DEL.50-100.0.0001.bcf - varlociraptor-manta/simulated-bwa.DEL.50-100.0.001.bcf - varlociraptor-manta/simulated-bwa.DEL.50-100.0.01.bcf - varlociraptor-manta/simulated-bwa.DEL.50-100.0.05.bcf - varlociraptor-manta/simulated-bwa.DEL.50-100.0.1.bcf - varlociraptor-manta/simulated-bwa.DEL.50-100.0.2.bcf - varlociraptor-manta/simulated-bwa.DEL.50-100.0.3.bcf - varlociraptor-manta/simulated-bwa.DEL.50-100.0.4.bcf - varlociraptor-manta/simulated-bwa.DEL.50-100.0.5.bcf - varlociraptor-manta/simulated-bwa.DEL.50-100.0.6.bcf - varlociraptor-manta/simulated-bwa.DEL.50-100.0.7.bcf - varlociraptor-manta/simulated-bwa.DEL.50-100.0.8.bcf - varlociraptor-manta/simulated-bwa.DEL.50-100.0.9.bcf - varlociraptor-manta/simulated-bwa.DEL.100-250.1e-05.bcf - varlociraptor-manta/simulated-bwa.DEL.100-250.0.0001.bcf - varlociraptor-manta/simulated-bwa.DEL.100-250.0.001.bcf - varlociraptor-manta/simulated-bwa.DEL.100-250.0.01.bcf - varlociraptor-manta/simulated-bwa.DEL.100-250.0.05.bcf - varlociraptor-manta/simulated-bwa.DEL.100-250.0.1.bcf - varlociraptor-manta/simulated-bwa.DEL.100-250.0.2.bcf - varlociraptor-manta/simulated-bwa.DEL.100-250.0.3.bcf - varlociraptor-manta/simulated-bwa.DEL.100-250.0.4.bcf - varlociraptor-manta/simulated-bwa.DEL.100-250.0.5.bcf - varlociraptor-manta/simulated-bwa.DEL.100-250.0.6.bcf - varlociraptor-manta/simulated-bwa.DEL.100-250.0.7.bcf - varlociraptor-manta/simulated-bwa.DEL.100-250.0.8.bcf - varlociraptor-manta/simulated-bwa.DEL.100-250.0.9.bcf - varlociraptor-strelka/simulated-bwa.DEL.1-30.1e-05.bcf - varlociraptor-strelka/simulated-bwa.DEL.1-30.0.0001.bcf - varlociraptor-strelka/simulated-bwa.DEL.1-30.0.001.bcf - varlociraptor-strelka/simulated-bwa.DEL.1-30.0.01.bcf - varlociraptor-strelka/simulated-bwa.DEL.1-30.0.05.bcf - varlociraptor-strelka/simulated-bwa.DEL.1-30.0.1.bcf - varlociraptor-strelka/simulated-bwa.DEL.1-30.0.2.bcf - varlociraptor-strelka/simulated-bwa.DEL.1-30.0.3.bcf - varlociraptor-strelka/simulated-bwa.DEL.1-30.0.4.bcf - varlociraptor-strelka/simulated-bwa.DEL.1-30.0.5.bcf - varlociraptor-strelka/simulated-bwa.DEL.1-30.0.6.bcf - varlociraptor-strelka/simulated-bwa.DEL.1-30.0.7.bcf - varlociraptor-strelka/simulated-bwa.DEL.1-30.0.8.bcf - varlociraptor-strelka/simulated-bwa.DEL.1-30.0.9.bcf - varlociraptor-strelka/simulated-bwa.DEL.30-50.1e-05.bcf - varlociraptor-strelka/simulated-bwa.DEL.30-50.0.0001.bcf - varlociraptor-strelka/simulated-bwa.DEL.30-50.0.001.bcf - varlociraptor-strelka/simulated-bwa.DEL.30-50.0.01.bcf - varlociraptor-strelka/simulated-bwa.DEL.30-50.0.05.bcf - varlociraptor-strelka/simulated-bwa.DEL.30-50.0.1.bcf - varlociraptor-strelka/simulated-bwa.DEL.30-50.0.2.bcf - varlociraptor-strelka/simulated-bwa.DEL.30-50.0.3.bcf - varlociraptor-strelka/simulated-bwa.DEL.30-50.0.4.bcf - varlociraptor-strelka/simulated-bwa.DEL.30-50.0.5.bcf - varlociraptor-strelka/simulated-bwa.DEL.30-50.0.6.bcf - varlociraptor-strelka/simulated-bwa.DEL.30-50.0.7.bcf - varlociraptor-strelka/simulated-bwa.DEL.30-50.0.8.bcf - varlociraptor-strelka/simulated-bwa.DEL.30-50.0.9.bcf - varlociraptor-strelka/simulated-bwa.DEL.50-100.1e-05.bcf - varlociraptor-strelka/simulated-bwa.DEL.50-100.0.0001.bcf - varlociraptor-strelka/simulated-bwa.DEL.50-100.0.001.bcf - varlociraptor-strelka/simulated-bwa.DEL.50-100.0.01.bcf - varlociraptor-strelka/simulated-bwa.DEL.50-100.0.05.bcf - varlociraptor-strelka/simulated-bwa.DEL.50-100.0.1.bcf - varlociraptor-strelka/simulated-bwa.DEL.50-100.0.2.bcf - varlociraptor-strelka/simulated-bwa.DEL.50-100.0.3.bcf - varlociraptor-strelka/simulated-bwa.DEL.50-100.0.4.bcf - varlociraptor-strelka/simulated-bwa.DEL.50-100.0.5.bcf - varlociraptor-strelka/simulated-bwa.DEL.50-100.0.6.bcf - varlociraptor-strelka/simulated-bwa.DEL.50-100.0.7.bcf - varlociraptor-strelka/simulated-bwa.DEL.50-100.0.8.bcf - varlociraptor-strelka/simulated-bwa.DEL.50-100.0.9.bcf - varlociraptor-strelka/simulated-bwa.DEL.100-250.1e-05.bcf - varlociraptor-strelka/simulated-bwa.DEL.100-250.0.0001.bcf - varlociraptor-strelka/simulated-bwa.DEL.100-250.0.001.bcf - varlociraptor-strelka/simulated-bwa.DEL.100-250.0.01.bcf - varlociraptor-strelka/simulated-bwa.DEL.100-250.0.05.bcf - varlociraptor-strelka/simulated-bwa.DEL.100-250.0.1.bcf - varlociraptor-strelka/simulated-bwa.DEL.100-250.0.2.bcf - varlociraptor-strelka/simulated-bwa.DEL.100-250.0.3.bcf - varlociraptor-strelka/simulated-bwa.DEL.100-250.0.4.bcf - varlociraptor-strelka/simulated-bwa.DEL.100-250.0.5.bcf - varlociraptor-strelka/simulated-bwa.DEL.100-250.0.6.bcf - varlociraptor-strelka/simulated-bwa.DEL.100-250.0.7.bcf - varlociraptor-strelka/simulated-bwa.DEL.100-250.0.8.bcf - varlociraptor-strelka/simulated-bwa.DEL.100-250.0.9.bcf - varlociraptor-delly/simulated-bwa.INS.1-250.1.0.bcf - varlociraptor-lancet/simulated-bwa.INS.1-250.1.0.bcf - varlociraptor-manta/simulated-bwa.INS.1-250.1.0.bcf - varlociraptor-strelka/simulated-bwa.INS.1-250.1.0.bcf - varlociraptor-delly/simulated-bwa.DEL.1-250.1.0.bcf - varlociraptor-lancet/simulated-bwa.DEL.1-250.1.0.bcf - varlociraptor-manta/simulated-bwa.DEL.1-250.1.0.bcf - varlociraptor-strelka/simulated-bwa.DEL.1-250.1.0.bcf |  |  | ``` 1 ``` \| ``` varlociraptor filter-calls control-fdr {input} --events SOMATIC_TUMOR --var {wildcards.type} --minlen {wildcards.minlen} --maxlen {wildcards.maxlen} --fdr {wildcards.fdr} > {output} ``` |
| adhoc\_varlociraptor | 40 | - varlociraptor-delly/COLO\_829-GSC.adhoc.0.9.bcf - varlociraptor-delly/COLO\_829-Ill.adhoc.0.9.bcf - varlociraptor-delly/COLO\_829-TGen.adhoc.0.9.bcf - varlociraptor-delly/COLO\_829-EBI.adhoc.0.9.bcf - varlociraptor-lancet/COLO\_829-GSC.adhoc.0.9.bcf - varlociraptor-lancet/COLO\_829-Ill.adhoc.0.9.bcf - varlociraptor-lancet/COLO\_829-TGen.adhoc.0.9.bcf - varlociraptor-lancet/COLO\_829-EBI.adhoc.0.9.bcf - varlociraptor-manta/COLO\_829-GSC.adhoc.0.9.bcf - varlociraptor-manta/COLO\_829-Ill.adhoc.0.9.bcf - varlociraptor-manta/COLO\_829-TGen.adhoc.0.9.bcf - varlociraptor-manta/COLO\_829-EBI.adhoc.0.9.bcf - varlociraptor-strelka/COLO\_829-GSC.adhoc.0.9.bcf - varlociraptor-strelka/COLO\_829-Ill.adhoc.0.9.bcf - varlociraptor-strelka/COLO\_829-TGen.adhoc.0.9.bcf - varlociraptor-strelka/COLO\_829-EBI.adhoc.0.9.bcf - varlociraptor-bpi/COLO\_829-GSC.adhoc.0.9.bcf - varlociraptor-bpi/COLO\_829-Ill.adhoc.0.9.bcf - varlociraptor-bpi/COLO\_829-TGen.adhoc.0.9.bcf - varlociraptor-bpi/COLO\_829-EBI.adhoc.0.9.bcf - varlociraptor-delly/COLO\_829-GSC.adhoc.0.98.bcf - varlociraptor-delly/COLO\_829-Ill.adhoc.0.98.bcf - varlociraptor-delly/COLO\_829-TGen.adhoc.0.98.bcf - varlociraptor-delly/COLO\_829-EBI.adhoc.0.98.bcf - varlociraptor-lancet/COLO\_829-GSC.adhoc.0.98.bcf - varlociraptor-lancet/COLO\_829-Ill.adhoc.0.98.bcf - varlociraptor-lancet/COLO\_829-TGen.adhoc.0.98.bcf - varlociraptor-lancet/COLO\_829-EBI.adhoc.0.98.bcf - varlociraptor-manta/COLO\_829-GSC.adhoc.0.98.bcf - varlociraptor-manta/COLO\_829-Ill.adhoc.0.98.bcf - varlociraptor-manta/COLO\_829-TGen.adhoc.0.98.bcf - varlociraptor-manta/COLO\_829-EBI.adhoc.0.98.bcf - varlociraptor-strelka/COLO\_829-GSC.adhoc.0.98.bcf - varlociraptor-strelka/COLO\_829-Ill.adhoc.0.98.bcf - varlociraptor-strelka/COLO\_829-TGen.adhoc.0.98.bcf - varlociraptor-strelka/COLO\_829-EBI.adhoc.0.98.bcf - varlociraptor-bpi/COLO\_829-GSC.adhoc.0.98.bcf - varlociraptor-bpi/COLO\_829-Ill.adhoc.0.98.bcf - varlociraptor-bpi/COLO\_829-TGen.adhoc.0.98.bcf - varlociraptor-bpi/COLO\_829-EBI.adhoc.0.98.bcf |  | - bcftools ==1.5 | ``` 1 2 3 4 5 6 7 8 9 10 11 12 ``` \| ``` __author__ = "Johannes Köster" __copyright__ = "Copyright 2016, Johannes Köster" __email__ = "" __license__ = "MIT" from snakemake.shell import shell shell( "bcftools view {snakemake.params} {snakemake.input[0]} " "-o {snakemake.output[0]}") ``` |
| varlociraptor\_call | 600 | - varlociraptor-delly/COLO\_829-GSC.1.bcf - varlociraptor-delly/COLO\_829-GSC.2.bcf - varlociraptor-delly/COLO\_829-GSC.3.bcf - varlociraptor-delly/COLO\_829-GSC.4.bcf - varlociraptor-delly/COLO\_829-GSC.5.bcf - varlociraptor-delly/COLO\_829-GSC.6.bcf - varlociraptor-delly/COLO\_829-GSC.7.bcf - varlociraptor-delly/COLO\_829-GSC.8.bcf - varlociraptor-delly/COLO\_829-GSC.9.bcf - varlociraptor-delly/COLO\_829-GSC.10.bcf - varlociraptor-delly/COLO\_829-GSC.11.bcf - varlociraptor-delly/COLO\_829-GSC.12.bcf - varlociraptor-delly/COLO\_829-GSC.13.bcf - varlociraptor-delly/COLO\_829-GSC.14.bcf - varlociraptor-delly/COLO\_829-GSC.15.bcf - varlociraptor-delly/COLO\_829-GSC.16.bcf - varlociraptor-delly/COLO\_829-GSC.17.bcf - varlociraptor-delly/COLO\_829-GSC.18.bcf - varlociraptor-delly/COLO\_829-GSC.19.bcf - varlociraptor-delly/COLO\_829-GSC.20.bcf - varlociraptor-delly/COLO\_829-GSC.21.bcf - varlociraptor-delly/COLO\_829-GSC.22.bcf - varlociraptor-delly/COLO\_829-GSC.M.bcf - varlociraptor-delly/COLO\_829-GSC.X.bcf - varlociraptor-delly/COLO\_829-GSC.Y.bcf - varlociraptor-delly/COLO\_829-Ill.1.bcf - varlociraptor-delly/COLO\_829-Ill.2.bcf - varlociraptor-delly/COLO\_829-Ill.3.bcf - varlociraptor-delly/COLO\_829-Ill.4.bcf - varlociraptor-delly/COLO\_829-Ill.5.bcf - varlociraptor-delly/COLO\_829-Ill.6.bcf - varlociraptor-delly/COLO\_829-Ill.7.bcf - varlociraptor-delly/COLO\_829-Ill.8.bcf - varlociraptor-delly/COLO\_829-Ill.9.bcf - varlociraptor-delly/COLO\_829-Ill.10.bcf - varlociraptor-delly/COLO\_829-Ill.11.bcf - varlociraptor-delly/COLO\_829-Ill.12.bcf - varlociraptor-delly/COLO\_829-Ill.13.bcf - varlociraptor-delly/COLO\_829-Ill.14.bcf - varlociraptor-delly/COLO\_829-Ill.15.bcf - varlociraptor-delly/COLO\_829-Ill.16.bcf - varlociraptor-delly/COLO\_829-Ill.17.bcf - varlociraptor-delly/COLO\_829-Ill.18.bcf - varlociraptor-delly/COLO\_829-Ill.19.bcf - varlociraptor-delly/COLO\_829-Ill.20.bcf - varlociraptor-delly/COLO\_829-Ill.21.bcf - varlociraptor-delly/COLO\_829-Ill.22.bcf - varlociraptor-delly/COLO\_829-Ill.M.bcf - varlociraptor-delly/COLO\_829-Ill.X.bcf - varlociraptor-delly/COLO\_829-Ill.Y.bcf - varlociraptor-delly/COLO\_829-TGen.1.bcf - varlociraptor-delly/COLO\_829-TGen.2.bcf - varlociraptor-delly/COLO\_829-TGen.3.bcf - varlociraptor-delly/COLO\_829-TGen.4.bcf - varlociraptor-delly/COLO\_829-TGen.5.bcf - varlociraptor-delly/COLO\_829-TGen.6.bcf - varlociraptor-delly/COLO\_829-TGen.7.bcf - varlociraptor-delly/COLO\_829-TGen.8.bcf - varlociraptor-delly/COLO\_829-TGen.9.bcf - varlociraptor-delly/COLO\_829-TGen.10.bcf - varlociraptor-delly/COLO\_829-TGen.11.bcf - varlociraptor-delly/COLO\_829-TGen.12.bcf - varlociraptor-delly/COLO\_829-TGen.13.bcf - varlociraptor-delly/COLO\_829-TGen.14.bcf - varlociraptor-delly/COLO\_829-TGen.15.bcf - varlociraptor-delly/COLO\_829-TGen.16.bcf - varlociraptor-delly/COLO\_829-TGen.17.bcf - varlociraptor-delly/COLO\_829-TGen.18.bcf - varlociraptor-delly/COLO\_829-TGen.19.bcf - varlociraptor-delly/COLO\_829-TGen.20.bcf - varlociraptor-delly/COLO\_829-TGen.21.bcf - varlociraptor-delly/COLO\_829-TGen.22.bcf - varlociraptor-delly/COLO\_829-TGen.M.bcf - varlociraptor-delly/COLO\_829-TGen.X.bcf - varlociraptor-delly/COLO\_829-TGen.Y.bcf - varlociraptor-delly/COLO\_829-EBI.1.bcf - varlociraptor-delly/COLO\_829-EBI.2.bcf - varlociraptor-delly/COLO\_829-EBI.3.bcf - varlociraptor-delly/COLO\_829-EBI.4.bcf - varlociraptor-delly/COLO\_829-EBI.5.bcf - varlociraptor-delly/COLO\_829-EBI.6.bcf - varlociraptor-delly/COLO\_829-EBI.7.bcf - varlociraptor-delly/COLO\_829-EBI.8.bcf - varlociraptor-delly/COLO\_829-EBI.9.bcf - varlociraptor-delly/COLO\_829-EBI.10.bcf - varlociraptor-delly/COLO\_829-EBI.11.bcf - varlociraptor-delly/COLO\_829-EBI.12.bcf - varlociraptor-delly/COLO\_829-EBI.13.bcf - varlociraptor-delly/COLO\_829-EBI.14.bcf - varlociraptor-delly/COLO\_829-EBI.15.bcf - varlociraptor-delly/COLO\_829-EBI.16.bcf - varlociraptor-delly/COLO\_829-EBI.17.bcf - varlociraptor-delly/COLO\_829-EBI.18.bcf - varlociraptor-delly/COLO\_829-EBI.19.bcf - varlociraptor-delly/COLO\_829-EBI.20.bcf - varlociraptor-delly/COLO\_829-EBI.21.bcf - varlociraptor-delly/COLO\_829-EBI.22.bcf - varlociraptor-delly/COLO\_829-EBI.M.bcf - varlociraptor-delly/COLO\_829-EBI.X.bcf - varlociraptor-delly/COLO\_829-EBI.Y.bcf - varlociraptor-lancet/COLO\_829-GSC.1.bcf - varlociraptor-lancet/COLO\_829-GSC.2.bcf - varlociraptor-lancet/COLO\_829-GSC.3.bcf - varlociraptor-lancet/COLO\_829-GSC.4.bcf - varlociraptor-lancet/COLO\_829-GSC.5.bcf - varlociraptor-lancet/COLO\_829-GSC.6.bcf - varlociraptor-lancet/COLO\_829-GSC.7.bcf - varlociraptor-lancet/COLO\_829-GSC.8.bcf - varlociraptor-lancet/COLO\_829-GSC.9.bcf - varlociraptor-lancet/COLO\_829-GSC.10.bcf - varlociraptor-lancet/COLO\_829-GSC.11.bcf - varlociraptor-lancet/COLO\_829-GSC.12.bcf - varlociraptor-lancet/COLO\_829-GSC.13.bcf - varlociraptor-lancet/COLO\_829-GSC.14.bcf - varlociraptor-lancet/COLO\_829-GSC.15.bcf - varlociraptor-lancet/COLO\_829-GSC.16.bcf - varlociraptor-lancet/COLO\_829-GSC.17.bcf - varlociraptor-lancet/COLO\_829-GSC.18.bcf - varlociraptor-lancet/COLO\_829-GSC.19.bcf - varlociraptor-lancet/COLO\_829-GSC.20.bcf - varlociraptor-lancet/COLO\_829-GSC.21.bcf - varlociraptor-lancet/COLO\_829-GSC.22.bcf - varlociraptor-lancet/COLO\_829-GSC.M.bcf - varlociraptor-lancet/COLO\_829-GSC.X.bcf - varlociraptor-lancet/COLO\_829-GSC.Y.bcf - varlociraptor-lancet/COLO\_829-Ill.1.bcf - varlociraptor-lancet/COLO\_829-Ill.2.bcf - varlociraptor-lancet/COLO\_829-Ill.3.bcf - varlociraptor-lancet/COLO\_829-Ill.4.bcf - varlociraptor-lancet/COLO\_829-Ill.5.bcf - varlociraptor-lancet/COLO\_829-Ill.6.bcf - varlociraptor-lancet/COLO\_829-Ill.7.bcf - varlociraptor-lancet/COLO\_829-Ill.8.bcf - varlociraptor-lancet/COLO\_829-Ill.9.bcf - varlociraptor-lancet/COLO\_829-Ill.10.bcf - varlociraptor-lancet/COLO\_829-Ill.11.bcf - varlociraptor-lancet/COLO\_829-Ill.12.bcf - varlociraptor-lancet/COLO\_829-Ill.13.bcf - varlociraptor-lancet/COLO\_829-Ill.14.bcf - varlociraptor-lancet/COLO\_829-Ill.15.bcf - varlociraptor-lancet/COLO\_829-Ill.16.bcf - varlociraptor-lancet/COLO\_829-Ill.17.bcf - varlociraptor-lancet/COLO\_829-Ill.18.bcf - varlociraptor-lancet/COLO\_829-Ill.19.bcf - varlociraptor-lancet/COLO\_829-Ill.20.bcf - varlociraptor-lancet/COLO\_829-Ill.21.bcf - varlociraptor-lancet/COLO\_829-Ill.22.bcf - varlociraptor-lancet/COLO\_829-Ill.M.bcf - varlociraptor-lancet/COLO\_829-Ill.X.bcf - varlociraptor-lancet/COLO\_829-Ill.Y.bcf - varlociraptor-lancet/COLO\_829-TGen.1.bcf - varlociraptor-lancet/COLO\_829-TGen.2.bcf - varlociraptor-lancet/COLO\_829-TGen.3.bcf - varlociraptor-lancet/COLO\_829-TGen.4.bcf - varlociraptor-lancet/COLO\_829-TGen.5.bcf - varlociraptor-lancet/COLO\_829-TGen.6.bcf - varlociraptor-lancet/COLO\_829-TGen.7.bcf - varlociraptor-lancet/COLO\_829-TGen.8.bcf - varlociraptor-lancet/COLO\_829-TGen.9.bcf - varlociraptor-lancet/COLO\_829-TGen.10.bcf - varlociraptor-lancet/COLO\_829-TGen.11.bcf - varlociraptor-lancet/COLO\_829-TGen.12.bcf - varlociraptor-lancet/COLO\_829-TGen.13.bcf - varlociraptor-lancet/COLO\_829-TGen.14.bcf - varlociraptor-lancet/COLO\_829-TGen.15.bcf - varlociraptor-lancet/COLO\_829-TGen.16.bcf - varlociraptor-lancet/COLO\_829-TGen.17.bcf - varlociraptor-lancet/COLO\_829-TGen.18.bcf - varlociraptor-lancet/COLO\_829-TGen.19.bcf - varlociraptor-lancet/COLO\_829-TGen.20.bcf - varlociraptor-lancet/COLO\_829-TGen.21.bcf - varlociraptor-lancet/COLO\_829-TGen.22.bcf - varlociraptor-lancet/COLO\_829-TGen.M.bcf - varlociraptor-lancet/COLO\_829-TGen.X.bcf - varlociraptor-lancet/COLO\_829-TGen.Y.bcf - varlociraptor-lancet/COLO\_829-EBI.1.bcf - varlociraptor-lancet/COLO\_829-EBI.2.bcf - varlociraptor-lancet/COLO\_829-EBI.3.bcf - varlociraptor-lancet/COLO\_829-EBI.4.bcf - varlociraptor-lancet/COLO\_829-EBI.5.bcf - varlociraptor-lancet/COLO\_829-EBI.6.bcf - varlociraptor-lancet/COLO\_829-EBI.7.bcf - varlociraptor-lancet/COLO\_829-EBI.8.bcf - varlociraptor-lancet/COLO\_829-EBI.9.bcf - varlociraptor-lancet/COLO\_829-EBI.10.bcf - varlociraptor-lancet/COLO\_829-EBI.11.bcf - varlociraptor-lancet/COLO\_829-EBI.12.bcf - varlociraptor-lancet/COLO\_829-EBI.13.bcf - varlociraptor-lancet/COLO\_829-EBI.14.bcf - varlociraptor-lancet/COLO\_829-EBI.15.bcf - varlociraptor-lancet/COLO\_829-EBI.16.bcf - varlociraptor-lancet/COLO\_829-EBI.17.bcf - varlociraptor-lancet/COLO\_829-EBI.18.bcf - varlociraptor-lancet/COLO\_829-EBI.19.bcf - varlociraptor-lancet/COLO\_829-EBI.20.bcf - varlociraptor-lancet/COLO\_829-EBI.21.bcf - varlociraptor-lancet/COLO\_829-EBI.22.bcf - varlociraptor-lancet/COLO\_829-EBI.M.bcf - varlociraptor-lancet/COLO\_829-EBI.X.bcf - varlociraptor-lancet/COLO\_829-EBI.Y.bcf - varlociraptor-manta/COLO\_829-GSC.1.bcf - varlociraptor-manta/COLO\_829-GSC.2.bcf - varlociraptor-manta/COLO\_829-GSC.3.bcf - varlociraptor-manta/COLO\_829-GSC.4.bcf - varlociraptor-manta/COLO\_829-GSC.5.bcf - varlociraptor-manta/COLO\_829-GSC.6.bcf - varlociraptor-manta/COLO\_829-GSC.7.bcf - varlociraptor-manta/COLO\_829-GSC.8.bcf - varlociraptor-manta/COLO\_829-GSC.9.bcf - varlociraptor-manta/COLO\_829-GSC.10.bcf - varlociraptor-manta/COLO\_829-GSC.11.bcf - varlociraptor-manta/COLO\_829-GSC.12.bcf - varlociraptor-manta/COLO\_829-GSC.13.bcf - varlociraptor-manta/COLO\_829-GSC.14.bcf - varlociraptor-manta/COLO\_829-GSC.15.bcf - varlociraptor-manta/COLO\_829-GSC.16.bcf - varlociraptor-manta/COLO\_829-GSC.17.bcf - varlociraptor-manta/COLO\_829-GSC.18.bcf - varlociraptor-manta/COLO\_829-GSC.19.bcf - varlociraptor-manta/COLO\_829-GSC.20.bcf - varlociraptor-manta/COLO\_829-GSC.21.bcf - varlociraptor-manta/COLO\_829-GSC.22.bcf - varlociraptor-manta/COLO\_829-GSC.M.bcf - varlociraptor-manta/COLO\_829-GSC.X.bcf - varlociraptor-manta/COLO\_829-GSC.Y.bcf - varlociraptor-manta/COLO\_829-Ill.1.bcf - varlociraptor-manta/COLO\_829-Ill.2.bcf - varlociraptor-manta/COLO\_829-Ill.3.bcf - varlociraptor-manta/COLO\_829-Ill.4.bcf - varlociraptor-manta/COLO\_829-Ill.5.bcf - varlociraptor-manta/COLO\_829-Ill.6.bcf - varlociraptor-manta/COLO\_829-Ill.7.bcf - varlociraptor-manta/COLO\_829-Ill.8.bcf - varlociraptor-manta/COLO\_829-Ill.9.bcf - varlociraptor-manta/COLO\_829-Ill.10.bcf - varlociraptor-manta/COLO\_829-Ill.11.bcf - varlociraptor-manta/COLO\_829-Ill.12.bcf - varlociraptor-manta/COLO\_829-Ill.13.bcf - varlociraptor-manta/COLO\_829-Ill.14.bcf - varlociraptor-manta/COLO\_829-Ill.15.bcf - varlociraptor-manta/COLO\_829-Ill.16.bcf - varlociraptor-manta/COLO\_829-Ill.17.bcf - varlociraptor-manta/COLO\_829-Ill.18.bcf - varlociraptor-manta/COLO\_829-Ill.19.bcf - varlociraptor-manta/COLO\_829-Ill.20.bcf - varlociraptor-manta/COLO\_829-Ill.21.bcf - varlociraptor-manta/COLO\_829-Ill.22.bcf - varlociraptor-manta/COLO\_829-Ill.M.bcf - varlociraptor-manta/COLO\_829-Ill.X.bcf - varlociraptor-manta/COLO\_829-Ill.Y.bcf - varlociraptor-manta/COLO\_829-TGen.1.bcf - varlociraptor-manta/COLO\_829-TGen.2.bcf - varlociraptor-manta/COLO\_829-TGen.3.bcf - varlociraptor-manta/COLO\_829-TGen.4.bcf - varlociraptor-manta/COLO\_829-TGen.5.bcf - varlociraptor-manta/COLO\_829-TGen.6.bcf - varlociraptor-manta/COLO\_829-TGen.7.bcf - varlociraptor-manta/COLO\_829-TGen.8.bcf - varlociraptor-manta/COLO\_829-TGen.9.bcf - varlociraptor-manta/COLO\_829-TGen.10.bcf - varlociraptor-manta/COLO\_829-TGen.11.bcf - varlociraptor-manta/COLO\_829-TGen.12.bcf - varlociraptor-manta/COLO\_829-TGen.13.bcf - varlociraptor-manta/COLO\_829-TGen.14.bcf - varlociraptor-manta/COLO\_829-TGen.15.bcf - varlociraptor-manta/COLO\_829-TGen.16.bcf - varlociraptor-manta/COLO\_829-TGen.17.bcf - varlociraptor-manta/COLO\_829-TGen.18.bcf - varlociraptor-manta/COLO\_829-TGen.19.bcf - varlociraptor-manta/COLO\_829-TGen.20.bcf - varlociraptor-manta/COLO\_829-TGen.21.bcf - varlociraptor-manta/COLO\_829-TGen.22.bcf - varlociraptor-manta/COLO\_829-TGen.M.bcf - varlociraptor-manta/COLO\_829-TGen.X.bcf - varlociraptor-manta/COLO\_829-TGen.Y.bcf - varlociraptor-manta/COLO\_829-EBI.1.bcf - varlociraptor-manta/COLO\_829-EBI.2.bcf - varlociraptor-manta/COLO\_829-EBI.3.bcf - varlociraptor-manta/COLO\_829-EBI.4.bcf - varlociraptor-manta/COLO\_829-EBI.5.bcf - varlociraptor-manta/COLO\_829-EBI.6.bcf - varlociraptor-manta/COLO\_829-EBI.7.bcf - varlociraptor-manta/COLO\_829-EBI.8.bcf - varlociraptor-manta/COLO\_829-EBI.9.bcf - varlociraptor-manta/COLO\_829-EBI.10.bcf - varlociraptor-manta/COLO\_829-EBI.11.bcf - varlociraptor-manta/COLO\_829-EBI.12.bcf - varlociraptor-manta/COLO\_829-EBI.13.bcf - varlociraptor-manta/COLO\_829-EBI.14.bcf - varlociraptor-manta/COLO\_829-EBI.15.bcf - varlociraptor-manta/COLO\_829-EBI.16.bcf - varlociraptor-manta/COLO\_829-EBI.17.bcf - varlociraptor-manta/COLO\_829-EBI.18.bcf - varlociraptor-manta/COLO\_829-EBI.19.bcf - varlociraptor-manta/COLO\_829-EBI.20.bcf - varlociraptor-manta/COLO\_829-EBI.21.bcf - varlociraptor-manta/COLO\_829-EBI.22.bcf - varlociraptor-manta/COLO\_829-EBI.M.bcf - varlociraptor-manta/COLO\_829-EBI.X.bcf - varlociraptor-manta/COLO\_829-EBI.Y.bcf - varlociraptor-strelka/COLO\_829-GSC.1.bcf - varlociraptor-strelka/COLO\_829-GSC.2.bcf - varlociraptor-strelka/COLO\_829-GSC.3.bcf - varlociraptor-strelka/COLO\_829-GSC.4.bcf - varlociraptor-strelka/COLO\_829-GSC.5.bcf - varlociraptor-strelka/COLO\_829-GSC.6.bcf - varlociraptor-strelka/COLO\_829-GSC.7.bcf - varlociraptor-strelka/COLO\_829-GSC.8.bcf - varlociraptor-strelka/COLO\_829-GSC.9.bcf - varlociraptor-strelka/COLO\_829-GSC.10.bcf - varlociraptor-strelka/COLO\_829-GSC.11.bcf - varlociraptor-strelka/COLO\_829-GSC.12.bcf - varlociraptor-strelka/COLO\_829-GSC.13.bcf - varlociraptor-strelka/COLO\_829-GSC.14.bcf - varlociraptor-strelka/COLO\_829-GSC.15.bcf - varlociraptor-strelka/COLO\_829-GSC.16.bcf - varlociraptor-strelka/COLO\_829-GSC.17.bcf - varlociraptor-strelka/COLO\_829-GSC.18.bcf - varlociraptor-strelka/COLO\_829-GSC.19.bcf - varlociraptor-strelka/COLO\_829-GSC.20.bcf - varlociraptor-strelka/COLO\_829-GSC.21.bcf - varlociraptor-strelka/COLO\_829-GSC.22.bcf - varlociraptor-strelka/COLO\_829-GSC.M.bcf - varlociraptor-strelka/COLO\_829-GSC.X.bcf - varlociraptor-strelka/COLO\_829-GSC.Y.bcf - varlociraptor-strelka/COLO\_829-Ill.1.bcf - varlociraptor-strelka/COLO\_829-Ill.2.bcf - varlociraptor-strelka/COLO\_829-Ill.3.bcf - varlociraptor-strelka/COLO\_829-Ill.4.bcf - varlociraptor-strelka/COLO\_829-Ill.5.bcf - varlociraptor-strelka/COLO\_829-Ill.6.bcf - varlociraptor-strelka/COLO\_829-Ill.7.bcf - varlociraptor-strelka/COLO\_829-Ill.8.bcf - varlociraptor-strelka/COLO\_829-Ill.9.bcf - varlociraptor-strelka/COLO\_829-Ill.10.bcf - varlociraptor-strelka/COLO\_829-Ill.11.bcf - varlociraptor-strelka/COLO\_829-Ill.12.bcf - varlociraptor-strelka/COLO\_829-Ill.13.bcf - varlociraptor-strelka/COLO\_829-Ill.14.bcf - varlociraptor-strelka/COLO\_829-Ill.15.bcf - varlociraptor-strelka/COLO\_829-Ill.16.bcf - varlociraptor-strelka/COLO\_829-Ill.17.bcf - varlociraptor-strelka/COLO\_829-Ill.18.bcf - varlociraptor-strelka/COLO\_829-Ill.19.bcf - varlociraptor-strelka/COLO\_829-Ill.20.bcf - varlociraptor-strelka/COLO\_829-Ill.21.bcf - varlociraptor-strelka/COLO\_829-Ill.22.bcf - varlociraptor-strelka/COLO\_829-Ill.M.bcf - varlociraptor-strelka/COLO\_829-Ill.X.bcf - varlociraptor-strelka/COLO\_829-Ill.Y.bcf - varlociraptor-strelka/COLO\_829-TGen.1.bcf - varlociraptor-strelka/COLO\_829-TGen.2.bcf - varlociraptor-strelka/COLO\_829-TGen.3.bcf - varlociraptor-strelka/COLO\_829-TGen.4.bcf - varlociraptor-strelka/COLO\_829-TGen.5.bcf - varlociraptor-strelka/COLO\_829-TGen.6.bcf - varlociraptor-strelka/COLO\_829-TGen.7.bcf - varlociraptor-strelka/COLO\_829-TGen.8.bcf - varlociraptor-strelka/COLO\_829-TGen.9.bcf - varlociraptor-strelka/COLO\_829-TGen.10.bcf - varlociraptor-strelka/COLO\_829-TGen.11.bcf - varlociraptor-strelka/COLO\_829-TGen.12.bcf - varlociraptor-strelka/COLO\_829-TGen.13.bcf - varlociraptor-strelka/COLO\_829-TGen.14.bcf - varlociraptor-strelka/COLO\_829-TGen.15.bcf - varlociraptor-strelka/COLO\_829-TGen.16.bcf - varlociraptor-strelka/COLO\_829-TGen.17.bcf - varlociraptor-strelka/COLO\_829-TGen.18.bcf - varlociraptor-strelka/COLO\_829-TGen.19.bcf - varlociraptor-strelka/COLO\_829-TGen.20.bcf - varlociraptor-strelka/COLO\_829-TGen.21.bcf - varlociraptor-strelka/COLO\_829-TGen.22.bcf - varlociraptor-strelka/COLO\_829-TGen.M.bcf - varlociraptor-strelka/COLO\_829-TGen.X.bcf - varlociraptor-strelka/COLO\_829-TGen.Y.bcf - varlociraptor-strelka/COLO\_829-EBI.1.bcf - varlociraptor-strelka/COLO\_829-EBI.2.bcf - varlociraptor-strelka/COLO\_829-EBI.3.bcf - varlociraptor-strelka/COLO\_829-EBI.4.bcf - varlociraptor-strelka/COLO\_829-EBI.5.bcf - varlociraptor-strelka/COLO\_829-EBI.6.bcf - varlociraptor-strelka/COLO\_829-EBI.7.bcf - varlociraptor-strelka/COLO\_829-EBI.8.bcf - varlociraptor-strelka/COLO\_829-EBI.9.bcf - varlociraptor-strelka/COLO\_829-EBI.10.bcf - varlociraptor-strelka/COLO\_829-EBI.11.bcf - varlociraptor-strelka/COLO\_829-EBI.12.bcf - varlociraptor-strelka/COLO\_829-EBI.13.bcf - varlociraptor-strelka/COLO\_829-EBI.14.bcf - varlociraptor-strelka/COLO\_829-EBI.15.bcf - varlociraptor-strelka/COLO\_829-EBI.16.bcf - varlociraptor-strelka/COLO\_829-EBI.17.bcf - varlociraptor-strelka/COLO\_829-EBI.18.bcf - varlociraptor-strelka/COLO\_829-EBI.19.bcf - varlociraptor-strelka/COLO\_829-EBI.20.bcf - varlociraptor-strelka/COLO\_829-EBI.21.bcf - varlociraptor-strelka/COLO\_829-EBI.22.bcf - varlociraptor-strelka/COLO\_829-EBI.M.bcf - varlociraptor-strelka/COLO\_829-EBI.X.bcf - varlociraptor-strelka/COLO\_829-EBI.Y.bcf - varlociraptor-bpi/COLO\_829-GSC.1.bcf - varlociraptor-bpi/COLO\_829-GSC.2.bcf - varlociraptor-bpi/COLO\_829-GSC.3.bcf - varlociraptor-bpi/COLO\_829-GSC.4.bcf - varlociraptor-bpi/COLO\_829-GSC.5.bcf - varlociraptor-bpi/COLO\_829-GSC.6.bcf - varlociraptor-bpi/COLO\_829-GSC.7.bcf - varlociraptor-bpi/COLO\_829-GSC.8.bcf - varlociraptor-bpi/COLO\_829-GSC.9.bcf - varlociraptor-bpi/COLO\_829-GSC.10.bcf - varlociraptor-bpi/COLO\_829-GSC.11.bcf - varlociraptor-bpi/COLO\_829-GSC.12.bcf - varlociraptor-bpi/COLO\_829-GSC.13.bcf - varlociraptor-bpi/COLO\_829-GSC.14.bcf - varlociraptor-bpi/COLO\_829-GSC.15.bcf - varlociraptor-bpi/COLO\_829-GSC.16.bcf - varlociraptor-bpi/COLO\_829-GSC.17.bcf - varlociraptor-bpi/COLO\_829-GSC.18.bcf - varlociraptor-bpi/COLO\_829-GSC.19.bcf - varlociraptor-bpi/COLO\_829-GSC.20.bcf - varlociraptor-bpi/COLO\_829-GSC.21.bcf - varlociraptor-bpi/COLO\_829-GSC.22.bcf - varlociraptor-bpi/COLO\_829-GSC.M.bcf - varlociraptor-bpi/COLO\_829-GSC.X.bcf - varlociraptor-bpi/COLO\_829-GSC.Y.bcf - varlociraptor-bpi/COLO\_829-Ill.1.bcf - varlociraptor-bpi/COLO\_829-Ill.2.bcf - varlociraptor-bpi/COLO\_829-Ill.3.bcf - varlociraptor-bpi/COLO\_829-Ill.4.bcf - varlociraptor-bpi/COLO\_829-Ill.5.bcf - varlociraptor-bpi/COLO\_829-Ill.6.bcf - varlociraptor-bpi/COLO\_829-Ill.7.bcf - varlociraptor-bpi/COLO\_829-Ill.8.bcf - varlociraptor-bpi/COLO\_829-Ill.9.bcf - varlociraptor-bpi/COLO\_829-Ill.10.bcf - varlociraptor-bpi/COLO\_829-Ill.11.bcf - varlociraptor-bpi/COLO\_829-Ill.12.bcf - varlociraptor-bpi/COLO\_829-Ill.13.bcf - varlociraptor-bpi/COLO\_829-Ill.14.bcf - varlociraptor-bpi/COLO\_829-Ill.15.bcf - varlociraptor-bpi/COLO\_829-Ill.16.bcf - varlociraptor-bpi/COLO\_829-Ill.17.bcf - varlociraptor-bpi/COLO\_829-Ill.18.bcf - varlociraptor-bpi/COLO\_829-Ill.19.bcf - varlociraptor-bpi/COLO\_829-Ill.20.bcf - varlociraptor-bpi/COLO\_829-Ill.21.bcf - varlociraptor-bpi/COLO\_829-Ill.22.bcf - varlociraptor-bpi/COLO\_829-Ill.M.bcf - varlociraptor-bpi/COLO\_829-Ill.X.bcf - varlociraptor-bpi/COLO\_829-Ill.Y.bcf - varlociraptor-bpi/COLO\_829-TGen.1.bcf - varlociraptor-bpi/COLO\_829-TGen.2.bcf - varlociraptor-bpi/COLO\_829-TGen.3.bcf - varlociraptor-bpi/COLO\_829-TGen.4.bcf - varlociraptor-bpi/COLO\_829-TGen.5.bcf - varlociraptor-bpi/COLO\_829-TGen.6.bcf - varlociraptor-bpi/COLO\_829-TGen.7.bcf - varlociraptor-bpi/COLO\_829-TGen.8.bcf - varlociraptor-bpi/COLO\_829-TGen.9.bcf - varlociraptor-bpi/COLO\_829-TGen.10.bcf - varlociraptor-bpi/COLO\_829-TGen.11.bcf - varlociraptor-bpi/COLO\_829-TGen.12.bcf - varlociraptor-bpi/COLO\_829-TGen.13.bcf - varlociraptor-bpi/COLO\_829-TGen.14.bcf - varlociraptor-bpi/COLO\_829-TGen.15.bcf - varlociraptor-bpi/COLO\_829-TGen.16.bcf - varlociraptor-bpi/COLO\_829-TGen.17.bcf - varlociraptor-bpi/COLO\_829-TGen.18.bcf - varlociraptor-bpi/COLO\_829-TGen.19.bcf - varlociraptor-bpi/COLO\_829-TGen.20.bcf - varlociraptor-bpi/COLO\_829-TGen.21.bcf - varlociraptor-bpi/COLO\_829-TGen.22.bcf - varlociraptor-bpi/COLO\_829-TGen.M.bcf - varlociraptor-bpi/COLO\_829-TGen.X.bcf - varlociraptor-bpi/COLO\_829-TGen.Y.bcf - varlociraptor-bpi/COLO\_829-EBI.1.bcf - varlociraptor-bpi/COLO\_829-EBI.2.bcf - varlociraptor-bpi/COLO\_829-EBI.3.bcf - varlociraptor-bpi/COLO\_829-EBI.4.bcf - varlociraptor-bpi/COLO\_829-EBI.5.bcf - varlociraptor-bpi/COLO\_829-EBI.6.bcf - varlociraptor-bpi/COLO\_829-EBI.7.bcf - varlociraptor-bpi/COLO\_829-EBI.8.bcf - varlociraptor-bpi/COLO\_829-EBI.9.bcf - varlociraptor-bpi/COLO\_829-EBI.10.bcf - varlociraptor-bpi/COLO\_829-EBI.11.bcf - varlociraptor-bpi/COLO\_829-EBI.12.bcf - varlociraptor-bpi/COLO\_829-EBI.13.bcf - varlociraptor-bpi/COLO\_829-EBI.14.bcf - varlociraptor-bpi/COLO\_829-EBI.15.bcf - varlociraptor-bpi/COLO\_829-EBI.16.bcf - varlociraptor-bpi/COLO\_829-EBI.17.bcf - varlociraptor-bpi/COLO\_829-EBI.18.bcf - varlociraptor-bpi/COLO\_829-EBI.19.bcf - varlociraptor-bpi/COLO\_829-EBI.20.bcf - varlociraptor-bpi/COLO\_829-EBI.21.bcf - varlociraptor-bpi/COLO\_829-EBI.22.bcf - varlociraptor-bpi/COLO\_829-EBI.M.bcf - varlociraptor-bpi/COLO\_829-EBI.X.bcf - varlociraptor-bpi/COLO\_829-EBI.Y.bcf - varlociraptor-delly/simulated-bwa.1.bcf - varlociraptor-delly/simulated-bwa.2.bcf - varlociraptor-delly/simulated-bwa.3.bcf - varlociraptor-delly/simulated-bwa.4.bcf - varlociraptor-delly/simulated-bwa.5.bcf - varlociraptor-delly/simulated-bwa.6.bcf - varlociraptor-delly/simulated-bwa.7.bcf - varlociraptor-delly/simulated-bwa.8.bcf - varlociraptor-delly/simulated-bwa.9.bcf - varlociraptor-delly/simulated-bwa.10.bcf - varlociraptor-delly/simulated-bwa.11.bcf - varlociraptor-delly/simulated-bwa.12.bcf - varlociraptor-delly/simulated-bwa.13.bcf - varlociraptor-delly/simulated-bwa.14.bcf - varlociraptor-delly/simulated-bwa.15.bcf - varlociraptor-delly/simulated-bwa.16.bcf - varlociraptor-delly/simulated-bwa.17.bcf - varlociraptor-delly/simulated-bwa.18.bcf - varlociraptor-delly/simulated-bwa.19.bcf - varlociraptor-delly/simulated-bwa.20.bcf - varlociraptor-delly/simulated-bwa.21.bcf - varlociraptor-delly/simulated-bwa.22.bcf - varlociraptor-delly/simulated-bwa.M.bcf - varlociraptor-delly/simulated-bwa.X.bcf - varlociraptor-delly/simulated-bwa.Y.bcf - varlociraptor-lancet/simulated-bwa.1.bcf - varlociraptor-lancet/simulated-bwa.2.bcf - varlociraptor-lancet/simulated-bwa.3.bcf - varlociraptor-lancet/simulated-bwa.4.bcf - varlociraptor-lancet/simulated-bwa.5.bcf - varlociraptor-lancet/simulated-bwa.6.bcf - varlociraptor-lancet/simulated-bwa.7.bcf - varlociraptor-lancet/simulated-bwa.8.bcf - varlociraptor-lancet/simulated-bwa.9.bcf - varlociraptor-lancet/simulated-bwa.10.bcf - varlociraptor-lancet/simulated-bwa.11.bcf - varlociraptor-lancet/simulated-bwa.12.bcf - varlociraptor-lancet/simulated-bwa.13.bcf - varlociraptor-lancet/simulated-bwa.14.bcf - varlociraptor-lancet/simulated-bwa.15.bcf - varlociraptor-lancet/simulated-bwa.16.bcf - varlociraptor-lancet/simulated-bwa.17.bcf - varlociraptor-lancet/simulated-bwa.18.bcf - varlociraptor-lancet/simulated-bwa.19.bcf - varlociraptor-lancet/simulated-bwa.20.bcf - varlociraptor-lancet/simulated-bwa.21.bcf - varlociraptor-lancet/simulated-bwa.22.bcf - varlociraptor-lancet/simulated-bwa.M.bcf - varlociraptor-lancet/simulated-bwa.X.bcf - varlociraptor-lancet/simulated-bwa.Y.bcf - varlociraptor-manta/simulated-bwa.1.bcf - varlociraptor-manta/simulated-bwa.2.bcf - varlociraptor-manta/simulated-bwa.3.bcf - varlociraptor-manta/simulated-bwa.4.bcf - varlociraptor-manta/simulated-bwa.5.bcf - varlociraptor-manta/simulated-bwa.6.bcf - varlociraptor-manta/simulated-bwa.7.bcf - varlociraptor-manta/simulated-bwa.8.bcf - varlociraptor-manta/simulated-bwa.9.bcf - varlociraptor-manta/simulated-bwa.10.bcf - varlociraptor-manta/simulated-bwa.11.bcf - varlociraptor-manta/simulated-bwa.12.bcf - varlociraptor-manta/simulated-bwa.13.bcf - varlociraptor-manta/simulated-bwa.14.bcf - varlociraptor-manta/simulated-bwa.15.bcf - varlociraptor-manta/simulated-bwa.16.bcf - varlociraptor-manta/simulated-bwa.17.bcf - varlociraptor-manta/simulated-bwa.18.bcf - varlociraptor-manta/simulated-bwa.19.bcf - varlociraptor-manta/simulated-bwa.20.bcf - varlociraptor-manta/simulated-bwa.21.bcf - varlociraptor-manta/simulated-bwa.22.bcf - varlociraptor-manta/simulated-bwa.M.bcf - varlociraptor-manta/simulated-bwa.X.bcf - varlociraptor-manta/simulated-bwa.Y.bcf - varlociraptor-strelka/simulated-bwa.1.bcf - varlociraptor-strelka/simulated-bwa.2.bcf - varlociraptor-strelka/simulated-bwa.3.bcf - varlociraptor-strelka/simulated-bwa.4.bcf - varlociraptor-strelka/simulated-bwa.5.bcf - varlociraptor-strelka/simulated-bwa.6.bcf - varlociraptor-strelka/simulated-bwa.7.bcf - varlociraptor-strelka/simulated-bwa.8.bcf - varlociraptor-strelka/simulated-bwa.9.bcf - varlociraptor-strelka/simulated-bwa.10.bcf - varlociraptor-strelka/simulated-bwa.11.bcf - varlociraptor-strelka/simulated-bwa.12.bcf - varlociraptor-strelka/simulated-bwa.13.bcf - varlociraptor-strelka/simulated-bwa.14.bcf - varlociraptor-strelka/simulated-bwa.15.bcf - varlociraptor-strelka/simulated-bwa.16.bcf - varlociraptor-strelka/simulated-bwa.17.bcf - varlociraptor-strelka/simulated-bwa.18.bcf - varlociraptor-strelka/simulated-bwa.19.bcf - varlociraptor-strelka/simulated-bwa.20.bcf - varlociraptor-strelka/simulated-bwa.21.bcf - varlociraptor-strelka/simulated-bwa.22.bcf - varlociraptor-strelka/simulated-bwa.M.bcf - varlociraptor-strelka/simulated-bwa.X.bcf - varlociraptor-strelka/simulated-bwa.Y.bcf |  |  | ``` 1 ``` \| ``` bcftools view -Ou {input.calls} {params.chrom_prefix} \| varlociraptor call variants {input.ref} {config[caller][varlociraptor][params]} {params.caller} tumor-normal {input.bams} --purity {params.purity} > {output} 2> {log} ``` |
| delly\_adhoc | 4 | - adhoc-delly/COLO\_829-GSC.all.bcf - adhoc-delly/COLO\_829-Ill.all.bcf - adhoc-delly/COLO\_829-TGen.all.bcf - adhoc-delly/COLO\_829-EBI.all.bcf |  | - delly =0.7.7 - bcftools =1.6 | ``` 1 ``` \| ``` delly filter -m 0 -r 1.0 --samples {input.samples} -o {params.tmp} {input.bcf}; bcftools view -i INFO/SOMATIC -f PASS -Ob {params.tmp} > {output} ``` |
| lancet\_adhoc | 4 | - adhoc-lancet/COLO\_829-GSC.all.bcf - adhoc-lancet/COLO\_829-Ill.all.bcf - adhoc-lancet/COLO\_829-TGen.all.bcf - adhoc-lancet/COLO\_829-EBI.all.bcf |  | - bcftools ==1.5 | ``` 1 2 3 4 5 6 7 8 9 10 11 12 ``` \| ``` __author__ = "Johannes Köster" __copyright__ = "Copyright 2016, Johannes Köster" __email__ = "" __license__ = "MIT" from snakemake.shell import shell shell( "bcftools view {snakemake.params} {snakemake.input[0]} " "-o {snakemake.output[0]}") ``` |
| manta\_adhoc | 4 | - adhoc-manta/COLO\_829-GSC.all.bcf - adhoc-manta/COLO\_829-Ill.all.bcf - adhoc-manta/COLO\_829-TGen.all.bcf - adhoc-manta/COLO\_829-EBI.all.bcf |  | - bcftools ==1.5 | ``` 1 2 3 4 5 6 7 8 9 10 11 12 ``` \| ``` __author__ = "Johannes Köster" __copyright__ = "Copyright 2016, Johannes Köster" __email__ = "" __license__ = "MIT" from snakemake.shell import shell shell( "bcftools view {snakemake.params} {snakemake.input[0]} " "-o {snakemake.output[0]}") ``` |
| strelka\_adhoc | 4 | - adhoc-strelka/COLO\_829-GSC.all.bcf - adhoc-strelka/COLO\_829-Ill.all.bcf - adhoc-strelka/COLO\_829-TGen.all.bcf - adhoc-strelka/COLO\_829-EBI.all.bcf |  | - bcftools ==1.5 | ``` 1 2 3 4 5 6 7 8 9 10 11 12 ``` \| ``` __author__ = "Johannes Köster" __copyright__ = "Copyright 2016, Johannes Köster" __email__ = "" __license__ = "MIT" from snakemake.shell import shell shell( "bcftools view {snakemake.params} {snakemake.input[0]} " "-o {snakemake.output[0]}") ``` |
| bpi\_adhoc | 4 | - adhoc-bpi/COLO\_829-GSC.all.bcf - adhoc-bpi/COLO\_829-Ill.all.bcf - adhoc-bpi/COLO\_829-TGen.all.bcf - adhoc-bpi/COLO\_829-EBI.all.bcf |  | - bcftools ==1.5 | ``` 1 2 3 4 5 6 7 8 9 10 11 12 ``` \| ``` __author__ = "Johannes Köster" __copyright__ = "Copyright 2016, Johannes Köster" __email__ = "" __license__ = "MIT" from snakemake.shell import shell shell( "bcftools view {snakemake.params} {snakemake.input[0]} " "-o {snakemake.output[0]}") ``` |
| delly\_concat | 4 | - delly/COLO\_829-GSC.all.bcf - delly/COLO\_829-Ill.all.bcf - delly/COLO\_829-TGen.all.bcf - delly/COLO\_829-EBI.all.bcf |  | - bcftools ==1.5 | ``` 1 2 3 4 5 6 7 8 9 10 11 12 ``` \| ``` __author__ = "Johannes Köster" __copyright__ = "Copyright 2016, Johannes Köster" __email__ = "" __license__ = "MIT" from snakemake.shell import shell shell( "bcftools concat {snakemake.params} -o {snakemake.output[0]} " "{snakemake.input}") ``` |
| index\_bcf | 20 | - delly/COLO\_829-GSC.all.bcf.csi - delly/COLO\_829-Ill.all.bcf.csi - delly/COLO\_829-TGen.all.bcf.csi - delly/COLO\_829-EBI.all.bcf.csi - default-lancet/COLO\_829-GSC.all.bcf.csi - default-lancet/COLO\_829-Ill.all.bcf.csi - default-lancet/COLO\_829-TGen.all.bcf.csi - default-lancet/COLO\_829-EBI.all.bcf.csi - manta/COLO\_829-GSC.all.bcf.csi - manta/COLO\_829-Ill.all.bcf.csi - manta/COLO\_829-TGen.all.bcf.csi - manta/COLO\_829-EBI.all.bcf.csi - default-strelka/COLO\_829-GSC.all.bcf.csi - default-strelka/COLO\_829-Ill.all.bcf.csi - default-strelka/COLO\_829-TGen.all.bcf.csi - default-strelka/COLO\_829-EBI.all.bcf.csi - bpi/COLO\_829-GSC.all.bcf.csi - bpi/COLO\_829-Ill.all.bcf.csi - bpi/COLO\_829-TGen.all.bcf.csi - bpi/COLO\_829-EBI.all.bcf.csi |  | - bcftools =1.6 - samtools =1.6 | ``` 1 ``` \| ``` bcftools index {input} ``` |
| mark\_duplicates | 8 | - mapped-bwa/COLO\_829-GSC.tumor.hg38.sorted.bam - mapped-bwa/COLO\_829-GSC.tumor.hg38.markdup.metrics.txt - mapped-bwa/COLO\_829-GSC.normal.hg38.sorted.bam - mapped-bwa/COLO\_829-GSC.normal.hg38.markdup.metrics.txt - mapped-bwa/COLO\_829-Ill.tumor.hg38.sorted.bam - mapped-bwa/COLO\_829-Ill.tumor.hg38.markdup.metrics.txt - mapped-bwa/COLO\_829-Ill.normal.hg38.sorted.bam - mapped-bwa/COLO\_829-Ill.normal.hg38.markdup.metrics.txt - mapped-bwa/COLO\_829-TGen.tumor.hg38.sorted.bam - mapped-bwa/COLO\_829-TGen.tumor.hg38.markdup.metrics.txt - mapped-bwa/COLO\_829-TGen.normal.hg38.sorted.bam - mapped-bwa/COLO\_829-TGen.normal.hg38.markdup.metrics.txt - mapped-bwa/COLO\_829-EBI.tumor.hg38.sorted.bam - mapped-bwa/COLO\_829-EBI.tumor.hg38.markdup.metrics.txt - mapped-bwa/COLO\_829-EBI.normal.hg38.sorted.bam - mapped-bwa/COLO\_829-EBI.normal.hg38.markdup.metrics.txt |  | - picard ==2.9.2 | ``` 1 2 3 4 5 6 7 8 9 10 11 12 ``` \| ``` __author__ = "Johannes Köster" __copyright__ = "Copyright 2016, Johannes Köster" __email__ = "" __license__ = "MIT" from snakemake.shell import shell shell("picard MarkDuplicates {snakemake.params} INPUT={snakemake.input} " "OUTPUT={snakemake.output.bam} METRICS_FILE={snakemake.output.metrics} " "&> {snakemake.log}") ``` |
| samtools\_index | 8 | - mapped-bwa/COLO\_829-GSC.tumor.hg38.sorted.bam.bai - mapped-bwa/COLO\_829-GSC.normal.hg38.sorted.bam.bai - mapped-bwa/COLO\_829-Ill.tumor.hg38.sorted.bam.bai - mapped-bwa/COLO\_829-Ill.normal.hg38.sorted.bam.bai - mapped-bwa/COLO\_829-TGen.tumor.hg38.sorted.bam.bai - mapped-bwa/COLO\_829-TGen.normal.hg38.sorted.bam.bai - mapped-bwa/COLO\_829-EBI.tumor.hg38.sorted.bam.bai - mapped-bwa/COLO\_829-EBI.normal.hg38.sorted.bam.bai |  | - samtools ==1.6 | ``` 1 2 3 4 5 6 7 8 9 10 ``` \| ``` __author__ = "Johannes Köster" __copyright__ = "Copyright 2016, Johannes Köster" __email__ = "" __license__ = "MIT" from snakemake.shell import shell shell("samtools index {snakemake.params} {snakemake.input[0]} {snakemake.output[0]}") ``` |
| merge\_lancet | 4 | - default-lancet/COLO\_829-GSC.all.bcf - default-lancet/COLO\_829-Ill.all.bcf - default-lancet/COLO\_829-TGen.all.bcf - default-lancet/COLO\_829-EBI.all.bcf |  | - bcftools =1.6 - samtools =1.6 | ``` 1 ``` \| ``` bcftools concat -Ob {input} > {output} ``` |
| manta\_raw | 4 | - manta/COLO\_829-GSC.all.bcf - manta/COLO\_829-Ill.all.bcf - manta/COLO\_829-TGen.all.bcf - manta/COLO\_829-EBI.all.bcf |  | - bcftools ==1.5 | ``` 1 2 3 4 5 6 7 8 9 10 11 12 ``` \| ``` __author__ = "Johannes Köster" __copyright__ = "Copyright 2016, Johannes Köster" __email__ = "" __license__ = "MIT" from snakemake.shell import shell shell( "bcftools view {snakemake.params} {snakemake.input[0]} " "-o {snakemake.output[0]}") ``` |
| strelka\_default | 4 | - default-strelka/COLO\_829-GSC.all.bcf - default-strelka/COLO\_829-Ill.all.bcf - default-strelka/COLO\_829-TGen.all.bcf - default-strelka/COLO\_829-EBI.all.bcf |  | - bcftools ==1.6 | ``` 1 2 3 4 5 6 7 8 9 10 11 12 ``` \| ``` __author__ = "Johannes Köster" __copyright__ = "Copyright 2016, Johannes Köster" __email__ = "" __license__ = "MIT" from snakemake.shell import shell shell( "bcftools concat {snakemake.params} -o {snakemake.output[0]} " "{snakemake.input.calls}") ``` |
| bpi\_convert | 4 | - bpi/COLO\_829-GSC.all.bcf - bpi/COLO\_829-Ill.all.bcf - bpi/COLO\_829-TGen.all.bcf - bpi/COLO\_829-EBI.all.bcf |  | - bcftools ==1.5 | ``` 1 2 3 4 5 6 7 8 9 10 11 12 ``` \| ``` __author__ = "Johannes Köster" __copyright__ = "Copyright 2016, Johannes Köster" __email__ = "" __license__ = "MIT" from snakemake.shell import shell shell( "bcftools view {snakemake.params} {snakemake.input[0]} " "-o {snakemake.output[0]}") ``` |
| manta\_default | 4 | - default-manta/COLO\_829-GSC.all.bcf - default-manta/COLO\_829-Ill.all.bcf - default-manta/COLO\_829-TGen.all.bcf - default-manta/COLO\_829-EBI.all.bcf |  | - bcftools ==1.5 | ``` 1 2 3 4 5 6 7 8 9 10 11 12 ``` \| ``` __author__ = "Johannes Köster" __copyright__ = "Copyright 2016, Johannes Köster" __email__ = "" __license__ = "MIT" from snakemake.shell import shell shell( "bcftools view {snakemake.params} {snakemake.input[0]} " "-o {snakemake.output[0]}") ``` |
| delly | 8 | - delly/COLO\_829-GSC.DEL.bcf - delly/COLO\_829-GSC.INS.bcf - delly/COLO\_829-Ill.DEL.bcf - delly/COLO\_829-Ill.INS.bcf - delly/COLO\_829-TGen.DEL.bcf - delly/COLO\_829-TGen.INS.bcf - delly/COLO\_829-EBI.DEL.bcf - delly/COLO\_829-EBI.INS.bcf |  | - delly ==0.7.7 | ``` 1 2 3 4 5 6 7 8 9 10 11 12 13 14 15 16 17 18 19 20 21 22 ``` \| ``` __author__ = "Johannes Köster" __copyright__ = "Copyright 2016, Johannes Köster" __email__ = "" __license__ = "MIT" from snakemake.shell import shell try: exclude = "-x " + snakemake.input.exclude except AttributeError: exclude = "" extra = snakemake.params.get("extra", "") log = snakemake.log_fmt_shell(stdout=True, stderr=True) shell( "OMP_NUM_THREADS={snakemake.threads} delly call {extra} " "{exclude} -t {snakemake.params.vartype} -g {snakemake.input.ref} " "-o {snakemake.output[0]} {snakemake.input.samples} {log}") ``` |
| samtools\_sort | 8 | - mapped-bwa/COLO\_829-GSC.tumor.hg38.sorted.pre.bam - mapped-bwa/COLO\_829-GSC.normal.hg38.sorted.pre.bam - mapped-bwa/COLO\_829-Ill.tumor.hg38.sorted.pre.bam - mapped-bwa/COLO\_829-Ill.normal.hg38.sorted.pre.bam - mapped-bwa/COLO\_829-TGen.tumor.hg38.sorted.pre.bam - mapped-bwa/COLO\_829-TGen.normal.hg38.sorted.pre.bam - mapped-bwa/COLO\_829-EBI.tumor.hg38.sorted.pre.bam - mapped-bwa/COLO\_829-EBI.normal.hg38.sorted.pre.bam |  | - samtools ==1.6 | ``` 1 2 3 4 5 6 7 8 9 10 11 12 13 14 15 ``` \| ``` __author__ = "Johannes Köster" __copyright__ = "Copyright 2016, Johannes Köster" __email__ = "" __license__ = "MIT" import os from snakemake.shell import shell prefix = os.path.splitext(snakemake.output[0])[0] shell( "samtools sort {snakemake.params} -@ {snakemake.threads} -o {snakemake.output[0]} " "-T {prefix} {snakemake.input[0]}") ``` |
| fix\_lancet | 100 | - lancet/COLO\_829-GSC/chr1.fixed.vcf - lancet/COLO\_829-GSC/chr2.fixed.vcf - lancet/COLO\_829-GSC/chr3.fixed.vcf - lancet/COLO\_829-GSC/chr4.fixed.vcf - lancet/COLO\_829-GSC/chr5.fixed.vcf - lancet/COLO\_829-GSC/chr6.fixed.vcf - lancet/COLO\_829-GSC/chr7.fixed.vcf - lancet/COLO\_829-GSC/chr8.fixed.vcf - lancet/COLO\_829-GSC/chr9.fixed.vcf - lancet/COLO\_829-GSC/chr10.fixed.vcf - lancet/COLO\_829-GSC/chr11.fixed.vcf - lancet/COLO\_829-GSC/chr12.fixed.vcf - lancet/COLO\_829-GSC/chr13.fixed.vcf - lancet/COLO\_829-GSC/chr14.fixed.vcf - lancet/COLO\_829-GSC/chr15.fixed.vcf - lancet/COLO\_829-GSC/chr16.fixed.vcf - lancet/COLO\_829-GSC/chr17.fixed.vcf - lancet/COLO\_829-GSC/chr18.fixed.vcf - lancet/COLO\_829-GSC/chr19.fixed.vcf - lancet/COLO\_829-GSC/chr20.fixed.vcf - lancet/COLO\_829-GSC/chr21.fixed.vcf - lancet/COLO\_829-GSC/chr22.fixed.vcf - lancet/COLO\_829-GSC/chrM.fixed.vcf - lancet/COLO\_829-GSC/chrX.fixed.vcf - lancet/COLO\_829-GSC/chrY.fixed.vcf - lancet/COLO\_829-Ill/chr1.fixed.vcf - lancet/COLO\_829-Ill/chr2.fixed.vcf - lancet/COLO\_829-Ill/chr3.fixed.vcf - lancet/COLO\_829-Ill/chr4.fixed.vcf - lancet/COLO\_829-Ill/chr5.fixed.vcf - lancet/COLO\_829-Ill/chr6.fixed.vcf - lancet/COLO\_829-Ill/chr7.fixed.vcf - lancet/COLO\_829-Ill/chr8.fixed.vcf - lancet/COLO\_829-Ill/chr9.fixed.vcf - lancet/COLO\_829-Ill/chr10.fixed.vcf - lancet/COLO\_829-Ill/chr11.fixed.vcf - lancet/COLO\_829-Ill/chr12.fixed.vcf - lancet/COLO\_829-Ill/chr13.fixed.vcf - lancet/COLO\_829-Ill/chr14.fixed.vcf - lancet/COLO\_829-Ill/chr15.fixed.vcf - lancet/COLO\_829-Ill/chr16.fixed.vcf - lancet/COLO\_829-Ill/chr17.fixed.vcf - lancet/COLO\_829-Ill/chr18.fixed.vcf - lancet/COLO\_829-Ill/chr19.fixed.vcf - lancet/COLO\_829-Ill/chr20.fixed.vcf - lancet/COLO\_829-Ill/chr21.fixed.vcf - lancet/COLO\_829-Ill/chr22.fixed.vcf - lancet/COLO\_829-Ill/chrM.fixed.vcf - lancet/COLO\_829-Ill/chrX.fixed.vcf - lancet/COLO\_829-Ill/chrY.fixed.vcf - lancet/COLO\_829-TGen/chr1.fixed.vcf - lancet/COLO\_829-TGen/chr2.fixed.vcf - lancet/COLO\_829-TGen/chr3.fixed.vcf - lancet/COLO\_829-TGen/chr4.fixed.vcf - lancet/COLO\_829-TGen/chr5.fixed.vcf - lancet/COLO\_829-TGen/chr6.fixed.vcf - lancet/COLO\_829-TGen/chr7.fixed.vcf - lancet/COLO\_829-TGen/chr8.fixed.vcf - lancet/COLO\_829-TGen/chr9.fixed.vcf - lancet/COLO\_829-TGen/chr10.fixed.vcf - lancet/COLO\_829-TGen/chr11.fixed.vcf - lancet/COLO\_829-TGen/chr12.fixed.vcf - lancet/COLO\_829-TGen/chr13.fixed.vcf - lancet/COLO\_829-TGen/chr14.fixed.vcf - lancet/COLO\_829-TGen/chr15.fixed.vcf - lancet/COLO\_829-TGen/chr16.fixed.vcf - lancet/COLO\_829-TGen/chr17.fixed.vcf - lancet/COLO\_829-TGen/chr18.fixed.vcf - lancet/COLO\_829-TGen/chr19.fixed.vcf - lancet/COLO\_829-TGen/chr20.fixed.vcf - lancet/COLO\_829-TGen/chr21.fixed.vcf - lancet/COLO\_829-TGen/chr22.fixed.vcf - lancet/COLO\_829-TGen/chrM.fixed.vcf - lancet/COLO\_829-TGen/chrX.fixed.vcf - lancet/COLO\_829-TGen/chrY.fixed.vcf - lancet/COLO\_829-EBI/chr1.fixed.vcf - lancet/COLO\_829-EBI/chr2.fixed.vcf - lancet/COLO\_829-EBI/chr3.fixed.vcf - lancet/COLO\_829-EBI/chr4.fixed.vcf - lancet/COLO\_829-EBI/chr5.fixed.vcf - lancet/COLO\_829-EBI/chr6.fixed.vcf - lancet/COLO\_829-EBI/chr7.fixed.vcf - lancet/COLO\_829-EBI/chr8.fixed.vcf - lancet/COLO\_829-EBI/chr9.fixed.vcf - lancet/COLO\_829-EBI/chr10.fixed.vcf - lancet/COLO\_829-EBI/chr11.fixed.vcf - lancet/COLO\_829-EBI/chr12.fixed.vcf - lancet/COLO\_829-EBI/chr13.fixed.vcf - lancet/COLO\_829-EBI/chr14.fixed.vcf - lancet/COLO\_829-EBI/chr15.fixed.vcf - lancet/COLO\_829-EBI/chr16.fixed.vcf - lancet/COLO\_829-EBI/chr17.fixed.vcf - lancet/COLO\_829-EBI/chr18.fixed.vcf - lancet/COLO\_829-EBI/chr19.fixed.vcf - lancet/COLO\_829-EBI/chr20.fixed.vcf - lancet/COLO\_829-EBI/chr21.fixed.vcf - lancet/COLO\_829-EBI/chr22.fixed.vcf - lancet/COLO\_829-EBI/chrM.fixed.vcf - lancet/COLO\_829-EBI/chrX.fixed.vcf - lancet/COLO\_829-EBI/chrY.fixed.vcf |  | - bcftools =1.6 - samtools =1.6 | ``` 1 ``` \| ``` sed -r 's/MS\=[0-9]+[ACGT]+/MS/g' {input.vcf} \| bcftools annotate -o {output} -h {input.header} - ``` |
| manta | 4 | - manta/COLO\_829-GSC/results/variants/candidateSV.vcf.gz - manta/COLO\_829-GSC/results/variants/somaticSV.vcf.gz - manta/COLO\_829-GSC/results/variants/candidateSmallIndels.vcf.gz - manta/COLO\_829-Ill/results/variants/candidateSV.vcf.gz - manta/COLO\_829-Ill/results/variants/somaticSV.vcf.gz - manta/COLO\_829-Ill/results/variants/candidateSmallIndels.vcf.gz - manta/COLO\_829-TGen/results/variants/candidateSV.vcf.gz - manta/COLO\_829-TGen/results/variants/somaticSV.vcf.gz - manta/COLO\_829-TGen/results/variants/candidateSmallIndels.vcf.gz - manta/COLO\_829-EBI/results/variants/candidateSV.vcf.gz - manta/COLO\_829-EBI/results/variants/somaticSV.vcf.gz - manta/COLO\_829-EBI/results/variants/candidateSmallIndels.vcf.gz |  | - manta =1.3.0 | ``` 1 ``` \| ``` rm -rf {params.dir}; (configManta.py {params.extra} --tumorBam {input.samples[0]} --normalBam {input.samples[1]} --referenceFasta {input.ref} --runDir {params.dir}; {params.dir}/runWorkflow.py -m local -j {threads}) > {log} 2>&1 ``` |
| strelka | 4 | - strelka/COLO\_829-GSC/results/variants/somatic.snvs.vcf.gz - strelka/COLO\_829-GSC/results/variants/somatic.indels.vcf.gz - strelka/COLO\_829-Ill/results/variants/somatic.snvs.vcf.gz - strelka/COLO\_829-Ill/results/variants/somatic.indels.vcf.gz - strelka/COLO\_829-TGen/results/variants/somatic.snvs.vcf.gz - strelka/COLO\_829-TGen/results/variants/somatic.indels.vcf.gz - strelka/COLO\_829-EBI/results/variants/somatic.snvs.vcf.gz - strelka/COLO\_829-EBI/results/variants/somatic.indels.vcf.gz |  | - strelka =2.8.4 | ``` 1 ``` \| ``` rm -rf {params.dir}; (configureStrelkaSomaticWorkflow.py {params.extra} --tumorBam {input.samples[0]} --normalBam {input.samples[1]} --referenceFasta {input.ref} --runDir {params.dir} --indelCandidates {input.manta}; {params.dir}/runWorkflow.py -m local -j {threads}) > {log} 2>&1 ``` |
| bpi | 4 | - bpi/COLO\_829-GSC.all.vcf - bpi/COLO\_829-Ill.all.vcf - bpi/COLO\_829-TGen.all.vcf - bpi/COLO\_829-EBI.all.vcf |  | - break-point-inspector =1.5 | ``` 1 ``` \| ``` (break-point-inspector -vcf {input.manta} -ref {input.samples[1]} -tumor {input.samples[0]} -output_vcf {output}) > {log} 2>&1 ``` |
| bwa | 8 | - mapped-bwa/COLO\_829-GSC.tumor.hg38.bam - mapped-bwa/COLO\_829-GSC.normal.hg38.bam - mapped-bwa/COLO\_829-Ill.tumor.hg38.bam - mapped-bwa/COLO\_829-Ill.normal.hg38.bam - mapped-bwa/COLO\_829-TGen.tumor.hg38.bam - mapped-bwa/COLO\_829-TGen.normal.hg38.bam - mapped-bwa/COLO\_829-EBI.tumor.hg38.bam - mapped-bwa/COLO\_829-EBI.normal.hg38.bam |  | - python - samtools =1.6 - numpy - scikit-learn - pandas - setuptools | ``` 1 ``` \| ``` (resources/bwa mem -t {threads} {params.extra} {params.index} {input.sample} \| samtools view -Sb - > {output}) 2> {log} ``` |
| lancet | 75 | - lancet/COLO\_829-GSC/chr1.vcf - lancet/COLO\_829-GSC/chr2.vcf - lancet/COLO\_829-GSC/chr3.vcf - lancet/COLO\_829-GSC/chr4.vcf - lancet/COLO\_829-GSC/chr5.vcf - lancet/COLO\_829-GSC/chr6.vcf - lancet/COLO\_829-GSC/chr7.vcf - lancet/COLO\_829-GSC/chr8.vcf - lancet/COLO\_829-GSC/chr9.vcf - lancet/COLO\_829-GSC/chr10.vcf - lancet/COLO\_829-GSC/chr11.vcf - lancet/COLO\_829-GSC/chr12.vcf - lancet/COLO\_829-GSC/chr13.vcf - lancet/COLO\_829-GSC/chr14.vcf - lancet/COLO\_829-GSC/chr15.vcf - lancet/COLO\_829-GSC/chr16.vcf - lancet/COLO\_829-GSC/chr17.vcf - lancet/COLO\_829-GSC/chr18.vcf - lancet/COLO\_829-GSC/chr19.vcf - lancet/COLO\_829-GSC/chr20.vcf - lancet/COLO\_829-GSC/chr21.vcf - lancet/COLO\_829-GSC/chr22.vcf - lancet/COLO\_829-GSC/chrM.vcf - lancet/COLO\_829-GSC/chrX.vcf - lancet/COLO\_829-GSC/chrY.vcf - lancet/COLO\_829-Ill/chr1.vcf - lancet/COLO\_829-Ill/chr2.vcf - lancet/COLO\_829-Ill/chr3.vcf - lancet/COLO\_829-Ill/chr4.vcf - lancet/COLO\_829-Ill/chr5.vcf - lancet/COLO\_829-Ill/chr6.vcf - lancet/COLO\_829-Ill/chr7.vcf - lancet/COLO\_829-Ill/chr8.vcf - lancet/COLO\_829-Ill/chr9.vcf - lancet/COLO\_829-Ill/chr10.vcf - lancet/COLO\_829-Ill/chr11.vcf - lancet/COLO\_829-Ill/chr12.vcf - lancet/COLO\_829-Ill/chr13.vcf - lancet/COLO\_829-Ill/chr14.vcf - lancet/COLO\_829-Ill/chr15.vcf - lancet/COLO\_829-Ill/chr16.vcf - lancet/COLO\_829-Ill/chr17.vcf - lancet/COLO\_829-Ill/chr18.vcf - lancet/COLO\_829-Ill/chr19.vcf - lancet/COLO\_829-Ill/chr20.vcf - lancet/COLO\_829-Ill/chr21.vcf - lancet/COLO\_829-Ill/chr22.vcf - lancet/COLO\_829-Ill/chrM.vcf - lancet/COLO\_829-Ill/chrX.vcf - lancet/COLO\_829-Ill/chrY.vcf - lancet/COLO\_829-TGen/chr1.vcf - lancet/COLO\_829-TGen/chr2.vcf - lancet/COLO\_829-TGen/chr3.vcf - lancet/COLO\_829-TGen/chr4.vcf - lancet/COLO\_829-TGen/chr5.vcf - lancet/COLO\_829-TGen/chr6.vcf - lancet/COLO\_829-TGen/chr7.vcf - lancet/COLO\_829-TGen/chr8.vcf - lancet/COLO\_829-TGen/chr9.vcf - lancet/COLO\_829-TGen/chr10.vcf - lancet/COLO\_829-TGen/chr11.vcf - lancet/COLO\_829-TGen/chr12.vcf - lancet/COLO\_829-TGen/chr13.vcf - lancet/COLO\_829-TGen/chr14.vcf - lancet/COLO\_829-TGen/chr15.vcf - lancet/COLO\_829-TGen/chr16.vcf - lancet/COLO\_829-TGen/chr17.vcf - lancet/COLO\_829-TGen/chr18.vcf - lancet/COLO\_829-TGen/chr19.vcf - lancet/COLO\_829-TGen/chr20.vcf - lancet/COLO\_829-TGen/chr21.vcf - lancet/COLO\_829-TGen/chr22.vcf - lancet/COLO\_829-TGen/chrM.vcf - lancet/COLO\_829-TGen/chrX.vcf - lancet/COLO\_829-TGen/chrY.vcf |  | - bamtools =2.3.0 | ``` 1 ``` \| ``` LD_LIBRARY_PATH=$CONDA_PREFIX/lib resources/lancet --tumor {input.bams[0]} --normal {input.bams[1]} --ref {input.ref} --reg {params.region} --num-threads {threads} {params.extra} > {output} 2> {log} ``` |
| lancet | 25 | - lancet/COLO\_829-EBI/chr1.vcf - lancet/COLO\_829-EBI/chr2.vcf - lancet/COLO\_829-EBI/chr3.vcf - lancet/COLO\_829-EBI/chr4.vcf - lancet/COLO\_829-EBI/chr5.vcf - lancet/COLO\_829-EBI/chr6.vcf - lancet/COLO\_829-EBI/chr7.vcf - lancet/COLO\_829-EBI/chr8.vcf - lancet/COLO\_829-EBI/chr9.vcf - lancet/COLO\_829-EBI/chr10.vcf - lancet/COLO\_829-EBI/chr11.vcf - lancet/COLO\_829-EBI/chr12.vcf - lancet/COLO\_829-EBI/chr13.vcf - lancet/COLO\_829-EBI/chr14.vcf - lancet/COLO\_829-EBI/chr15.vcf - lancet/COLO\_829-EBI/chr16.vcf - lancet/COLO\_829-EBI/chr17.vcf - lancet/COLO\_829-EBI/chr18.vcf - lancet/COLO\_829-EBI/chr19.vcf - lancet/COLO\_829-EBI/chr20.vcf - lancet/COLO\_829-EBI/chr21.vcf - lancet/COLO\_829-EBI/chr22.vcf - lancet/COLO\_829-EBI/chrM.vcf - lancet/COLO\_829-EBI/chrX.vcf - lancet/COLO\_829-EBI/chrY.vcf |  |  | ``` 1 ``` \| ``` LD_LIBRARY_PATH=$CONDA_PREFIX/lib resources/lancet --tumor {input.bams[0]} --normal {input.bams[1]} --ref {input.ref} --reg {params.region} --num-threads {threads} {params.extra} > {output} 2> {log} ``` |
| bam2fq | 4 | - reads/COLO\_829-Ill.tumor.1.fastq.gz - reads/COLO\_829-Ill.tumor.2.fastq.gz - reads/COLO\_829-Ill.tumor.fastq.gz - reads/COLO\_829-Ill.normal.1.fastq.gz - reads/COLO\_829-Ill.normal.2.fastq.gz - reads/COLO\_829-Ill.normal.fastq.gz - reads/COLO\_829-TGen.tumor.1.fastq.gz - reads/COLO\_829-TGen.tumor.2.fastq.gz - reads/COLO\_829-TGen.tumor.fastq.gz - reads/COLO\_829-TGen.normal.1.fastq.gz - reads/COLO\_829-TGen.normal.2.fastq.gz - reads/COLO\_829-TGen.normal.fastq.gz |  | - bcftools =1.6 - samtools =1.6 | ``` 1 ``` \| ``` samtools bam2fq {input} -1 {output.m1} -2 {output.m2} -0 {output.mixed} ``` |
| prepare\_bam | 8 | - reads/COLO\_829-GSC.tumor.namesorted.bam - reads/COLO\_829-GSC.normal.namesorted.bam - reads/COLO\_829-Ill.tumor.namesorted.bam - reads/COLO\_829-Ill.normal.namesorted.bam - reads/COLO\_829-TGen.tumor.namesorted.bam - reads/COLO\_829-TGen.normal.namesorted.bam - reads/COLO\_829-EBI.tumor.namesorted.bam - reads/COLO\_829-EBI.normal.namesorted.bam |  | - samtools ==1.6 | ``` 1 2 3 4 5 6 7 8 9 10 11 12 13 14 15 ``` \| ``` __author__ = "Johannes Köster" __copyright__ = "Copyright 2016, Johannes Köster" __email__ = "" __license__ = "MIT" import os from snakemake.shell import shell prefix = os.path.splitext(snakemake.output[0])[0] shell( "samtools sort {snakemake.params} -@ {snakemake.threads} -o {snakemake.output[0]} " "-T {prefix} {snakemake.input[0]}") ``` |
